# Chromosome silencing *in vitro* reveals trisomy 21 causes cell-autonomous deficits in angiogenesis and early dysregulation in Notch signaling

**DOI:** 10.1101/2022.06.01.494361

**Authors:** Jennifer E. Moon, Jeanne B. Lawrence

**Affiliations:** Department of Neurology, University of Massachusetts Medical School, Worcester, MA, 01655, USA; Department of Pediatrics, University of Massachusetts Medical School, Worcester, MA, 01655, USA

## Abstract

Despite the prevalence and clinical importance of Down syndrome, little is known as to the specific cell pathologies that underlie this multi-system disorder. To understand which cell types and pathways are more directly impacted by trisomy 21, we used an inducible-*XIST* system to silence the extra chromosome 21 in a panel of patient-derived iPSCs. Transcriptomic analysis showed significant dysregulation of Notch signaling occurring as early as pluripotent stem cells, potentially impacting programming of multiple cell-types. Unbiased analysis from iPSCs revealed prominent dysregulation in two major cell type processes: neurogenesis and angiogenesis. Angiogenesis is important for many systems impacted in Down syndrome but has been understudied; therefore, we focused on investigating whether trisomy 21 impacts endothelial cells. An *in vitro* assay for microvasculature formation used in a tightly controlled system reveals a novel cellular pathology involving delays in angiogenic response during tube formation. Results demonstrate that this is a cell-autonomous effect of trisomy 21, and transcriptomic analysis of differentiated endothelial cells shows deficits in known angiogenesis regulators. This study reveals a major unknown cell pathology caused by trisomy 21 and highlights the importance of endothelial cell function for Down syndrome comorbidities, with wide reaching implications for development and disease progression.

## INTRODUCTION

Down syndrome (DS), caused by trisomy of chromosome 21 (chr21) occurs in 1 in every 700 live births in the US and conceptions have a high rate of spontaneous loss. Despite the clinical and societal impact of DS, the cellular pathologies that underlie this important syndrome remain poorly understood. Trisomy 21 (T21) consistently leads to mild to moderate intellectual disability in children which can progress in severity in adults. T21 sharply increases risks for other medical conditions, particularly congenital heart defects, early-onset Alzheimer’s disease (AD) and acute megakaryoblastic leukemia (Antonarakis et al., 2020; Startin et al., 2020). Despite higher risk of leukemia, T21 results in reduced rates of solid tumors (Hasle et al., 2016). Unlike monogenic disorders, understanding DS biology is a particular challenge given it involves an extra copy of ∼300 genes on chr21 (∼1% of human genes). In fact, with the exception of the well-established hematopoietic cell pathologies, the specific tissues and cell pathologies that underlie human DS phenotypes remain unclear. Research on mice carrying orthologous segments of chr21 have provided some insights, but questions remain as to how well pathologies in these mice reflect human DS. Studies of patient samples, including post mortem tissues or primary cells, are important but are often limited by small sample sizes, general variation between people, and sample preparations. Importantly, such studies--in mice and humans—only examine the complex outcomes of T21 far downstream of the initial cellular changes that lead to broad impacts on system development or function. Thus, approaches are needed that can illuminate not only how, but when, T21 impacts the development or function of many different systems. Hence, we sought to identify the earliest changes in transcriptome-wide expression and cell development caused by overexpression of chr21 genes, taking an unbiased approach beginning from pluripotent stem cells rather than a specific cell-type. We sought to do so using an experimental strategy designed to minimize sources of variation other than T21 overexpression. Despite variability in DS, consistent DS phenotypes are indicative of a core impact of chr21 dosage on a common set of pathways and cell types.

Our lab developed a novel strategy to study trisomy 21 by translating the unique biology of *XIST* RNA, which coats one female X chromosome in *cis (Brown et al., 1992; Clemson et al., 1996)*, epigenetic modifications which stably silence the chromosome (Creamer and Lawrence, 2017; Loda and Heard, 2019; Sahakyan et al., 2018). We targeted an *XIST* transgene into one chr21 in DS patient-derived induced pluripotent stem cells (iPSCs), and demonstrated robust gene silencing across one chr21 in *cis* (Figure 1A) (Jiang et al., 2013). By making the system inducible (with doxycycline), this allows tightly-controlled comparison of the same cell populations with and without chr21 over-expression. This circumvents inter-cell line variation that can confound identification of trisomy-specific differences, especially for subtle or kinetic differences in cell differentiation or function (Hussein et al., 2013; Liang and Zhang, 2013; Soldner and Jaenisch, 2012). Even isogenic iPSC of hESC lines, during subcloning, passaging, or freeze-thaw, evolve epigenetic drift, as well as occasional chromosomal/genetic changes (Hall et al., 2008; Laurent et al., 2011; Mayshar et al., 2010). The inducible manipulation of chr21 can avoid inter-line sources of variation.

**Figure 1:**
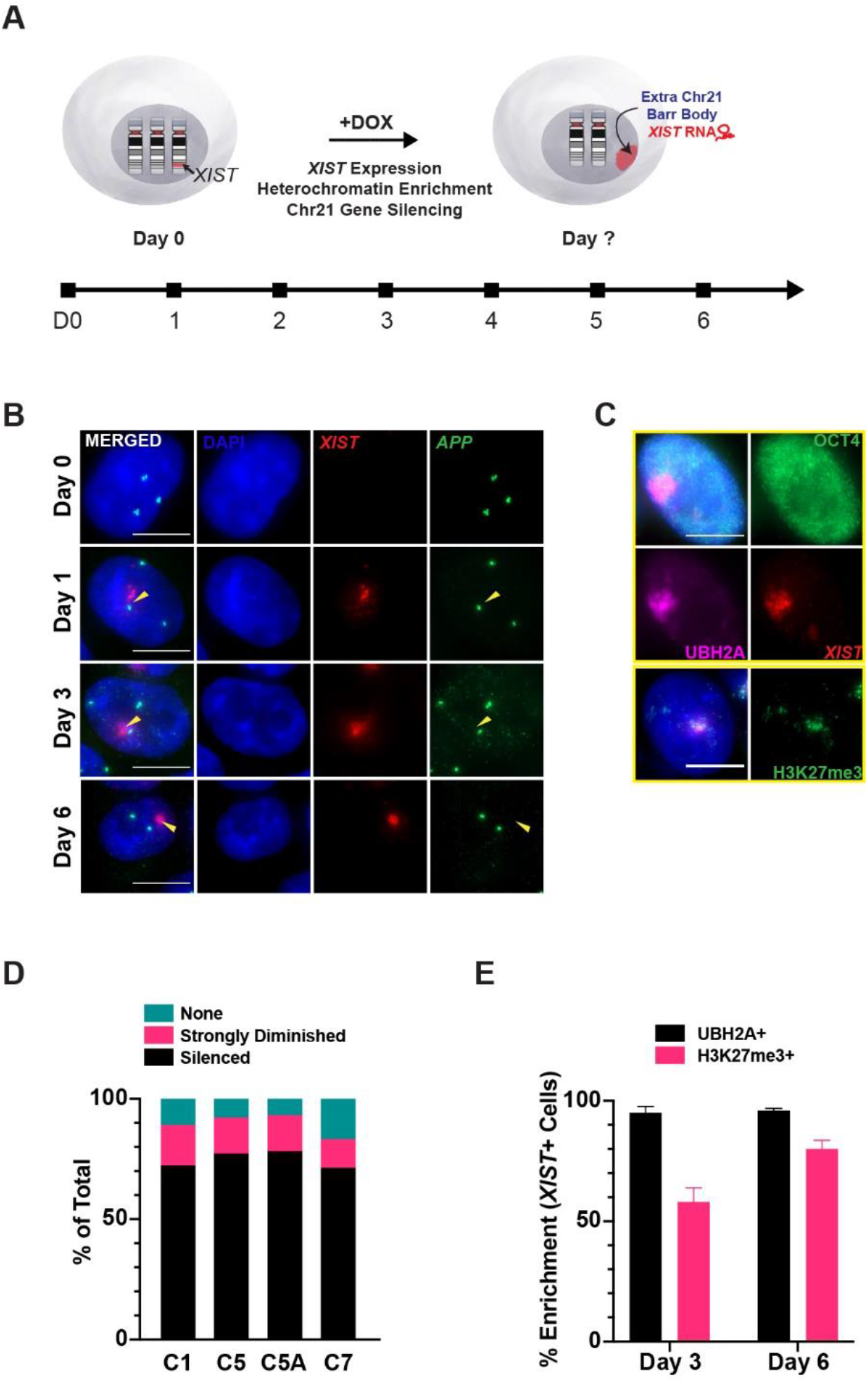
Silencing of one chr21 in human DS iPSCs is essentially complete by day 6. (A) Schematic of approach and kinetic analysis of XIST-mediated chr21 silencing of DS iPSCs. (B) RNA FISH detects three *APP* transcription foci at early time points and later shows late-silencing gene *APP (*green) is essentially silenced by XIST RNA (red) at day 6. (C) Immunofluorescence (top four panels) shows H2Ak119Ub induced by XIST RNA (red) in the nucleus with OCT4 as a marker for pluripotency. Bottom two panels show H3K27me3 (green) with XIST RNA (red). Scale bar = 10 um. (D) Quantification of *APP* gene silencing shows transcription foci from the XIST-coated allele are essentially silenced in ∼90% of cells at Day 6 (n = 300). (E) Enrichment of heterochromatin marks (H3K27me3 and H2Ak119Ub) in XIST+ cells in four transgenic lines (n = 100/line; mean ± SD).

Recently, we demonstrated that trisomy silencing prevented developmental pathogenesis *in vitro* for the best-established DS cell pathology, overproduction of certain hematopoietic cell types (Chiang et al., 2018). This result supports that correction of chr21 overexpression is sufficient to normalize a known DS cell pathogenesis. Subsequently, we used this approach to examine effects of chr21 dosage compensation on *in vitro* neurogenesis. Results showed that chr21 silencing enhanced formation of neurons from neural stem cells, and implicated delayed terminal differentiation of neurons due to elevated Notch signaling in T21 (Czermiński and Lawrence, 2020).

In contrast to studies that focus on a particular tissue or cell type, here we begin by using RNAseq analysis in iPSCs to determine how correcting T21 expression levels impact genome-wide pathways while maintaining pluripotency. The pluripotent stem cell state--in some sense, a theoretical tabula rasa--allows analysis of any effects of chr21 dosage on the gene expression landscape and developmental programming from the onset. Pluripotent cells express an unusually large fraction of the genome including many genes expressed later in specific cell-types (Efroni et al., 2008; Kobayashi and Kikyo, 2015; Postovit et al., 2007), thus potentially providing insight into later developmental programming. Moreover, we purposefully explored transcriptomic effects shortly after chr21 silencing in order to infer the most immediate and direct effects of chr21 gene dosage changes without *a priori* assumptions or emphasis.

Pathway analysis revealed changes in specific genes relevant to current or new hypotheses of DS research, leading us to investigate endothelial cells (ECs) and angiogenesis. While results shown here are relevant to neurodevelopment, we focused more on angiogenesis because this was unanticipated and less studied, and any impact on angiogenesis could affect multiple systems, during development and adulthood. For example, vascular deficits could contribute to pulmonary hypertension, early-onset AD, low incidence of solid tumors, neurodevelopment, or even the higher risk of autism in DS (Asim et al., 2015; Colvin and Yeager, 2017; Ouellette et al., 2020). Limited mouse studies or clinical observations have suggested vasculature deficits may occur in DS (Bush et al., 2019; Galambos et al., 2016; Trevino et al., 2020), although it remains unclear if DS is characterized by an impaired vasculature. Furthermore, such studies cannot address whether any differences in vasculature are a root effect of T21 on angiogenesis or secondary to other pathologies.

In summary, results of initial transcriptomic analysis in a panel of DS iPSCs identified gene pathways impacted by chr21 over-expression, which then led us to examine angiogenesis in DS iPSC-derived ECs. Results reveal a previously unreported *in vitro* cellular phenotype, demonstrating for the first time that T21 expression impacts ECs and angiogenesis.

## RESULTS

Many DS studies describe extensive genome-wide transcriptome changes in cells/tissues; however, these changes will often reflect downstream impact of T21 on development, cell-type representation, or pathological state. Here, our first goal was to compare otherwise identical cell cultures with and without dox-induced *XIST* and identify the more direct transcriptome-wide changes caused by chr21 dosage, distinct from changes in developmental cell state. Jiang et al (2013) described the panel of human transgenic lines used here and demonstrated that *XIST* could silence one chr 21 in iPSC cultures after 20 days of induction (Figure S1). The microarray data generated was not used for genome-wide pathway analysis due to uncertainty regarding whether some cultures differentiated over prolonged culture. Because pluripotent cells are prone to differentiate and change epigenetic state, and to maximize detection of more direct effects of T21, we began by identifying the earliest time after *XIST* induction when chr21 silencing was essentially complete, while maintaining cultures in an undifferentiated state. This shorter timeframe was employed in a panel of four isogenic *XIST*-transgenic clones followed by quantitative analysis of gene expression changes genome-wide.

### Analysis of Chromosome Silencing Kinetics Shows Chr21 Effectively Silenced by Day 6

To determine the optimal timepoint after chr21 is silenced, we examined changes in specific gene expression and heterochromatin marks over several days (Figure 1A). Using RNA FISH, we quantified transcription foci for the chr21 gene, *APP*, which we recently found is one of the last genes silenced (Czermiński and Lawrence, 2020). To indicate heterochromatin formation chromosome wide, we scored enrichment of *XIST*-associated heterochromatin marks, H2Ak119Ub and H3K27me3 (Kohlmaier et al., 2004; Plath et al., 2003; Smith et al., 2004).

Almost all uninduced cells show three *APP* transcription foci. At day 1 of dox, *XIST* RNA accumulates over one chr21 territory but all three *APP* alleles continue to express. By day 6, the vast majority of *XIST*+ cells show complete silencing of that *APP* allele (Figure 1B and Figure 1D). The level of chr21 silencing was consistent for all four transgenic clones. Due to the stochastic silencing of the *TET3G* transactivator transgene, not all cells (∼70%) induced expression of *XIST*, but by day six *XIST+* cells overwhelmingly show that *APP* allele was silenced (Figure 1B). Immunofluorescence (IF) for H2AK119ub and H3K27me3 showed heterochromatin formation coincident with *XIST* RNA (Figure 1E). Hence, this analysis identified day six as the earliest time point when the extra chr21 is effectively silenced, as affirmed below by RNA sequencing, and the shorter time frame was optimal to maintain and compare cultures in a homogeneous pluripotent state.

### Detection of Robust Chr21 Silencing and Transcriptome-wide Changes in Pluripotent Cells

We quantified expression changes in parallel cultures with and without dox-induced *XIST* expression, for each of the four isogenic transgenic lines (*XIST*-/+ cells; Figure 2A). Importantly, this experimental design involves comparison between parallel cultures (-/+ dox-induced *XIST*) rather than between different isogenic cell lines. This avoids epigenetic and potentially genetic variability between lines, as evidenced by PC2 in Figure 2B, which can confound or dampen identification of trisomy-related differences (Kim et al., 2010; Koyanagi-Aoi et al., 2013). The PCA plot (Figure 2B) affirms that in this system the largest variance (PC1) between samples is due to trisomy versus functional disomy (*via* chr21 silencing). We included dox treatment of isogenic lines with the same *TET3G* transgene but lacking the *XIST* transgene (Figure 2A), as a control to minimize artifacts of dox or transactivator interactions. Any differentially expressed genes (DEGs) in the *XIST*-transgenic lines that were similarly affected by dox treatment in control lines were excluded from consideration as DEGs.

**Figure 2:**
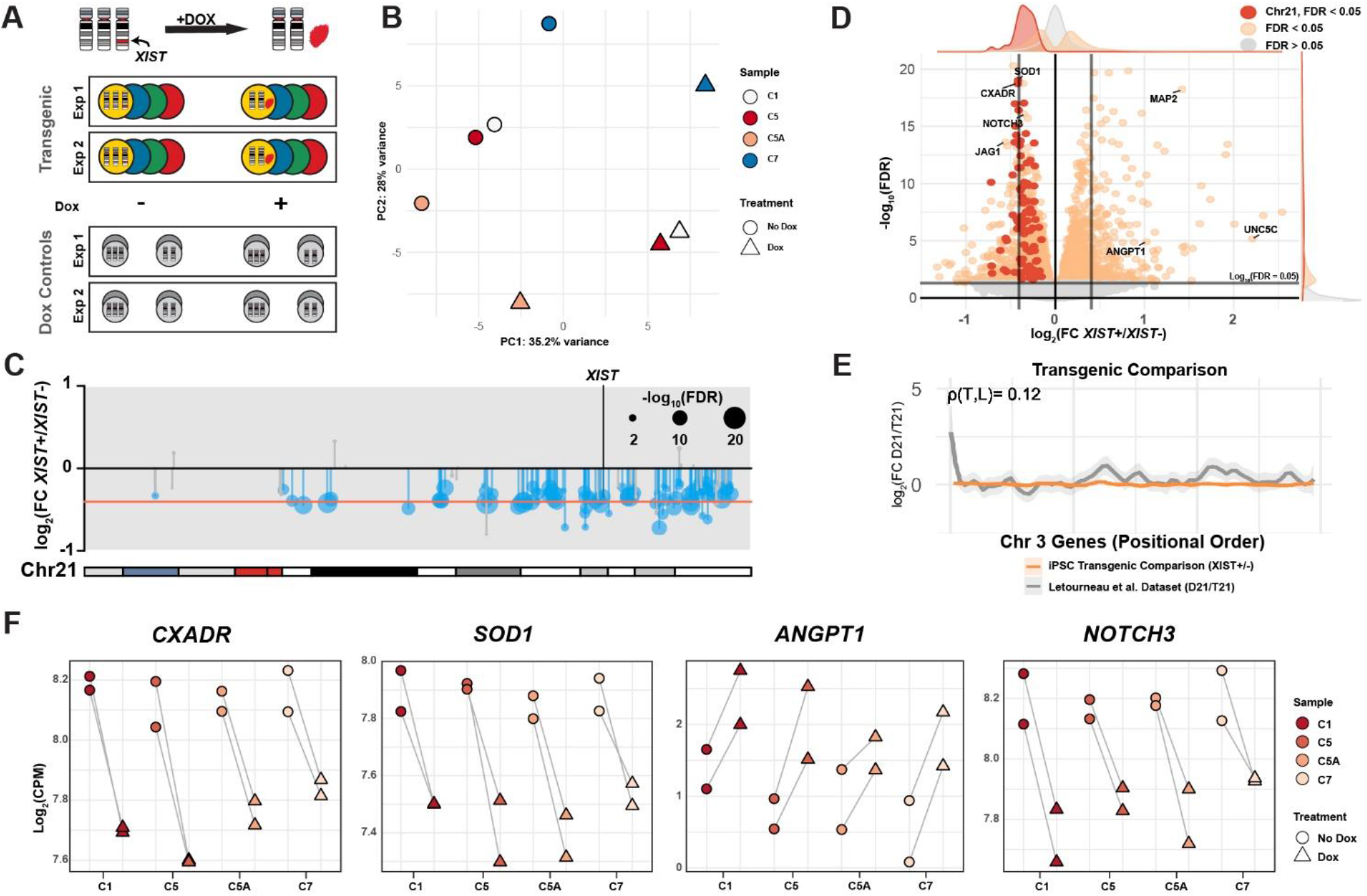
Transcriptomic analysis of chr21 and genome-wide changes rapidly induced by silencing of extra chr21. (A) Experimental design in which parallel cultures of four isogenic *XIST* transgenic DS iPSC lines are examined with and without dox-induction of “trisomy silencing”. Isogenic subclones carrying the *TET3G* transgene but not *XIST* with and without dox were used as treatment controls and analyzed in parallel. (B) Principal component analysis of four transgenic lines of PC1 (x-axis) and PC2 (y-axis). *XIST* and *TET3G* RNAs were excluded from this analysis. (C) Ideogram of chr21 and expression of genes across the chromosome. Theoretical one third reduction in expression (adjusted for 70% of *XIST* expressing cells) in red and each dot is scaled to -log10 (FDR) values. (D) Volcano plot of all expressed genes detected. Horizontal dashed lines represent the theoretical one third reduction or increase in fold change; horizontal dashed line is the FDR value of 0.05 cutoff for differential expression. Dots represent individual genes with chr21 genes in black, FDR < 0.05 in dark gray, and FDR > 0.05 in light gray. (E) Example of local regression plot showing expression changes across chr3 compared to the Letourneau dataset. Pearson correlation (Rho) are reported above each plot. Additional comparisons are graphed in Supplemental Figure 1B. (F) Chr21 genes *CXADR* and *SOD1* expression before and after silencing (left). Non-chr21 genes *ANGPT1* and *NOTCH3* expression before and after silencing (right).

Figure 2C summarizes results for silencing chr21 genes based on analysis of all four clones, showing consistent downregulation of genes across that chromosome. Even though in these initial experiments only ∼70% of cells are induced to express *XIST* (due to technical silencing of the *TET3G* transgene, see STAR Methods), this approach detected an average reduction across chr21 that aligns with the 33% reduction expected (corrected for 70% *XIST*-expressing cells). While some studies have suggested not all chr21 genes are overexpressed due to potential feedback regulation, these results indicate that all or most expressed mRNAs approximate the theoretical one third reduction.

We next examined genome-wide (non-chr21) expression changes in the four clones just after completion of chr21 silencing. As shown in Figure 2D, changes in ∼2500 genes met statistical significance at FDR < 0.05 (Table S1). Many genes important to developmental processes consistently changed in all lines upon chr21 silencing (Figure 2F). This includes genes later expressed in a cell-type specific manner, as illustrated by MAP2, a non-chr21 gene commonly used as a neuronal marker (Figure S2), which consistently increased in all four lines with chr21 silenced. A). In data from our earlier study (Jiang et al., 2013), we noted a similar increase in MAP2 expression, but given the propensity of pluripotent cells to differentiate, we were uncertain whether cultures had diverged in differentiation status during the 20 day dox treatment. However, in this study, we examined parallel cultures just six days after inducing *XIST* and rigorously maintained pluripotency as evidenced by OCT4 staining (Figure 1C). A subsequent study showed that a single iPSC colony could stain brightly for both MAP2 and Oct4 (Gonzales et al., 2018). That study noted higher MAP2 in one trisomic versus disomic isogenic comparison, but this was not consistent between lines, suggesting inter-line variability. Results here using inducible Chr21 silencing indicate that one or more chr21 genes affects MAP2 expression in pluripotent cells, further illustrating iPSCs may transiently express genes which later function in specific developmental programs (Efroni et al., 2008; VanOudenhove et al., 2016).

Before examining potential changes to developmental pathways, we first addressed a distinct hypothesis: that broad transcriptomic changes reflect the general impact of T21 on genome architecture, resulting in large contiguous segments with alternating up- or down-regulated genes across all autosomes. Letourneau et al. reported gene expression dysregulation domains (GEDDs) due to T21, reflecting broad changes to the chromatin landscape impacting the global transcriptome, distinct from more specific pathways. As shown in Figures 2D and S2B, our results show no evidence of GEDDs in the *XIST*-/+ comparisons, and gene expression across each chromosome did not strongly correlate with the Letourneau dataset. While not examined extensively, we also did not detect GEDDs comparing disomic and trisomic isogenic lines, consistent with other results (Do et al., 2015; Sullivan et al., 2016). Since XY, XX, or XXX karyotypes do not physically impact nuclear transcriptome architecture, it is highly improbable that tiny chr21 would do so, hence we focused on investigating whether specific pathways are impacted by chr21 over-expression.

### Chr21 Overexpression Causes Early Dysregulation of Neural Pathways and Notch Pathway

Using the list of non-chr21 DEGs, we performed enrichment analysis using the Gene Ontology (GO) catalog for biological processes. When visualizing the GO terms of upregulated genes in hierarchical order, four major categories--angiogenesis, neurogenesis, cytoskeleton organization, and response to stimulus--were apparent (Figure 3A-B). Even at the pluripotent stage, trisomy silencing significantly impacted genes involved in neurogenesis, with enrichments in GO terms relating to neuronal cell fate, migration, proliferation, and differentiation (Figure 3C and Table S2). These results are in keeping with and significant for the known clinical impact of T21 on cognitive development, and support that chr21 silencing ameliorates these effects *in vitro*.

**Figure 3:**
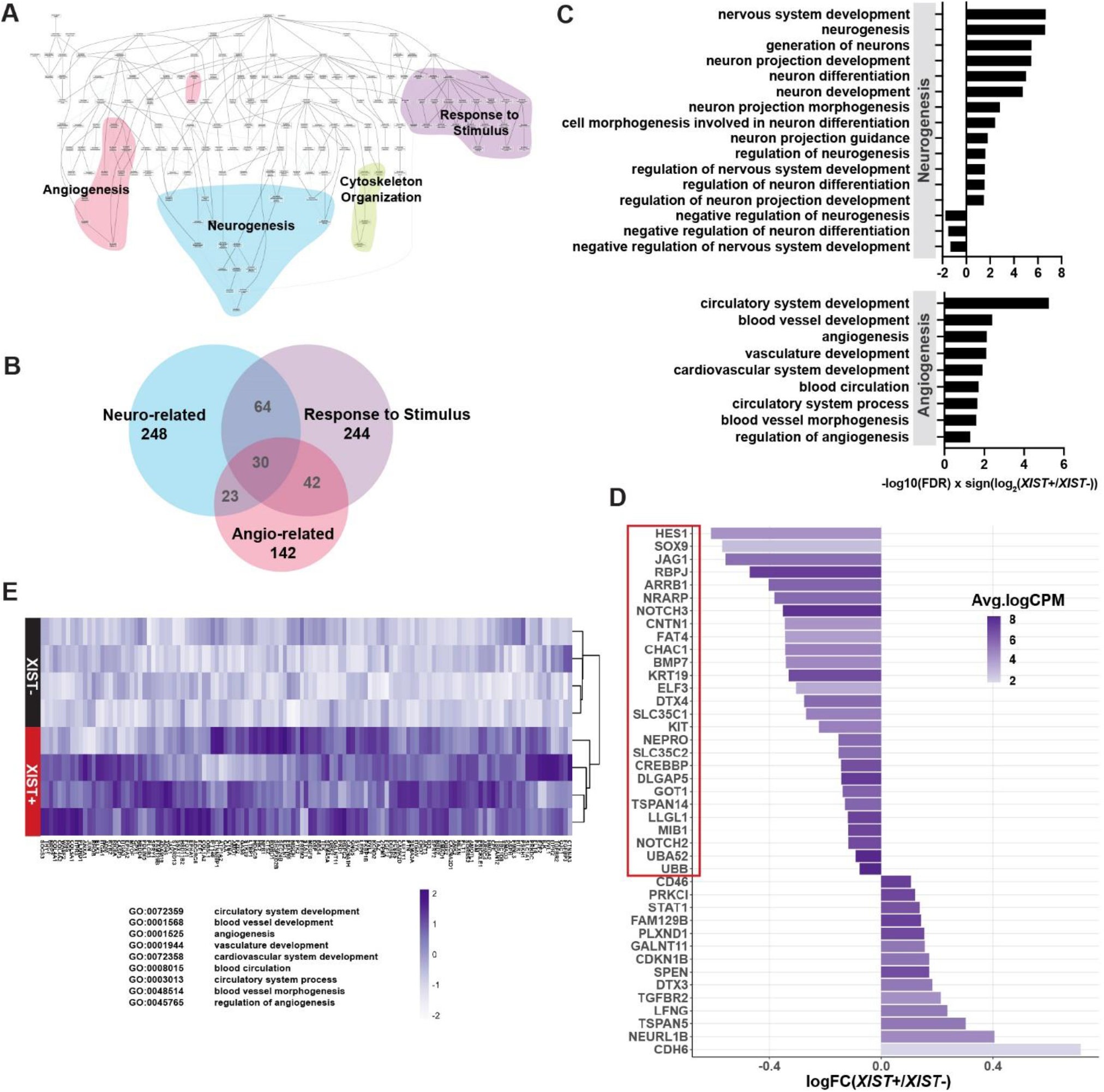
Correcting chr21 dosage upregulates neuro- and angio-related gene sets, and downregulates Notch signaling pathway. (A) Hierarchical tree of biological processes with enriched GO terms among upregulated genes. Colors cluster by similar terms (pink = angiogenesis; blue = neurogenesis; green = cytoskeleton organization and ECM; purple = response to stimulus). (B) Largest GO term clusters were relating to neurogenesis, response to stimulus, and angiogenesis. Genes enriched in these terms show large overlaps which are depicted in the Venn diagram. (C) Subset of significant GO terms relating to neuro- and angio-related terms (*XIST*+/*XIST*-; FDR < 0.05). Notably, *XIST*+ cells show enhancement of neuro- and angio-related pathways and downregulate negative regulation of neural development. (D) Notch genes significantly up- and down-regulated after silencing the extra chr21 (*XIST*+/*XIST*-). Downregulated genes that were significantly enriched in the Notch Pathway in *XIST*+ cells are highlighted in the red box (GO:0007219, GO:0008593; FDR < 0.05). (E) Heatmap of z-score normalized gene expression enriched in angiogenesis-related gene sets (row = clone, column = gene). Rows were clustered by expression. Relevant gene ontology ID and terms are listed below heatmap.

Of particular interest is the overexpression of Notch pathway genes in trisomic iPSCs relative to the same cells with trisomy 21 silencing (Figure 3D; GO:0007219, GO:0008593; FDR<0.05). Overactive Notch signaling in iPSCs supports and extends key findings in our recent study of T21 effects on day 28 neural cells (Czermiński and Lawrence, 2020). That study showed increased Notch signaling and also upregulation of *TTYH1*, a non-chr21 gene recently reported to upregulate Notch via gamma-secretase (Kim et al., 2018). This was linked to delayed terminal differentiation of cycling neural stem cells to form neurons. Here we find that trisomic pluripotent cells also have elevated Notch signaling and also *TTYH1*. This supports findings in the neural cells, but the short time frame of silencing studied here strongly suggests a more direct impact of T21 on Notch signaling, potentially via TTYH1. Results support previous hypotheses that neuronal deficits in DS fetal and adult brains may be due to impaired Notch signaling (Golden and Hyman, 1994; Karlsen and Pakkenberg, 2011), and further suggest that the impact on Notch signaling could occur earlier in development and may not be specific to neural cells.

### Correction of Chr21 Overexpression Upregulates Angiogenesis Pathways

As shown in Figure 3A, neural pathways were most prominently impacted by T21 in iPSCs, but results also implicated a second major cell-type. Surprisingly, the second prominent pathway cluster pointed to endothelial cells (ECs) and their involvement in vascular development and angiogenesis (Figure 3E). Notably, several Semaphorin genes (*SEMA3A, SEMA5A, SEMA4F*) along with its binding receptors--*NRP2, PLXNB1, PLXND1*--were differentially expressed in our dataset; these genes are known to promote angiogenesis, endothelial proliferation, and survival, as well as functions related to cell extension and migrational guidance in ECs. *ADM*--a gene encoding two post-translationally modified proteins, PAMP and AM--is involved in angiogenesis and EC differentiation, and may play an important role in memory retention. In addition, expression of VEGF receptor *FLT1* changes significantly after trisomy silencing. This gene plays an important role in regulating the Notch-VEGF pathway feedback loop in angiogenesis and maintains dynamic endothelial tip and stalk cell subtype identities (Hellström et al., 2007; Phng and Gerhardt, 2009). Both *FLT1* and *NPR1* play critical roles in angiogenesis but are also important in cardiovascular development and neovascular formation (Kitsukawa et al., 1997; Krueger et al., 2011; Oh et al., 2002). Collectively, many genes in angiogenic pathways have overlapping functions in cytoskeletal changes and signaling, many of which were enriched in GO terms relating to response to stimulus (Figure 3B).

Overall, 125 DE genes upregulated after trisomy silencing were enriched in angiogenic GO terms (Figure 3E). As for Notch and neurogenesis, the results indicate that it is an essentially direct effect of chr21 over-expression on pathways involved in angiogenesis, discernible even in pluripotent cells, just after chromosome silencing.

### Trisomic and Chr21-silenced Cells Show Similar Production of Endothelial Progenitor Cells

Since analysis of iPSCs suggested that T21 impacts pathways related to angiogenesis, we further investigated these findings by examining effects of T21 expression on endothelial cells, differentiated from DS iPSCs. First, we examined whether dosage correction of chr21 genes would impact the production of endothelial progenitor cells (EPCs). As outlined in Figure 4A-B, EPCs have a close developmental relationship to hematopoietic cells, both arising from hemogenic epithelium. T21 is known to cause excess production of CD43+ hematopoietic progenitor cells, leading to excess production of megakaryocyte-erythroid progenitor cells (Chiang et al., 2018; Roy et al., 2012; Tunstall-Pedoe et al., 2008), and Chiang et al. (2018) affirmed that transcriptional silencing of chr21 corrected hematopoietic pathogenesis for this known DS cellular phenotype. Therefore, we examined whether T21 impacted the proportions of progenitor cells produced during EC differentiation, examined in all four transgenic lines. As above, we included trisomic and disomic isogenic subclones lacking the *XIST* transgene to control for dox effects. We quantified CD31+CD34+ which mark endothelial progenitor cells or EPCs in parallel cultures differentiated for five days, with/ without chr21 silencing. In this timeframe, no significant difference in the proportion of EPCs was seen (Figure 4C & 4D, Figure S3A), nor did dox affect the EPC populations in controls. This is in line with our previous finding that over-production of hematopoietic cells arises after the hemogenic endothelium is established (Chiang et al., 2018). We also considered whether T21 impacted cell proliferation by quantifying the fraction of S-phase cells in each population. Results show rapid cycling of both cell populations, although a subtle increase in proliferation may be suggested by a marginal increase in S phase cells in *XIST*+ cultures (Figure S3B). Taken together, T21 did not substantially affect developmental programming of EPC production.

**Figure 4:**
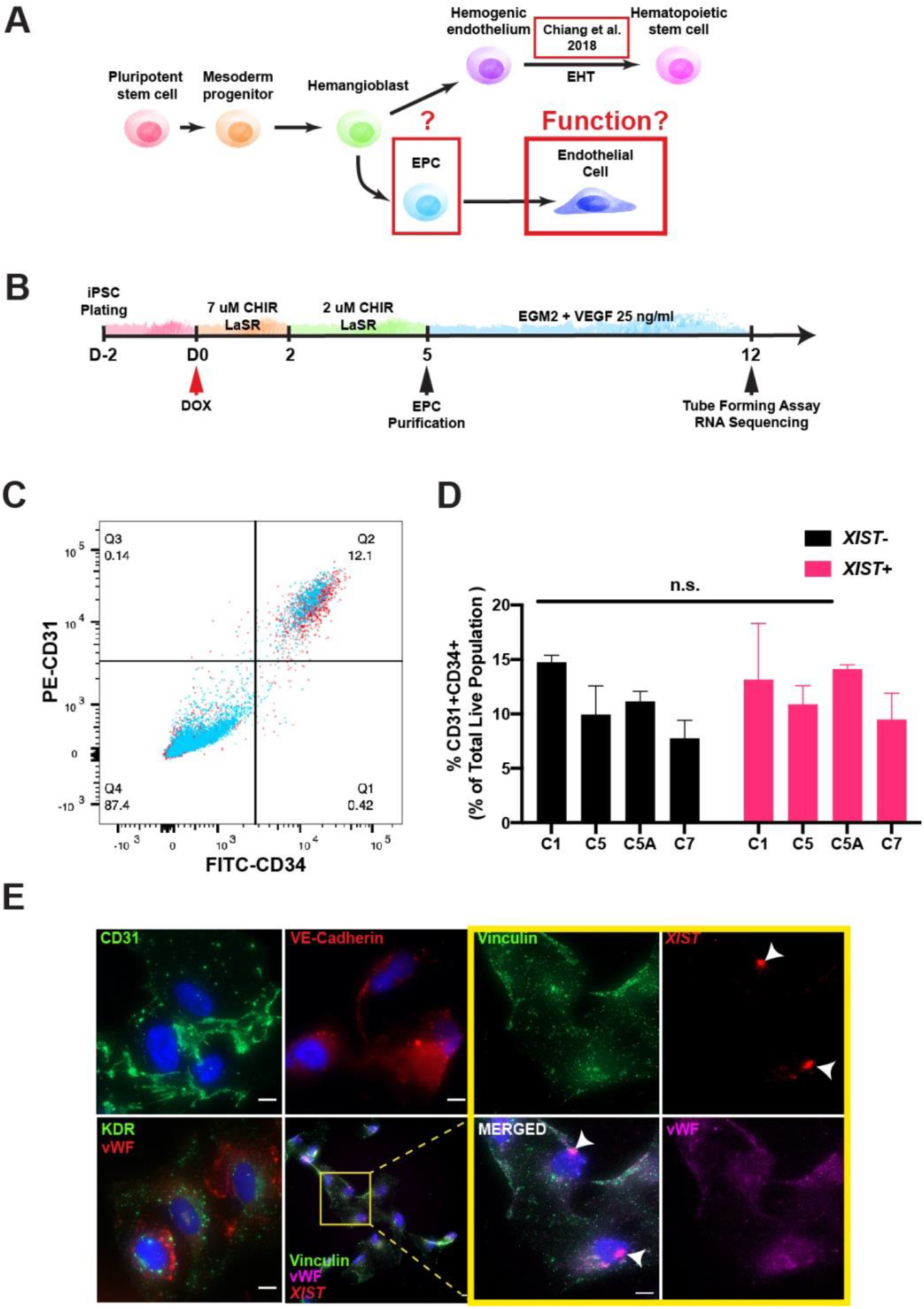
Endothelial Cell Differentiation of DS-iPSCs. (A) Diagram showing the close developmental relationship between hematopoietic and endothelial cell developmental pathways. Defects in hematopoietic lineage cells were characterized by Chiang et al. (2018). Areas of interest (i.e., production of EPCs from the hemangioblast and EC function) highlighted in red box. (B) Experimental schematic of endothelial cell differentiation and downstream assays. Dox is treated at Day 0 of differentiation and EPC population is assessed before purification. After purification, EPCs are matured and later tested for angiogenic function. (C) Representative flow analysis of CD31+ and CD34+ cell populations. Relevant CD31+CD34+ endothelial progenitor cells are in quadrant 2. *XIST-(*red) and *XIST*+ cells (blue) are overlaid. (D) Quantification of the CD31+CD34+ population of each sample (-/+dox; paired t-test; mean ± SD). (E) Immunofluorescence staining of endothelial cell specific markers and RNA FISH of *XIST (*scale bar = 10 um).

### Chr21 Silencing Enhances Endothelial Cell Response to Angiogenic Signals

While dosage compensation of chr21 did not significantly impact EPC production, a distinct question is whether functionality of endothelial cells is impacted (Figure 4A). Endothelial function in angiogenesis is heavily reliant on cell signaling and extension; consistent with this, GO terms relating to response to stimulus, cytoskeleton organization, and migration were strongly implicated in our iPSC data (Figure 3A and Table S2). Hence, we expanded CD31+CD34+ EPCs; after differentiation day 7, both *XIST-* and *XIST*+ cultures exhibited more mature characteristic cobblestone-like EC morphologies. All conditions displayed appropriate endothelial cell markers of vWF, KDR, CD31, and ve-Cadherin by IF (Figure 4E and Figure S3C).

To examine whether T21 impacts the angiogenic function of EC cells, we used an established *in vitro* tube forming assay (DeCicco-Skinner et al., 2014) as a readout to assess the ability of trisomic ECs to migrate and form network structures. In brief, equal numbers of ECs are seeded onto Matrigel-coated plates and treated with factors to promote tube-like formations. These structures form very rapidly as cells respond to signaling cues, and can then be maintained for several hours (Figure 5A). Computerized morphometrics analyses was used to address if there are functional deficits in angiogenic signaling and response in trisomic ECs, comparing cultures of equal cell density, independent of their proliferation.

**Figure 5:**
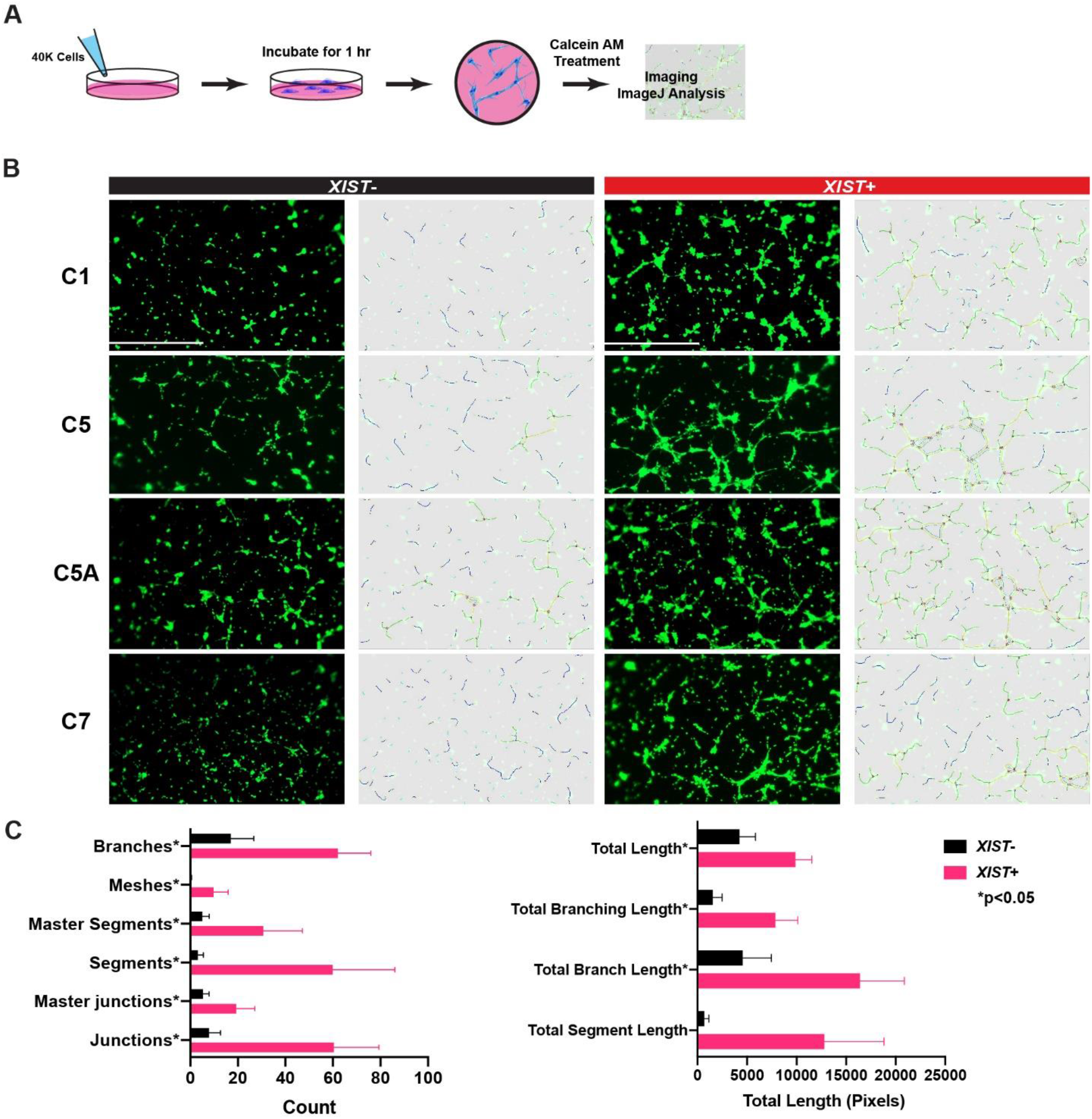
T21 in endothelial cells results in delayed early response to angiogenic signals. (A) Experimental schematic of tube formation assay and analysis. (B) Representative image of endothelial cells after 1 hr of incubation on Matrigel. CalceinAM was used to visualize tube formation. Dox treatment controls were performed in parallel (T21 and D21) in Figure S4B. Scale bar = 1000 um. (C) Quantification of features presented during the tube formation using Angio Analyzer of *XIST*-(black) and *XIST*+ (pink) cells. This experiment was conducted across all four transgenic lines and repeated three times (paired t-test; mean ± SD). (D) Representative image of tube formation from *XIST*- and *XIST*+ cells after 12 hours of incubation. Scale bar = 1000 um. (E) Quantification of tube formation from images using Angiogenesis Analyzer after 12 hours of incubation in *XIST*-(black) and *XIST*+ (pink) cells (n = 4; mean ± SD).

Remarkably, within one hour of plating, a significant difference was observed in network formations between ECs derived with and without *XIST* expression (Figure 5B). This analysis was performed on four different transgenic lines, comparing the same line with and without chr21 silencing, and each comparison repeated in three replicate experiments. Results were analyzed quantitatively by the Angiogenesis Analyzer package for ImageJ (Carpentier et al., 2012). This demonstrated repeatedly that *XIST*+ ECs displayed more branching and network formation in contrast to *XIST*-cells which had more isolated segments and less connections between segments. *XIST*+ cells were able to quickly extend further to form interconnected branches and meshes, consistently in all four transgenic lines (Figure 5C). Although subtle phenotypes can be difficult to assess by comparison between cell lines, we included one isogenic comparison of one disomic and trisomic subclone, which showed similar impairment in tube formation after one hour (Figure S4A-B), with no effect on dox-treated controls. Importantly, the impact of T21 revealed here is a kinetic deficit in signaling response and tubule formation, not a lack of trisomic ECs to undergo angiogenesis. By 12 hours, *XIST*-cells have formed networks similar to their dosage-corrected counterparts, with similar culture density. This is consistent with the fact that angiogenesis clearly occurs in individuals with DS (Figure 5D-E). While the rapid early kinetic response to angiogenic cues is impaired, trisomic cells remain fully viable and eventually fill cultures with similarly dense microvessels (Figure 5D).

Using this inducible system to closely compare trisomic and “euploid” cell function *in vitro*, findings indicate that, production of trisomic EPCs is normal and angiogenesis still occurs, but results revealed a delay in angiogenic response and early micro-vessel formation. This cellular impact demonstrated *in vitro* strongly suggests that effects on microvasculature may be an important, but largely unrecognized, contributor to DS phenotypes, as considered in the Discussion.

### Chr21 Silencing Increases Cell Signaling and Projection Pathways in Endothelial Cells

Given that ECs showed a difference in angiogenic response, we performed transcriptome analysis on isolated populations of ECs to study the underlying pathways and mechanisms impacted by T21. The rapidity of the angiogenic response indicates post-transcriptional control; however, given that ECs cycle rapidly, the mRNAs and proteins involved in signaling and cell projections would need to be actively produced and may be expressed at different levels with chr21 silencing. While in a subset of iPSCs (above) dox did not induce *XIST (*due to silencing of *TET3G*), we subsequently found that selecting for puromycin resistance (inserted with *TET3G* transgene) ensured that dox induced XIST in ∼100% of cells (and maintained expression through differentiation) (See STAR Methods). Figure 6A-B shows the comprehensive repression of genes across chr21, demonstrating power to detect even small (∼33%) decreases in mRNA (unadjusted p < 0.05). The strong trend for all138 chr21 expressed genes is highly significant, and for 62 this small modest expected change met significance even as individual genes (FDR < 0.05; Table S3). Silencing was also confirmed by SNP analysis (Figure S5A); notably, by comparing transgenic lines that silence different chr21 homologs, this approach can examine effects of chr21 SNPs on gene expression or cell phenotypes (Figure S5A). We note that the RNAseq data confirms IF results (Figure 4E) showing EC identity, as we detected expression of EC-specific markers such as CD34, PECAM1, vWF, and CDH5 (VE-cadherin) (Figure S5B) (Goncharov et al., 2017). In addition, principal component analysis showed these ECs clustered well with publicly available iPSC-derived and primary EC datasets (Figure S5C).

**Figure 6:**
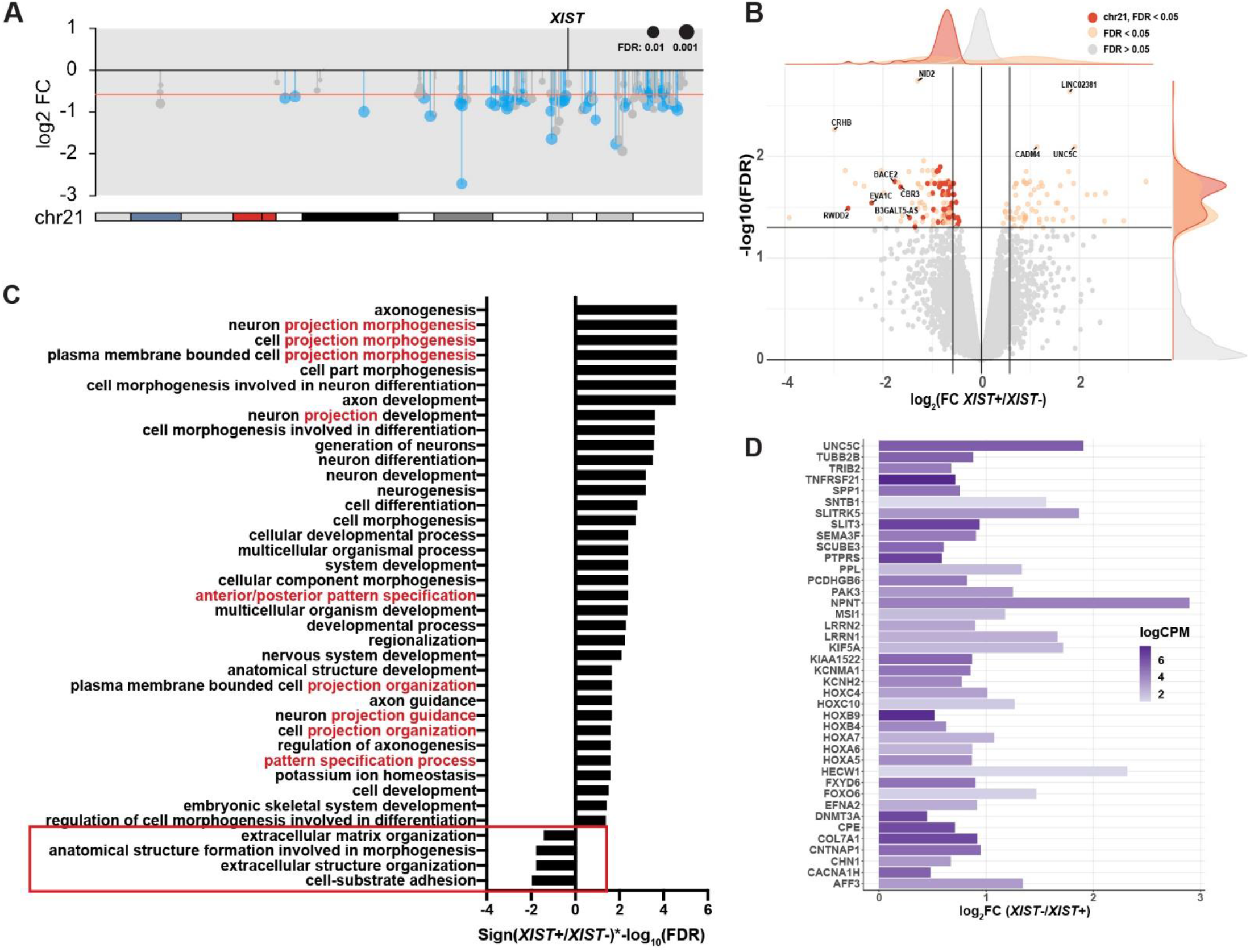
Silencing the extra chr21 in DS endothelial cells perturbs genes relating to cell projections and cell adhesion. (A) Ideogram of chr21 gene expression in DS endothelial cells (*XIST*+/*XIST*-). Red line represents the theoretical one-third reduction in gene expression. Significant genes are in blue (FDR < 0.05), with other expressed genes in gray. (B) Volcano plot of all genes detected. Horizontal lines flanking zero on the x-axis represent the theoretical one third reduction or increase in fold change; horizontal line is the value for cutoff for differential expression (FDR < 0.05). Dots represent individual genes. FDR < 0.05 in orange with chr21 genes in red, and FDR > 0.05 in light gray. (C) Significant GO Terms enriched in the endothelial dataset. Terms are ordered by the direction of fold change and log10 (FDR) value. Many terms shared words relating to projection or patterning among the upregulated gene sets and the few down regulated gene sets were all relating to cell adhesion (red). (D) Genes that were enriched in upregulated GO terms involving branching morphogenesis. All genes met significance threshold (FDR < 0.05).

While chr21 genes were strongly detected, just 120 non-chr21 DEGs were found across the genome. This is substantially fewer than in our iPSC dataset, which may reflect less open euchromatin and more restricted expression profile of a differentiated cell type (see Discussion). Importantly, many genes relating to EC function, including signaling, and migration were upregulated after chr21 silencing, consistent with the more rapid angiogenic response and tube formation in functional assays (Figure 6C) (Wang et al., 2020). In contrast, we saw a significant downregulation of genes involved in cell adhesion and the extracellular matrix (such as *RAC2, PARVB, FMOD*, and some collagen genes). We note that we did not see any gene sets enriched for apoptosis or cell death, consistent with similar vitality of trisomic cultures (Figure S5D; GO:0031589, GO:0043062, GO:0048646, GO:0030198; FDR<0.05). Notably *RAC2* and *PARVB* are important for endothelial cell polarity during neovascular patterning as well as vascular morphogenesis, and dynamic cell migration and projection are important for vascular remodeling and regression during early development (De et al., 2009; Pitter et al., 2018). Therefore, downregulation of these genes after chr21 silencing could facilitate more rapid response to angiogenic cues. *TEK* and *ANGPT2*, associated with hypovascularity in the lungs of DS patients, and thought to be due to elevated levels of anti-angiogenic factors from chr21 (Galambos et al., 2016), were more highly expressed in *XIST*-cells at nominal significance (unadjusted p < 0.05). Corresponding with results here and elsewhere, DS patients have been found to have elevated levels of *ANGPT1* and *ANGPT2*, which are protective against tumor metastasis, providing insight into the less solid tumors in DS (Galambos et al., 2016; Michael et al., 2017).

Similar to our iPSC results, *SEMA3A, EFNA2, UNC5C*, and *PLXN1B* were upregulated in *XIST*+ cells (Figure 6D, Figure S5E). While these genes are enriched in GO terms relating to neuronal processes and axon guidance, they are among a common set of genes involved in cooperative early development of neurons and ECs, with common functions for chemotaxis and branching morphogenesis (Adams and Eichmann, 2010). Additionally, they have a parallel role in outgrowth, using common cues, and are important for neovascularization and structural stability during vascularization (Carmeliet and Tessier-Lavigne, 2005). Similarly, several HOX genes, involved in cell migration and angiogenesis and linked to tumor metastasis (Chung et al., 2009; Toshner et al., 2014), were upregulated with chr21 silencing.

Our tube forming assay demonstrates a functional phenotype in ECs with T21, similar to studies of general Notch-VEGF dynamics and dysregulation (Benedito et al., 2009; Chappell et al., 2013). We detected statistically significant changes in a few Notch genes in ECs, with some trends similar to iPSCs (unadjusted p < 0.05); however, we did not see the same degree of Notch network perturbation in the EC and iPSC transcriptomes. Angiogenic sprouting is guided by two subtypes called tip and stalk cells (Gerhardt, 2013; Ochsenbein et al., 2016), determined and maintained by anti-correlated Notch and VEGF signaling levels, which ensure healthy angiogenic outgrowth (Benedito et al., 2009; Blanco and Gerhardt, 2013). If our EC populations contain both sub-types and proportions are impacted by T21, effects on Notch and VEGF signaling in either cell-type could be somewhat muted (unlike more homogeneous iPSCs).

Overall, silencing the extra ch21 in ECs reduces expression of cell adhesion genes but elevates genes responding to signaling and branching morphogenesis. This fits well with results of *in vitro* functional assays, showing that *XIST*+ ECs (essentially disomic) are more responsive to angiogenic cues and more readily form tubes within the first hour (Figure 5). These kinetic differences in rapid angiogenic response (evident in the first hour) are supported by higher expression levels of genes that support these functions. In total, both transcriptomic data (on ECs and iPSCs) and functional cell assays reveal overexpression of chr21 caused a cell-autonomous deficit in angiogenic response impacting early micro-vessel formation.

## DISCUSSION

Findings here contribute significantly to advancing the poorly understood cell biology of Trisomy 21. Connecting T21 to ECs and angiogenesis has important implications for the multi-systemic impacts of DS, including several comorbidities, but also the beneficial reduction in solid tumors.

By capitalizing on a system to manipulate expression of one chr21 in DS stem cells, we identified the more immediate and direct effects of chr21 overexpression on the transcriptome. Interestingly, we detect less genome-wide DEGs in differentiated cells than pluripotent cells, which have uniquely open chromatin and broad expression (Efroni et al., 2008; Kobayashi and Kikyo, 2015; Postovit et al., 2007), hence they are likely particularly sensitive to levels of regulatory factors. We detect less non-Chr21 DEGs in ECs than would be predicted by recent studies reporting global genomic impact. This may be because our system strives to minimize uncontrolled differences unrelated to T21. While we aimed to compare “homogeneous” cultures of ECs cells, we do not rule out that induced-silencing of trisomy 21 subtly impacts EC status (i.e., proportions of tip and stalk sub-types). Nonetheless, results suggest the important possibility that transcriptome-wide impact of T21 in a given cell-type is more limited than is currently widely thought. Such questions are fundamental to designing potential therapeutic strategies.

Analysis of iPSCs just after chr21 silencing provides a window into more direct transcriptomic impacts, predominantly revealing pathways important to neurogenesis and angiogenesis. The two other main pathway clusters impacted were cell signaling/response to stimulus and cytoskeleton organization, largely shared processes important for both neural and angiogenic function (cell extension, migration, and response). The impact on Notch signaling was notable, even in undifferentiated iPSCs. A recent study using this system also found increased Notch signaling (including *TTYH1*) as the predominant impact, and connected this to reduced terminal differentiation of neurons (Czermiński and Lawrence, 2020). Findings here go further to show rapid impact of chr21 silencing on the Notch pathway in iPSCs, indicating one or more chr21 genes likely impacts Notch regulation more directly, and independent of differentiation state. Importantly, the current study again finds over-expression of *TTYH1* in trisomy, which other studies report regulates Notch via gamma-secretase (Kim et al., 2018). While the pathways we show impacted here are of significant interest for neurogenesis, we focused much of the study on the less anticipated findings regarding angiogenesis.

Most significantly, this study reveals a novel cell type specific phenotype in DS iPSC-derived ECs, demonstrating that T21 affects EC’s early response to form tube-like structures *in vitro*, using an established assay. Clinical reports have observed less dense vasculature in the lungs and thicker vascular walls in DS patients and suggest that this contributes to risks for pulmonary arterial hypertension (Coultas et al., 2010; Galambos et al., 2016). In addition, DS individuals were found to have elevated levels of endostatin, a C-terminus fragment of chr21 gene *COL18A1*, known to inhibit angiogenesis and tumor growth (Seppinen and Pihlajaniemi, 2011; Zorick et al., 2001). DS patients have a lower incidence of solid tumors, potentially due to decreased angiogenesis. Some candidate genes implicated were studied in mice, although single gene perturbations did not explain all angiogenic phenotypes in Ts65Dn mice; overexpression of *RCAN1* and *DYRK1A* were reported to affect proliferation, vascular structures, and tumor allografts (Baek et al., 2009; Minami et al., 2004). However, there are limited studies in humans and there has not been a way to address whether any vascular changes reflect T21 impact on ECs themselves (Hasle et al., 2016). Results here show *in vitro* a direct, cell-autonomous impact of T21 over-expression on EC function in angiogenesis, providing a foundation for future studies in this under-studied area, including pursuit of chr21 genes involved.

Even slight impact on angiogenesis and vasculature could contribute to the pleiotropic effects of DS, potentially during both development and adulthood. This finding is clearly relevant for increased risk of pulmonary hypertension, a substantial DS comorbidity, and could exacerbate other pathologies, such as early-onset AD. In addition, ECs also function in other ways, such as EC crosstalk with inflammatory cells that is critical for immune response. This could contribute to the immune dysregulation and inflammation recently shown to be prominent in DS (Sullivan et al., 2016). As discussed below, vascular deficits during development could have earlier and widespread consequences.

If Notch signaling and ECs are impacted in early embryos, this could have a significant, potentially stochastic, impact on T21 conceptions, which have high rates of spontaneous loss and defects in heart and lung organogenesis (Llurba et al., 2014; Morris et al., 1999). Our data suggests T21 impacts genes involved in neovascularization important in embryonic development. Further, impaired Notch signaling in DS would likely disrupt the cooperative function of the neurovascular unit (NVU)--made up of ECs, pericytes, glia and neurons--that maintain the stem cell niche, important for early cognitive development, facilitating neovascularization and glia/neuron positioning and maturation (Bell et al., 2020; Shen et al., 2004). The importance of the vascular contribution to neurodevelopment was recently illustrated by the finding that autism (linked to 16p deletion) is associated with deficits in brain microvasculature (Ouellette et al., 2020) and we note that a substantial fraction of children with DS, while sociable, have autistic repetitive behaviors and are diagnosed with ASD (Capone et al., 2005).

Finally, the impact on ECs and angiogenic function shown here has important implications for early-onset AD, which occurs in ∼80% of DS individuals, 20-30 years before AD in the non-DS population (Wiseman et al., 2015). This is clearly related to trisomy for the *APP* gene, but other evidence suggests other chr21 encoded factors likely influence this. Notably, cerebral amyloid angiopathy and microbleeds are more frequent in AD-DS, and NVU impairment may reduce amyloid plaque clearance or increase vascular damage (Helman et al., 2019; Sagare et al., 2013). Hence, it has been hypothesized that deficits in angiogenesis might play a critical role in AD in DS, and possibly cognitive development (Carmona-Iragui et al., 2019; Drachman et al., 2017). It has also been speculated that Notch signaling may be impacted and contribute to DS AD-related vascular pathology (Cho et al., 2019; Drachman, 2014), however such changes could be downstream of other pathologies. Our findings now provide evidence of a direct impact of T21 on angiogenesis. Finally, we note a small subset of individuals with DS undergo cognitive regression as younger adults, distinct from AD dementia (Mircher et al., 2017). Any deficit of brain vascularization, which may vary between individuals, could have progressive consequences beyond early development. Given that angiogenesis also plays an important role in NPC migration after injury, angiogenic impairments could contribute to any cognitive decline during adulthood and aging.

For all these reasons, it will be important for future studies to further examine the effects of T21 on ECs and angiogenesis in DS and the interplay between angiogenesis and other systems affected in DS.

## LIMITATIONS OF THE STUDY

This study uses a panel of all-isogenic sublines of iPS cells and relies mostly on direct comparison of identical cell populations (just different culture dishes), with and without induced Chr21 silencing. While this approach is designed to minimize variables between people (or even isogenic lines), comparison of all-isogenic cells limits findings to the genetic background of one individual. Using this “reductionist” approach allowed us to show a direct, cell-autonomous impact to Chr21 expression on certain pathways and angiogenic function; however, clinical studies in diverse individuals are a key complementary approach, required to draw conclusions about how generalizable findings here are to the broader DS population. Angiogenesis is under-studied in DS; however, we have cited several clinical studies which suggest vascular changes are more broadly evident. Finally, we incorporated controls to correct for effects of doxycycline on individual genes (independent of XIST-induction), but statistical corrections are imperfect and could introduce some error.

## ACKNOWLEDGEMENTS

The authors would like to acknowledge other members of the Lawrence lab, in particular Jen-Chieh Chiang and Jan Czermiński and Oliver King for feedback and guidance during this work or manuscript preparation. This work was supported by the RF1 AG056302 and R01 HD091357 to J.B.L. and F31 HD095588 to J.E.M.

## AUTHOR CONTRIBUTIONS

This study was conceptualized by J.E.M. and J.B.L. Experiments and data analysis were performed by J.E.M. The manuscript was written by J.E.M. and subsequently revised by J.E.M. and J.B.L.

## METHODS

### RESOURCE AVAILABILITY

#### Lead Contact

Further information and requests for resources and reagents should be directed to and will be fulfilled by the lead contact, Jeanne B. Lawrence.

#### Materials Availability

This study did not generate new unique reagents.

#### Data and code availability

RNA sequencing datasets generated by this study have been deposited at GEO (GSE166849).

### EXPERIMENTAL MODEL AND SUBJECT DETAILS

#### Human Cell Lines

Jiang et al. (2013) generated and characterized the transgenic DS hiPSC lines used in this study. The transgenic DS hiPSC lines (C1, C5, C5A, C7) were generated from the original male DS hiPSC parental line, which was obtained by George Q. Daley (Park et al., 2008). All DS hiPSC lines in this study contain the transcriptional activator gene, *TET3G*, at the AAVS1 safe harbor locus. Each transgene line with the exception of C5A was generated from an independent *XIST* transgene integration event into an intron of the *DYRK1A* locus. *XIST* is expressed when treated with dox, which was described by Jiang et al (2013). Isogenic trisomic and disomic lines containing *TET3G* but not the *XIST* transgene were included as dox controls (“Tri” and “Dis”). All iPSCs were cultured with Essential 8™ medium (ThermoFisher) in a feeder-free condition on vitronectin-coated plates. Cells were maintained at 37°C, 20% O,_2_, and 5% CO_2_, and passaged every 3-5 days (∼80% confluency) using 500 uM EDTA in PBS. Expression of *XIST* was induced by adding doxycycline at a final concentration of 500 ng/ml while maintaining cultures in the pluripotent stage or directly upon differentiation. Experiments for dox treatment in iPSC cultures were performed twice.

### METHODS DETAILS

#### Monolayer Endothelial Cell Differentiation

The endothelial cell differentiation of DS hiPSC lines were adapted from Lian et al. (2014) and Patsch et al (2015). In brief, two days before differentiation, hiPSCs were dissociated into a single cell suspension using TrypLE Express (ThermoFisher). The cells were resuspended in Essential 8 media with 10 uM Rock inhibitor Y-27632 (Tocris Bioscience) and seeded at 40,000 cells/well in a vitronectin-coated 12-well plate. Cells were fed with Essential 8 media the next day. On Day 0 and Day 1, the media was replaced with LaSR media as described in Lian et al (2014) supplemented with 8 uM CHIR99021 (Tocris Bioscience). On Day 2 and 4, the media was replaced with LaSR media supplemented with 2 uM CHIR99021. On Day 5, cells were dissociated using TrypLE Express and enriched for CD34+ endothelial progenitor cells using the CD34 MicroBead Kit (Miltenyi Biotec). Purified cells resuspended in EGM-2 media (Lonza) supplemented with 25 ng/ml VEGF165 (PEPROTECH). Cells were seeded onto Collagen I coated plates (50-100 ng/ml in 0.02 M acetic acid) and expanded for downstream experiments. Samples included all transgenic lines with or without dox and one isogenic disomic and trisomic line as a dox control. Three independent differentiations were performed.

#### Cell fixation, RNA fluorescence in situ hybridization, and immunofluorescence staining

Coverslips prepared for RNA fluorescence *in situ* hybridization (FISH) staining were adapted from previously published protocols (Byron et al., 2013; Jiang et al., 2013). In brief, coverslips were coated with appropriate attachment protein solution and seeded with cells. After cell attachment, coverslips were fixed with 4% paraformaldehyde in 1X phosphate buffer saline solution (PBS) for 10 minutes then extracted with 0.5% Triton-X in 10mM vanadyl ribonuclease complex (VRC) for 3 minutes and stored in cold 1X PBS or 70% ethanol. For IF, coverslips were fixed as described for RNA FISH or with 100% cold methanol for 10 minutes. For RNA FISH, we used a Stellaris probe (Biosearch Technologies, SMF-2038-1) to detect *XIST* RNA. The *APP* probe was generated using a BAC from BACPAC Resources (RP11-910G8) and labeled via nick translation with Digoxigenin-dUTP (Roche). To dual stain for protein and RNA, RNAsin (Promega) was added to the primary and secondary antibody stains as described by Byron et al (2013). To assess proliferation BrdU was incubated for two hours in day 10 endothelial cells and fixed as described above. Coverslips were incubated in 70% formamide in 2x SSC for 5 minutes followed by a 5 min dehydration step in 70% and 100% cold ethanol, then stained for IF. All antibodies used are listed in the Key Resources Table.

#### Microscopy

IF and RNA FISH images used a Zeiss AxioObserver 7 with a Flash 4.0 LT CMOS camera (Hamamatsu). Brightness and contrast were corrected in *Fiji (Schindelin et al., 2012, 2015)* to best represent what was observed by eye.

#### RNA isolation and library preparation

RNA samples were extracted using Trizol® Reagent (ThermoFisher) adapted from the manufacturer’s instructions, using 100% ethanol in place of isopropanol. Precipitated RNA was resuspended in RNase- and DNase-free water and DNA digestion was performed using DNase I (Roche) with 2U/ul of RNasin® Plus (Promega) for 1 hr at room temperature. RNeasy® Mini Kit was used for RNA purification and DNase I removal following the RNA Cleanup protocol from the manufacturer’s instructions. RNA quality and purity was assessed via Fragment Analyzer (Advanced Analytical Technologies, Inc). All samples received an RQN score of >8.0.

The NEBNext® Ultra™ II Directional RNA Library Prep Kit for Illumina®, NEBNext® Poly (A) mRNA Magnetic Isolation Module, and NEBNext® Multiplex Oligos for Illumina® were used to prepare sequencing libraries following the manufacturer’s instructions (New England Biolabs, Inc). All libraries were prepared with 1 ug of starting RNA. Sequencing was performed by the University of Massachusetts Medical School Deep Sequencing Core with 7-10 million reads per sample.

#### Flow cytometry

Cells from monolayer endothelial cell differentiation were dissociated using TrypLE Express (ThermoFisher) according to manufacturer’s instructions, dissociated in a wash buffer (1x HBSS, 2 mM EDTA, 0.5% BSA). Dissociated cells were run through a 30 uM MACS SmartStrainer (Miltenyi Biotec). One million cells were resuspended in the wash buffer. Conjugated antibodies were added to each sample and incubated for 1 hr. Cells were washed and resuspended with 300 ul of flow cytometry running buffer (1x HBSS, 2mM EDTA, 2% BSA, 20 mM HEPES pH 7.0). The Flow Cytometry Core Facility detected the staining on BD LSR II Flow Cytometer and analysis was completed using FlowJo™ software (Becton, Dickinson and Company, 2019).

#### Tube Forming Assay and analysis

The tube forming assay was adapted and optimized from DeCicco-Skinner et al (2014). After enrichment for CD34+ endothelial progenitor cells (CD34 MicroBead Kit, Miltenyi Biotec), cells were expanded for one week in a T75 flask. When confluent, the cells were detached using TrypLE Express. The cells were resuspended at 40,000 cells/well in a 12-well plate coated with Matrigel and incubated for 1 hour. Each condition had three replicate wells. Cells were treated with CalceinAM (5 uM working concentration; ThermoFisher) and incubated for at least 15 minutes before imaging. Experiment was repeated for three separate EC differentiations. Analysis of tube forming assay was conducted using the plugin *Angiogenesis Analyzer (Carpentier et al., 2012)* in *Fiji (Schindelin et al., 2012, 2015)*.

### QUANTIFICATION AND STATISTICAL ANALYSIS

#### RNA sequencing analysis

Libraries were aligned to genome build GRch38 using *HISAT2 (Kim et al., 2019)* and mapped reads were counted using *featureCounts* in the *subread* package (Liao et al., 2014). Multi-mapped reads were excluded from analysis. Subsequent gene expression normalization and differential expression analysis was conducted using the *edgeR* package within R (McCarthy et al., 2012; R Core Team, 2021; Robinson et al., 2010). Gene detection cutoff was set to mean CPM > 1 across samples for differential expression analysis. Replicates were summed and p-values were generated via quasi-likelihood F-test in *edgeR*. Differential expression analysis was visualized by the R packages *ggplot2, karyoploteR*, and *pheatmap (Gel and Serra, 2017; Kolde, 2019; Wickham, 2016)*. For the EC datasets, surrogate variables were estimated using the *svaseq* command in the *sva* package (Leek et al., 2020) and incorporated into the model design. Isogenic trisomy and disomy lines (contain the transactivator, but does not contain the transgene *XIST*) were used as dox controls. Genes differentially expressed in the transgenic and control lines after treatment of dox (i.e., changing in the same direction and more than half the magnitude of logFC is explained by dox) were excluded from pathway analysis. We used *goana* for GO enrichment analysis from the R package *limma*, correcting for transcript length bias (Ritchie et al., 2015), and visualized hierarchical trees with the *AmiGO* web application (Ashburner et al., 2000; Carbon et al., 2009; Gene Ontology Consortium, 2021).

#### Statistics

Graphs were generated using GraphPad Prism 9 and R. Specific details regarding statistical tests, value of n, and other graph features are detailed in the figure legends. RNA sequencing and analysis methods are detailed above.

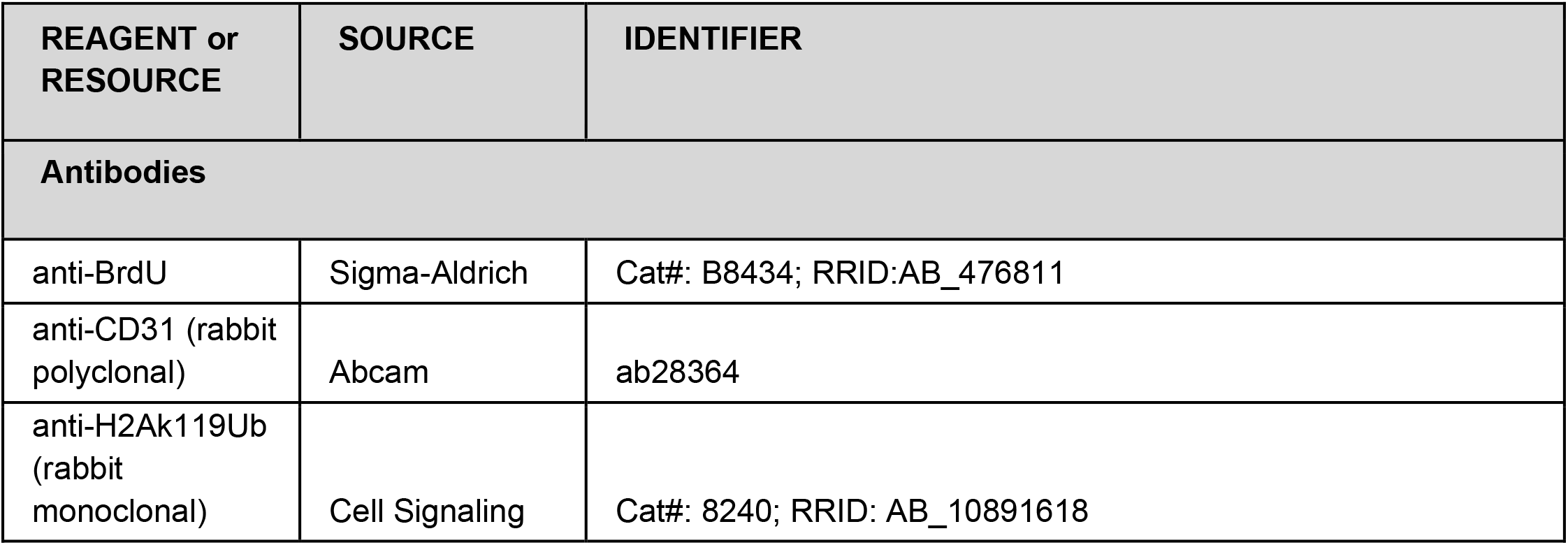
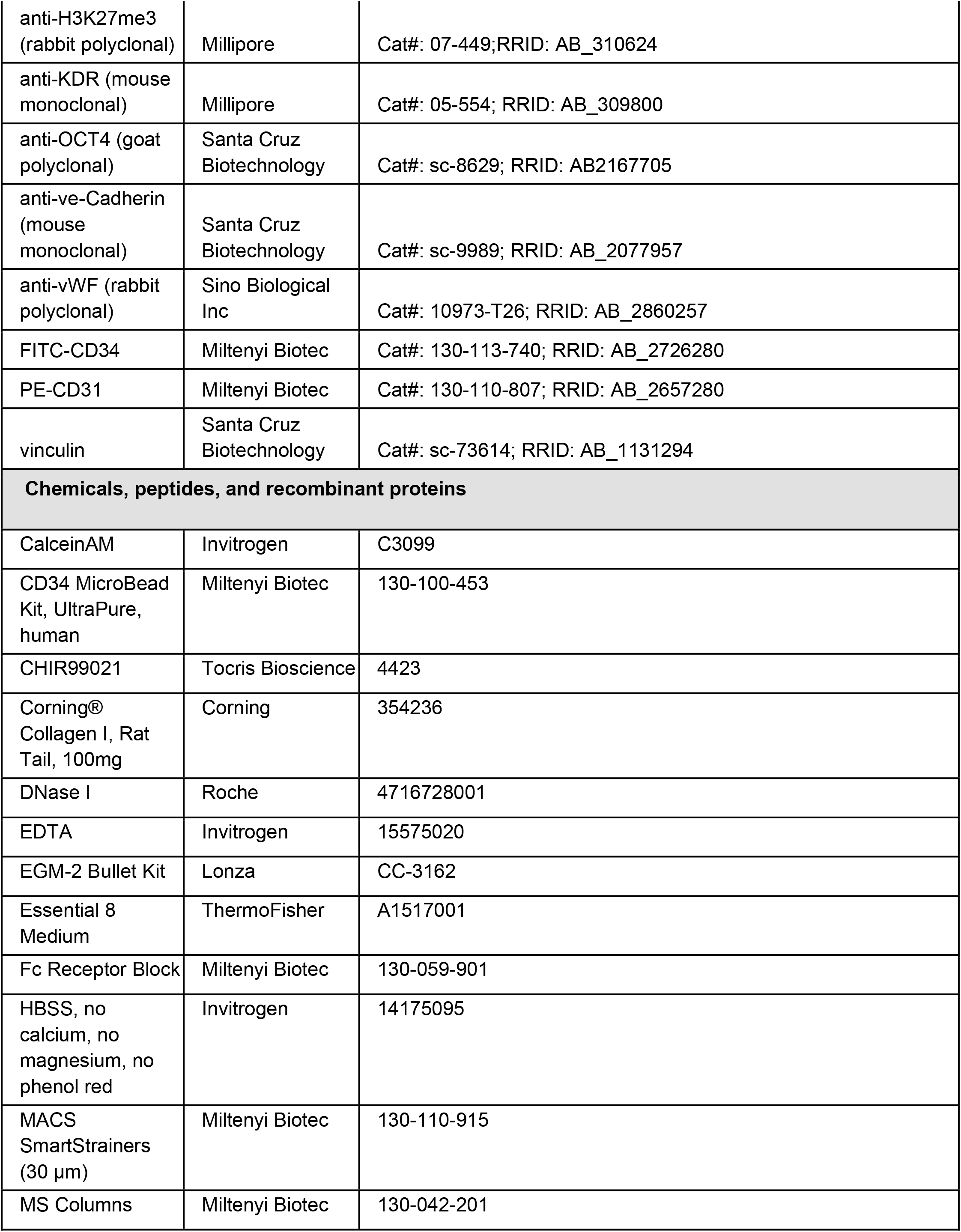
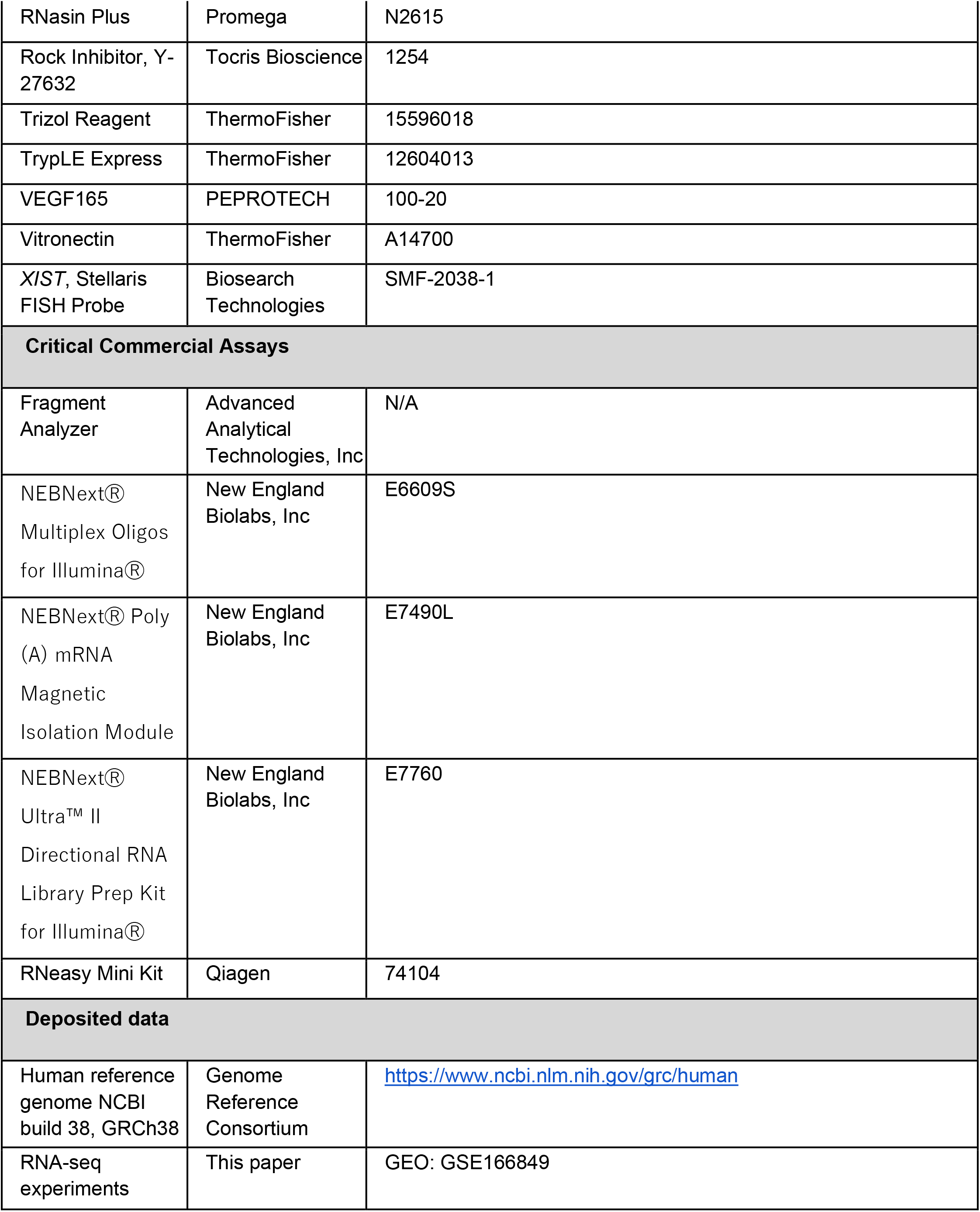
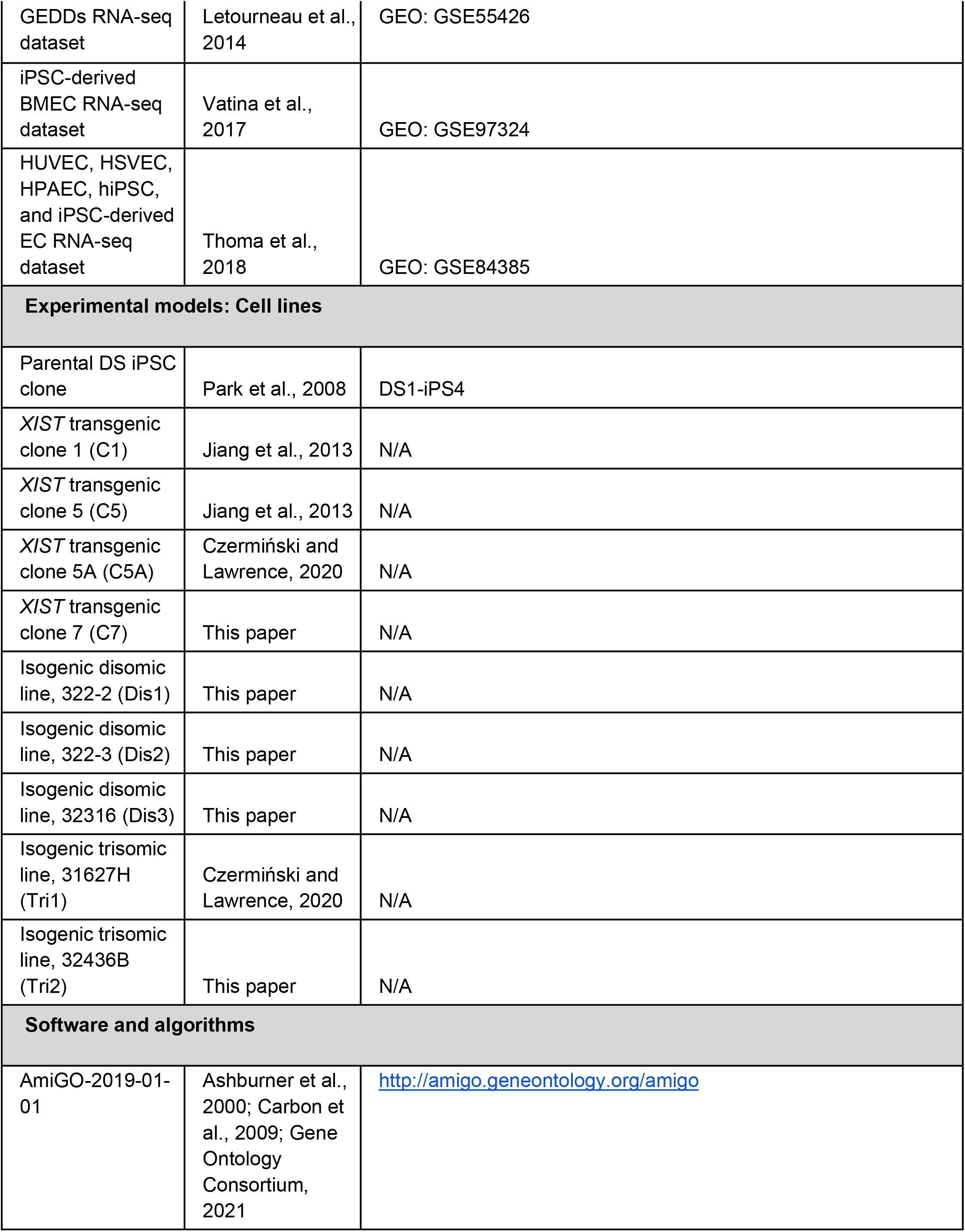
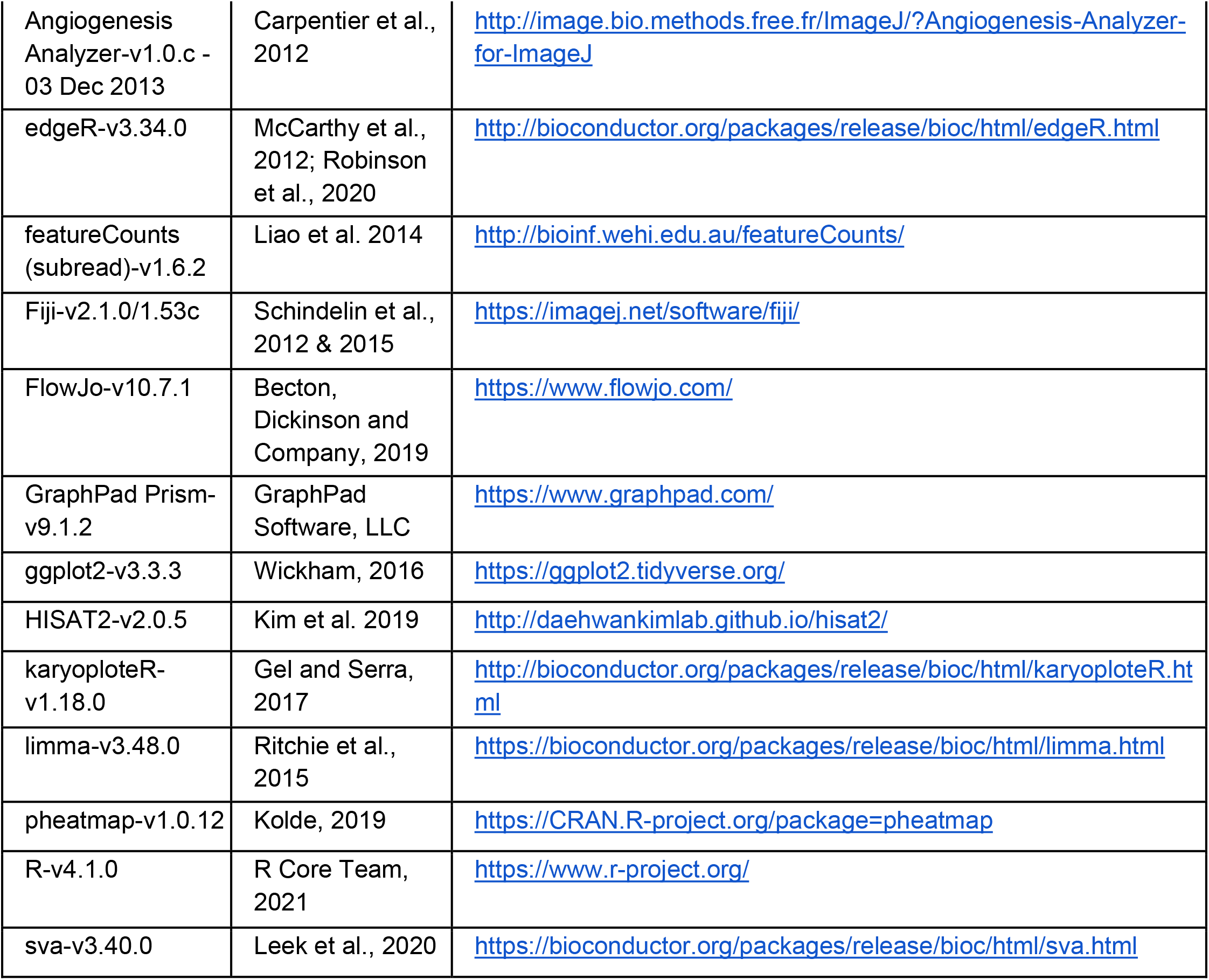

## REFERENCES

Adams, R.H., and Eichmann, A. (2010). Axon Guidance Molecules in Vascular Patterning. Cold Spring Harb. Perspect. Biol. 2. https://doi.org/10.1101/cshperspect.a001875.

Antonarakis, S.E., Skotko, B.G., Rafii, M.S., Strydom, A., Pape, S.E., Bianchi, D.W., Sherman, S.L., and Reeves, R.H. (2020). Down syndrome. Nat Rev Dis Primers 6, 9. https://doi.org/10.1038/s41572-019-0143-7.

Ashburner, M., Ball, C.A., Blake, J.A., Botstein, D., Butler, H., Cherry, J.M., Davis, A.P., Dolinski, K., Dwight, S.S., Eppig, J.T., et al. (2000). Gene ontology: tool for the unification of biology. The Gene Ontology Consortium. Nat. Genet. 25, 25–29. https://doi.org/10.1038/75556.

Asim, A., Kumar, A., Muthuswamy, S., Jain, S., and Agarwal, S. (2015). “Down syndrome: an insight of the disease.” J. Biomed. Sci. 22, 41. https://doi.org/10.1186/s12929-015-0138-y.

Baek, K.-H., Zaslavsky, A., Lynch, R.C., Britt, C., Okada, Y., Siarey, R.J., Lensch, M.W., Park, I.-H., Yoon, S.S., Minami, T., et al. (2009). Down’s syndrome suppression of tumour growth and the role of the calcineurin inhibitor DSCR1. Nature 459, 1126–1130. https://doi.org/10.1038/nature08062.

Becton, Dickinson and Company (2019). Flow Jo^™^ Software.

Bell, A.H., Miller, S.L., Castillo-Melendez, M., and Malhotra, A. (2020). The Neurovascular Unit: Effects of Brain Insults During the Perinatal Period. Front. Neurosci. 13, 1452. https://doi.org/10.3389/fnins.2019.01452.

Benedito, R., Roca, C., Sörensen, I., Adams, S., Gossler, A., Fruttiger, M., and Adams, R.H. (2009). The notch ligands Dll4 and Jagged1 have opposing effects on angiogenesis. Cell 137, 1124–1135. https://doi.org/10.1016/j.cell.2009.03.025.

Blanco, R., and Gerhardt, H. (2013). VEGF and Notch in tip and stalk cell selection. Cold Spring Harb. Perspect. Med. 3, a006569. https://doi.org/10.1101/cshperspect.a006569.

Brown, C.J., Hendrich, B.D., Rupert, J.L., Lafrenière, R.G., Xing, Y., Lawrence, J., and Willard, H.F. (1992). The human XIST gene: analysis of a 17 kb inactive X-specific RNA that contains conserved repeats and is highly localized within the nucleus. Cell 71, 527–542. https://doi.org/10.1016/0092-8674(92)90520-M.

Bush, D., Wolter-Warmerdam, K., Wagner, B.D., Galambos, C., Ivy, D.D., Abman, S., McMorrow, D., and Hickey, F. (2019). EXPRESS: Angiogenic Profile Identifies Pulmonary Hypertension in Children with Down Syndrome. Pulm. Circ. 2045894019866549. https://doi.org/10.1177/2045894019866549.

Byron, M., Hall, L.L., and Lawrence, J.B. (2013). A multifaceted FISH approach to study endogenous RNAs and DNAs in native nuclear and cell structures. Curr. Protoc. Hum. Genet. Chapter 4, Unit 4.15. https://doi.org/10.1002/0471142905.hg0415s76.

Capone, G.T., Grados, M.A., Kaufmann, W.E., Bernad-Ripoll, S., and Jewell, A. (2005). Down syndrome and comorbid autism-spectrum disorder: characterization using the aberrant behavior checklist. Am. J. Med. Genet. A 134, 373–380. https://doi.org/10.1002/ajmg.a.30622.

Carbon, S., Ireland, A., Mungall, C.J., Shu, S., Marshall, B., Lewis, S., AmiGO Hub, and Web Presence Working Group (2009). AmiGO: online access to ontology and annotation data. Bioinformatics 25, 288–289. https://doi.org/10.1093/bioinformatics/btn615.

Carmeliet, P., and Tessier-Lavigne, M. (2005). Common mechanisms of nerve and blood vessel wiring. Nature 436, 193–200. https://doi.org/10.1038/nature03875.

Carmona-Iragui, M., Videla, L., Lleó, A., and Fortea, J. (2019). Down syndrome, Alzheimer disease and cerebral amyloid angiopathy: the complex triangle of brain amyloidosis. Dev. Neurobiol. https://doi.org/10.1002/dneu.22709.

Carpentier, G., Martinelli, M., Courty, J., and and Cascone, I. (2012). Angiogenesis analyzer. 4th ImageJ User and Developer Conference Proceedings 198–201..

Chappell, J.C., Mouillesseaux, K.P., and Bautch, V.L. (2013). Flt-1 (vascular endothelial growth factor receptor-1) is essential for the vascular endothelial growth factor-Notch feedback loop during angiogenesis. Arterioscler. Thromb. Vasc. Biol. 33, 1952–1959. https://doi.org/10.1161/ATVBAHA.113.301805.

Chiang, J.-C., Jiang, J., Newburger, P.E., and Lawrence, J.B. (2018). Trisomy silencing by XIST normalizes Down syndrome cell pathogenesis demonstrated for hematopoietic defects in vitro. Nat. Commun. 9, 5180. https://doi.org/10.1038/s41467-018-07630-y.

Cho, S.-J., Yun, S.-M., Jo, C., Jeong, J., Park, M.H., Han, C., and Koh, Y.H. (2019). Altered expression of Notch1 in Alzheimer’s disease. PLoS One 14, e0224941. https://doi.org/10.1371/journal.pone.0224941.

Chung, N., Jee, B.K., Chae, S.W., Jeon, Y.-W., Lee, K.H., and Rha, H.K. (2009). HOX gene analysis of endothelial cell differentiation in human bone marrow-derived mesenchymal stem cells. Mol. Biol. Rep. 36, 227–235. https://doi.org/10.1007/s11033-007-9171-6.

Clemson, C.M., McNeil, J.A., Willard, H.F., and Lawrence, J.B. (1996). XIST RNA paints the inactive X chromosome at interphase: evidence for a novel RNA involved in nuclear/chromosome structure. J. Cell Biol. 132, 259–275. https://doi.org/10.1083/jcb.132.3.259.

Colvin, K.L., and Yeager, M.E. (2017). What people with Down Syndrome can teach us about cardiopulmonary disease. Eur. Respir. Rev. 26. https://doi.org/10.1183/16000617.0098-2016.

Coultas, L., Nieuwenhuis, E., Anderson, G.A., Cabezas, J., Nagy, A., Henkelman, R.M., Hui, C.-C., and Rossant, J. (2010). Hedgehog regulates distinct vascular patterning events through VEGF-dependent and - independent mechanisms. Blood 116, 653–660. https://doi.org/10.1182/blood-2009-12-256644.

Creamer, K.M., and Lawrence, J.B. (2017). XIST RNA: a window into the broader role of RNA in nuclear chromosome architecture. Philos. Trans. R. Soc. Lond. B Biol. Sci. 372. https://doi.org/10.1098/rstb.2016.0360.

Czermiński, J.T., and Lawrence, J.B. (2020). Silencing Trisomy 21 with XIST in Neural Stem Cells Promotes Neuronal Differentiation. Dev. Cell https://doi.org/10.1016/j.devcel.2019.12.015.

De, P., Peng, Q., Traktuev, D.O., Li, W., Yoder, M.C., March, K.L., and Durden, D.L. (2009). Expression of RAC2 in endothelial cells is required for the postnatal neovascular response. Exp. Cell Res. 315, 248–263. https://doi.org/10.1016/j.yexcr.2008.10.003.

DeCicco-Skinner, K.L., Henry, G.H., Cataisson, C., Tabib, T., Gwilliam, J.C., Watson, N.J., Bullwinkle, E.M., Falkenburg, L., O’Neill, R.C., Morin, A., et al. (2014). Endothelial cell tube formation assay for the in vitro study of angiogenesis. J. Vis. Exp. e51312. https://doi.org/10.3791/51312.

Do, L.H., Mobley, W.C., and Singhal, N. (2015). Questioned validity of Gene Expression Dysregulated Domains in Down’s Syndrome. F1000Res. 4, 269. https://doi.org/10.12688/f1000research.6735.1.

Drachman, D.A. (2014). The amyloid hypothesis, time to move on: Amyloid is the downstream result, not cause, of Alzheimer’s disease. Alzheimers. Dement. 10, 372–380. https://doi.org/10.1016/j.jalz.2013.11.003.

Drachman, D.A., Smith, T.W., Alkamachi, B., and Kane, K. (2017). Microvascular changes in Down syndrome with Alzheimer’s-type pathology: Insights into a potential vascular mechanism for Down syndrome and Alzheimer’s disease. Alzheimers. Dement. 13, 1389–1396. https://doi.org/10.1016/j.jalz.2017.05.003.

Efroni, S., Duttagupta, R., Cheng, J., Dehghani, H., Hoeppner, D.J., Dash, C., Bazett-Jones, D.P., Le Grice, S., McKay, R.D.G., Buetow, K.H., et al. (2008). Global transcription in pluripotent embryonic stem cells. Cell Stem Cell 2, 437–447. https://doi.org/10.1016/j.stem.2008.03.021.

Galambos, C., Minic, A.D., Bush, D., Nguyen, D., Dodson, B., Seedorf, G., and Abman, S.H. (2016). Increased Lung Expression of Anti-Angiogenic Factors in Down Syndrome: Potential Role in Abnormal Lung Vascular Growth and the Risk for Pulmonary Hypertension. PLoS One 11, e0159005. https://doi.org/10.1371/journal.pone.0159005.

Gel, B., and Serra, E. (2017). karyoploteR: an R/Bioconductor package to plot customizable genomes displaying arbitrary data. Bioinformatics 33, 3088–3090. https://doi.org/10.1093/bioinformatics/btx346.

Gene Ontology Consortium (2021). The Gene Ontology resource: enriching a GOld mine. Nucleic Acids Res. 49, D325–D334. https://doi.org/10.1093/nar/gkaa1113.

Gerhardt, H. (2013). VEGF and Endothelial Guidance in Angiogenic Sprouting (Landes Bioscience).

Golden, J.A., and Hyman, B.T. (1994). Development of the superior temporal neocortex is anomalous in trisomy 21. J. Neuropathol. Exp. Neurol. 53, 513–520. https://doi.org/10.1097/00005072-199409000-00011.

Goncharov, N.V., Nadeev, A.D., Jenkins, R.O., and Avdonin, P.V. (2017). Markers and Biomarkers of Endothelium: When Something Is Rotten in the State. Oxid. Med. Cell. Longev. 2017, 9759735. https://doi.org/10.1155/2017/9759735.

Gonzales, P.K., Roberts, C.M., Fonte, V., Jacobsen, C., Stein, G.H., and Link, C.D. (2018). Transcriptome analysis of genetically matched human induced pluripotent stem cells disomic or trisomic for chromosome 21. PLoS One 13, e0194581. https://doi.org/10.1371/journal.pone.0194581.

Hall, L.L., Byron, M., Butler, J., Becker, K.A., Nelson, A., Amit, M., Itskovitz-Eldor, J., Stein, J., Stein, G., Ware, C., et al. (2008). X-inactivation reveals epigenetic anomalies in most hESC but identifies sublines that initiate as expected. J. Cell. Physiol. 216, 445–452. https://doi.org/10.1002/jcp.21411.

Hasle, H., Friedman, J.M., Olsen, J.H., and Rasmussen, S.A. (2016). Low risk of solid tumors in persons with Down syndrome. Genet. Med. 18, 1151–1157. https://doi.org/10.1038/gim.2016.23.

Hellström, M., Phng, L.-K., and Gerhardt, H. (2007). VEGF and Notch signaling: the yin and yang of angiogenic sprouting. Cell Adh. Migr. 1, 133–136. https://doi.org/10.4161/cam.1.3.4978.

Helman, A.M., Siever, M., McCarty, K.L., Lott, I.T., Doran, E., Abner, E.L., Schmitt, F.A., and Head, E. (2019). Microbleeds and Cerebral Amyloid Angiopathy in the Brains of People with Down Syndrome with Alzheimer’s Disease. J. Alzheimers. Dis. 67, 103–112. https://doi.org/10.3233/JAD-180589.

Hussein, S.M.I., Elbaz, J., and Nagy, A.A. (2013). Genome damage in induced pluripotent stem cells: assessing the mechanisms and their consequences. Bioessays 35, 152–162. https://doi.org/10.1002/bies.201200114.

Jiang, J., Jing, Y., Cost, G.J., Chiang, J.-C., Kolpa, H.J., Cotton, A.M., Carone, D.M., Carone, B.R., Shivak, D. a., Guschin, D.Y., et al. (2013). Translating dosage compensation to trisomy 21. Nature 500, 296–300. https://doi.org/10.1038/nature12394.

Karlsen, A.S., and Pakkenberg, B. (2011). Total numbers of neurons and glial cells in cortex and basal ganglia of aged brains with Down syndrome--a stereological study. Cereb. Cortex 21, 2519–2524. https://doi.org/10.1093/cercor/bhr033.

Kim, D., Paggi, J.M., Park, C., Bennett, C., and Salzberg, S.L. (2019). Graph-based genome alignment and genotyping with HISAT2 and HISAT-genotype. Nat. Biotechnol. 37, 907–915. https://doi.org/10.1038/s41587-019-0201-4.

Kim, J., Han, D., Byun, S.-H., Kwon, M., Cho, J.Y., Pleasure, S.J., and Yoon, K. (2018). Ttyh1 regulates embryonic neural stem cell properties by enhancing the Notch signaling pathway. EMBO Rep. 19. https://doi.org/10.15252/embr.201745472.

Kim, K., Doi, A., Wen, B., Ng, K., Zhao, R., Cahan, P., Kim, J., Aryee, M.J., Ji, H., Ehrlich, L.I.R., et al. (2010). Epigenetic memory in induced pluripotent stem cells. Nature 467, 285–290. https://doi.org/10.1038/nature09342.

Kitsukawa, T., Shimizu, M., Sanbo, M., Hirata, T., Taniguchi, M., Bekku, Y., Yagi, T., and Fujisawa, H. (1997). Neuropilin-semaphorin III/D-mediated chemorepulsive signals play a crucial role in peripheral nerve projection in mice. Neuron 19, 995–1005. https://doi.org/10.1016/s0896-6273(00)80392-x.

Kobayashi, H., and Kikyo, N. (2015). Epigenetic regulation of open chromatin in pluripotent stem cells. Transl. Res. 165, 18–27. https://doi.org/10.1016/j.trsl.2014.03.004.

Kohlmaier, A., Savarese, F., Lachner, M., Martens, J., Jenuwein, T., and Wutz, A. (2004). A chromosomal memory triggered by Xist regulates histone methylation in X inactivation. PLoS Biol. 2, E171. https://doi.org/10.1371/journal.pbio.0020171.

Kolde, R. (2019). pheatmap: Pretty Heatmaps.

Koyanagi-Aoi, M., Ohnuki, M., Takahashi, K., Okita, K., Noma, H., Sawamura, Y., Teramoto, I., Narita, M., Sato, Y., Ichisaka, T., et al. (2013). Differentiation-defective phenotypes revealed by large-scale analyses of human pluripotent stem cells. Proc. Natl. Acad. Sci. U. S. A. 110, 20569–20574. https://doi.org/10.1073/pnas.1319061110.

Krueger, J., Liu, D., Scholz, K., Zimmer, A., Shi, Y., Klein, C., Siekmann, A., Schulte-Merker, S., Cudmore, M., Ahmed, A., et al. (2011). Flt1 acts as a negative regulator of tip cell formation and branching morphogenesis in the zebrafish embryo. Development 138, 2111–2120. https://doi.org/10.1242/dev.063933.

Larrayoz, I.M., Ferrero, H., Martisova, E., Gil-Bea, F.J., Ramírez, M.J., and Martínez, A. (2017). Adrenomedullin Contributes to Age-Related Memory Loss in Mice and Is Elevated in Aging Human Brains. Front. Mol. Neurosci. 10, 384. https://doi.org/10.3389/fnmol.2017.00384.

Laurent, L.C., Ulitsky, I., Slavin, I., Tran, H., Schork, A., Morey, R., Lynch, C., Harness, J.V., Lee, S., Barrero, M.J., et al. (2011). Dynamic changes in the copy number of pluripotency and cell proliferation genes in human ESCs and iPSCs during reprogramming and time in culture. Cell Stem Cell 8, 106–118. https://doi.org/10.1016/j.stem.2010.12.003.

Leek, J.T., Johnson, W.E., Parker, H.S., Fertig, E.J., Jaffe, A.E., Zhang, Y., Storey, J.D., and Torres, L.C. (2020). sva: Surrogate Variable Analysis.

Lian, X., Bao, X., Al-Ahmad, A., Liu, J., Wu, Y., Dong, W., Dunn, K.K., Shusta, E.V., and Palecek, S.P. (2014). Efficient differentiation of human pluripotent stem cells to endothelial progenitors via small-molecule activation of WNT signaling. Stem Cell Reports 3, 804–816. https://doi.org/10.1016/j.stemcr.2014.09.005.

Liang, G., and Zhang, Y. (2013). Genetic and epigenetic variations in iPSCs: potential causes and implications for application. Cell Stem Cell 13, 149–159. https://doi.org/10.1016/j.stem.2013.07.001.

Liao, Y., Smyth, G.K., and Shi, W. (2014). featureCounts: an efficient general purpose program for assigning sequence reads to genomic features. Bioinformatics 30, 923–930. https://doi.org/10.1093/bioinformatics/btt656.

Llurba, E., Sánchez, O., Ferrer, Q., Nicolaides, K.H., Ruíz, A., Domínguez, C., Sánchez-de-Toledo, J., García-García, B., Soro, G., Arévalo, S., et al. (2014). Maternal and foetal angiogenic imbalance in congenital heart defects. Eur. Heart J. 35, 701–707. https://doi.org/10.1093/eurheartj/eht389.

Loda, A., and Heard, E. (2019). Xist RNA in action: Past, present, and future. PLoS Genet. 15, e1008333. https://doi.org/10.1371/journal.pgen.1008333.

Mayshar, Y., Ben-David, U., Lavon, N., Biancotti, J.-C., Yakir, B., Clark, A.T., Plath, K., Lowry, W.E., and Benvenisty, N. (2010). Identification and classification of chromosomal aberrations in human induced pluripotent stem cells. Cell Stem Cell 7, 521–531. https://doi.org/10.1016/j.stem.2010.07.017.

McCarthy, D.J., Chen, Y., and Smyth, G.K. (2012). Differential expression analysis of multifactor RNA-Seq experiments with respect to biological variation. Nucleic Acids Res. 40, 4288–4297. https://doi.org/10.1093/nar/gks042.

Michael, I.P., Orebrand, M., Lima, M., Pereira, B., Volpert, O., Quaggin, S.E., and Jeansson, M. (2017). Angiopoietin-1 deficiency increases tumor metastasis in mice. BMC Cancer 17, 539. https://doi.org/10.1186/s12885-017-3531-y.

Minami, T., Horiuchi, K., Miura, M., Abid, M.R., Takabe, W., Noguchi, N., Kohro, T., Ge, X., Aburatani, H., Hamakubo, T., et al. (2004). Vascular endothelial growth factor- and thrombin-induced termination factor, Down syndrome critical region-1, attenuates endothelial cell proliferation and angiogenesis. J. Biol. Chem. 279, 50537–50554. https://doi.org/10.1074/jbc.M406454200.

Mircher, C., Cieuta-Walti, C., Marey, I., Rebillat, A.-S., Cretu, L., Milenko, E., Conte, M., Sturtz, F., Rethore, M.-O., and Ravel, A. (2017). Acute Regression in Young People with Down Syndrome. Brain Sci 7. https://doi.org/10.3390/brainsci7060057.

Morris, J.K., Wald, N.J., and Watt, H.C. (1999). Fetal loss in Down syndrome pregnancies. Prenat. Diagn. 19, 142–145. https://doi.org/10.1002/(SICI)1097-0223(199902)19:2<142::AID-PD486>3.0.CO;2-7.

Neufeld, G., and Kessler, O. (2008). The semaphorins: versatile regulators of tumour progression and tumour angiogenesis. Nat. Rev. Cancer 8, 632–645. https://doi.org/10.1038/nrc2404.

Ochsenbein, A.M., Karaman, S., Proulx, S.T., Berchtold, M., Jurisic, G., Stoeckli, E.T., and Detmar, M. (2016). Endothelial cell-derived semaphorin 3A inhibits filopodia formation by blood vascular tip cells. Development 143, 589–594. https://doi.org/10.1242/dev.127670.

Oh, H., Takagi, H., Otani, A., Koyama, S., Kemmochi, S., Uemura, A., and Honda, Y. (2002). Selective induction of neuropilin-1 by vascular endothelial growth factor (VEGF): a mechanism contributing to VEGF-induced angiogenesis. Proc. Natl. Acad. Sci. U. S. A. 99, 383–388. https://doi.org/10.1073/pnas.012074399.

Ouellette, J., Toussay, X., Comin, C.H., Costa, L. da F., Ho, M., Lacalle-Aurioles, M., Freitas-Andrade, M., Liu, Q.Y., Leclerc, S., Pan, Y., et al. (2020). Vascular contributions to 16p11.2 deletion autism syndrome modeled in mice. Nat. Neurosci. 23, 1090–1101. https://doi.org/10.1038/s41593-020-0663-1.

Park, I.-H., Arora, N., Huo, H., Maherali, N., Ahfeldt, T., Shimamura, A., Lensch, M.W., Cowan, C., Hochedlinger, K., and Daley, G.Q. (2008). Disease-specific induced pluripotent stem cells. Cell 134, 877–886. https://doi.org/10.1016/j.cell.2008.07.041.

Patsch, C., Challet-Meylan, L., Thoma, E.C., Urich, E., Heckel, T., O’Sullivan, J.F., Grainger, S.J., Kapp, F.G., Sun, L., Christensen, K., et al. (2015). Generation of vascular endothelial and smooth muscle cells from human pluripotent stem cells. Nat. Cell Biol. 17, 994–1003. https://doi.org/10.1038/ncb3205.

Phng, L.-K., and Gerhardt, H. (2009). Angiogenesis: a team effort coordinated by notch. Dev. Cell 16, 196– 208. https://doi.org/10.1016/j.devcel.2009.01.015.

Pitter, B., Werner, A.-C., and Montanez, E. (2018). Parvins Are Required for Endothelial Cell-Cell Junctions and Cell Polarity During Embryonic Blood Vessel Formation. Arterioscler. Thromb. Vasc. Biol. 38, 1147–1158. https://doi.org/10.1161/ATVBAHA.118.310840.

Plath, K., Fang, J., Mlynarczyk-Evans, S.K., Cao, R., Worringer, K.A., Wang, H., de la Cruz, C.C., Otte, A.P., Panning, B., and Zhang, Y. (2003). Role of histone H3 lysine 27 methylation in X inactivation. Science 300, 131–135. https://doi.org/10.1126/science.1084274.

Postovit, L.-M., Costa, F.F., Bischof, J.M., Seftor, E.A., Wen, B., Seftor, R.E.B., Feinberg, A.P., Soares, M.B., and Hendrix, M.J.C. (2007). The commonality of plasticity underlying multipotent tumor cells and embryonic stem cells. J. Cell. Biochem. 101, 908–917. https://doi.org/10.1002/jcb.21227.

R Core Team (2021). R: A Language and Environment for Statistical Computing.

Ritchie, M.E., Phipson, B., Wu, D., Hu, Y., Law, C.W., Shi, W., and Smyth, G.K. (2015). limma powers differential expression analyses for RNA-sequencing and microarray studies. Nucleic Acids Res. 43, e47. https://doi.org/10.1093/nar/gkv007.

Robinson, M.D., McCarthy, D.J., and Smyth, G.K. (2010). edgeR: a Bioconductor package for differential expression analysis of digital gene expression data. Bioinformatics 26, 139–140. https://doi.org/10.1093/bioinformatics/btp616.

Roy, A., Cowan, G., Mead, A.J., Filippi, S., Bohn, G., Chaidos, A., Tunstall, O., Chan, J.K.Y., Choolani, M., Bennett, P., et al. (2012). Perturbation of fetal liver hematopoietic stem and progenitor cell development by trisomy 21. Proc. Natl. Acad. Sci. U. S. A. 109, 17579–17584. https://doi.org/10.1073/pnas.1211405109.

Sagare, A.P., Bell, R.D., and Zlokovic, B.V. (2013). Neurovascular defects and faulty amyloid-β vascular clearance in Alzheimer’s disease. J. Alzheimers. Dis.33 Suppl 1, S87–S100. https://doi.org/10.3233/JAD-2012-129037.

Sahakyan, A., Yang, Y., and Plath, K. (2018). The Role of Xist in X-Chromosome Dosage Compensation. Trends Cell Biol. https://doi.org/10.1016/j.tcb.2018.05.005.

Schindelin, J., Arganda-Carreras, I., Frise, E., Kaynig, V., Longair, M., Pietzsch, T., Preibisch, S., Rueden, C., Saalfeld, S., Schmid, B., et al. (2012). Fiji: an open-source platform for biological-image analysis. Nat. Methods 9, 676–682. https://doi.org/10.1038/nmeth.2019.

Schindelin, J., Rueden, C.T., Hiner, M.C., and Eliceiri, K.W. (2015). The ImageJ ecosystem: An open platform for biomedical image analysis. Mol. Reprod. Dev. 82, 518–529. https://doi.org/10.1002/mrd.22489.

Seppinen, L., and Pihlajaniemi, T. (2011). The multiple functions of collagen XVIII in development and disease. Matrix Biol. 30, 83–92. https://doi.org/10.1016/j.matbio.2010.11.001.

Shen, Q., Goderie, S.K., Jin, L., Karanth, N., Sun, Y., Abramova, N., Vincent, P., Pumiglia, K., and Temple, S. (2004). Endothelial cells stimulate self-renewal and expand neurogenesis of neural stem cells. Science 304, 1338–1340. https://doi.org/10.1126/science.1095505.

Smith, K.P., Byron, M., Clemson, C.M., and Lawrence, J.B. (2004). Ubiquitinated proteins including uH2A on the human and mouse inactive X chromosome: enrichment in gene rich bands. Chromosoma 113, 324–335. https://doi.org/10.1007/s00412-004-0325-1.

Soldner, F., and Jaenisch, R. (2012). Medicine. iPSC disease modeling. Science 338, 1155–1156. https://doi.org/10.1126/science.1227682.

Startin, C.M., D’Souza, H., Ball, G., Hamburg, S., Hithersay, R., Hughes, K.M.O., Massand, E., Karmiloff-Smith, A., Thomas, M.S.C., LonDownS Consortium, et al. (2020). Health comorbidities and cognitive abilities across the lifespan in Down syndrome. J. Neurodev. Disord. 12, 4. https://doi.org/10.1186/s11689-019-9306-9.

Sullivan, K.D., Lewis, H.C., Hill, A.A., Pandey, A., Jackson, L.P., Cabral, J.M., Smith, K.P., Liggett, L.A., Gomez, E.B., Galbraith, M.D., et al. (2016). Trisomy 21 consistently activates the interferon response. Elife 5. https://doi.org/10.7554/eLife.16220.

Toshner, M., Dunmore, B.J., McKinney, E.F., Southwood, M., Caruso, P., Upton, P.D., Waters, J.P., Ormiston, M.L., Skepper, J.N., Nash, G., et al. (2014). Transcript analysis reveals a specific HOX signature associated with positional identity of human endothelial cells. PLoS One 9, e91334. https://doi.org/10.1371/journal.pone.0091334.

Trevino, C.E., Holleman, A.M., Corbitt, H., Maslen, C.L., Rosser, T.C., Cutler, D.J., Johnston, H.R., Rambo-Martin, B.L., Oberoi, J., Dooley, K.J., et al. (2020). Identifying genetic factors that contribute to the increased risk of congenital heart defects in infants with Down syndrome. Sci. Rep. 10, 18051. https://doi.org/10.1038/s41598-020-74650-4.

Tunstall-Pedoe, O., Roy, A., Karadimitris, A., de la Fuente, J., Fisk, N.M., Bennett, P., Norton, A., Vyas, P., and Roberts, I. (2008). Abnormalities in the myeloid progenitor compartment in Down syndrome fetal liver precede acquisition of GATA1 mutations. Blood 112, 4507–4511. https://doi.org/10.1182/blood-2008-04-152967.

VanOudenhove, J.J., Medina, R., Ghule, P.N., Lian, J.B., Stein, J.L., Zaidi, S.K., and Stein, G.S. (2016). Transient RUNX1 Expression during Early Mesendodermal Differentiation of hESCs Promotes Epithelial to Mesenchymal Transition through TGFB2 Signaling. Stem Cell Reports 7, 884–896. https://doi.org/10.1016/j.stemcr.2016.09.006.

Wang, C., Liu, H., Yang, M., Bai, Y., Ren, H., Zou, Y., Yao, Z., Zhang, B., and Li, Y. (2020). RNA-Seq Based Transcriptome Analysis of Endothelial Differentiation of Bone Marrow Mesenchymal Stem Cells. Eur. J. Vasc. Endovasc. Surg. 59, 834–842. https://doi.org/10.1016/j.ejvs.2019.11.003.

Wickham, H. (2016). ggplot2: Elegant Graphics for Data Analysis (Springer International Publishing).

Wiseman, F.K., Al-Janabi, T., Hardy, J., Karmiloff-Smith, A., Nizetic, D., Tybulewicz, V.L.J., Fisher, E.M.C., and Strydom, A. (2015). A genetic cause of Alzheimer disease: mechanistic insights from Down syndrome. Nature Publishing Group 16, 564–574. https://doi.org/10.1038/nrn3983.

Yurugi-Kobayashi, T., Itoh, H., Schroeder, T., Nakano, A., Narazaki, G., Kita, F., Yanagi, K., Hiraoka-Kanie, M., Inoue, E., Ara, T., et al. (2006). Adrenomedullin/cyclic AMP pathway induces Notch activation and differentiation of arterial endothelial cells from vascular progenitors. Arterioscler. Thromb. Vasc. Biol. 26, 1977– 1984. https://doi.org/10.1161/01.ATV.0000234978.10658.41.

Zorick, T.S., Mustacchi, Z., Bando, S.Y., Zatz, M., Moreira-Filho, C.A., Olsen, B., and Passos-Bueno, M.R. (2001). High serum endostatin levels in Down syndrome: implications for improved treatment and prevention of solid tumours. Eur. J. Hum. Genet. 9, 811–814. https://doi.org/10.1038/sj.ejhg.5200721.

