## Supplemental Material for "Chromosome silencing *in vitro* reveals trisomy 21 causes cell-autonomous deficits in angiogenesis and early dysregulation in Notch signaling"

**SUPPLEMENTAL MATERIALS**

**Figure S1: *DYRK1A* expression is not disrupted by XIST insertion**
**Figure S2: Gene expression patterns seen in DS iPSCs.**
**Figure S3: Characterization of XIST- and XIST+ endothelial cells.**
**Figure S4: Isogenic Trisomy/Disomy comparison after 1 hours.**
**Figure S5: Gene expression analysis of iPSCs and ECs.**

**Table S1: DEGs of iPSCs (XIST+/XIST-)**
**Table S2: DEGs of iPSC-derived ECs (XIST+/XIST-)**

A

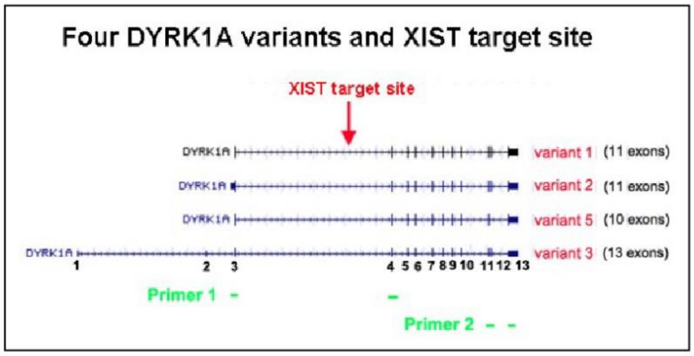

B

| Primer sets | product size | amplified variants |
|---|---|---|
| set 1 | 202 bp | variants 1,2,3,5 |
| set 2 | 324 bp | variant 5 |
| | 449 bp | variants 1,2 |
| | 580 bp | variant 3 |

C

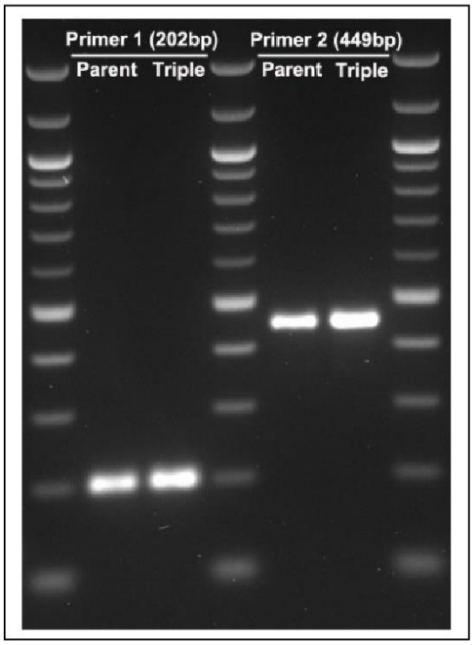

**Figure S1: *DYRK1A* expression is not disrupted by XIST insertion**

(A) *DYRK1A* splicing isoforms with differing 5'UTR and 3' coding regions. The *XIST* transgene (red) was inserted into the intron of variants 1, 2, 5, or intron 3 of variant 3 shown here. Primers were designed to detect for *DYRK1A* expression (green).

(B) Anticipated product sizes for each variant.

(C) RT-PCR results shows the first set of primers generates a 202 bp band, and the second set of primers generates only one 449 bp of single band in both parental and triple target lines (*XIST* inserted in all three alleles of *DYRK1A*). Sequencing results confirm the 202 bp product from the first set of primers in both lines is the sequence spanning exon 1 and exon 2 of variants 1, 2, 5, or spanning exon 3 and exon 4 of variant 3.

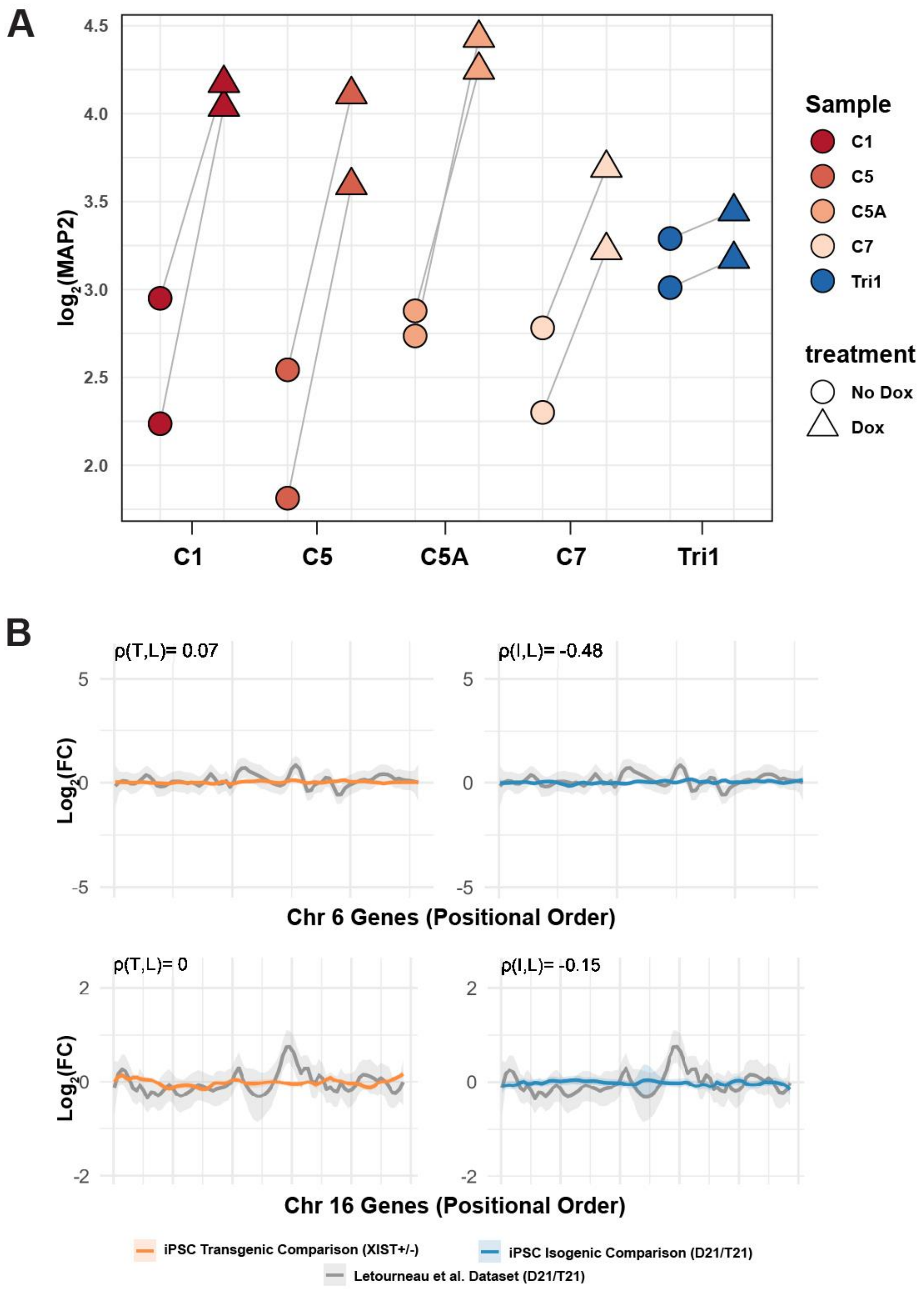

**Figure S2: Gene expression patterns seen in DS iPSCs.**

(A) MAP2 expression of all cell lines in counts per million (CPM). Non-dox treated cells are in dark gray and dox treated cells are in light gray.

(B) Representative local regression plots of chr 6 and chr 16. Letourneau et al. dataset (gray) is plotted against our transgenic or isogenic comparison. Pearson correlation (Rho) is reported above each comparison. P > 0.05.

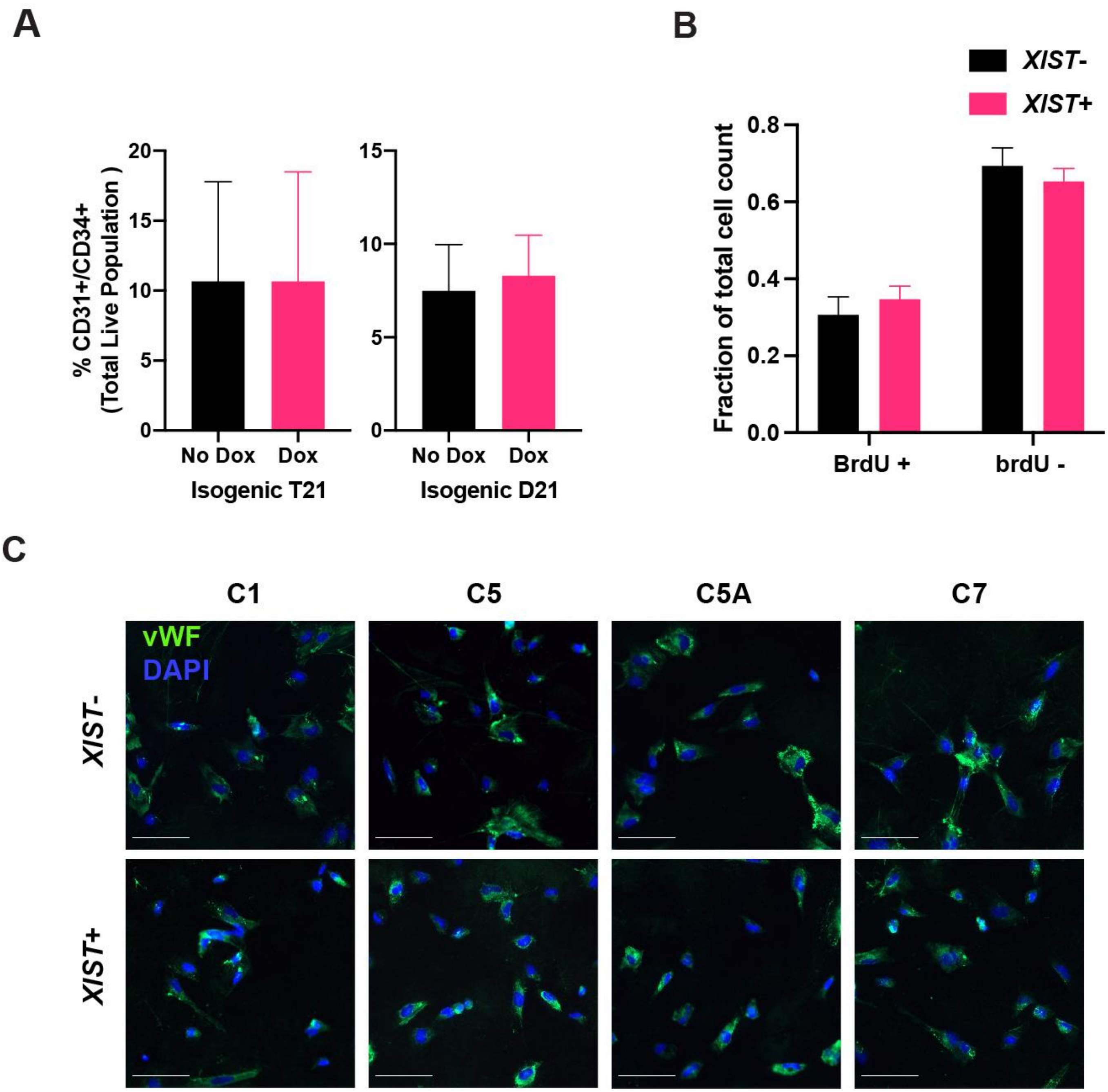

**Figure S3: Characterization of XIST- and XIST+ endothelial cells.**

(A) Quantification of CD31+CD34+ cells in one isogenic trisomic (T21) and disomic lines (D21) with *TET3G* but lacking the *XIST* transgene. Experiment was conducted in parallel with the transgenic lines and repeated three times (mean ± SD).

(B) Representative immunofluorescence images of each condition for vWF (endothelial cell marker). Scale bar = 100 um.

(C) After endothelial progenitor cell enrichment and expansion (day 10), cells were incubated with BrdU for 2 hours. Quantification of BrdU positive and negative cells (400-500 observations per condition; n = 4, mean ± SD; paired t- test p-value = 0.02).

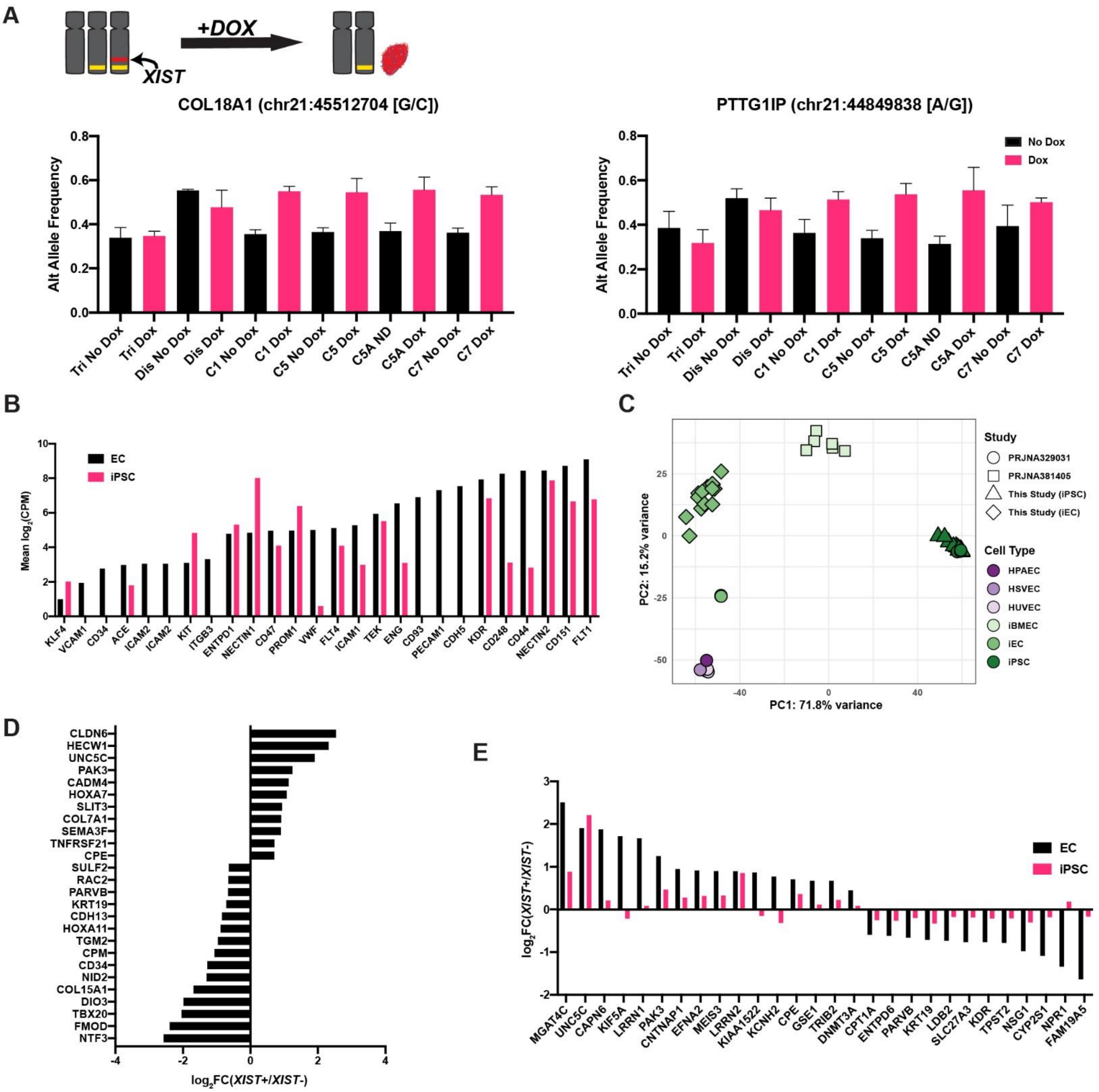

**Figure S4: Isogenic Trisomy/Disomy comparison after 1 hours.**

(A) Representative images of tube formation after one hour of incubation in the dox control lines (Tri1 and Dis1) run in parallel with the transgenic lines seen in Figure II-5B and repeated three times. Scale bar = 1000 um.

(B) The two panels on the right are quantification of tube formation from images represented in (A) using the dox control lines (mean ± SD). The third (left) panel is the comparison between individual features detected by Angiogenesis Analyzer in one isogenic T21 (Tri1) and D21 (Dis1) sample after one hour of incubation (mean ± SD).

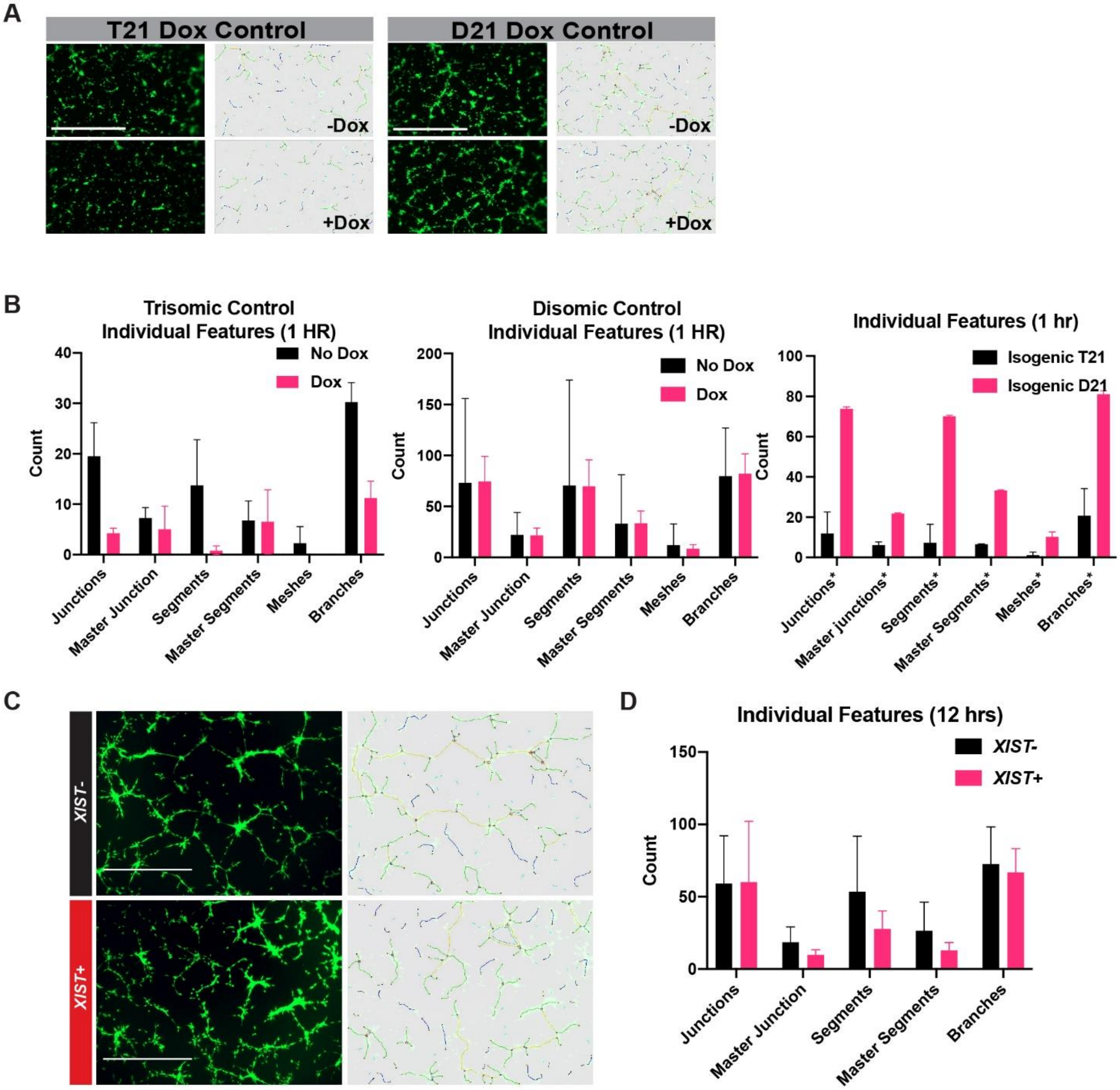

**Figure S5: Gene expression analysis of iPSCs and ECs.**

(A) Illustration of allele frequency changes due to *XIST*-mediated silencing (the allele on the extra chr21 is depicted in yellow and *XIST* locus in red) and two representative SNPs from the endothelial data set after treatment of dox. Each graph shows the gene symbol, its chromosomal position, and reference/alternate alleles. Both genes are ~8 Mb away from the *DYRK1A* locus.

(B) Mean log2 (CPM) expression of common endothelial cell markers are plotted for both EC (black) and iPSC (pink) datasets.

(C) Principal component analysis of the endothelial cells generated in this study against publicly available RNA-seq datasets from iPSC derived endothelial cells (iEC and iBMEC) and primary endothelial cells (HPAEC, HSVEC, HUVEC). iECs generated from this study clustered well with other ECs across principal component 1.

(D) The logFC of DEGs relating to cell migration, adhesion, and extracellular matrix (FDR < 0.05).

(E) Genes differentially expressed in both iPSC and EC datasets after silencing the extra chr21 (FDR < 0.05).

**Table S1. iPSC DEGs**

| geneid | chr | start | end | strand | length | description | logFC | logCPM | F | PValue | FDR |
|---|---|---|---|---|---|---|---|---|---|---|---|
| RBPJ | 4 | 2.6E+07 | 2.6E+07 | + | 9253 | recombination | -0.4706 | 7.54872 | 226.921 | 4.06E-22 | 1.94E-18 |
| FZD2 | 17 | 4.5E+07 | 4.5E+07 | + | 2112 | frizzled class re | 0.81681 | 5.33701 | 216.713 | 1.22E-21 | 4.37E-18 |
| AMOTL1 | 11 | 9.5E+07 | 9.5E+07 | + | 10030 | angiomotin like | 0.60021 | 6.14127 | 214.561 | 1.54E-21 | 4.42E-18 |
| CHD7 | 8 | 6.1E+07 | 6.1E+07 | + | 17020 | chromodomain | 0.43682 | 7.5862 | 212.16 | 2.01E-21 | 4.81E-18 |
| SOD1 | 21 | 3.2E+07 | 3.2E+07 | + | 2019 | superoxide dis | -0.4185 | 7.68841 | 199.559 | 8.44E-21 | 1.73E-17 |
| UGP2 | 2 | 6.4E+07 | 6.4E+07 | + | 6501 | UDP-glucose p | -0.3599 | 8.97171 | 194.798 | 2.10E-20 | 3.35E-17 |
| CXADR | 21 | 1.8E+07 | 1.8E+07 | + | 6021 | CXADR, Ig-like | -0.4221 | 7.9525 | 200.544 | 2.76E-20 | 3.97E-17 |
| MAP2 | 2 | 2.1E+08 | 2.1E+08 | + | 11559 | microtubule as | 1.41947 | 3.43175 | 182.806 | 8.88E-20 | 1.16E-16 |
| TENM3 | 4 | 1.8E+08 | 1.8E+08 | + | 12010 | teneurin trans | 0.42888 | 7.51095 | 180.494 | 3.98E-19 | 4.40E-16 |
| CSTB | 21 | 4.4E+07 | 4.4E+07 | - | 3935 | cystatin B [Sou | -0.4524 | 6.72424 | 164.574 | 6.79E-19 | 6.50E-16 |
| HMGN1 | 21 | 3.9E+07 | 3.9E+07 | - | 5042 | high mobility g | -0.3514 | 8.92199 | 174.984 | 1.47E-18 | 1.24E-15 |
| TMEM123 | 11 | 1E+08 | 1E+08 | - | 4442 | transmembran | 0.50183 | 6.35389 | 156.506 | 2.06E-18 | 1.64E-15 |
| PDXK | 21 | 4.4E+07 | 4.4E+07 | + | 11912 | pyridoxal kinas | -0.4112 | 7.08188 | 154.034 | 2.91E-18 | 2.20E-15 |
| GPC4 | X | 1.3E+08 | 1.3E+08 | - | 4959 | glypican 4 [Sou | 0.34639 | 8.99966 | 170.797 | 4.42E-18 | 3.18E-15 |
| STOX2 | 4 | 1.8E+08 | 1.8E+08 | + | 11365 | storkhead box | 0.6507 | 5.46301 | 150.16 | 5.07E-18 | 3.47E-15 |
| NOTCH3 | 19 | 1.5E+07 | 1.5E+07 | - | 9408 | notch 3 [Sourc | -0.3519 | 8.02246 | 147.67 | 7.28E-18 | 4.75E-15 |
| GART | CHR21_4_ | 3.4E+07 | 3.4E+07 | - | 7751 | phosphoribosy | -0.3079 | 8.69315 | 144.508 | 1.16E-17 | 7.24E-15 |
| SIPA1L2 | 1 | 2.3E+08 | 2.3E+08 | - | 7641 | signal induced | 0.96965 | 4.87473 | 198.32 | 1.59E-17 | 9.51E-15 |
| KLHL4 | X | 8.8E+07 | 8.8E+07 | + | 6158 | kelch like famil | 0.75709 | 4.70498 | 138.025 | 3.08E-17 | 1.77E-14 |
| ADM | 11 | 1E+07 | 1E+07 | + | 2325 | adrenomedulli | 0.56654 | 6.14715 | 160.541 | 3.62E-17 | 2.00E-14 |
| USP25 | 21 | 1.6E+07 | 1.6E+07 | + | 7823 | ubiquitin speci | -0.4476 | 6.62012 | 136.649 | 8.47E-17 | 4.51E-14 |
| NSD2 | 4 | 1871393 | 1982207 | + | 19807 | nuclear recept | 0.37047 | 7.50114 | 136.124 | 9.15E-17 | 4.70E-14 |
| AC064802 | 8 | 1.1E+08 | 1.1E+08 | + | 1192 | novel transcrip | -0.4033 | 7.42759 | 148.325 | 1.04E-16 | 5.14E-14 |
| ZIC2 | 13 | 1E+08 | 1E+08 | + | 3290 | Zic family mem | 0.51239 | 7.30196 | 206.912 | 1.16E-16 | 5.57E-14 |
| ARHGEF40 | 14 | 2.1E+07 | 2.1E+07 | + | 8194 | Rho guanine n | 0.43877 | 6.26907 | 125.851 | 2.11E-16 | 9.79E-14 |
| RRP1B | 21 | 4.4E+07 | 4.4E+07 | + | 5750 | ribosomal RNA | -0.33 | 7.84654 | 124.535 | 2.62E-16 | 1.18E-13 |
| NME4 | 16 | 396725 | 410367 | + | 3665 | NME/NM23 nu | 0.33142 | 7.53576 | 123.687 | 3.01E-16 | 1.31E-13 |
| VPS26C | 21 | 3.7E+07 | 3.7E+07 | - | 9301 | VPS26 endoso | -0.4365 | 6.86997 | 137.932 | 6.21E-16 | 2.55E-13 |
| SCAF4 | 21 | 3.2E+07 | 3.2E+07 | - | 5901 | SR-related CTD | -0.3899 | 6.74332 | 116.395 | 1.03E-15 | 4.11E-13 |
| TIAM1 | 21 | 3.1E+07 | 3.2E+07 | - | 9216 | T cell lymphom | -0.4553 | 6.05472 | 116.071 | 1.09E-15 | 4.23E-13 |
| APP | 21 | 2.6E+07 | 2.6E+07 | - | 6316 | amyloid beta p | -0.2367 | 10.7376 | 114.867 | 1.34E-15 | 5.08E-13 |
| CD99 | X | 2691187 | 2741309 | + | 4858 | CD99 molecule | 0.6373 | 5.08 | 114.655 | 1.39E-15 | 5.13E-13 |
| JAG1 | 20 | 1.1E+07 | 1.1E+07 | - | 9298 | jagged 1 [Sourc | -0.5565 | 5.55934 | 113.924 | 1.76E-15 | 6.33E-13 |
| BCAT1 | 12 | 2.5E+07 | 2.5E+07 | - | 11415 | branched chair | 0.28447 | 8.84984 | 116.768 | 1.92E-15 | 6.72E-13 |
| NFASC | 1 | 2E+08 | 2.1E+08 | + | 21271 | neurofascin [So | 0.71114 | 4.60279 | 112.437 | 2.05E-15 | 6.94E-13 |
| XIST | X | 7.4E+07 | 7.4E+07 | - | 25264 | X inactive spec | 9.36311 | 10.0166 | 14533.4 | 2.08E-15 | 6.94E-13 |
| SLC5A3 | 21 | 3.4E+07 | 3.4E+07 | + | 11566 | solute carrier f | -0.3993 | 6.48413 | 111.757 | 2.32E-15 | 7.56E-13 |
| PURO | PURO | 1 | 1353 | - | 1353 | Puromycin sele | -0.4197 | 6.6701 | 119.247 | 2.53E-15 | 8.07E-13 |
| AC007950 | 15 | 6.4E+07 | 6.4E+07 | + | 7764 | ubiquitin speci | 1.14572 | 3.14286 | 110.813 | 2.74E-15 | 8.27E-13 |
| NPFFR2 | 4 | 7.2E+07 | 7.2E+07 | + | 2359 | neuropeptide | 1.93208 | 1.75416 | 110.76 | 2.76E-15 | 8.27E-13 |
| DUSP6 | 12 | 8.9E+07 | 8.9E+07 | - | 3859 | dual specificity | 0.5222 | 7.19754 | 196.741 | 3.13E-15 | 9.19E-13 |

| | | | | | | | | | | |
|---|---|---|---|---|---|---|---|---|---|---|
| H2AFY | 5 | 1.4E+08 | 1.4E+08 | - | 17730 | H2A histone fa | 0.34916 | 7.01158 | 108.368 | 4.24E-15 | 1.22E-12 |
| PCSK9 | 1 | 5.5E+07 | 5.5E+07 | + | 4981 | proprotein cor | -0.434 | 6.31661 | 110.725 | 4.67E-15 | 1.29E-12 |
| GBX2 | 2 | 2.4E+08 | 2.4E+08 | - | 2332 | gastrulation br | 1.22349 | 2.89794 | 106.046 | 6.47E-15 | 1.75E-12 |
| ANOS1 | X | 8528874 | 8732137 | - | 7082 | anosmin 1 [Sou | 0.37363 | 6.66055 | 102.882 | 1.16E-14 | 3.09E-12 |
| PMAIP1 | 18 | 6E+07 | 6E+07 | + | 2235 | phorbol-12-my | -0.3876 | 6.74389 | 107.243 | 1.41E-14 | 3.67E-12 |
| GPRC5B | 16 | 2E+07 | 2E+07 | - | 6654 | G protein-coup | -0.3285 | 7.17134 | 101.391 | 1.54E-14 | 3.94E-12 |
| ATP5PF | 21 | 2.6E+07 | 2.6E+07 | - | 2043 | ATP synthase p | -0.3894 | 6.44538 | 100.061 | 1.97E-14 | 4.89E-12 |
| LITAF | 16 | 1.2E+07 | 1.2E+07 | - | 5060 | lipopolysaccha | -0.5394 | 7.29027 | 202.281 | 2.14E-14 | 5.10E-12 |
| GJA1 | 6 | 1.2E+08 | 1.2E+08 | + | 3413 | gap junction pı | -0.3063 | 9.72672 | 139.395 | 2.16E-14 | 5.10E-12 |
| LSS | SCHR21_5 | 4.6E+07 | 4.6E+07 | - | 7238 | lanosterol synt | -0.2765 | 8.12501 | 98.8924 | 2.47E-14 | 5.72E-12 |
| SLC16A12 | 10 | 8.9E+07 | 9E+07 | - | 4873 | solute carrier f | 1.90946 | 1.77327 | 98.1277 | 2.86E-14 | 6.52E-12 |
| CALB1 | 8 | 9E+07 | 9E+07 | - | 7138 | calbindin 1 [So | 0.78539 | 4.08497 | 97.6135 | 3.15E-14 | 7.08E-12 |
| ITSN1 | 21 | 3.4E+07 | 3.4E+07 | + | 21748 | intersectin 1 [S | -0.2792 | 7.89856 | 96.3565 | 4.02E-14 | 8.90E-12 |
| ODC1 | 2 | 1E+07 | 1E+07 | - | 2791 | ornithine deca | 0.39461 | 6.43994 | 98.7532 | 4.29E-14 | 9.35E-12 |
| NLGN4X | X | 5840637 | 6228867 | - | 7439 | neuroligin 4 X- | 0.31008 | 7.23087 | 94.5014 | 5.79E-14 | 1.24E-11 |
| PLA2G16 | 11 | 6.4E+07 | 6.4E+07 | - | 3116 | phospholipase | -0.4257 | 6.18462 | 97.4122 | 6.31E-14 | 1.33E-11 |
| NFE2L3 | 7 | 2.6E+07 | 2.6E+07 | + | 4303 | nuclear factor, | 0.27625 | 7.75797 | 93.5165 | 7.03E-14 | 1.46E-11 |
| PAM | 5 | 1E+08 | 1E+08 | + | 8846 | peptidylglycine | 0.36131 | 6.57452 | 92.849 | 8.02E-14 | 1.62E-11 |
| BRWD1 | 21 | 3.9E+07 | 3.9E+07 | - | 19794 | bromodomain | -0.3194 | 7.07928 | 92.4307 | 8.72E-14 | 1.72E-11 |
| PTPRZ1 | 7 | 1.2E+08 | 1.2E+08 | + | 10137 | protein tyrosin | 0.24478 | 8.80359 | 92.387 | 8.80E-14 | 1.72E-11 |
| SALL4 | 20 | 5.2E+07 | 5.2E+07 | - | 5773 | spalt like trans | -0.3375 | 7.35606 | 101.786 | 9.11E-14 | 1.75E-11 |
| SON | 21 | 3.4E+07 | 3.4E+07 | + | 12785 | SON DNA bind | -0.2312 | 9.24018 | 90.8392 | 1.20E-13 | 2.27E-11 |
| SAMHD1 | 20 | 3.7E+07 | 3.7E+07 | - | 11261 | SAM and HD d | -0.4329 | 5.9882 | 91.483 | 1.21E-13 | 2.27E-11 |
| KRT8 | 12 | 5.3E+07 | 5.3E+07 | - | 6881 | keratin 8 [Sour | -0.2343 | 9.00786 | 89.8379 | 1.47E-13 | 2.71E-11 |
| TFRC | 3 | 2E+08 | 2E+08 | - | 7794 | transferrin rec | 0.2894 | 7.40055 | 88.9271 | 1.77E-13 | 3.12E-11 |
| PTCH1 | 9 | 9.5E+07 | 9.6E+07 | - | 16591 | patched 1 [Sou | -0.4199 | 6.04571 | 89.5209 | 1.78E-13 | 3.12E-11 |
| FRAS1 | 4 | 7.8E+07 | 7.9E+07 | + | 19207 | Fraser extracel | 0.24861 | 8.54677 | 88.583 | 1.90E-13 | 3.29E-11 |
| KRT19 | 17 | 4.2E+07 | 4.2E+07 | - | 2336 | keratin 19 [Sou | -0.3304 | 7.05267 | 90.4283 | 2.46E-13 | 4.16E-11 |
| ATP5PO | 21 | 3.4E+07 | 3.4E+07 | - | 3170 | ATP synthase p | -0.3889 | 6.28865 | 88.2084 | 2.71E-13 | 4.53E-11 |
| IGFBP2 | 2 | 2.2E+08 | 2.2E+08 | + | 4742 | insulin like gro | 0.29499 | 8.70097 | 108.692 | 3.07E-13 | 5.07E-11 |
| USP11 | X | 4.7E+07 | 4.7E+07 | + | 5269 | ubiquitin speci | -0.3168 | 6.85119 | 85.2257 | 3.81E-13 | 6.22E-11 |
| ITGAV | 2 | 1.9E+08 | 1.9E+08 | + | 8220 | integrin subun | 0.32892 | 6.8099 | 84.3975 | 4.53E-13 | 7.24E-11 |
| PYCR1 | 17 | 8.2E+07 | 8.2E+07 | - | 2886 | pyrroline-5-car | 0.32829 | 6.76684 | 84.3924 | 4.54E-13 | 7.24E-11 |
| ROCK1 | 18 | 2.1E+07 | 2.1E+07 | - | 11075 | Rho associated | -0.3145 | 6.97872 | 84.2842 | 4.64E-13 | 7.33E-11 |
| EMD | X | 1.5E+08 | 1.5E+08 | + | 2020 | emerin [Source | -0.371 | 6.20426 | 84.1902 | 4.73E-13 | 7.39E-11 |
| ESRG | 3 | 5.5E+07 | 5.5E+07 | - | 3140 | embryonic ster | -0.2179 | 9.69157 | 84.065 | 4.86E-13 | 7.51E-11 |
| POLE | 12 | 1.3E+08 | 1.3E+08 | - | 16609 | DNA polymera | 0.30282 | 6.97935 | 83.8632 | 5.07E-13 | 7.65E-11 |
| MIS18A | 21 | 3.2E+07 | 3.2E+07 | - | 1694 | MIS18 kinetocl | -0.4124 | 6.64027 | 105.112 | 5.11E-13 | 7.65E-11 |
| TP53I11 | 11 | 4.5E+07 | 4.5E+07 | - | 7345 | tumor protein | -0.3541 | 6.45384 | 82.9903 | 6.10E-13 | 8.86E-11 |
| SLC37A1 | 21 | 4.2E+07 | 4.3E+07 | + | 6854 | solute carrier f | -0.4773 | 5.39243 | 81.9024 | 7.70E-13 | 1.11E-10 |
| MT-ND3 | MT | 10059 | 10404 | + | 346 | mitochondriall | -0.2146 | 9.37871 | 81.6784 | 8.08E-13 | 1.15E-10 |
| ROBO2 | 3 | 7.6E+07 | 7.8E+07 | + | 16663 | roundabout gu | 1.02085 | 2.98463 | 81.6287 | 8.17E-13 | 1.15E-10 |
| FAM20C | HSCHR7_1 | 196274 | 199037 | + | 5054 | FAM20C, golgi | 0.54709 | 4.99621 | 81.2246 | 8.91E-13 | 1.24E-10 |

| | | | | | | | | | | | |
|---|---|---|---|---|---|---|---|---|---|---|---|
| PYCR2 | 1 | 2.3E+08 | 2.3E+08 | - | 3602 | pyrroline-5-car | 0.3906 | 6.1606 | 82.889 | 9.11E-13 | 1.26E-10 |
| PTPRG | 3 | 6.2E+07 | 6.2E+07 | + | 14625 | protein tyrosin | 0.2581 | 7.83785 | 81.0557 | 9.24E-13 | 1.26E-10 |
| COL5A2 | 2 | 1.9E+08 | 1.9E+08 | - | 7319 | collagen type V | 0.85932 | 6.09956 | 241.423 | 1.12E-12 | 1.51E-10 |
| HSP90AA1 | 14 | 1E+08 | 1E+08 | - | 5248 | heat shock pro | -0.1972 | 10.6463 | 79.3332 | 1.34E-12 | 1.80E-10 |
| CADM2 | 3 | 8.5E+07 | 8.6E+07 | + | 9990 | cell adhesion n | 0.80155 | 3.72001 | 78.638 | 1.57E-12 | 2.08E-10 |
| ROBO1 | 3 | 7.9E+07 | 8E+07 | - | 11059 | roundabout gu | 0.28493 | 7.33794 | 78.134 | 1.80E-12 | 2.37E-10 |
| DST | 6 | 5.6E+07 | 5.7E+07 | - | 47883 | dystonin [Sour | 0.26435 | 8.13828 | 82.1234 | 1.86E-12 | 2.42E-10 |
| ARRB1 | 11 | 7.5E+07 | 7.5E+07 | - | 9888 | arrestin beta 1 | -0.4025 | 6.05613 | 80.1561 | 2.14E-12 | 2.74E-10 |
| ASS1 | 9 | 1.3E+08 | 1.3E+08 | + | 3050 | argininosuccin | 0.43347 | 5.60148 | 77.1656 | 2.17E-12 | 2.76E-10 |
| FARP1 | 13 | 9.8E+07 | 9.8E+07 | + | 36228 | FERM, ARH/Rh | 0.30003 | 6.82176 | 77.0182 | 2.24E-12 | 2.82E-10 |
| NECTIN2 | 19 | 4.5E+07 | 4.5E+07 | + | 3742 | nectin cell adh | -0.2497 | 7.87364 | 76.8986 | 2.30E-12 | 2.87E-10 |
| TMPRSS2 | 21 | 4.1E+07 | 4.2E+07 | - | 4818 | transmembran | -0.7193 | 3.96754 | 75.4481 | 3.18E-12 | 3.94E-10 |
| DYRK1A | 21 | 3.7E+07 | 3.8E+07 | + | 26763 | dual specificity | -0.3463 | 6.27331 | 75.3802 | 3.23E-12 | 3.97E-10 |
| UNC5D | 8 | 3.5E+07 | 3.6E+07 | + | 10798 | unc-5 netrin re | 0.31138 | 6.76535 | 75.1147 | 3.43E-12 | 4.18E-10 |
| ANK3 | 10 | 6E+07 | 6.1E+07 | - | 25759 | ankyrin 3 [Sou | 0.346 | 6.25618 | 74.8468 | 3.64E-12 | 4.40E-10 |
| POU3F1 | 1 | 3.8E+07 | 3.8E+07 | - | 2184 | POU class 3 ho | 0.59284 | 4.68801 | 75.9587 | 4.19E-12 | 5.02E-10 |
| SPATC1L | SCHR21_5 | 4.6E+07 | 4.6E+07 | - | 2333 | spermatogene | -0.3424 | 6.20366 | 73.183 | 5.32E-12 | 6.23E-10 |
| MMP24 | 20 | 3.5E+07 | 3.5E+07 | + | 4376 | matrix metallo | -0.5004 | 5.06937 | 72.7919 | 5.82E-12 | 6.74E-10 |
| PRMT2 | 21 | 4.7E+07 | 4.7E+07 | + | 7603 | protein argini | -0.3135 | 6.80318 | 74.1102 | 6.23E-12 | 7.16E-10 |
| ID3 | 1 | 2.4E+07 | 2.4E+07 | - | 1496 | inhibitor of DN | 0.26994 | 7.23004 | 72.4286 | 6.33E-12 | 7.21E-10 |
| B3GALT5 | 21 | 4E+07 | 4E+07 | + | 14319 | beta-1,3-galac | -0.2994 | 6.76994 | 72.3893 | 6.38E-12 | 7.22E-10 |
| MCM3AP | 21 | 4.6E+07 | 4.6E+07 | - | 9570 | minichromoso | -0.2384 | 7.98386 | 72.1301 | 6.78E-12 | 7.61E-10 |
| NRARP | 9 | 1.4E+08 | 1.4E+08 | - | 2641 | NOTCH regulat | -0.3817 | 5.81029 | 70.9713 | 8.87E-12 | 9.66E-10 |
| TBC1D9B | 5 | 1.8E+08 | 1.8E+08 | - | 11473 | TBC1 domain f | 0.29943 | 6.73294 | 70.9432 | 8.92E-12 | 9.66E-10 |
| COL4A5 | X | 1.1E+08 | 1.1E+08 | + | 11865 | collagen type I | 0.42891 | 5.47221 | 70.9343 | 8.94E-12 | 9.66E-10 |
| C21orf91 | 21 | 1.8E+07 | 1.8E+07 | - | 6228 | chromosome 2 | -0.3747 | 5.88613 | 70.8442 | 9.13E-12 | 9.79E-10 |
| NOVA2 | 19 | 4.6E+07 | 4.6E+07 | - | 9003 | NOVA alternat | -0.4773 | 5.13563 | 70.8144 | 9.20E-12 | 9.79E-10 |
| PSMG1 | 21 | 3.9E+07 | 3.9E+07 | - | 2515 | proteasome as | -0.3686 | 5.96666 | 70.7374 | 9.36E-12 | 9.89E-10 |
| IGFBP5 | 2 | 2.2E+08 | 2.2E+08 | - | 6466 | insulin like gro | 0.94262 | 3.53958 | 81.7976 | 9.61E-12 | 1.01E-09 |
| NNAT | 20 | 3.8E+07 | 3.8E+07 | + | 1329 | neuronatin [So | 0.49341 | 5.29403 | 73.787 | 1.13E-11 | 1.18E-09 |
| EFNB3 | 17 | 7705202 | 7711372 | + | 3216 | ephrin B3 [Sou | -0.5063 | 4.94717 | 69.6684 | 1.20E-11 | 1.24E-09 |
| SLC38A5 | X | 4.8E+07 | 4.8E+07 | - | 4018 | solute carrier f | -0.4515 | 5.72689 | 77.2309 | 1.30E-11 | 1.34E-09 |
| CAMK2D | 4 | 1.1E+08 | 1.1E+08 | - | 7363 | calcium/calmo | 0.55323 | 4.60343 | 68.9025 | 1.44E-11 | 1.46E-09 |
| LPAR6 | 13 | 4.8E+07 | 4.8E+07 | - | 4348 | lysophosphatic | 1.63495 | 1.64112 | 68.8888 | 1.45E-11 | 1.46E-09 |
| GFPT2 | 5 | 1.8E+08 | 1.8E+08 | - | 4721 | glutamine-fruc | 0.24966 | 7.54091 | 68.5621 | 1.56E-11 | 1.57E-09 |
| GABPA | 21 | 2.6E+07 | 2.6E+07 | + | 5429 | GA binding pro | -0.3954 | 5.84052 | 69.1424 | 1.68E-11 | 1.68E-09 |
| MRPL39 | 21 | 2.6E+07 | 2.6E+07 | - | 1199 | mitochondrial | -0.3881 | 5.78021 | 67.9783 | 1.79E-11 | 1.78E-09 |
| NQO1 | 16 | 7E+07 | 7E+07 | - | 2971 | NAD(P)H quinc | 0.45057 | 5.42691 | 68.8229 | 2.15E-11 | 2.12E-09 |
| PGGHG | 11 | 289135 | 296107 | + | 5260 | protein-glucos | 0.4164 | 5.47906 | 66.8463 | 2.35E-11 | 2.30E-09 |
| FGF2 | 4 | 1.2E+08 | 1.2E+08 | + | 6775 | fibroblast grow | -0.2563 | 7.37036 | 66.6114 | 2.49E-11 | 2.42E-09 |
| PTGIS | 20 | 5E+07 | 5E+07 | - | 5570 | prostaglandin | -0.3632 | 6.09006 | 67.9371 | 2.56E-11 | 2.47E-09 |
| WEE1 | 11 | 9573670 | 9593457 | + | 9251 | WEE1 G2 chec | 0.27948 | 6.9697 | 66.3585 | 2.65E-11 | 2.54E-09 |
| LARGE2 | 11 | 4.6E+07 | 4.6E+07 | + | 3151 | LARGE xylosyl- | -0.2898 | 6.82856 | 65.7059 | 3.10E-11 | 2.95E-09 |

| | | | | | | | | | | | |
|---|---|---|---|---|---|---|---|---|---|---|---|
| UNC5B | 10 | 7.1E+07 | 7.1E+07 | + | 6841 | unc-5 netrin re | -0.3022 | 6.49888 | 65.4833 | 3.27E-11 | 3.09E-09 |
| EMX1 | 2 | 7.3E+07 | 7.3E+07 | + | 5516 | empty spiracle | 0.67577 | 3.96245 | 64.825 | 3.84E-11 | 3.59E-09 |
| COBL | 7 | 5.1E+07 | 5.1E+07 | - | 14349 | cordon-bleu W | -0.2619 | 7.14105 | 64.3431 | 4.33E-11 | 4.01E-09 |
| EMP2 | 16 | 1.1E+07 | 1.1E+07 | - | 8652 | epithelial mem | -0.5284 | 4.8719 | 65.8102 | 4.80E-11 | 4.42E-09 |
| FAT1 | 4 | 1.9E+08 | 1.9E+08 | - | 16167 | FAT atypical ca | 0.22038 | 8.13095 | 63.2756 | 5.63E-11 | 5.12E-09 |
| SCOC | 4 | 1.4E+08 | 1.4E+08 | + | 6294 | short coiled-co | -0.4684 | 5.21402 | 64.5133 | 5.90E-11 | 5.33E-09 |
| TTC3 | 21 | 3.7E+07 | 3.7E+07 | + | 15883 | tetratricopepti | -0.368 | 5.80537 | 62.8381 | 6.27E-11 | 5.56E-09 |
| ZDHHC22 | 14 | 7.7E+07 | 7.7E+07 | - | 3862 | zinc finger DHH | -0.2904 | 6.66912 | 60.9752 | 1.00E-10 | 8.77E-09 |
| AES | 19 | 3052910 | 3063107 | - | 5298 | amino-termina | 0.23255 | 7.57478 | 60.3997 | 1.16E-10 | 1.01E-08 |
| MRPS6 | 21 | 3.4E+07 | 3.4E+07 | + | 4273 | mitochondrial | -0.3072 | 6.25717 | 60.061 | 1.26E-10 | 1.09E-08 |
| PODXL | 7 | 1.3E+08 | 1.3E+08 | - | 6893 | podocalyxin lik | -0.1651 | 10.9279 | 59.8972 | 1.32E-10 | 1.13E-08 |
| HMX2 | 10 | 1.2E+08 | 1.2E+08 | + | 1628 | H6 family hom | 1.73749 | 1.21037 | 59.6856 | 1.39E-10 | 1.18E-08 |
| UBE2G2 | 21 | 4.5E+07 | 4.5E+07 | - | 7195 | ubiquitin conju | -0.2417 | 7.29279 | 59.6558 | 1.40E-10 | 1.18E-08 |
| FOXO4 | X | 7.1E+07 | 7.1E+07 | + | 3447 | forkhead box ( | -0.3327 | 5.98679 | 59.5144 | 1.45E-10 | 1.21E-08 |
| TFAP2C | 20 | 5.7E+07 | 5.7E+07 | + | 3001 | transcription fa | -0.7203 | 3.65583 | 59.3129 | 1.53E-10 | 1.27E-08 |
| LRRN2 | 1 | 2E+08 | 2E+08 | - | 5686 | leucine rich re | 0.85487 | 3.41002 | 64.2888 | 1.55E-10 | 1.28E-08 |
| SUMO3 | 21 | 4.5E+07 | 4.5E+07 | - | 2732 | small ubiquitin | -0.2367 | 7.61035 | 59.0507 | 1.63E-10 | 1.34E-08 |
| PRKCB | 16 | 2.4E+07 | 2.4E+07 | + | 11086 | protein kinase | -0.4783 | 4.81519 | 58.2805 | 1.99E-10 | 1.63E-08 |
| PDE9A | 21 | 4.3E+07 | 4.3E+07 | + | 5281 | phosphodieste | -0.3604 | 5.72633 | 58.1239 | 2.07E-10 | 1.67E-08 |
| CACNA2D | 7 | 8.2E+07 | 8.2E+07 | - | 8989 | calcium voltag | 0.36807 | 5.6634 | 58.1195 | 2.08E-10 | 1.67E-08 |
| TMEM132 | 11 | 6.1E+07 | 6.1E+07 | + | 7271 | transmembran | 0.39028 | 5.66098 | 59.5514 | 2.08E-10 | 1.67E-08 |
| TUBB4A | 19 | 6494319 | 6502848 | - | 3220 | tubulin beta 4A | -0.4148 | 5.24719 | 58.0213 | 2.13E-10 | 1.70E-08 |
| CHAF1B | 21 | 3.6E+07 | 3.6E+07 | + | 5104 | chromatin asse | -0.3544 | 5.91838 | 58.4302 | 2.27E-10 | 1.80E-08 |
| CXCL5 | 4 | 7.4E+07 | 7.4E+07 | - | 2436 | C-X-C motif ch | 0.98566 | 5.67293 | 204.118 | 2.42E-10 | 1.90E-08 |
| CGNL1 | 15 | 5.7E+07 | 5.8E+07 | + | 8663 | cingulin like 1 | -0.2438 | 7.25016 | 57.3327 | 2.55E-10 | 1.99E-08 |
| ARHGEF2 | 12 | 5.8E+07 | 5.8E+07 | + | 3503 | Rho guanine n | 1.03358 | 2.46364 | 56.9588 | 2.81E-10 | 2.18E-08 |
| HES1 | 3 | 1.9E+08 | 1.9E+08 | + | 2059 | hes family bHL | -0.6092 | 4.13913 | 56.6266 | 3.22E-10 | 2.46E-08 |
| FLT1 | 13 | 2.8E+07 | 2.8E+07 | - | 12565 | fms related tyr | 0.2659 | 6.7753 | 56.429 | 3.23E-10 | 2.46E-08 |
| MALAT1 | 11 | 6.5E+07 | 6.6E+07 | + | 8829 | metastasis ass | -0.3642 | 7.82299 | 102.98 | 3.27E-10 | 2.47E-08 |
| NAV2 | 11 | 1.9E+07 | 2E+07 | + | 15174 | neuron naviga | -0.286 | 6.35692 | 56.3434 | 3.30E-10 | 2.48E-08 |
| TMEM47 | X | 3.5E+07 | 3.5E+07 | - | 4040 | transmembran | -0.3571 | 5.71388 | 55.8631 | 3.75E-10 | 2.80E-08 |
| HIST1H2A | 6 | 2.6E+07 | 2.6E+07 | + | 1668 | histone cluster | -0.5017 | 4.57467 | 55.5897 | 4.03E-10 | 2.98E-08 |
| PRDM14 | 8 | 7E+07 | 7E+07 | - | 2443 | PR/SET domain | -0.3074 | 6.3468 | 56.2945 | 4.50E-10 | 3.30E-08 |
| CDK14 | 7 | 9E+07 | 9.1E+07 | + | 7017 | cyclin depende | 0.32316 | 6.10452 | 55.3661 | 4.78E-10 | 3.48E-08 |
| RBBP7 | X | 1.7E+07 | 1.7E+07 | - | 7392 | RB binding pro | -0.2551 | 7.17199 | 56.5713 | 4.98E-10 | 3.60E-08 |
| FKBP9 | 7 | 3.3E+07 | 3.3E+07 | + | 5169 | FKBP prolyl iso | 0.65389 | 3.70533 | 54.7179 | 5.08E-10 | 3.65E-08 |
| LTN1 | 21 | 2.9E+07 | 2.9E+07 | - | 8876 | listerin E3 ubiq | -0.2828 | 6.36914 | 54.6503 | 5.17E-10 | 3.70E-08 |
| HAPLN3 | 15 | 8.9E+07 | 8.9E+07 | - | 3524 | hyaluronan an | -0.3026 | 6.15189 | 54.4874 | 5.40E-10 | 3.84E-08 |
| TMEM132 | 12 | 1.3E+08 | 1.3E+08 | - | 7434 | transmembran | 0.50938 | 4.47668 | 54.258 | 5.74E-10 | 4.04E-08 |
| HMCN1 | 1 | 1.9E+08 | 1.9E+08 | + | 18739 | hemicentin 1 [ | 0.48613 | 4.73623 | 53.8077 | 6.71E-10 | 4.68E-08 |
| SEMA3A | 7 | 8.4E+07 | 8.4E+07 | - | 9259 | semaphorin 3A | 0.34232 | 5.7197 | 53.538 | 6.97E-10 | 4.82E-08 |
| CDK16 | X | 4.7E+07 | 4.7E+07 | + | 6531 | cyclin depende | -0.2412 | 7.00232 | 53.5339 | 6.98E-10 | 4.82E-08 |
| LRRC8A | 9 | 1.3E+08 | 1.3E+08 | + | 5230 | leucine rich re | 0.22089 | 7.48176 | 53.2584 | 7.51E-10 | 5.16E-08 |

| | | | | | | | | | | | |
|---|---|---|---|---|---|---|---|---|---|---|---|
| COL4A2 | 13 | 1.1E+08 | 1.1E+08 | + | 19293 | collagen type I | 0.2059 | 8.24248 | 53.5709 | 7.54E-10 | 5.16E-08 |
| FTH1 | 11 | 6.2E+07 | 6.2E+07 | - | 2004 | ferritin heavy c | 0.18444 | 8.62953 | 53.0172 | 8.02E-10 | 5.46E-08 |
| RPS2 | 16 | 1962058 | 1964841 | - | 2388 | ribosomal prot | 0.15326 | 10.9114 | 52.9935 | 8.07E-10 | 5.47E-08 |
| SYP | X | 4.9E+07 | 4.9E+07 | - | 5215 | synaptophysin | -0.7855 | 3.20298 | 51.9364 | 1.08E-09 | 7.23E-08 |
| RRP1 | 21 | 4.4E+07 | 4.4E+07 | + | 4251 | ribosomal RNA | -0.27 | 6.58421 | 51.8853 | 1.09E-09 | 7.30E-08 |
| LINC0096 | 3 | 7.6E+07 | 7.6E+07 | + | 3181 | long intergenic | 0.73677 | 3.22543 | 51.6381 | 1.17E-09 | 7.77E-08 |
| TCAF1 | 7 | 1.4E+08 | 1.4E+08 | - | 6310 | TRPM8 channe | 0.42828 | 4.92786 | 51.1436 | 1.34E-09 | 8.82E-08 |
| FAM207A | 21 | 4.5E+07 | 4.5E+07 | + | 1400 | family with sec | -0.3296 | 5.87905 | 50.8795 | 1.44E-09 | 9.44E-08 |
| CBR1 | 21 | 3.6E+07 | 3.6E+07 | + | 2450 | carbonyl reduc | -0.2821 | 6.32789 | 50.8659 | 1.44E-09 | 9.44E-08 |
| PCNT | 21 | 4.6E+07 | 4.6E+07 | + | 11641 | pericentrin [So | -0.2574 | 6.72871 | 50.7738 | 1.48E-09 | 9.64E-08 |
| MMP15 | 16 | 5.8E+07 | 5.8E+07 | + | 4231 | matrix metallo | -0.2541 | 6.8745 | 50.9477 | 1.53E-09 | 9.90E-08 |
| KNTC1 | 12 | 1.2E+08 | 1.2E+08 | + | 11325 | kinetochore as | 0.25931 | 6.59848 | 50.6444 | 1.54E-09 | 9.90E-08 |
| CYP2S1 | 19 | 4.1E+07 | 4.1E+07 | + | 2713 | cytochrome P4 | -0.1851 | 8.44004 | 50.4841 | 1.61E-09 | 1.03E-07 |
| RHOBTB3 | 5 | 9.6E+07 | 9.6E+07 | + | 8827 | Rho related BT | 0.58039 | 3.91918 | 50.4788 | 1.61E-09 | 1.03E-07 |
| BACH1 | 21 | 2.9E+07 | 3E+07 | + | 7666 | BTB domain ar | -0.3132 | 6.20059 | 52.4141 | 1.62E-09 | 1.03E-07 |
| IER5 | 1 | 1.8E+08 | 1.8E+08 | + | 4188 | immediate ear | 0.4619 | 4.93018 | 52.1408 | 1.74E-09 | 1.10E-07 |
| IGFBP4 | 17 | 4E+07 | 4E+07 | + | 2205 | insulin like gro | -0.2643 | 7.13864 | 55.8565 | 1.86E-09 | 1.17E-07 |
| DAG1 | 3 | 4.9E+07 | 5E+07 | + | 6923 | dystroglycan 1 | -0.2033 | 7.84805 | 49.9196 | 1.88E-09 | 1.17E-07 |
| TNIK | 3 | 1.7E+08 | 1.7E+08 | - | 11448 | TRAF2 and NCI | 0.23758 | 7.12207 | 50.0335 | 1.91E-09 | 1.19E-07 |
| CRABP2 | 1 | 1.6E+08 | 1.6E+08 | - | 1082 | cellular retinoi | 0.29712 | 6.3509 | 51.5986 | 1.96E-09 | 1.21E-07 |
| DOP1B | 21 | 3.6E+07 | 3.6E+07 | + | 9030 | DOP1 leucine z | -0.4769 | 4.54712 | 49.6393 | 2.03E-09 | 1.25E-07 |
| ITPRID2 | 2 | 1.8E+08 | 1.8E+08 | + | 9685 | ITPR interactin | 0.55767 | 4.02226 | 49.6037 | 2.05E-09 | 1.25E-07 |
| COL23A1 | 5 | 1.8E+08 | 1.8E+08 | - | 3604 | collagen type X | 0.57458 | 3.94621 | 49.5794 | 2.07E-09 | 1.26E-07 |
| MT-ATP8 | MT | 8366 | 8572 | + | 207 | mitochondriall | -0.1827 | 9.80335 | 54.0061 | 2.07E-09 | 1.26E-07 |
| MT-CYB | MT | 14747 | 15887 | + | 1141 | mitochondriall | -0.1421 | 11.7035 | 49.509 | 2.11E-09 | 1.27E-07 |
| PRUNE2 | 9 | 7.7E+07 | 7.7E+07 | - | 14683 | prune homolog | 0.45609 | 4.78987 | 49.1395 | 2.35E-09 | 1.40E-07 |
| ATP2A3 | 17 | 3923870 | 3964464 | - | 6893 | ATPase sarcop | -0.3151 | 5.91639 | 48.9526 | 2.46E-09 | 1.47E-07 |
| NT5DC2 | 3 | 5.3E+07 | 5.3E+07 | - | 4799 | 5'-nucleotidase | 0.19533 | 8.12456 | 48.9385 | 2.47E-09 | 1.47E-07 |
| AC009055 | 16 | 6.5E+07 | 6.5E+07 | - | 1464 | novel transcrip | -0.2873 | 6.27338 | 48.8845 | 2.51E-09 | 1.48E-07 |
| C2CD2 | 21 | 4.2E+07 | 4.2E+07 | - | 8730 | C2 calcium dep | -0.3798 | 5.25717 | 48.5341 | 2.77E-09 | 1.63E-07 |
| AC005062 | 7 | 2E+07 | 2E+07 | - | 2564 | novel transcrip | -0.3753 | 5.33355 | 48.4312 | 2.85E-09 | 1.67E-07 |
| RND3 | 2 | 1.5E+08 | 1.5E+08 | - | 3594 | Rho family GTF | 0.41433 | 5.10623 | 48.6013 | 3.09E-09 | 1.80E-07 |
| PITX2 | 4 | 1.1E+08 | 1.1E+08 | - | 5787 | paired like hor | 2.33402 | 0.91061 | 58.145 | 3.96E-09 | 2.30E-07 |
| MACF1 | 1 | 3.9E+07 | 3.9E+07 | + | 40739 | microtubule-ac | 0.19742 | 9.45621 | 55.9563 | 4.14E-09 | 2.39E-07 |
| CHL1 | 3 | 196763 | 409417 | + | 11694 | cell adhesion n | 2.54365 | 0.17565 | 47.0836 | 4.19E-09 | 2.41E-07 |
| SLC39A14 | 8 | 2.2E+07 | 2.2E+07 | + | 6937 | solute carrier f | -0.1784 | 8.42859 | 46.7491 | 4.61E-09 | 2.64E-07 |
| SLC1A1 | 9 | 4490468 | 4587469 | + | 3974 | solute carrier f | 0.5182 | 4.13098 | 46.6228 | 4.78E-09 | 2.73E-07 |
| TTYH1 | 19 | 5.4E+07 | 5.4E+07 | + | 7068 | tweety family r | -0.444 | 5.06876 | 50.0335 | 4.80E-09 | 2.73E-07 |
| COL6A2 | 21 | 4.6E+07 | 4.6E+07 | + | 5350 | collagen type V | -0.3151 | 7.21773 | 68.19 | 4.97E-09 | 2.81E-07 |
| RBM5 | 3 | 5E+07 | 5E+07 | + | 10368 | RNA binding m | 0.27003 | 6.2448 | 46.4532 | 5.02E-09 | 2.83E-07 |
| URB1 | 21 | 3.2E+07 | 3.2E+07 | - | 11447 | URB1 ribosome | -0.2228 | 7.03862 | 46.4278 | 5.06E-09 | 2.84E-07 |
| HUNK | 21 | 3.2E+07 | 3.2E+07 | + | 8589 | hormonally up | -0.431 | 4.74307 | 46.3631 | 5.15E-09 | 2.88E-07 |
| MXD4 | 4 | 2247432 | 2262109 | - | 6873 | MAX dimerizat | 0.3321 | 5.68566 | 46.0787 | 5.59E-09 | 3.11E-07 |

| | | | | | | | | | | | |
|---|---|---|---|---|---|---|---|---|---|---|---|
| MT-ND4L | MT | 10470 | 10766 | + | 297 | mitochondriall | -0.2177 | 9.97614 | 65.8218 | 5.72E-09 | 3.17E-07 |
| SNX27 | 1 | 1.5E+08 | 1.5E+08 | + | 16711 | sorting nexin f | -0.3274 | 5.61486 | 45.9956 | 5.73E-09 | 3.17E-07 |
| CECR2 | 22 | 1.7E+07 | 1.8E+07 | + | 10759 | CECR2, histone | -0.2146 | 7.79636 | 48.0059 | 6.17E-09 | 3.38E-07 |
| ADCY2 | 5 | 7396208 | 7830081 | + | 17538 | adenylate cycl | 0.20979 | 7.40771 | 45.501 | 6.61E-09 | 3.61E-07 |
| MAP7D1 | 1 | 3.6E+07 | 3.6E+07 | + | 5271 | MAP7 domain | 0.23998 | 6.794 | 45.36 | 6.89E-09 | 3.75E-07 |
| SPRY2 | 13 | 8E+07 | 8E+07 | - | 3213 | sprouty RTK sig | 0.34693 | 5.55265 | 45.2978 | 7.02E-09 | 3.80E-07 |
| MEIS3 | 19 | 4.7E+07 | 4.7E+07 | - | 5122 | Meis homeobo | 0.32515 | 5.5799 | 45.2259 | 7.17E-09 | 3.87E-07 |
| CDH1 | 16 | 6.9E+07 | 6.9E+07 | + | 6103 | cadherin 1 [So | -0.2018 | 8.08904 | 46.9354 | 7.18E-09 | 3.87E-07 |
| INSR | 19 | 7112255 | 7294414 | - | 10831 | insulin recepto | -0.289 | 5.99596 | 45.083 | 7.47E-09 | 3.99E-07 |
| HLCS | 21 | 3.7E+07 | 3.7E+07 | - | 7889 | holocarboxylas | -0.4225 | 4.81224 | 44.765 | 8.20E-09 | 4.36E-07 |
| METRN | 16 | 715118 | 719655 | + | 3689 | meteorin, glial | 0.48749 | 4.43965 | 44.7076 | 9.14E-09 | 4.84E-07 |
| IFNGR2 | 21 | 3.3E+07 | 3.3E+07 | + | 2960 | interferon gam | -0.349 | 5.40548 | 44.2233 | 9.61E-09 | 5.08E-07 |
| LSM4 | 19 | 1.8E+07 | 1.8E+07 | - | 3029 | LSM4 homolog | 0.20057 | 7.6205 | 44.175 | 9.75E-09 | 5.13E-07 |
| ZHX2 | 8 | 1.2E+08 | 1.2E+08 | + | 4453 | zinc fingers an | 0.38366 | 5.05976 | 44.0833 | 1.00E-08 | 5.25E-07 |
| NSG1 | 4 | 4348140 | 4419058 | + | 4355 | neuronal vesic | -0.3062 | 5.78616 | 44.0484 | 1.01E-08 | 5.29E-07 |
| LOXL3 | 2 | 7.5E+07 | 7.5E+07 | - | 4519 | lysyl oxidase li | 0.70957 | 3.23318 | 43.8946 | 1.06E-08 | 5.50E-07 |
| GRID2 | 4 | 9.2E+07 | 9.4E+07 | + | 10617 | glutamate ion | 0.32452 | 5.71395 | 43.8369 | 1.08E-08 | 5.57E-07 |
| SOX3 | X | 1.4E+08 | 1.4E+08 | - | 2132 | SRY-box 3 [Sou | 0.34287 | 5.53088 | 43.8097 | 1.09E-08 | 5.59E-07 |
| CCND2 | 12 | 4273762 | 4305353 | + | 7170 | cyclin D2 [Sour | -0.2319 | 6.89367 | 43.7766 | 1.10E-08 | 5.62E-07 |
| PRXL2B | HSCHR1_1 | 2586491 | 2591469 | + | 3371 | peroxiredoxin | -0.277 | 6.13639 | 43.7685 | 1.10E-08 | 5.62E-07 |
| ETV1 | 7 | 1.4E+07 | 1.4E+07 | - | 9615 | ETS variant 1 [ | 0.22457 | 6.88579 | 43.7122 | 1.12E-08 | 5.70E-07 |
| COL4A6 | X | 1.1E+08 | 1.1E+08 | - | 9282 | collagen type I | 0.45559 | 4.57275 | 43.6357 | 1.14E-08 | 5.81E-07 |
| DUSP16 | 12 | 1.2E+07 | 1.3E+07 | - | 5724 | dual specificity | 0.22192 | 6.85638 | 43.5041 | 1.19E-08 | 6.02E-07 |
| UHRF2 | 9 | 6413151 | 6507054 | + | 9919 | ubiquitin like w | 0.31152 | 5.73784 | 43.3846 | 1.23E-08 | 6.21E-07 |
| GALNT7 | 4 | 1.7E+08 | 1.7E+08 | + | 6807 | polypeptide N- | 0.23678 | 6.79221 | 43.2274 | 1.29E-08 | 6.47E-07 |
| HADH | 4 | 1.1E+08 | 1.1E+08 | + | 12086 | hydroxyacyl-Co | 0.28808 | 5.92287 | 43.2224 | 1.29E-08 | 6.47E-07 |
| SHROOM3 | 4 | 7.6E+07 | 7.7E+07 | + | 13829 | shroom family | 0.22094 | 6.94768 | 43.1685 | 1.31E-08 | 6.53E-07 |
| OPN3 | 1 | 2.4E+08 | 2.4E+08 | - | 3724 | opsin 3 [Sourc | 0.74754 | 2.91492 | 43.0842 | 1.35E-08 | 6.68E-07 |
| MT-CO2 | MT | 7586 | 8269 | + | 684 | mitochondriall | -0.1309 | 11.8658 | 42.8375 | 1.45E-08 | 7.14E-07 |
| CRMP1 | 4 | 5748084 | 5893058 | - | 5823 | collapsin respo | -0.2101 | 7.23594 | 42.5556 | 1.58E-08 | 7.74E-07 |
| KLHDC7A | 1 | 1.8E+07 | 1.8E+07 | + | 5145 | kelch domain | 0.36153 | 5.16841 | 42.4785 | 1.62E-08 | 7.89E-07 |
| FAM107B | 10 | 1.5E+07 | 1.5E+07 | - | 7019 | family with se | -0.4197 | 4.85707 | 42.4986 | 1.71E-08 | 8.34E-07 |
| AMD1 | 6 | 1.1E+08 | 1.1E+08 | + | 7027 | adenosylmethi | -0.2085 | 8.03516 | 46.5877 | 1.72E-08 | 8.36E-07 |
| CD9 | 12 | 6199715 | 6238271 | + | 4165 | CD9 molecule | -0.2223 | 6.95171 | 42.254 | 1.73E-08 | 8.36E-07 |
| PHC2 | 1 | 3.3E+07 | 3.3E+07 | - | 4975 | polyhomeotic | 0.59422 | 3.59179 | 42.2228 | 1.74E-08 | 8.41E-07 |
| CD24 | 6 | 1.1E+08 | 1.1E+08 | - | 3096 | CD24 molecule | -0.154 | 9.62248 | 41.9526 | 1.89E-08 | 9.06E-07 |
| DOT1L | 19 | 2163933 | 2232578 | + | 11488 | DOT1 like histo | 0.23967 | 6.60809 | 41.8535 | 1.95E-08 | 9.30E-07 |
| NRIP1 | 21 | 1.5E+07 | 1.5E+07 | - | 8497 | nuclear recept | -0.3954 | 4.93281 | 41.8445 | 1.95E-08 | 9.30E-07 |
| CXCL12 | 10 | 4.4E+07 | 4.4E+07 | - | 6077 | C-X-C motif ch | -0.3371 | 5.67262 | 43.3186 | 1.96E-08 | 9.31E-07 |
| MLLT6 | 17 | 3.9E+07 | 3.9E+07 | + | 13315 | MLLT6, PHD fi | 0.75131 | 2.86015 | 41.8007 | 1.98E-08 | 9.36E-07 |
| TENM3-A | 4 | 1.8E+08 | 1.8E+08 | - | 5551 | TENM3 antiser | 0.55088 | 3.82804 | 41.6273 | 2.09E-08 | 9.83E-07 |
| KCND2 | 7 | 1.2E+08 | 1.2E+08 | + | 6054 | potassium volt | 0.48725 | 4.15216 | 41.5824 | 2.12E-08 | 9.93E-07 |
| NUCB2 | 11 | 1.7E+07 | 1.7E+07 | + | 9025 | nucleobindin 2 | -0.6364 | 3.51929 | 41.5737 | 2.12E-08 | 9.93E-07 |

| | | | | | | | | | | | |
|---|---|---|---|---|---|---|---|---|---|---|---|
| CD82 | 11 | 4.5E+07 | 4.5E+07 | + | 4039 | CD82 molecule | -0.6991 | 3.21057 | 41.503 | 2.17E-08 | 1.01E-06 |
| PAXBP1 | 21 | 3.3E+07 | 3.3E+07 | - | 7146 | PAX3 and PAX7 | -0.3123 | 7.01584 | 60.3836 | 2.18E-08 | 1.01E-06 |
| NEK6 | 9 | 1.2E+08 | 1.2E+08 | + | 4419 | NIMA related k | 0.39653 | 5.03141 | 42.1466 | 2.18E-08 | 1.01E-06 |
| ZIC5 | 13 | 1E+08 | 1E+08 | - | 4639 | Zic family mem | 0.30714 | 5.67751 | 41.4221 | 2.22E-08 | 1.03E-06 |
| KHDRBS2 | 6 | 6.2E+07 | 6.2E+07 | - | 2329 | KH RNA bindin | 0.82798 | 2.72446 | 41.5733 | 2.26E-08 | 1.04E-06 |
| ATP10B | 5 | 1.6E+08 | 1.6E+08 | - | 11310 | ATPase phosph | 1.61942 | 0.8826 | 41.3234 | 2.29E-08 | 1.05E-06 |
| PALLD | 4 | 1.7E+08 | 1.7E+08 | + | 10128 | palladin, cytos | 0.41221 | 5.38369 | 48.7878 | 2.36E-08 | 1.08E-06 |
| MT-ND2 | MT | 4470 | 5511 | + | 1042 | mitochondriall | -0.1412 | 10.8132 | 41.1288 | 2.43E-08 | 1.11E-06 |
| HDAC9 | 7 | 1.8E+07 | 1.9E+07 | + | 16827 | histone deacet | 0.50275 | 4.08121 | 41.0702 | 2.47E-08 | 1.12E-06 |
| ZMIZ2 | 7 | 4.5E+07 | 4.5E+07 | + | 7354 | zinc finger MIZ | 0.28926 | 5.81535 | 40.9974 | 2.53E-08 | 1.15E-06 |
| PHLDB2 | 3 | 1.1E+08 | 1.1E+08 | + | 12309 | pleckstrin hom | 0.34483 | 5.29463 | 40.9584 | 2.56E-08 | 1.16E-06 |
| DDX6 | 11 | 1.2E+08 | 1.2E+08 | - | 11462 | DEAD-box helic | -0.2039 | 7.12311 | 40.8935 | 2.61E-08 | 1.17E-06 |
| PCNX2 | 1 | 2.3E+08 | 2.3E+08 | - | 13827 | pecanex 2 [Sou | 0.26279 | 6.18534 | 40.7962 | 2.69E-08 | 1.21E-06 |
| PFKL | 21 | 4.4E+07 | 4.4E+07 | + | 8926 | phosphofructo | -0.1831 | 7.9177 | 40.6164 | 2.84E-08 | 1.27E-06 |
| HTR7 | 10 | 9.1E+07 | 9.1E+07 | - | 3472 | 5-hydroxytrypt | 0.30036 | 5.85673 | 40.8996 | 2.86E-08 | 1.28E-06 |
| HSPA13 | 21 | 1.4E+07 | 1.4E+07 | - | 4153 | heat shock pro | -0.2559 | 6.18096 | 40.4872 | 2.95E-08 | 1.31E-06 |
| RABL6 | 9 | 1.4E+08 | 1.4E+08 | + | 10178 | RAB, member | 0.19472 | 7.42724 | 40.4166 | 3.02E-08 | 1.34E-06 |
| DBI | 2 | 1.2E+08 | 1.2E+08 | + | 2509 | diazepam bind | -0.2257 | 6.6681 | 40.4004 | 3.03E-08 | 1.34E-06 |
| ID4 | 6 | 2E+07 | 2E+07 | + | 3873 | inhibitor of DN | -0.6087 | 4.1042 | 47.1984 | 3.10E-08 | 1.36E-06 |
| SYT13 | 11 | 4.5E+07 | 4.5E+07 | - | 5445 | synaptotagmin | -0.4153 | 4.73599 | 40.1549 | 3.27E-08 | 1.43E-06 |
| RBM15B | 3 | 5.1E+07 | 5.1E+07 | + | 6624 | RNA binding m | 0.21313 | 6.87079 | 40.1288 | 3.30E-08 | 1.44E-06 |
| SCMH1 | 1 | 4.1E+07 | 4.1E+07 | - | 4447 | Scm polycomb | 0.4742 | 4.30823 | 40.1094 | 3.32E-08 | 1.44E-06 |
| UBE2H | 7 | 1.3E+08 | 1.3E+08 | - | 6503 | ubiquitin conju | 0.1972 | 7.39504 | 40.015 | 3.41E-08 | 1.48E-06 |
| BACE2 | 21 | 4.1E+07 | 4.1E+07 | + | 9631 | beta-secretase | -0.4267 | 4.54471 | 39.9483 | 3.48E-08 | 1.51E-06 |
| JDP2 | 14 | 7.5E+07 | 7.5E+07 | + | 6259 | Jun dimerizatio | -0.3954 | 4.77732 | 39.9162 | 3.52E-08 | 1.52E-06 |
| LHX8 | 1 | 7.5E+07 | 7.5E+07 | + | 3977 | LIM homeobox | 2.006 | 0.53201 | 40.3731 | 3.64E-08 | 1.56E-06 |
| ARGLU1 | 13 | 1.1E+08 | 1.1E+08 | - | 5783 | arginine and gl | 0.17639 | 7.96537 | 39.7631 | 3.69E-08 | 1.58E-06 |
| OR52A1 | 11 | 5143219 | 5154757 | - | 9783 | olfactory recep | -1.0237 | 2.04435 | 39.7218 | 3.74E-08 | 1.60E-06 |
| EEF2K | 16 | 2.2E+07 | 2.2E+07 | + | 8030 | eukaryotic elon | -0.337 | 5.36637 | 39.6877 | 3.78E-08 | 1.61E-06 |
| PROSER1 | 13 | 3.9E+07 | 3.9E+07 | - | 7329 | proline and ser | -0.2153 | 7.32101 | 41.3775 | 3.84E-08 | 1.63E-06 |
| ACAA2 | 18 | 5E+07 | 5E+07 | - | 4920 | acetyl-CoA acy | -0.2029 | 7.0604 | 39.6269 | 3.85E-08 | 1.63E-06 |
| PIEZO1 | 16 | 8.9E+07 | 8.9E+07 | - | 11292 | piezo type mec | 0.25371 | 6.34676 | 39.687 | 3.89E-08 | 1.64E-06 |
| RN7SL1 | 14 | 5E+07 | 5E+07 | + | 299 | RNA, 7SL, cyto | -0.7298 | 5.51644 | 99.7559 | 4.18E-08 | 1.76E-06 |
| VAT1 | 17 | 4.3E+07 | 4.3E+07 | - | 3466 | vesicle amine t | -0.156 | 8.85982 | 39.3183 | 4.23E-08 | 1.78E-06 |
| COL26A1 | 7 | 1E+08 | 1E+08 | + | 3101 | collagen type X | 0.20165 | 7.23008 | 39.3009 | 4.26E-08 | 1.78E-06 |
| STK35 | 20 | 2101611 | 2177038 | + | 7055 | serine/threoni | -0.2585 | 6.14272 | 39.1859 | 4.41E-08 | 1.84E-06 |
| SCGB3A2 | 5 | 1.5E+08 | 1.5E+08 | + | 1075 | secretoglobin f | -0.3158 | 5.56448 | 39.1818 | 4.42E-08 | 1.84E-06 |
| RYR2 | 1 | 2.4E+08 | 2.4E+08 | + | 17798 | ryanodine rece | 0.44403 | 4.64778 | 40.3598 | 4.49E-08 | 1.87E-06 |
| SETD4 | 21 | 3.6E+07 | 3.6E+07 | - | 7312 | SET domain co | -0.3494 | 5.17922 | 39.072 | 4.57E-08 | 1.89E-06 |
| HMGCS1 | 5 | 4.3E+07 | 4.3E+07 | - | 6627 | 3-hydroxy-3-m | -0.1647 | 8.5099 | 38.93 | 4.78E-08 | 1.96E-06 |
| POFUT2 | 21 | 4.5E+07 | 4.5E+07 | - | 5862 | protein O-fuco | -0.2837 | 5.78705 | 38.7898 | 4.99E-08 | 2.04E-06 |
| GNA14 | 9 | 7.7E+07 | 7.8E+07 | - | 2736 | G protein subu | 0.91568 | 3.9219 | 70.297 | 5.26E-08 | 2.15E-06 |
| VWCE | 11 | 6.1E+07 | 6.1E+07 | - | 4766 | von Willebrand | -0.466 | 4.23429 | 38.3834 | 5.66E-08 | 2.31E-06 |

| | | | | | | | | | | | | |
|---|---|---|---|---|---|---|---|---|---|---|---|---|
| RCC2 | 1 | 1.7E+07 | 1.7E+07 | - | | 4079 | regulator of ch | -0.1466 | 9.2973 | 38.3206 | 5.78E-08 | 2.34E-06 |
| NUP210 | 3 | 1.3E+07 | 1.3E+07 | - | | 8195 | nucleoporin 21 | 0.17767 | 7.8166 | 38.3168 | 5.78E-08 | 2.34E-06 |
| EIF2S3 | X | 2.4E+07 | 2.4E+07 | + | | 4081 | eukaryotic tran | -0.1753 | 7.94875 | 38.2522 | 5.90E-08 | 2.38E-06 |
| RND2 | 17 | 4.3E+07 | 4.3E+07 | + | | 4368 | Rho family GTP | -0.2151 | 6.82591 | 38.0792 | 6.23E-08 | 2.50E-06 |
| DONSON | 21 | 3.4E+07 | 3.4E+07 | - | | 3511 | downstream n | -0.3006 | 5.62935 | 38.0645 | 6.26E-08 | 2.51E-06 |
| VCAN | 5 | 8.3E+07 | 8.4E+07 | + | | 14678 | versican [Sourc | -0.1431 | 9.6205 | 37.8771 | 6.64E-08 | 2.65E-06 |
| GALNT17 | 7 | 7.1E+07 | 7.2E+07 | + | | 4895 | polypeptide N- | 0.24108 | 6.25999 | 37.7897 | 6.82E-08 | 2.72E-06 |
| GAS7 | 17 | 9910609 | 1E+07 | - | | 10100 | growth arrest s | -0.4282 | 4.65321 | 38.1761 | 7.21E-08 | 2.85E-06 |
| TET1 | 10 | 6.9E+07 | 6.9E+07 | + | | 9612 | tet methylcyto | -0.1934 | 7.31888 | 37.5951 | 7.26E-08 | 2.86E-06 |
| LAMA1 | 18 | 6941742 | 7117797 | - | | 15857 | laminin subuni | -0.1993 | 7.1275 | 37.3432 | 7.86E-08 | 3.08E-06 |
| SALL3 | 18 | 7.9E+07 | 7.9E+07 | + | | 7397 | spalt like trans | -0.3434 | 5.26201 | 37.3339 | 7.88E-08 | 3.09E-06 |
| SORBS2 | 4 | 1.9E+08 | 1.9E+08 | - | | 17161 | sorbin and SH3 | 0.65958 | 3.09803 | 37.2754 | 8.03E-08 | 3.13E-06 |
| KDR | 4 | 5.5E+07 | 5.5E+07 | - | | 6563 | kinase insert d | -0.211 | 6.8401 | 37.1569 | 8.34E-08 | 3.25E-06 |
| ANO6 | 12 | 4.5E+07 | 4.5E+07 | + | | 8235 | anoctamin 6 [S | 0.20242 | 6.94837 | 37.1041 | 8.48E-08 | 3.28E-06 |
| DHRS3 | 1 | 1.3E+07 | 1.3E+07 | - | | 3515 | dehydrogenase | -0.4683 | 4.14243 | 37.1035 | 8.48E-08 | 3.28E-06 |
| GAS6 | 13 | 1.1E+08 | 1.1E+08 | - | | 6015 | growth arrest s | 0.37255 | 4.82553 | 37.0146 | 8.72E-08 | 3.37E-06 |
| USP16 | 21 | 2.9E+07 | 2.9E+07 | + | | 5694 | ubiquitin speci | -0.4085 | 5.21614 | 42.9369 | 8.76E-08 | 3.38E-06 |
| HAS3 | 16 | 6.9E+07 | 6.9E+07 | + | | 5200 | hyaluronan syn | 0.21381 | 6.73074 | 36.9885 | 8.79E-08 | 3.38E-06 |
| TENT5B | 1 | 2.7E+07 | 2.7E+07 | - | | 2382 | terminal nucle | -0.197 | 7.19226 | 36.9652 | 8.86E-08 | 3.39E-06 |
| SEH1L | 18 | 1.3E+07 | 1.3E+07 | + | | 10481 | SEH1 like nucle | -0.2747 | 6.06642 | 37.7049 | 9.24E-08 | 3.52E-06 |
| ID1 | 20 | 3.2E+07 | 3.2E+07 | + | | 1233 | inhibitor of DN | -0.2521 | 6.35581 | 37.6957 | 9.26E-08 | 3.52E-06 |
| GNL3L | X | 5.5E+07 | 5.5E+07 | + | | 8807 | G protein nucle | -0.2684 | 5.96808 | 36.707 | 9.62E-08 | 3.65E-06 |
| ATP11A | 13 | 1.1E+08 | 1.1E+08 | + | | 13733 | ATPase phosph | 0.21957 | 6.6043 | 36.6759 | 9.71E-08 | 3.67E-06 |
| PTPN13 | 4 | 8.7E+07 | 8.7E+07 | + | | 9496 | protein tyrosin | 0.19818 | 7.14888 | 36.6551 | 9.78E-08 | 3.69E-06 |
| SLC7A8 | 14 | 2.3E+07 | 2.3E+07 | - | | 5991 | solute carrier f | -0.1682 | 8.04249 | 36.6441 | 9.81E-08 | 3.69E-06 |
| ANK2 | 4 | 1.1E+08 | 1.1E+08 | + | | 18224 | ankyrin 2 [Sou | 0.29487 | 5.70634 | 36.6222 | 9.88E-08 | 3.70E-06 |
| PCSK5 | 9 | 7.6E+07 | 7.6E+07 | + | | 12705 | proprotein con | 0.39368 | 4.6692 | 36.6161 | 9.90E-08 | 3.70E-06 |
| UBE2J1 | 6 | 8.9E+07 | 8.9E+07 | - | | 4164 | ubiquitin conju | 0.32152 | 5.33137 | 36.5854 | 1.00E-07 | 3.73E-06 |
| TNKS1BP1 | 11 | 5.7E+07 | 5.7E+07 | - | | 7149 | tankyrase 1 bin | -0.1796 | 7.66593 | 36.4823 | 1.03E-07 | 3.85E-06 |
| EFS | 14 | 2.3E+07 | 2.3E+07 | - | | 3236 | embryonal Fyn | 0.35915 | 4.89813 | 36.4669 | 1.04E-07 | 3.86E-06 |
| CEBPZ | 2 | 3.7E+07 | 3.7E+07 | - | | 3512 | CCAAT enhanc | -0.1821 | 7.62765 | 36.2669 | 1.11E-07 | 4.08E-06 |
| PIM1 | 6 | 3.7E+07 | 3.7E+07 | + | | 3030 | Pim-1 proto-or | -0.196 | 7.05923 | 36.2278 | 1.12E-07 | 4.12E-06 |
| ETV5 | 3 | 1.9E+08 | 1.9E+08 | - | | 5773 | ETS variant 5 [S | 0.28065 | 5.84756 | 36.1768 | 1.14E-07 | 4.18E-06 |
| ENTPD6 | 20 | 2.5E+07 | 2.5E+07 | + | | 5327 | ectonucleoside | -0.2642 | 5.95532 | 36.0832 | 1.17E-07 | 4.29E-06 |
| MORC3 | 21 | 3.6E+07 | 3.6E+07 | + | | 6408 | MORC family C | -0.3501 | 5.10311 | 35.9103 | 1.24E-07 | 4.53E-06 |
| FGD5 | 3 | 1.5E+07 | 1.5E+07 | + | | 8093 | FYVE, RhoGEF | 0.30597 | 5.87133 | 38.8758 | 1.32E-07 | 4.80E-06 |
| EZH2 | 7 | 1.5E+08 | 1.5E+08 | - | | 4522 | enhancer of ze | 0.25196 | 6.08254 | 35.7051 | 1.33E-07 | 4.80E-06 |
| IGFBP7 | 4 | 5.7E+07 | 5.7E+07 | - | | 1930 | insulin like gro | 1.1161 | 1.76858 | 36.0782 | 1.38E-07 | 4.98E-06 |
| DTX4 | 11 | 5.9E+07 | 5.9E+07 | + | | 6433 | deltex E3 ubiqu | -0.2752 | 5.77942 | 35.485 | 1.42E-07 | 5.11E-06 |
| CGN | 1 | 1.5E+08 | 1.5E+08 | + | | 6616 | cingulin [Sourc | -0.1974 | 6.96935 | 35.4763 | 1.43E-07 | 5.12E-06 |
| ZNF286A | 17 | 1.6E+07 | 1.6E+07 | + | | 6754 | zinc finger prot | -0.3653 | 4.90683 | 35.3804 | 1.47E-07 | 5.26E-06 |
| RPL7 | 8 | 7.3E+07 | 7.3E+07 | - | | 2700 | ribosomal prot | -0.1401 | 9.41305 | 35.3643 | 1.48E-07 | 5.28E-06 |
| PROM1 | 4 | 1.6E+07 | 1.6E+07 | - | | 7434 | prominin 1 [So | 0.23323 | 6.38948 | 35.3336 | 1.49E-07 | 5.32E-06 |

| | | | | | | | | | | | |
|---|---|---|---|---|---|---|---|---|---|---|---|
| ZFP36L2 | 2 | 4.3E+07 | 4.3E+07 | - | 3693 | ZFP36 ring fing | -0.2101 | 6.78738 | 35.3109 | 1.51E-07 | 5.34E-06 |
| TMEM63A | 1 | 2.3E+08 | 2.3E+08 | - | 10350 | transmembran | 0.53281 | 6.10649 | 80.8545 | 1.51E-07 | 5.34E-06 |
| CORO1B | 11 | 6.7E+07 | 6.7E+07 | - | 5327 | coronin 1B [So | 0.19776 | 6.89712 | 35.2409 | 1.54E-07 | 5.44E-06 |
| PSMA4 | 15 | 7.9E+07 | 7.9E+07 | + | 6994 | proteasome su | -0.1729 | 7.66181 | 35.1148 | 1.60E-07 | 5.65E-06 |
| CENPT | 16 | 6.8E+07 | 6.8E+07 | - | 5660 | centromere pr | 0.32982 | 5.221 | 35.0168 | 1.66E-07 | 5.82E-06 |
| PLEKHG4E | 5 | 92151 | 189972 | + | 13304 | pleckstrin hom | 0.24109 | 6.12823 | 35.004 | 1.66E-07 | 5.83E-06 |
| RAPGEF5 | 7 | 2.2E+07 | 2.2E+07 | - | 10245 | Rap guanine nu | 0.57026 | 3.40593 | 34.821 | 1.77E-07 | 6.17E-06 |
| RABGAP1 | 1 | 1.7E+08 | 1.7E+08 | + | 16071 | RAB GTPase ac | -0.2306 | 6.94169 | 38.2948 | 1.94E-07 | 6.70E-06 |
| MYO10 | 5 | 1.7E+07 | 1.7E+07 | - | 13982 | myosin X [Sou | -0.1659 | 8.14999 | 34.5248 | 1.94E-07 | 6.72E-06 |
| PPDPF | 20 | 6.4E+07 | 6.4E+07 | + | 1053 | pancreatic pro | 0.22185 | 6.42852 | 34.4708 | 1.98E-07 | 6.82E-06 |
| MID1 | X | 1E+07 | 1.1E+07 | - | 9295 | midline 1 [Sou | -0.2816 | 6.22549 | 39.0141 | 2.00E-07 | 6.87E-06 |
| FAM83H | HSCHR8_3 | 1.4E+08 | 1.4E+08 | - | 6220 | family with seq | 0.19917 | 6.86982 | 34.4156 | 2.02E-07 | 6.91E-06 |
| LARP1 | 5 | 1.5E+08 | 1.5E+08 | + | 8549 | La ribonucleop | -0.1423 | 8.98546 | 34.3289 | 2.07E-07 | 7.09E-06 |
| EPB41L1 | 20 | 3.6E+07 | 3.6E+07 | + | 9676 | erythrocyte me | -0.2706 | 5.7935 | 34.2287 | 2.14E-07 | 7.31E-06 |
| LINC0064 | 14 | 2.1E+07 | 2.1E+07 | - | 5715 | long intergenic | 0.41804 | 4.37586 | 34.1245 | 2.22E-07 | 7.55E-06 |
| SMCHD1 | 18 | 2655738 | 2805017 | + | 12435 | structural mair | 0.22193 | 6.51083 | 33.9854 | 2.32E-07 | 7.86E-06 |
| TMEM200 | 6 | 1.3E+08 | 1.3E+08 | + | 6137 | transmembran | 0.83792 | 4.91033 | 86.9602 | 2.32E-07 | 7.86E-06 |
| ACVR2B | 3 | 3.8E+07 | 3.8E+07 | + | 15705 | activin A recep | -0.174 | 7.7112 | 33.9781 | 2.33E-07 | 7.86E-06 |
| ADD1 | 4 | 2843857 | 2930076 | + | 13227 | adducin 1 [Sou | 0.17045 | 7.64778 | 33.9433 | 2.35E-07 | 7.94E-06 |
| TECR | 19 | 1.5E+07 | 1.5E+07 | + | 5899 | trans-2,3-enoy | -0.2063 | 6.65363 | 33.9339 | 2.36E-07 | 7.94E-06 |
| NEURL1B | 5 | 1.7E+08 | 1.7E+08 | + | 6424 | neuralized E3 u | 0.40516 | 4.45465 | 33.878 | 2.40E-07 | 8.07E-06 |
| PM20D2 | 6 | 8.9E+07 | 8.9E+07 | + | 4703 | peptidase M20 | 0.56636 | 3.44683 | 33.8452 | 2.43E-07 | 8.14E-06 |
| SNHG6 | 8 | 6.7E+07 | 6.7E+07 | - | 1247 | small nucleolar | 0.25238 | 5.97671 | 33.7659 | 2.49E-07 | 8.33E-06 |
| LRIG1 | 3 | 6.6E+07 | 6.7E+07 | - | 7032 | leucine rich rep | 0.19148 | 7.29966 | 33.9911 | 2.69E-07 | 8.97E-06 |
| UNC5C | 4 | 9.5E+07 | 9.6E+07 | - | 10685 | unc-5 netrin re | 2.2123 | 0.88261 | 45.9978 | 2.75E-07 | 9.15E-06 |
| CLCN5 | X | 5E+07 | 5E+07 | + | 11457 | chloride voltag | -0.3434 | 4.92596 | 33.2722 | 2.94E-07 | 9.72E-06 |
| MXRA7 | 17 | 7.7E+07 | 7.7E+07 | - | 9257 | matrix remode | 0.42173 | 4.23587 | 33.2507 | 2.96E-07 | 9.77E-06 |
| RIPK4 | 21 | 4.2E+07 | 4.2E+07 | - | 4016 | receptor intera | -0.5552 | 3.45909 | 33.2268 | 2.98E-07 | 9.82E-06 |
| FBLN2 | 3 | 1.4E+07 | 1.4E+07 | + | 4818 | fibulin 2 [Sourc | 0.25791 | 5.93804 | 33.2176 | 2.99E-07 | 9.83E-06 |
| NOL4L | 20 | 3.2E+07 | 3.3E+07 | - | 9448 | nucleolar prote | -0.3107 | 5.30202 | 33.1748 | 3.03E-07 | 9.95E-06 |
| SEPHS1 | 10 | 1.3E+07 | 1.3E+07 | - | 3884 | selenophospha | -0.1333 | 9.79841 | 33.1434 | 3.06E-07 | 1.00E-05 |
| TLE3 | 15 | 7E+07 | 7E+07 | - | 12634 | TLE family mer | -0.2428 | 6.01768 | 33.0251 | 3.19E-07 | 1.04E-05 |
| TPT1 | 13 | 4.5E+07 | 4.5E+07 | - | 7269 | tumor protein, | 0.13185 | 9.59341 | 33.0242 | 3.19E-07 | 1.04E-05 |
| SCD | 10 | 1E+08 | 1E+08 | + | 5245 | stearoyl-CoA d | -0.1248 | 10.7521 | 32.9882 | 3.23E-07 | 1.05E-05 |
| MYO1B | 2 | 1.9E+08 | 1.9E+08 | + | 9533 | myosin IB [Sou | 0.2042 | 6.66869 | 32.929 | 3.29E-07 | 1.07E-05 |
| LRBA | 4 | 1.5E+08 | 1.5E+08 | - | 16757 | LPS responsive | 0.16092 | 8.00123 | 32.9158 | 3.30E-07 | 1.07E-05 |
| MTA3 | 2 | 4.2E+07 | 4.3E+07 | + | 10580 | metastasis ass | -0.1798 | 7.32692 | 32.913 | 3.31E-07 | 1.07E-05 |
| PTTG1IP | 21 | 4.5E+07 | 4.5E+07 | - | 3537 | PTTG1 interact | -0.1557 | 8.15413 | 32.7614 | 3.48E-07 | 1.12E-05 |
| OPLAH | 8 | 1.4E+08 | 1.4E+08 | - | 4441 | 5-oxoprolinase | 0.33126 | 5.02592 | 32.7325 | 3.51E-07 | 1.13E-05 |
| ARMCX2 | X | 1E+08 | 1E+08 | - | 3548 | armadillo repe | 0.18552 | 7.16119 | 32.7154 | 3.53E-07 | 1.13E-05 |
| SNHG29 | 17 | 1.6E+07 | 1.6E+07 | + | 3759 | small nucleolar | 0.16071 | 7.94016 | 32.6305 | 3.63E-07 | 1.16E-05 |
| MTA1 | 14 | 1.1E+08 | 1.1E+08 | + | 6966 | metastasis ass | 0.16804 | 7.60671 | 32.5723 | 3.71E-07 | 1.18E-05 |
| XYLT1 | 16 | 1.7E+07 | 1.7E+07 | - | 11004 | xylosyltransfer | -0.2745 | 5.67464 | 32.4563 | 3.85E-07 | 1.23E-05 |

| | | | | | | | | | | | |
|---|---|---|---|---|---|---|---|---|---|---|---|
| FJX1 | 11 | 3.6E+07 | 3.6E+07 | + | 2450 | four-jointed bo | 0.50475 | 3.706 | 32.4063 | 3.92E-07 | 1.25E-05 |
| HS3ST3B1 | 17 | 1.4E+07 | 1.4E+07 | + | 5369 | heparan sulfat | -0.6143 | 3.15338 | 32.3358 | 4.01E-07 | 1.27E-05 |
| HSPA2 | 14 | 6.5E+07 | 6.5E+07 | + | 5777 | heat shock pro | -0.2644 | 5.75983 | 32.3269 | 4.02E-07 | 1.27E-05 |
| FLCN | 17 | 1.7E+07 | 1.7E+07 | - | 9786 | folliculin [Sour | -0.3544 | 4.81391 | 32.2412 | 4.14E-07 | 1.31E-05 |
| POLR3G | 5 | 9E+07 | 9.1E+07 | + | 4023 | RNA polymera | 0.2187 | 7.12542 | 36.4815 | 4.20E-07 | 1.32E-05 |
| PNPLA6 | 19 | 7534004 | 7561764 | + | 7172 | patatin like ph | -0.1779 | 7.25157 | 32.1625 | 4.25E-07 | 1.33E-05 |
| CTBP2 | 10 | 1.2E+08 | 1.3E+08 | - | 11056 | C-terminal bin | 0.15825 | 7.90375 | 32.1423 | 4.28E-07 | 1.34E-05 |
| SPRY4 | 5 | 1.4E+08 | 1.4E+08 | - | 6565 | sprouty RTK sig | 0.17333 | 8.21693 | 34.3929 | 4.31E-07 | 1.35E-05 |
| PCDH11X | X | 9.2E+07 | 9.3E+07 | + | 11630 | protocadherin | 0.59228 | 3.18248 | 31.9079 | 4.63E-07 | 1.44E-05 |
| ADAM23 | 2 | 2.1E+08 | 2.1E+08 | + | 6425 | ADAM metallo | 0.33343 | 5.10174 | 31.8604 | 4.70E-07 | 1.46E-05 |
| SLC1A6 | 19 | 1.5E+07 | 1.5E+07 | - | 7646 | solute carrier f | -0.4787 | 3.81585 | 31.8368 | 4.74E-07 | 1.47E-05 |
| CCDC85B | 11 | 6.6E+07 | 6.6E+07 | + | 1524 | coiled-coil dom | 0.28957 | 5.41815 | 31.7317 | 4.91E-07 | 1.52E-05 |
| KCND1 | X | 4.9E+07 | 4.9E+07 | - | 6082 | potassium volt | -0.5968 | 3.15888 | 31.6038 | 5.13E-07 | 1.59E-05 |
| AP000688 | 21 | 3.6E+07 | 3.6E+07 | + | 2336 | novel transcrip | -0.5441 | 3.47926 | 31.5917 | 5.15E-07 | 1.59E-05 |
| LRP1 | 12 | 5.7E+07 | 5.7E+07 | + | 20839 | LDL receptor r | -0.1381 | 9.02293 | 31.5914 | 5.15E-07 | 1.59E-05 |
| UBE2G1 | 17 | 4269259 | 4366628 | - | 4539 | ubiquitin conju | -0.1912 | 7.2498 | 32.3916 | 5.17E-07 | 1.59E-05 |
| CLIP3 | 19 | 3.6E+07 | 3.6E+07 | - | 3765 | CAP-Gly domai | -0.3469 | 4.99355 | 31.6727 | 5.23E-07 | 1.60E-05 |
| LMAN1 | 18 | 5.9E+07 | 5.9E+07 | - | 6070 | lectin, mannos | -0.1768 | 7.33009 | 31.4327 | 5.44E-07 | 1.66E-05 |
| HK2 | 2 | 7.5E+07 | 7.5E+07 | + | 6064 | hexokinase 2 [ | 0.18716 | 6.98902 | 31.4283 | 5.44E-07 | 1.66E-05 |
| MLH1 | 3 | 3.7E+07 | 3.7E+07 | + | 3532 | mutL homolog | 0.23581 | 6.04938 | 31.3687 | 5.55E-07 | 1.69E-05 |
| SUN1 | 7 | 816615 | 896435 | + | 11850 | Sad1 and UNC | 0.19471 | 6.86343 | 31.2979 | 5.69E-07 | 1.73E-05 |
| TPST2 | 22 | 2.7E+07 | 2.7E+07 | - | 6913 | tyrosylprotein | -0.2079 | 6.45608 | 31.2645 | 5.75E-07 | 1.74E-05 |
| IGSF8 | 1 | 1.6E+08 | 1.6E+08 | - | 2592 | immunoglobul | -0.1927 | 6.9061 | 31.1663 | 5.95E-07 | 1.80E-05 |
| IGDCC3 | 15 | 6.5E+07 | 6.5E+07 | - | 5514 | immunoglobul | 0.17942 | 7.12985 | 31.1601 | 5.96E-07 | 1.80E-05 |
| KRT18 | 12 | 5.3E+07 | 5.3E+07 | + | 3069 | keratin 18 [Sou | -0.1442 | 8.63269 | 31.0783 | 6.13E-07 | 1.85E-05 |
| LCK | 1 | 3.2E+07 | 3.2E+07 | + | 3339 | LCK proto-onc | 0.20312 | 6.65637 | 31.0669 | 6.15E-07 | 1.85E-05 |
| MYH14 | 19 | 5E+07 | 5E+07 | + | 10392 | myosin heavy | -0.2348 | 6.0924 | 31.0447 | 6.20E-07 | 1.86E-05 |
| ROR1 | 1 | 6.4E+07 | 6.4E+07 | + | 6720 | receptor tyrosi | 0.18713 | 6.92942 | 31.0342 | 6.22E-07 | 1.86E-05 |
| MYBL2 | 20 | 4.4E+07 | 4.4E+07 | + | 2776 | MYB proto-onc | -0.1588 | 8.07829 | 30.9947 | 6.31E-07 | 1.88E-05 |
| NACC1 | 19 | 1.3E+07 | 1.3E+07 | + | 4736 | nucleus accum | -0.1487 | 8.3804 | 30.9677 | 6.36E-07 | 1.90E-05 |
| USP32 | 17 | 6E+07 | 6E+07 | - | 9522 | ubiquitin speci | -0.1757 | 7.18154 | 30.9266 | 6.45E-07 | 1.92E-05 |
| DSC2 | 18 | 3.1E+07 | 3.1E+07 | - | 12505 | desmocollin 2 | -0.2524 | 5.89046 | 30.9239 | 6.46E-07 | 1.92E-05 |
| ARHGAP2 | 18 | 6729718 | 6915716 | + | 11910 | Rho GTPase ac | 0.35341 | 4.71348 | 30.7729 | 6.80E-07 | 2.01E-05 |
| TSPAN5 | 4 | 9.8E+07 | 9.9E+07 | - | 7412 | tetraspanin 5 [ | 0.30205 | 5.2555 | 30.744 | 6.87E-07 | 2.03E-05 |
| UTF1 | 10 | 1.3E+08 | 1.3E+08 | + | 1214 | undifferentiate | -0.3486 | 4.91056 | 30.7361 | 6.89E-07 | 2.03E-05 |
| SALL2 | 14 | 2.2E+07 | 2.2E+07 | - | 5383 | spalt like trans | -0.1383 | 8.83902 | 30.7194 | 6.93E-07 | 2.04E-05 |
| SCNN1A | 12 | 6346843 | 6377730 | - | 5876 | sodium channe | -0.1861 | 7.05294 | 30.7167 | 6.93E-07 | 2.04E-05 |
| TRIP10 | 19 | 6737925 | 6751530 | + | 3986 | thyroid hormo | -0.2043 | 6.5587 | 30.6724 | 7.04E-07 | 2.06E-05 |
| YWHAB | 20 | 4.5E+07 | 4.5E+07 | + | 7907 | tyrosine 3-mor | -0.1412 | 8.6925 | 30.646 | 7.10E-07 | 2.08E-05 |
| MGAT4C | 12 | 8.6E+07 | 8.7E+07 | - | 27763 | MGAT4 family | 0.88181 | 1.98172 | 30.6264 | 7.15E-07 | 2.09E-05 |
| HIST1H2B | 6 | 2.7E+07 | 2.7E+07 | - | 381 | histone cluster | -0.5733 | 3.43756 | 31.0015 | 7.17E-07 | 2.09E-05 |
| BTAF1 | 10 | 9.2E+07 | 9.2E+07 | + | 7581 | B-TFIID TATA-b | 0.18365 | 6.98903 | 30.5471 | 7.35E-07 | 2.14E-05 |
| ARMCX1 | X | 1E+08 | 1E+08 | + | 2141 | armadillo repe | -0.2931 | 5.42726 | 30.499 | 7.47E-07 | 2.17E-05 |

| | | | | | | | | | | | |
|---|---|---|---|---|---|---|---|---|---|---|---|
| ACBD7 | 10 | 1.5E+07 | 1.5E+07 | - | 3546 | acyl-CoA bindi | -0.4308 | 4.18995 | 30.492 | 7.49E-07 | 2.17E-05 |
| CDYL2 | 16 | 8.1E+07 | 8.1E+07 | - | 8825 | chromodomai | -0.5807 | 3.19916 | 30.3503 | 7.86E-07 | 2.27E-05 |
| PTMA | 2 | 2.3E+08 | 2.3E+08 | + | 3378 | prothymosin a | -0.1187 | 10.7422 | 30.3422 | 7.88E-07 | 2.27E-05 |
| BST2 | 19 | 1.7E+07 | 1.7E+07 | - | 1101 | bone marrow s | 0.25608 | 5.7801 | 30.238 | 8.17E-07 | 2.35E-05 |
| ZFHX4 | 8 | 7.7E+07 | 7.7E+07 | + | 15875 | zinc finger hon | 0.75515 | 2.49136 | 30.2305 | 8.19E-07 | 2.35E-05 |
| SHISA9 | 16 | 1.3E+07 | 1.3E+07 | + | 10180 | shisa family m | -0.2694 | 5.58866 | 30.1205 | 8.50E-07 | 2.44E-05 |
| HCFC1 | X | 1.5E+08 | 1.5E+08 | - | 9011 | host cell facto | -0.1528 | 8.02098 | 30.0763 | 8.63E-07 | 2.47E-05 |
| ANGPT1 | 8 | 1.1E+08 | 1.1E+08 | - | 7128 | angiopoietin 1 | 1.02087 | 1.61107 | 30.0646 | 8.67E-07 | 2.48E-05 |
| UQCRC2 | 16 | 2.2E+07 | 2.2E+07 | + | 5217 | ubiquinol-cyto | -0.1623 | 7.52663 | 30.0568 | 8.69E-07 | 2.48E-05 |
| TGIF2 | 20 | 3.7E+07 | 3.7E+07 | + | 3973 | TGFB induced | -0.232 | 6.18121 | 30.0262 | 8.80E-07 | 2.50E-05 |
| SSB | 2 | 1.7E+08 | 1.7E+08 | + | 5054 | Sjogren syndro | -0.1498 | 8.14417 | 29.9814 | 8.92E-07 | 2.53E-05 |
| BTBD2 | 19 | 1985438 | 2034881 | - | 5944 | BTB domain co | 0.17027 | 7.28941 | 29.9369 | 9.06E-07 | 2.57E-05 |
| SATB1 | 3 | 1.8E+07 | 1.8E+07 | - | 13419 | SATB homeob | 0.25444 | 5.81719 | 29.9056 | 9.16E-07 | 2.59E-05 |
| BOK | 2 | 2.4E+08 | 2.4E+08 | + | 2680 | BCL2 family ap | 0.28411 | 5.49736 | 29.898 | 9.18E-07 | 2.59E-05 |
| DENND1C | 19 | 6467207 | 6482557 | - | 4647 | DENN domain | -0.3748 | 4.55113 | 29.8975 | 9.18E-07 | 2.59E-05 |
| DYNLL2 | 17 | 5.8E+07 | 5.8E+07 | + | 6807 | dynein light ch | -0.1607 | 7.67587 | 29.8854 | 9.22E-07 | 2.59E-05 |
| PMEL | 12 | 5.6E+07 | 5.6E+07 | - | 3901 | premelanosom | -0.1946 | 7.04277 | 30.6756 | 9.25E-07 | 2.60E-05 |
| NID1 | 1 | 2.4E+08 | 2.4E+08 | - | 5811 | nidogen 1 [Sou | 0.32666 | 6.85894 | 50.8631 | 9.28E-07 | 2.60E-05 |
| PLPP3 | 1 | 5.6E+07 | 5.7E+07 | - | 5272 | phospholipid p | -0.2629 | 5.7116 | 29.8561 | 9.31E-07 | 2.60E-05 |
| B4GALT6 | 18 | 3.2E+07 | 3.2E+07 | - | 5306 | beta-1,4-galac | -0.2407 | 5.91377 | 29.8105 | 9.46E-07 | 2.64E-05 |
| SAAL1 | 11 | 1.8E+07 | 1.8E+07 | - | 3066 | serum amyloid | -0.2862 | 5.54558 | 29.9498 | 9.65E-07 | 2.69E-05 |
| NIPSNAP1 | 22 | 3E+07 | 3E+07 | - | 3067 | nipsnap homo | 0.16552 | 7.46574 | 29.6845 | 9.88E-07 | 2.75E-05 |
| TLE2 | 19 | 2997639 | 3047635 | - | 4150 | TLE family mer | 0.42432 | 4.04088 | 29.6278 | 1.01E-06 | 2.80E-05 |
| TRIB2 | 2 | 1.3E+07 | 1.3E+07 | + | 4588 | tribbles pseud | 0.223 | 6.18609 | 29.6173 | 1.01E-06 | 2.80E-05 |
| SAMM50 | 22 | 4.4E+07 | 4.4E+07 | + | 6410 | SAMM50 sorti | -0.1987 | 6.5438 | 29.5591 | 1.03E-06 | 2.85E-05 |
| FOXJ1 | 17 | 7.6E+07 | 7.6E+07 | - | 2587 | forkhead box J | 0.48037 | 3.75304 | 29.5247 | 1.04E-06 | 2.88E-05 |
| DHFR | 5 | 8.1E+07 | 8.1E+07 | - | 4411 | dihydrofolate | 0.21742 | 6.18724 | 29.4915 | 1.06E-06 | 2.91E-05 |
| PAFAH1B | 11 | 1.2E+08 | 1.2E+08 | + | 7084 | platelet activa | -0.1786 | 6.98729 | 29.4801 | 1.06E-06 | 2.91E-05 |
| OLFM2 | 19 | 9853718 | 9936552 | - | 2500 | olfactomedin 2 | -0.1896 | 6.81924 | 29.4716 | 1.06E-06 | 2.91E-05 |
| SIPA1L1 | 14 | 7.1E+07 | 7.2E+07 | + | 12412 | signal induced | 0.18247 | 6.90788 | 29.4684 | 1.06E-06 | 2.91E-05 |
| CTTNBP2 | 7 | 1.2E+08 | 1.2E+08 | - | 6969 | cortactin bindi | 0.32511 | 5.03003 | 29.4003 | 1.09E-06 | 2.97E-05 |
| TRIM14 | 9 | 9.8E+07 | 9.8E+07 | - | 5481 | tripartite moti | -0.3127 | 5.0701 | 29.395 | 1.09E-06 | 2.97E-05 |
| LBH | 2 | 3E+07 | 3E+07 | + | 4532 | limb bud and h | -0.3086 | 5.15703 | 29.3724 | 1.10E-06 | 2.99E-05 |
| NIPAL1 | 4 | 4.8E+07 | 4.8E+07 | + | 5983 | NIPA like doma | -0.6132 | 3.18809 | 29.6975 | 1.14E-06 | 3.09E-05 |
| SKIL | 3 | 1.7E+08 | 1.7E+08 | + | 7782 | SKI like proto- | -0.1701 | 7.31827 | 29.2621 | 1.14E-06 | 3.09E-05 |
| CDC42EP4 | 17 | 7.3E+07 | 7.3E+07 | - | 4837 | CDC42 effecto | -0.3518 | 4.70512 | 29.0891 | 1.21E-06 | 3.27E-05 |
| LYAR | 4 | 4267701 | 4290154 | - | 2304 | Ly1 antibody r | -0.2421 | 5.81982 | 29.0041 | 1.25E-06 | 3.36E-05 |
| HIF3A | 19 | 4.6E+07 | 4.6E+07 | + | 8371 | hypoxia induci | -0.3777 | 4.43645 | 29.0028 | 1.25E-06 | 3.36E-05 |
| KLF6 | 10 | 3775996 | 3785281 | - | 6139 | Kruppel like fa | -0.3163 | 5.03461 | 28.9948 | 1.26E-06 | 3.37E-05 |
| LMNB2 | 19 | 2427638 | 2456959 | - | 5395 | lamin B2 [Sour | 0.12943 | 9.11397 | 28.9562 | 1.27E-06 | 3.40E-05 |
| DLD | 7 | 1.1E+08 | 1.1E+08 | + | 5102 | dihydrolipoam | -0.1995 | 6.6357 | 28.9439 | 1.28E-06 | 3.41E-05 |
| ISOC1 | 5 | 1.3E+08 | 1.3E+08 | + | 1942 | isochorismatas | 0.30398 | 5.29915 | 29.2334 | 1.30E-06 | 3.47E-05 |
| GPI | CHR19_2_ | 3.4E+07 | 3.4E+07 | + | 10680 | glucose-6-phos | -0.1458 | 8.09371 | 28.8571 | 1.32E-06 | 3.50E-05 |

| | | | | | | | | | | |
|---|---|---|---|---|---|---|---|---|---|---|
| MMP25 | 16 | 3046561 | 3060726 | + | 4246 | matrix metallo | 0.30512 | 5.09859 | 28.8175 | 1.34E-06 | 3.55E-05 |
| ARFGEF3 | 6 | 1.4E+08 | 1.4E+08 | + | 14859 | ARFGEF family | 0.22748 | 6.04374 | 28.7876 | 1.35E-06 | 3.58E-05 |
| PRTG | 15 | 5.6E+07 | 5.6E+07 | - | 14630 | protogenin [So | 0.61425 | 2.9145 | 28.7462 | 1.37E-06 | 3.62E-05 |
| LINC0020 | 21 | 4.5E+07 | 4.5E+07 | + | 7619 | long intergenic | -0.3489 | 4.84502 | 28.9915 | 1.38E-06 | 3.64E-05 |
| CDK4 | 12 | 5.8E+07 | 5.8E+07 | - | 3094 | cyclin depende | 0.18138 | 6.88693 | 28.6989 | 1.39E-06 | 3.66E-05 |
| RNF130 | 5 | 1.8E+08 | 1.8E+08 | - | 12817 | ring finger prot | 0.19259 | 6.61491 | 28.6977 | 1.39E-06 | 3.66E-05 |
| IDH2 | 15 | 9E+07 | 9E+07 | - | 2814 | isocitrate dehy | 0.20358 | 6.48758 | 28.6446 | 1.42E-06 | 3.71E-05 |
| PLEKHO1 | 1 | 1.5E+08 | 1.5E+08 | + | 6629 | pleckstrin hom | 0.30561 | 5.09468 | 28.6281 | 1.43E-06 | 3.71E-05 |
| PDHA1 | X | 1.9E+07 | 1.9E+07 | + | 4493 | pyruvate dehy | -0.1864 | 6.75533 | 28.6277 | 1.43E-06 | 3.71E-05 |
| ZC3H15 | 2 | 1.9E+08 | 1.9E+08 | + | 3026 | zinc finger CCC | -0.2028 | 6.44564 | 28.6094 | 1.44E-06 | 3.73E-05 |
| EIF3E | 8 | 1.1E+08 | 1.1E+08 | - | 6005 | eukaryotic tran | -0.1432 | 8.45171 | 28.5248 | 1.48E-06 | 3.84E-05 |
| GNAL | 18 | 1.2E+07 | 1.2E+07 | + | 8137 | G protein subu | -0.3507 | 4.66895 | 28.4479 | 1.52E-06 | 3.93E-05 |
| DLGAP3 | 1 | 3.5E+07 | 3.5E+07 | - | 4042 | DLG associated | -0.3644 | 5.08159 | 31.7897 | 1.54E-06 | 3.97E-05 |
| NRP2 | 2 | 2.1E+08 | 2.1E+08 | + | 13896 | neuropilin 2 [S | 0.5109 | 5.04824 | 45.023 | 1.54E-06 | 3.97E-05 |
| GNAO1 | 16 | 5.6E+07 | 5.6E+07 | + | 13542 | G protein subu | -0.2887 | 5.37526 | 28.4023 | 1.54E-06 | 3.97E-05 |
| TRAPPC10 | 21 | 4.4E+07 | 4.4E+07 | + | 11951 | trafficking prot | -0.2389 | 5.86358 | 28.323 | 1.59E-06 | 4.07E-05 |
| CENPH | 5 | 6.9E+07 | 6.9E+07 | + | 2218 | centromere pr | 0.22129 | 6.0574 | 28.2759 | 1.61E-06 | 4.12E-05 |
| PFN2 | 3 | 1.5E+08 | 1.5E+08 | - | 3709 | profilin 2 [Sour | 0.15081 | 7.80909 | 28.2607 | 1.62E-06 | 4.14E-05 |
| FOXI3 | 2 | 8.8E+07 | 8.8E+07 | - | 2804 | forkhead box I | -0.4617 | 3.84037 | 28.1938 | 1.66E-06 | 4.23E-05 |
| WIPF1 | 2 | 1.7E+08 | 1.7E+08 | - | 9121 | WAS/WASL int | 0.57959 | 3.10392 | 28.1873 | 1.67E-06 | 4.23E-05 |
| AXL | 19 | 4.1E+07 | 4.1E+07 | + | 5134 | AXL receptor ty | -0.231 | 5.98956 | 28.182 | 1.67E-06 | 4.23E-05 |
| RN7SL3 | 14 | 5E+07 | 5E+07 | - | 299 | RNA, 7SL, cyto | -0.9371 | 1.94647 | 28.5036 | 1.67E-06 | 4.23E-05 |
| GAS5 | 1 | 1.7E+08 | 1.7E+08 | - | 4842 | growth arrest s | 0.16565 | 7.64472 | 28.3463 | 1.81E-06 | 4.57E-05 |
| LSR | 19 | 3.5E+07 | 3.5E+07 | + | 2916 | lipolysis stimul | -0.1624 | 7.37819 | 27.9088 | 1.84E-06 | 4.63E-05 |
| HSP90B1 | 12 | 1E+08 | 1E+08 | + | 4695 | heat shock pro | -0.1177 | 9.93943 | 27.7583 | 1.94E-06 | 4.87E-05 |
| TCEA1 | 8 | 5.4E+07 | 5.4E+07 | - | 4342 | transcription e | -0.1617 | 7.43765 | 27.6916 | 1.98E-06 | 4.98E-05 |
| AMOT | X | 1.1E+08 | 1.1E+08 | - | 8502 | angiomotin [So | 0.40613 | 4.18114 | 27.6308 | 2.03E-06 | 5.07E-05 |
| SLC4A7 | 3 | 2.7E+07 | 2.7E+07 | - | 9616 | solute carrier f | 0.19168 | 6.5461 | 27.593 | 2.05E-06 | 5.13E-05 |
| RPL11 | 1 | 2.4E+07 | 2.4E+07 | + | 2659 | ribosomal prot | -0.1235 | 9.57295 | 27.5471 | 2.09E-06 | 5.21E-05 |
| MON1B | 16 | 7.7E+07 | 7.7E+07 | + | 6942 | MON1 homolo | -0.2344 | 5.88481 | 27.5303 | 2.10E-06 | 5.23E-05 |
| ATP5F1C | 10 | 7788147 | 7807815 | + | 2308 | ATP synthase F | -0.1661 | 7.13875 | 27.509 | 2.12E-06 | 5.26E-05 |
| KIF1A | 2 | 2.4E+08 | 2.4E+08 | - | 16695 | kinesin family | 0.12919 | 9.22441 | 27.4265 | 2.18E-06 | 5.41E-05 |
| CA14 | 1 | 1.5E+08 | 1.5E+08 | + | 3296 | carbonic anhyd | 0.26628 | 5.48676 | 27.3674 | 2.23E-06 | 5.51E-05 |
| DEK | 6 | 1.8E+07 | 1.8E+07 | - | 7249 | DEK proto-onc | -0.142 | 8.29748 | 27.2033 | 2.36E-06 | 5.83E-05 |
| PSPC1 | 13 | 2E+07 | 2E+07 | - | 4817 | paraspeckle co | -0.2021 | 6.37031 | 27.1867 | 2.37E-06 | 5.85E-05 |
| NUDT11 | X | 5.1E+07 | 5.1E+07 | - | 2381 | nudix hydrolas | -0.2994 | 5.09195 | 27.1707 | 2.39E-06 | 5.87E-05 |
| UPF2 | 10 | 1.2E+07 | 1.2E+07 | - | 5881 | UPF2, regulato | -0.2136 | 6.14284 | 27.1131 | 2.44E-06 | 5.99E-05 |
| DTNA | 18 | 3.4E+07 | 3.5E+07 | + | 12634 | dystrobrevin a | 0.27235 | 5.37137 | 27.0997 | 2.45E-06 | 5.99E-05 |
| ZSWIM6 | 5 | 6.1E+07 | 6.2E+07 | + | 5518 | zinc finger SWI | 0.40055 | 4.37525 | 27.7839 | 2.45E-06 | 5.99E-05 |
| TNRC6A | 16 | 2.5E+07 | 2.5E+07 | + | 12968 | trinucleotide r | -0.1553 | 7.53255 | 27.0561 | 2.49E-06 | 6.08E-05 |
| ARVCF | 22 | 2E+07 | 2E+07 | - | 5636 | ARVCF, delta c | 0.2416 | 5.78114 | 27.0502 | 2.49E-06 | 6.08E-05 |
| AHSA2P | 2 | 6.1E+07 | 6.1E+07 | + | 9083 | activator of HS | 0.31124 | 4.96707 | 27.0263 | 2.51E-06 | 6.12E-05 |
| ETV4 | 17 | 4.4E+07 | 4.4E+07 | - | 3515 | ETS variant 4 [S | -0.1523 | 7.72446 | 26.9899 | 2.55E-06 | 6.19E-05 |

| | | | | | | | | | | | |
|---|---|---|---|---|---|---|---|---|---|---|---|
| ITPKB | 1 | 2.3E+08 | 2.3E+08 | - | 7036 | inositol-trisphо | 0.3757 | 4.49915 | 27.1944 | 2.64E-06 | 6.40E-05 |
| MARCH8 | SCHR10_1 | 4.5E+07 | 4.5E+07 | - | 7516 | membrane ass | -0.3091 | 5.04265 | 26.8751 | 2.65E-06 | 6.42E-05 |
| TSPAN3 | 15 | 7.7E+07 | 7.7E+07 | - | 8506 | tetraspanin 3 [ | 0.17805 | 6.76888 | 26.8724 | 2.66E-06 | 6.42E-05 |
| CHAC1 | 15 | 4.1E+07 | 4.1E+07 | + | 2051 | ChaC glutathio | -0.3439 | 4.76229 | 26.9336 | 2.69E-06 | 6.49E-05 |
| DPYSL2 | 8 | 2.7E+07 | 2.7E+07 | + | 7296 | dihydropyrimi | -0.1593 | 7.40179 | 26.8315 | 2.69E-06 | 6.50E-05 |
| ZNF697 | 1 | 1.2E+08 | 1.2E+08 | - | 5579 | zinc finger pro | -0.2941 | 5.17778 | 26.8161 | 2.71E-06 | 6.51E-05 |
| CSNK2A2 | 16 | 5.8E+07 | 5.8E+07 | - | 7091 | casein kinase 2 | -0.1726 | 6.99088 | 26.811 | 2.71E-06 | 6.51E-05 |
| PCDHA10 | 5 | 1.4E+08 | 1.4E+08 | + | 15227 | protocadherin | 0.36271 | 4.50021 | 26.8098 | 2.72E-06 | 6.51E-05 |
| PPM1J | 1 | 1.1E+08 | 1.1E+08 | - | 2874 | protein phosp | -0.3653 | 4.36766 | 26.6589 | 2.87E-06 | 6.84E-05 |
| DHCR7 | 11 | 7.1E+07 | 7.1E+07 | - | 4160 | 7-dehydrochol | -0.1542 | 7.56564 | 26.6422 | 2.88E-06 | 6.87E-05 |
| SSBP1 | HSCHR7_1 | 1.4E+08 | 1.4E+08 | + | 4975 | single strandec | -0.1713 | 6.9641 | 26.63 | 2.90E-06 | 6.89E-05 |
| PIMREG | 17 | 6444441 | 6451469 | + | 2797 | PICALM intera | -0.1897 | 6.6711 | 26.609 | 2.92E-06 | 6.93E-05 |
| TBL1X | X | 9463295 | 9741037 | + | 9092 | transducin bet | 0.24827 | 5.71446 | 26.6053 | 2.92E-06 | 6.93E-05 |
| HERC5 | 4 | 8.8E+07 | 8.9E+07 | + | 4764 | HECT and RLD | -0.2857 | 5.20811 | 26.5996 | 2.93E-06 | 6.93E-05 |
| SLC9A3R2 | 16 | 2025356 | 2039026 | + | 3516 | SLC9A3 regula | 0.33642 | 4.74979 | 26.5972 | 2.93E-06 | 6.93E-05 |
| INAFM2 | 15 | 4E+07 | 4E+07 | + | 3052 | InaF motif con | 0.29063 | 5.3099 | 26.8251 | 3.00E-06 | 7.08E-05 |
| DOCK7 | 1 | 6.2E+07 | 6.3E+07 | - | 24872 | dedicator of cy | 0.23451 | 5.78714 | 26.4951 | 3.04E-06 | 7.16E-05 |
| GALNT1 | 18 | 3.6E+07 | 3.6E+07 | + | 5126 | polypeptide N- | -0.186 | 6.65507 | 26.4271 | 3.12E-06 | 7.30E-05 |
| SH2D4A | 8 | 1.9E+07 | 1.9E+07 | + | 4072 | SH2 domain co | -0.5136 | 3.35771 | 26.4268 | 3.12E-06 | 7.30E-05 |
| SPEN | 1 | 1.6E+07 | 1.6E+07 | + | 13447 | spen family tra | 0.17175 | 6.97416 | 26.4044 | 3.14E-06 | 7.35E-05 |
| LRRC3 | 21 | 4.4E+07 | 4.4E+07 | + | 5845 | leucine rich re | -0.3192 | 4.85977 | 26.3781 | 3.17E-06 | 7.41E-05 |
| TMEM121 | 22 | 1.7E+07 | 1.7E+07 | - | 5069 | transmembran | -0.4403 | 3.78901 | 26.3279 | 3.23E-06 | 7.53E-05 |
| COL9A1 | 6 | 7E+07 | 7E+07 | - | 7240 | collagen type I | 0.39943 | 4.1488 | 26.3218 | 3.24E-06 | 7.54E-05 |
| DGKB | 7 | 1.4E+07 | 1.5E+07 | - | 9491 | diacylglycerol | 1.42685 | 0.61939 | 26.2916 | 3.27E-06 | 7.59E-05 |
| TYMS | 18 | 657653 | 673578 | + | 2421 | thymidylate sy | 0.20344 | 6.21045 | 26.2764 | 3.29E-06 | 7.62E-05 |
| TCIRG1 | 11 | 6.8E+07 | 6.8E+07 | + | 4375 | T cell immune | 0.49081 | 3.57651 | 26.2602 | 3.31E-06 | 7.66E-05 |
| DMD | X | 3.1E+07 | 3.3E+07 | - | 18910 | dystrophin [So | -0.5703 | 3.03774 | 26.2462 | 3.33E-06 | 7.68E-05 |
| FOCAD | 9 | 2.1E+07 | 2.1E+07 | + | 10036 | focadhesin [So | 0.19747 | 6.32747 | 26.2431 | 3.33E-06 | 7.68E-05 |
| KLHL7 | 7 | 2.3E+07 | 2.3E+07 | + | 7066 | kelch like fami | 0.17869 | 6.693 | 26.2371 | 3.34E-06 | 7.68E-05 |
| ADRA2C | 4 | 3766348 | 3768526 | + | 2179 | adrenoceptor a | -0.3474 | 4.55216 | 26.2109 | 3.37E-06 | 7.74E-05 |
| SH3PXD2B | 5 | 1.7E+08 | 1.7E+08 | - | 8244 | SH3 and PX do | 0.16645 | 7.20038 | 26.1545 | 3.44E-06 | 7.89E-05 |
| SGPL1 | 10 | 7.1E+07 | 7.1E+07 | + | 7486 | sphingosine-1- | -0.195 | 6.37022 | 26.1084 | 3.49E-06 | 8.01E-05 |
| CSMD1 | 8 | 2935353 | 4994972 | - | 18705 | CUB and Sushi | -0.7072 | 2.40797 | 26.0853 | 3.52E-06 | 8.05E-05 |
| SEM1 | 7 | 9.6E+07 | 9.7E+07 | - | 11490 | SEM1, 26S pro | -0.2132 | 6.07837 | 26.084 | 3.53E-06 | 8.05E-05 |
| ZNF578 | 19 | 5.2E+07 | 5.3E+07 | + | 11940 | zinc finger pro | -0.4837 | 3.49397 | 26.0786 | 3.53E-06 | 8.06E-05 |
| GPD2 | 2 | 1.6E+08 | 1.6E+08 | + | 7165 | glycerol-3-pho | 0.20689 | 6.23968 | 26.0575 | 3.56E-06 | 8.11E-05 |
| LIMD2 | 17 | 6.4E+07 | 6.4E+07 | - | 4751 | LIM domain co | 0.22785 | 5.82284 | 25.9472 | 3.70E-06 | 8.42E-05 |
| ETS2 | 21 | 3.9E+07 | 3.9E+07 | + | 4497 | ETS proto-onco | -0.2315 | 5.79391 | 25.878 | 3.80E-06 | 8.62E-05 |
| NPM1 | 5 | 1.7E+08 | 1.7E+08 | + | 3828 | nucleophosmin | -0.1059 | 11.1269 | 25.8447 | 3.84E-06 | 8.71E-05 |
| ZDHHC13 | 11 | 1.9E+07 | 1.9E+07 | + | 7759 | zinc finger DHH | -0.3153 | 4.83158 | 25.8231 | 3.87E-06 | 8.77E-05 |
| ADGRL1 | 19 | 1.4E+07 | 1.4E+07 | - | 8487 | adhesion G pro | -0.1439 | 7.90643 | 25.78 | 3.94E-06 | 8.89E-05 |
| HIST1H1C | 6 | 2.6E+07 | 2.6E+07 | - | 642 | histone cluster | -0.5898 | 2.9445 | 25.7571 | 3.97E-06 | 8.95E-05 |
| RNF40 | 16 | 3.1E+07 | 3.1E+07 | + | 6523 | ring finger pro | 0.21584 | 5.99182 | 25.7409 | 3.99E-06 | 8.98E-05 |

| | | | | | | | | | | | |
|---|---|---|---|---|---|---|---|---|---|---|---|
| SPATS2 | 12 | 4.9E+07 | 5E+07 | + | 6840 | spermatogene | 0.2827 | 5.14155 | 25.7388 | 4.00E-06 | 8.98E-05 |
| AP001505 | 21 | 4.5E+07 | 4.5E+07 | + | 443 | novel transcrip | -0.6173 | 2.8304 | 25.7313 | 4.01E-06 | 8.99E-05 |
| MZT2B | 2 | 1.3E+08 | 1.3E+08 | + | 1815 | mitotic spindle | 0.32042 | 4.73412 | 25.7285 | 4.01E-06 | 8.99E-05 |
| FXYD5 | 19 | 3.5E+07 | 3.5E+07 | + | 2993 | FXYD domain c | -0.2135 | 6.11968 | 25.6616 | 4.11E-06 | 9.20E-05 |
| SUCLA2 | 13 | 4.8E+07 | 4.8E+07 | - | 8928 | succinate-CoA | -0.1994 | 6.39681 | 25.6385 | 4.14E-06 | 9.26E-05 |
| SNX2 | 5 | 1.2E+08 | 1.2E+08 | + | 8606 | sorting nexin 2 | -0.2027 | 6.2073 | 25.5743 | 4.24E-06 | 9.46E-05 |
| WFDC2 | 20 | 4.5E+07 | 4.5E+07 | + | 1335 | WAP four-disu | -0.3089 | 4.90625 | 25.5532 | 4.27E-06 | 9.52E-05 |
| BAZ2A | 12 | 5.7E+07 | 5.7E+07 | - | 10918 | bromodomain | -0.1384 | 8.09808 | 25.5057 | 4.35E-06 | 9.66E-05 |
| STOM | 9 | 1.2E+08 | 1.2E+08 | - | 3198 | stomatin [Sour | -0.3177 | 5.02065 | 26.0104 | 4.40E-06 | 9.75E-05 |
| DELE1 | 5 | 1.4E+08 | 1.4E+08 | + | 6000 | DAP3 binding c | 0.21575 | 5.97019 | 25.3744 | 4.56E-06 | 1.01E-04 |
| DACH1 | 13 | 7.1E+07 | 7.2E+07 | - | 5401 | dachshund fan | 1.14647 | 1.11689 | 25.3537 | 4.60E-06 | 1.01E-04 |
| DSP | 6 | 7541617 | 7586714 | + | 10048 | desmoplakin [S | 0.14197 | 7.99026 | 25.3352 | 4.63E-06 | 1.02E-04 |
| DPEP1 | 16 | 9E+07 | 9E+07 | + | 2737 | dipeptidase 1 [ | 0.59291 | 2.94033 | 25.1601 | 4.93E-06 | 1.09E-04 |
| RASD2 | 22 | 3.6E+07 | 3.6E+07 | + | 3447 | RASD family m | -0.5628 | 3.14172 | 25.1493 | 4.96E-06 | 1.09E-04 |
| BMP7 | 20 | 5.7E+07 | 5.7E+07 | - | 5425 | bone morphog | -0.3416 | 4.49259 | 25.1388 | 4.97E-06 | 1.09E-04 |
| PLBD2 | 12 | 1.1E+08 | 1.1E+08 | + | 5264 | phospholipase | -0.1951 | 6.33992 | 25.1337 | 4.98E-06 | 1.09E-04 |
| SBF2 | 11 | 9778667 | 1E+07 | - | 16767 | SET binding fac | 0.20228 | 6.2564 | 25.1174 | 5.01E-06 | 1.10E-04 |
| USO1 | 4 | 7.6E+07 | 7.6E+07 | + | 4588 | USO1 vesicle tr | -0.1335 | 8.39613 | 25.0892 | 5.06E-06 | 1.11E-04 |
| CHML | 1 | 2.4E+08 | 2.4E+08 | - | 7661 | CHM like, Rab | -0.1503 | 7.4583 | 25.0648 | 5.11E-06 | 1.11E-04 |
| PCLAF | 15 | 6.4E+07 | 6.4E+07 | - | 3741 | PCNA clamp as | 0.38796 | 4.10291 | 25.0571 | 5.12E-06 | 1.12E-04 |
| NMRK2 | 19 | 3933103 | 3942416 | + | 1141 | nicotinamide r | -0.2118 | 6.01741 | 25.0389 | 5.16E-06 | 1.12E-04 |
| TXNRD1 | 12 | 1E+08 | 1E+08 | + | 6710 | thioredoxin red | 0.14492 | 7.74186 | 24.9553 | 5.32E-06 | 1.15E-04 |
| COL1A2 | 7 | 9.4E+07 | 9.4E+07 | + | 11156 | collagen type I | 0.17039 | 7.28646 | 25.6504 | 5.40E-06 | 1.17E-04 |
| POLR2A | 17 | 7484366 | 7514616 | + | 8355 | RNA polymera | -0.1248 | 8.72931 | 24.9113 | 5.40E-06 | 1.17E-04 |
| PLK1 | 16 | 2.4E+07 | 2.4E+07 | + | 4760 | polo like kinas | -0.149 | 7.73088 | 24.892 | 5.44E-06 | 1.18E-04 |
| FSIP2 | 2 | 1.9E+08 | 1.9E+08 | + | 23585 | fibrous sheath | 0.53872 | 3.13138 | 24.8761 | 5.47E-06 | 1.18E-04 |
| NARS | 18 | 5.8E+07 | 5.8E+07 | - | 4089 | asparaginyl-tR | -0.1361 | 8.28448 | 24.8501 | 5.53E-06 | 1.19E-04 |
| NPC2 | 14 | 7.4E+07 | 7.4E+07 | - | 2749 | NPC intracellul | -0.1816 | 6.5845 | 24.8001 | 5.63E-06 | 1.21E-04 |
| USP43 | 17 | 9644698 | 9729687 | + | 5084 | ubiquitin speci | -0.4662 | 3.62054 | 24.7839 | 5.66E-06 | 1.21E-04 |
| TANC1 | 2 | 1.6E+08 | 1.6E+08 | + | 14915 | tetratricopepti | 0.35102 | 4.44241 | 24.7767 | 5.68E-06 | 1.21E-04 |
| KIAA1522 | 1 | 3.3E+07 | 3.3E+07 | + | 6020 | KIAA1522 [Sou | -0.1526 | 7.56205 | 24.7748 | 5.68E-06 | 1.21E-04 |
| CACNA1H | 16 | 1153106 | 1221771 | + | 8620 | calcium voltag | 0.20759 | 6.17012 | 24.7539 | 5.73E-06 | 1.22E-04 |
| REST | 4 | 5.7E+07 | 5.7E+07 | + | 7983 | RE1 silencing t | -0.197 | 6.2529 | 24.7498 | 5.73E-06 | 1.22E-04 |
| USF3 | 3 | 1.1E+08 | 1.1E+08 | - | 13738 | upstream trans | -0.2971 | 4.9361 | 24.7233 | 5.79E-06 | 1.23E-04 |
| TBC1D16 | 17 | 8E+07 | 8E+07 | - | 11579 | TBC1 domain f | 0.13805 | 8.12497 | 24.6792 | 5.88E-06 | 1.25E-04 |
| PGM1 | 1 | 6.4E+07 | 6.4E+07 | + | 4077 | phosphogluco | -0.1899 | 6.36625 | 24.6757 | 5.89E-06 | 1.25E-04 |
| RASSF4 | 10 | 4.5E+07 | 4.5E+07 | + | 9481 | Ras association | -0.4866 | 3.54318 | 24.6687 | 5.92E-06 | 1.25E-04 |
| UVSSA | 4 | 1347266 | 1395992 | + | 15773 | UV stimulated | 0.43286 | 3.78481 | 24.551 | 6.17E-06 | 1.30E-04 |
| GABBR1 | R6_MHC_ | 3E+07 | 3E+07 | - | 8468 | gamma-amino | 0.23245 | 5.75407 | 24.497 | 6.29E-06 | 1.33E-04 |
| CYFIP2 | 5 | 1.6E+08 | 1.6E+08 | + | 10498 | cytoplasmic FM | 0.17425 | 6.96686 | 24.7305 | 6.34E-06 | 1.34E-04 |
| DDHD2 | 8 | 3.8E+07 | 3.8E+07 | + | 10686 | DDHD domain | 0.31689 | 4.78173 | 24.4714 | 6.35E-06 | 1.34E-04 |
| CLU | 8 | 2.8E+07 | 2.8E+07 | - | 6616 | clusterin [Sour | -0.1506 | 7.48444 | 24.4401 | 6.43E-06 | 1.35E-04 |
| PYCR3 | HSCHR8_3 | 1.4E+08 | 1.4E+08 | - | 3947 | pyrroline-5-ca | 0.48428 | 3.34782 | 24.4368 | 6.43E-06 | 1.35E-04 |

| | | | | | | | | | | | |
|---|---|---|---|---|---|---|---|---|---|---|---|
| NBEA | 13 | 3.5E+07 | 3.6E+07 | + | 11949 | neurobeachin | 0.21868 | 5.95673 | 24.4163 | 6.48E-06 | 1.36E-04 |
| LRAT | 4 | 1.5E+08 | 1.5E+08 | + | 7535 | lecithin retinol | 0.37668 | 4.10094 | 24.3671 | 6.60E-06 | 1.38E-04 |
| CFAP298 | 21 | 3.3E+07 | 3.3E+07 | - | 5403 | cilia and flagell | -0.2818 | 5.34053 | 24.8487 | 6.63E-06 | 1.38E-04 |
| SIX4 | 14 | 6.1E+07 | 6.1E+07 | - | 6771 | SIX homeobox | 0.23307 | 5.65962 | 24.3189 | 6.72E-06 | 1.40E-04 |
| HSPG2 | 1 | 2.2E+07 | 2.2E+07 | - | 18118 | heparan sulfat | 0.15365 | 7.98958 | 25.4807 | 6.76E-06 | 1.41E-04 |
| NYNRIN | 14 | 2.4E+07 | 2.4E+07 | + | 7733 | NYN domain a | -0.2493 | 6.399 | 29.7588 | 7.01E-06 | 1.45E-04 |
| COL4A1 | 13 | 1.1E+08 | 1.1E+08 | - | 14662 | collagen type I | 0.14365 | 8.21324 | 24.6703 | 7.24E-06 | 1.50E-04 |
| EPB41L2 | 6 | 1.3E+08 | 1.3E+08 | - | 10320 | erythrocyte m | 0.14982 | 7.52284 | 24.0636 | 7.39E-06 | 1.53E-04 |
| ELL2 | 5 | 9.6E+07 | 9.6E+07 | - | 7108 | elongation fact | -0.2734 | 5.1937 | 24.0505 | 7.42E-06 | 1.53E-04 |
| CRNKL1 | 20 | 2E+07 | 2E+07 | - | 4567 | crooked neck p | -0.245 | 5.55634 | 24.0322 | 7.47E-06 | 1.54E-04 |
| UGT3A2 | 5 | 3.6E+07 | 3.6E+07 | - | 2683 | UDP glycosyltr | -0.3014 | 4.92018 | 24.022 | 7.50E-06 | 1.54E-04 |
| ACOX1 | 17 | 7.6E+07 | 7.6E+07 | - | 8286 | acyl-CoA oxida | -0.3071 | 4.93338 | 24.0721 | 7.54E-06 | 1.55E-04 |
| PKP2 | 12 | 3.3E+07 | 3.3E+07 | - | 7043 | plakophilin 2 [S | -0.2038 | 6.06781 | 23.9909 | 7.59E-06 | 1.55E-04 |
| VPS13A | 9 | 7.7E+07 | 7.7E+07 | + | 20196 | vacuolar prote | 0.17818 | 6.61261 | 23.9886 | 7.60E-06 | 1.55E-04 |
| ST8SIA3 | 18 | 5.7E+07 | 5.7E+07 | + | 10324 | ST8 alpha-N-ac | -0.4943 | 3.40244 | 23.9231 | 7.78E-06 | 1.59E-04 |
| NPDC1 | 9 | 1.4E+08 | 1.4E+08 | - | 2720 | neural prolifer | 0.26796 | 5.26024 | 23.8858 | 7.89E-06 | 1.61E-04 |
| B3GALT6 | 1 | 1232226 | 1235041 | + | 2816 | beta-1,3-galac | 0.31566 | 4.70817 | 23.854 | 7.99E-06 | 1.63E-04 |
| B4GALT4 | 3 | 1.2E+08 | 1.2E+08 | - | 6988 | beta-1,4-galac | -0.2271 | 5.7511 | 23.85 | 8.00E-06 | 1.63E-04 |
| SRSF12 | 6 | 8.9E+07 | 8.9E+07 | - | 4538 | serine and argi | 0.64835 | 2.48191 | 23.7613 | 8.27E-06 | 1.68E-04 |
| EXOC6B | 2 | 7.2E+07 | 7.3E+07 | - | 10197 | exocyst compl | 0.20846 | 6.03515 | 23.7373 | 8.34E-06 | 1.69E-04 |
| SESTD1 | 2 | 1.8E+08 | 1.8E+08 | - | 11405 | SEC14 and spe | 0.23592 | 5.60587 | 23.6904 | 8.49E-06 | 1.72E-04 |
| CLC | 19 | 4E+07 | 4E+07 | - | 631 | Charcot-Leyde | -0.6585 | 2.52566 | 23.6849 | 8.50E-06 | 1.72E-04 |
| CA4 | 17 | 6E+07 | 6E+07 | + | 2418 | carbonic anhy | -1.002 | 1.38856 | 23.6623 | 8.58E-06 | 1.73E-04 |
| RIOK2 | 5 | 9.7E+07 | 9.7E+07 | - | 5489 | RIO kinase 2 [S | -0.3073 | 4.80415 | 23.6607 | 8.58E-06 | 1.73E-04 |
| TCF7L1 | 2 | 8.5E+07 | 8.5E+07 | + | 4431 | transcription f | -0.1366 | 7.87548 | 23.6564 | 8.60E-06 | 1.73E-04 |
| NPTXR | 22 | 3.9E+07 | 3.9E+07 | - | 5830 | neuronal pent | -0.354 | 4.2998 | 23.6444 | 8.63E-06 | 1.74E-04 |
| FAM84B | 8 | 1.3E+08 | 1.3E+08 | - | 5734 | family with sec | -0.1891 | 6.29867 | 23.5896 | 8.81E-06 | 1.77E-04 |
| ITPR2 | 12 | 2.6E+07 | 2.7E+07 | - | 13909 | inositol 1,4,5-t | 0.18073 | 6.78947 | 23.9848 | 8.92E-06 | 1.79E-04 |
| TXNIP | 1 | 1.5E+08 | 1.5E+08 | - | 3625 | thioredoxin int | 1.09792 | 2.56879 | 37.4586 | 9.03E-06 | 1.81E-04 |
| NABP2 | 12 | 5.6E+07 | 5.6E+07 | + | 2201 | nucleic acid bi | 0.25143 | 5.40657 | 23.5206 | 9.04E-06 | 1.81E-04 |
| MRPL51 | 12 | 6491886 | 6493841 | - | 1388 | mitochondrial | -0.1694 | 6.75434 | 23.5143 | 9.06E-06 | 1.81E-04 |
| KRBA1 | 7 | 1.5E+08 | 1.5E+08 | + | 5337 | KRAB-A domai | 0.42149 | 3.82986 | 23.5044 | 9.10E-06 | 1.82E-04 |
| SUGT1 | 13 | 5.3E+07 | 5.3E+07 | + | 15102 | SGT1 homolog | -0.2031 | 6.07735 | 23.4899 | 9.15E-06 | 1.82E-04 |
| HMCN2 | 9 | 1.3E+08 | 1.3E+08 | + | 20363 | hemicentin 2 [ | 0.42633 | 3.67642 | 23.4561 | 9.26E-06 | 1.84E-04 |
| TAP1 | R6_MHC_ | 3.3E+07 | 3.3E+07 | - | 4805 | transporter 1, | -0.3514 | 4.40762 | 23.4446 | 9.30E-06 | 1.85E-04 |
| PPP1R3B | 8 | 9136255 | 9151574 | - | 5738 | protein phosp | -0.2496 | 5.49723 | 23.4223 | 9.38E-06 | 1.86E-04 |
| PTAR1 | 9 | 7E+07 | 7E+07 | - | 12252 | protein prenyl | -0.1606 | 7.34087 | 23.7975 | 9.39E-06 | 1.86E-04 |
| SNX9 | 6 | 1.6E+08 | 1.6E+08 | + | 4637 | sorting nexin 9 | 0.22747 | 5.76942 | 23.3959 | 9.47E-06 | 1.88E-04 |
| ABCA3 | 16 | 2275881 | 2340746 | - | 7804 | ATP binding ca | 0.33133 | 4.49105 | 23.369 | 9.57E-06 | 1.89E-04 |
| TRIM6 | 11 | 5596109 | 5612958 | + | 3964 | tripartite moti | -0.3418 | 4.53281 | 23.3668 | 9.58E-06 | 1.89E-04 |
| TSKU | 11 | 7.7E+07 | 7.7E+07 | + | 4013 | tsukushi, small | 0.25139 | 5.36039 | 23.3361 | 9.69E-06 | 1.91E-04 |
| HNRNPA2 | 7 | 2.6E+07 | 2.6E+07 | - | 9267 | heterogeneous | -0.1037 | 10.7971 | 23.3203 | 9.75E-06 | 1.92E-04 |
| TMEM245 | 9 | 1.1E+08 | 1.1E+08 | - | 9503 | transmembran | 0.14898 | 7.2953 | 23.3173 | 9.76E-06 | 1.92E-04 |

| | | | | | | | | | | | |
|---|---|---|---|---|---|---|---|---|---|---|---|
| AC011462 | 19 | 4.1E+07 | 4.1E+07 | - | 3238 | transforming g | -0.3457 | 4.42046 | 23.3062 | 9.80E-06 | 1.92E-04 |
| USP51 | X | 5.5E+07 | 5.5E+07 | - | 4407 | ubiquitin speci | -0.4497 | 3.53178 | 23.2917 | 9.85E-06 | 1.93E-04 |
| EPOP | 17 | 3.9E+07 | 3.9E+07 | - | 3719 | elongin BC and | 0.64305 | 2.47699 | 23.2732 | 9.92E-06 | 1.94E-04 |
| DLGAP4 | 20 | 3.6E+07 | 3.7E+07 | + | 8789 | DLG associated | -0.1996 | 6.08668 | 23.2396 | 1.00E-05 | 1.96E-04 |
| CBWD1 | 9 | 121038 | 179147 | - | 14170 | COBW domain | -0.3467 | 4.3479 | 23.2394 | 1.00E-05 | 1.96E-04 |
| CTXN1 | 19 | 7924491 | 7926135 | - | 1237 | cortexin 1 [Sou | 0.22566 | 5.68239 | 23.2021 | 1.02E-05 | 1.99E-04 |
| HSPA14 | 10 | 1.5E+07 | 1.5E+07 | + | 7108 | heat shock pro | -0.2718 | 5.86655 | 27.0736 | 1.03E-05 | 2.00E-04 |
| NBEAL2 | 3 | 4.7E+07 | 4.7E+07 | + | 12041 | neurobeachin | 0.19465 | 6.21546 | 23.0619 | 1.07E-05 | 2.09E-04 |
| VPS13B | 8 | 9.9E+07 | 1E+08 | + | 16716 | vacuolar prote | 0.26001 | 5.41407 | 23.1491 | 1.08E-05 | 2.09E-04 |
| CEP170B | 14 | 1E+08 | 1E+08 | + | 7211 | centrosomal p | 0.16651 | 6.76931 | 23.0188 | 1.09E-05 | 2.12E-04 |
| EXOC7 | 17 | 7.6E+07 | 7.6E+07 | - | 10116 | exocyst compl | 0.20238 | 6.0154 | 22.9636 | 1.11E-05 | 2.16E-04 |
| STAT6 | 12 | 5.7E+07 | 5.7E+07 | - | 7298 | signal transduc | 0.1973 | 6.12303 | 22.9602 | 1.12E-05 | 2.16E-04 |
| CCDC47 | 17 | 6.4E+07 | 6.4E+07 | - | 4354 | coiled-coil dom | -0.1768 | 6.53878 | 22.9314 | 1.13E-05 | 2.18E-04 |
| NEGR1 | 1 | 7.1E+07 | 7.2E+07 | - | 13208 | neuronal grow | 0.80605 | 1.85414 | 22.916 | 1.13E-05 | 2.19E-04 |
| FANCI | 15 | 8.9E+07 | 8.9E+07 | + | 7766 | FA complemen | 0.16088 | 6.89525 | 22.916 | 1.13E-05 | 2.19E-04 |
| DOCK1 | 10 | 1.3E+08 | 1.3E+08 | + | 8142 | dedicator of cy | 0.16279 | 6.85449 | 22.907 | 1.14E-05 | 2.19E-04 |
| FEZF1 | 7 | 1.2E+08 | 1.2E+08 | - | 2942 | FEZ family zinc | 1.11609 | 1.03667 | 22.9009 | 1.14E-05 | 2.19E-04 |
| HIVEP1 | 6 | 1.2E+07 | 1.2E+07 | + | 11005 | human immun | 0.30594 | 4.77124 | 22.845 | 1.17E-05 | 2.24E-04 |
| DMXL2 | 15 | 5.1E+07 | 5.2E+07 | - | 13807 | Dmx like 2 [Sou | 0.28152 | 5.01435 | 22.8262 | 1.17E-05 | 2.25E-04 |
| PTN | 7 | 1.4E+08 | 1.4E+08 | - | 1713 | pleiotrophin [S | 0.2764 | 5.01763 | 22.8056 | 1.18E-05 | 2.26E-04 |
| MT-CO1 | MT | 5904 | 7445 | + | 1542 | mitochondriall | -0.0913 | 12.7512 | 22.8053 | 1.18E-05 | 2.26E-04 |
| PLAU | 10 | 7.4E+07 | 7.4E+07 | + | 2949 | plasminogen a | 0.31974 | 4.80087 | 23.2155 | 1.19E-05 | 2.28E-04 |
| FKBP4 | 12 | 2794970 | 2805423 | + | 5163 | FKBP prolyl iso | -0.1215 | 8.75438 | 22.7744 | 1.20E-05 | 2.28E-04 |
| TMEM121 | 14 | 1.1E+08 | 1.1E+08 | + | 1924 | transmembran | 1.4499 | 0.40956 | 22.7661 | 1.20E-05 | 2.28E-04 |
| STIM1 | 11 | 3854527 | 4093210 | + | 7130 | stromal intera | -0.2841 | 4.90915 | 22.7551 | 1.21E-05 | 2.29E-04 |
| STARD9 | 15 | 4.3E+07 | 4.3E+07 | + | 20776 | StAR related li | 0.42902 | 3.62875 | 22.7524 | 1.21E-05 | 2.29E-04 |
| EHBP1L1 | 11 | 6.6E+07 | 6.6E+07 | + | 6733 | EH domain bin | 0.4171 | 3.73396 | 22.7513 | 1.21E-05 | 2.29E-04 |
| RPS24 | 10 | 7.8E+07 | 7.8E+07 | + | 7857 | ribosomal prot | -0.1141 | 9.36918 | 22.6417 | 1.26E-05 | 0.000238 |
| USP28 | 11 | 1.1E+08 | 1.1E+08 | - | 5719 | ubiquitin speci | -0.1459 | 7.4837 | 22.5899 | 1.28E-05 | 2.42E-04 |
| BAP1 | 3 | 5.2E+07 | 5.2E+07 | - | 4718 | BRCA1 associat | 0.16228 | 6.79758 | 22.5877 | 1.28E-05 | 2.42E-04 |
| AHCTF1 | 1 | 2.5E+08 | 2.5E+08 | - | 9598 | AT-hook conta | 0.16733 | 6.79439 | 22.5802 | 1.29E-05 | 2.43E-04 |
| SFT2D2 | 1 | 1.7E+08 | 1.7E+08 | + | 11217 | SFT2 domain c | -0.4006 | 3.85226 | 22.5799 | 1.29E-05 | 2.43E-04 |
| AC007326 | 22 | 1.9E+07 | 1.9E+07 | + | 1664 | novel transcrip | -0.7517 | 2.15397 | 22.547 | 1.30E-05 | 2.45E-04 |
| FAM157A | 3 | 2E+08 | 2E+08 | + | 7091 | family with sec | 0.63676 | 2.56589 | 22.5153 | 1.32E-05 | 2.48E-04 |
| TMEM167 | 5 | 8.3E+07 | 8.3E+07 | - | 4865 | transmembran | -0.135 | 7.83801 | 22.5134 | 1.32E-05 | 2.48E-04 |
| ZNF100 | SCHR19_2 | 2.2E+07 | 2.2E+07 | - | 6558 | zinc finger pro | -0.388 | 4.10487 | 22.6185 | 1.33E-05 | 2.49E-04 |
| APLP2 | 11 | 1.3E+08 | 1.3E+08 | + | 7463 | amyloid beta p | 0.11449 | 9.13565 | 22.4908 | 1.33E-05 | 2.49E-04 |
| NAB1 | 2 | 1.9E+08 | 1.9E+08 | + | 4887 | NGFI-A binding | 0.25385 | 5.2852 | 22.4881 | 1.33E-05 | 2.49E-04 |
| ADAMTS1 | 5 | 1.3E+08 | 1.3E+08 | + | 5392 | ADAM metallo | 0.30771 | 5.71663 | 28.5323 | 1.34E-05 | 2.49E-04 |
| SGK1 | 6 | 1.3E+08 | 1.3E+08 | - | 9992 | serum/glucoco | -0.3631 | 4.19117 | 22.4678 | 1.34E-05 | 2.50E-04 |
| SLC20A1 | 2 | 1.1E+08 | 1.1E+08 | + | 5493 | solute carrier f | -0.1664 | 6.75281 | 22.465 | 1.35E-05 | 2.50E-04 |
| MRS2 | 6 | 2.4E+07 | 2.4E+07 | + | 5133 | magnesium tra | -0.1542 | 7.06404 | 22.4377 | 1.36E-05 | 2.52E-04 |
| ZNF770 | 15 | 3.5E+07 | 3.5E+07 | - | 5439 | zinc finger pro | -0.1836 | 6.57177 | 22.6531 | 1.36E-05 | 2.52E-04 |

| | | | | | | | | | | | |
|---|---|---|---|---|---|---|---|---|---|---|---|
| CHST1 | 11 | 4.6E+07 | 4.6E+07 | - | 4666 | carbohydrate s | -0.7103 | 2.16549 | 22.4004 | 1.38E-05 | 2.56E-04 |
| PEG10 | 7 | 9.5E+07 | 9.5E+07 | + | 6689 | paternally exp | 0.13818 | 7.831 | 22.3832 | 1.39E-05 | 2.57E-04 |
| CYBA | 16 | 8.9E+07 | 8.9E+07 | - | 3253 | cytochrome b- | 0.17078 | 6.61101 | 22.3823 | 1.39E-05 | 2.57E-04 |
| PQBP1 | X | 4.9E+07 | 4.9E+07 | + | 2835 | polyglutamine | -0.2163 | 5.86861 | 22.3612 | 1.40E-05 | 2.58E-04 |
| IL13RA1 | X | 1.2E+08 | 1.2E+08 | + | 4650 | interleukin 13 | 0.35821 | 4.25143 | 22.3586 | 1.40E-05 | 2.58E-04 |
| AP001324 | 11 | 7.5E+07 | 7.5E+07 | - | 399 | ribosomal prot | 0.26182 | 5.14414 | 22.3087 | 1.43E-05 | 2.63E-04 |
| SH3D19 | 4 | 1.5E+08 | 1.5E+08 | - | 7789 | SH3 domain co | 0.15837 | 6.95527 | 22.2937 | 1.44E-05 | 2.64E-04 |
| RELB | 19 | 4.5E+07 | 4.5E+07 | + | 3016 | RELB proto-on | -0.3918 | 4.00131 | 22.2914 | 1.44E-05 | 2.64E-04 |
| MCM7 | 7 | 1E+08 | 1E+08 | - | 4386 | minichromoso | 0.11552 | 8.88811 | 22.2877 | 1.44E-05 | 2.64E-04 |
| BCL2L1 | 20 | 3.2E+07 | 3.2E+07 | - | 3314 | BCL2 like 1 [So | -0.189 | 6.18339 | 22.2409 | 1.46E-05 | 2.68E-04 |
| RBM47 | 4 | 4E+07 | 4.1E+07 | - | 8544 | RNA binding m | 0.16315 | 6.76237 | 22.2372 | 1.47E-05 | 2.68E-04 |
| SNRPD1 | 18 | 2.2E+07 | 2.2E+07 | + | 4862 | small nuclear r | -0.1663 | 6.85778 | 22.2265 | 1.47E-05 | 2.69E-04 |
| PFKM | 12 | 4.8E+07 | 4.8E+07 | + | 7246 | phosphofructo | -0.1576 | 6.92653 | 22.2228 | 1.47E-05 | 2.69E-04 |
| DNMBP | 10 | 1E+08 | 1E+08 | - | 8489 | dynamin bindi | -0.2304 | 5.74692 | 22.26 | 1.52E-05 | 2.76E-04 |
| PCDHGB7 | 5 | 1.4E+08 | 1.4E+08 | + | 4966 | protocadherin | 0.50772 | 3.15452 | 22.1435 | 1.52E-05 | 2.76E-04 |
| CSE1L | 20 | 4.9E+07 | 4.9E+07 | + | 3800 | chromosome s | -0.122 | 8.70581 | 22.0855 | 1.55E-05 | 2.82E-04 |
| HM13 | 20 | 3.2E+07 | 3.2E+07 | + | 10805 | histocompatib | -0.1721 | 6.56894 | 22.0822 | 1.56E-05 | 2.82E-04 |
| CHORDC1 | 11 | 9E+07 | 9E+07 | - | 11099 | cysteine and h | -0.1866 | 6.3572 | 22.0774 | 1.56E-05 | 2.82E-04 |
| VGLL4 | 3 | 1.2E+07 | 1.2E+07 | - | 7228 | vestigial like fa | 0.27351 | 5.08011 | 22.057 | 1.57E-05 | 2.84E-04 |
| MDN1 | 6 | 9E+07 | 9E+07 | - | 18674 | midasin AAA A | 0.12347 | 9.30924 | 22.9 | 1.57E-05 | 2.84E-04 |
| FYTTD1 | 3 | 2E+08 | 2E+08 | + | 7897 | forty-two-thre | 0.2563 | 5.2454 | 22.0395 | 1.58E-05 | 2.85E-04 |
| PKD1 | 16 | 2088710 | 2135898 | - | 18532 | polycystin 1, tr | 0.22841 | 5.66581 | 22.0395 | 1.58E-05 | 2.85E-04 |
| SLC9B2 | 4 | 1E+08 | 1E+08 | - | 5374 | solute carrier f | 0.58682 | 2.61776 | 21.9109 | 1.66E-05 | 2.99E-04 |
| CELF1 | 11 | 4.7E+07 | 4.8E+07 | - | 10452 | CUGBP Elav-lik | -0.136 | 7.63613 | 21.822 | 1.72E-05 | 3.09E-04 |
| KIZ | 20 | 2.1E+07 | 2.1E+07 | + | 5207 | kizuna centros | -0.2708 | 5.01524 | 21.8133 | 1.72E-05 | 3.10E-04 |
| SFXN5 | 2 | 7.3E+07 | 7.3E+07 | - | 11087 | sideroflexin 5 [ | 0.2438 | 5.3528 | 21.8024 | 1.73E-05 | 3.10E-04 |
| FAM129B | 9 | 1.3E+08 | 1.3E+08 | - | 4617 | family with seq | 0.14253 | 7.46174 | 21.8023 | 1.73E-05 | 3.10E-04 |
| BEND4 | 4 | 4.2E+07 | 4.2E+07 | - | 8850 | BEN domain co | -0.2151 | 5.96747 | 22.0054 | 1.73E-05 | 3.10E-04 |
| PNKP | 19 | 5E+07 | 5E+07 | - | 4545 | polynucleotide | 0.26062 | 5.11142 | 21.8008 | 1.73E-05 | 3.10E-04 |
| DLAT | 11 | 1.1E+08 | 1.1E+08 | + | 4831 | dihydrolipoam | -0.1523 | 7.00218 | 21.7833 | 1.74E-05 | 3.11E-04 |
| DLGAP5 | 14 | 5.5E+07 | 5.5E+07 | - | 3597 | DLG associated | -0.142 | 7.65959 | 21.761 | 1.76E-05 | 3.13E-04 |
| CLGN | 4 | 1.4E+08 | 1.4E+08 | - | 2822 | calmegin [Sour | -0.7281 | 2.1184 | 21.7485 | 1.77E-05 | 3.14E-04 |
| CSNK1G2 | 19 | 1941172 | 1981338 | + | 3781 | casein kinase 1 | 0.13429 | 7.70627 | 21.7113 | 1.79E-05 | 3.18E-04 |
| KPNA2 | 17 | 6.8E+07 | 6.8E+07 | + | 2963 | karyopherin su | -0.1087 | 9.42999 | 21.7058 | 1.80E-05 | 3.19E-04 |
| ABLIM1 | 10 | 1.1E+08 | 1.1E+08 | - | 10231 | actin binding L | 0.16957 | 6.72482 | 21.6651 | 1.82E-05 | 3.22E-04 |
| PRKAR2B | 7 | 1.1E+08 | 1.1E+08 | + | 4262 | protein kinase | -0.2084 | 6.05617 | 21.8445 | 1.84E-05 | 3.25E-04 |
| SNHG12 | 1 | 2.9E+07 | 2.9E+07 | - | 4266 | small nucleolar | 0.32708 | 4.41394 | 21.5932 | 1.88E-05 | 3.30E-04 |
| MEGF8 | 19 | 4.2E+07 | 4.2E+07 | + | 11372 | multiple EGF li | 0.14233 | 7.40189 | 21.5932 | 1.88E-05 | 3.30E-04 |
| FGF4 | 11 | 7E+07 | 7E+07 | - | 3165 | fibroblast grow | -0.7571 | 1.93309 | 21.5698 | 1.89E-05 | 3.33E-04 |
| MT-TE | MT | 14674 | 14742 | - | 69 | mitochondriall | -0.7984 | 1.80478 | 21.5606 | 1.90E-05 | 3.34E-04 |
| CLCN4 | X | 1E+07 | 1E+07 | + | 6789 | chloride voltag | -0.38 | 3.96524 | 21.5239 | 1.93E-05 | 3.38E-04 |
| TMSB15A | X | 1E+08 | 1E+08 | - | 634 | thymosin beta | 0.30867 | 4.59269 | 21.4847 | 1.96E-05 | 3.42E-04 |
| RFLNB | 17 | 439978 | 445939 | - | 3621 | refilin B [Sourc | -0.1888 | 6.41023 | 21.7827 | 1.97E-05 | 3.45E-04 |

| | | | | | | | | | | | |
|---|---|---|---|---|---|---|---|---|---|---|---|
| CCDC169 | 13 | 3.6E+07 | 3.6E+07 | - | 8837 | coiled-coil dom | -0.4301 | 3.64073 | 21.4219 | 2.00E-05 | 3.50E-04 |
| AP2A2 | 11 | 924894 | 1012245 | + | 9638 | adaptor relate | 0.20429 | 5.87309 | 21.3944 | 2.02E-05 | 3.53E-04 |
| COL1A1 | 17 | 5E+07 | 5E+07 | - | 9819 | collagen type I | -0.2499 | 8.49297 | 40.8361 | 2.04E-05 | 3.56E-04 |
| ABHD12B | 14 | 5.1E+07 | 5.1E+07 | + | 4551 | abhydrolase do | -0.6469 | 2.43208 | 21.3634 | 2.05E-05 | 3.56E-04 |
| SLC6A6 | 3 | 1.4E+07 | 1.4E+07 | + | 8254 | solute carrier f | -0.1332 | 7.93489 | 21.3358 | 2.07E-05 | 3.60E-04 |
| CPNE7 | 16 | 9E+07 | 9E+07 | + | 3058 | copine 7 [Sour | 0.68597 | 2.17014 | 21.3255 | 2.08E-05 | 3.61E-04 |
| STX1B | 16 | 3.1E+07 | 3.1E+07 | - | 5206 | syntaxin 1B [So | -0.3844 | 3.92482 | 21.2791 | 2.12E-05 | 3.67E-04 |
| ACVR1B | 12 | 5.2E+07 | 5.2E+07 | + | 5658 | activin A recep | 0.27016 | 4.9632 | 21.2532 | 2.14E-05 | 3.70E-04 |
| DIXDC1 | 11 | 1.1E+08 | 1.1E+08 | + | 8509 | DIX domain co | -0.2749 | 4.94502 | 21.2452 | 2.14E-05 | 3.71E-04 |
| SEPT6 | X | 1.2E+08 | 1.2E+08 | - | 6245 | septin 6 [Sourc | -0.2537 | 5.16136 | 21.1804 | 2.20E-05 | 3.80E-04 |
| ABCG2 | 4 | 8.8E+07 | 8.8E+07 | - | 5602 | ATP binding ca | -0.7592 | 2.03501 | 21.173 | 2.20E-05 | 3.80E-04 |
| NDFIP1 | 5 | 1.4E+08 | 1.4E+08 | + | 5747 | Nedd4 family i | 0.21671 | 5.83739 | 21.1973 | 2.23E-05 | 3.84E-04 |
| WDR27 | 6 | 1.7E+08 | 1.7E+08 | - | 12610 | WD repeat dor | 0.281 | 4.84111 | 21.12 | 2.25E-05 | 3.87E-04 |
| NAGA | 22 | 4.2E+07 | 4.2E+07 | - | 3706 | alpha-N-acetyl | 0.27413 | 4.89915 | 21.1022 | 2.27E-05 | 3.89E-04 |
| FTSJ1 | X | 4.8E+07 | 4.8E+07 | + | 2872 | FtsJ RNA meth | -0.2211 | 5.66223 | 21.0837 | 2.28E-05 | 3.92E-04 |
| RAB17 | 2 | 2.4E+08 | 2.4E+08 | - | 4761 | RAB17, membe | -0.1544 | 7.50967 | 22.0434 | 2.29E-05 | 3.92E-04 |
| BUB3 | 10 | 1.2E+08 | 1.2E+08 | + | 8175 | BUB3, mitotic c | 0.13996 | 7.56313 | 21.0412 | 2.32E-05 | 3.97E-04 |
| TEDC1 | 14 | 1.1E+08 | 1.1E+08 | + | 6698 | tubulin epsilon | 0.37608 | 3.94068 | 21.0402 | 2.32E-05 | 3.97E-04 |
| TKFC | 11 | 6.1E+07 | 6.1E+07 | + | 8136 | triokinase and | 0.25291 | 5.16504 | 21.024 | 2.34E-05 | 3.99E-04 |
| ANKRD12 | 18 | 9136228 | 9285985 | + | 12248 | ankyrin repeat | -0.2043 | 5.922 | 20.9874 | 2.37E-05 | 4.04E-04 |
| POMP | 13 | 2.9E+07 | 2.9E+07 | + | 1462 | proteasome m | -0.172 | 6.54268 | 20.9804 | 2.38E-05 | 4.05E-04 |
| FBL | 19 | 4E+07 | 4E+07 | - | 2138 | fibrillarin [Sour | -0.115 | 8.71187 | 20.9703 | 2.38E-05 | 4.06E-04 |
| DTL | 1 | 2.1E+08 | 2.1E+08 | + | 4923 | denticleless E3 | 0.16748 | 6.60599 | 20.9695 | 2.39E-05 | 4.06E-04 |
| STAG3L4 | 7 | 6.7E+07 | 6.7E+07 | + | 3192 | stromal antige | -0.5952 | 2.70413 | 20.9848 | 2.39E-05 | 4.06E-04 |
| DVL2 | 17 | 7225342 | 7234517 | - | 5321 | dishevelled seg | 0.21528 | 5.71809 | 20.938 | 2.41E-05 | 4.10E-04 |
| ZNF641 | 12 | 4.8E+07 | 4.8E+07 | - | 8474 | zinc finger prot | -0.501 | 3.11522 | 20.9016 | 2.45E-05 | 4.14E-04 |
| GRIN2A | 16 | 9753404 | 1E+07 | - | 16718 | glutamate ionc | -0.6521 | 2.29862 | 20.8944 | 2.46E-05 | 4.15E-04 |
| REC8 | 14 | 2.4E+07 | 2.4E+07 | + | 3844 | REC8 meiotic r | 0.33003 | 4.27793 | 20.8845 | 2.46E-05 | 4.16E-04 |
| KLHL17 | 1 | 960584 | 965719 | + | 3402 | kelch like famil | 0.33537 | 4.27094 | 20.8757 | 2.47E-05 | 4.17E-04 |
| GRIK3 | 1 | 3.7E+07 | 3.7E+07 | - | 10497 | glutamate ionc | -0.5476 | 2.78942 | 20.8731 | 2.48E-05 | 4.17E-04 |
| MAGED2 | X | 5.5E+07 | 5.5E+07 | + | 2943 | MAGE family m | -0.1418 | 7.39379 | 20.8507 | 2.50E-05 | 4.20E-04 |
| ELOF1 | 19 | 1.2E+07 | 1.2E+07 | - | 4173 | elongation fact | -0.1904 | 6.11165 | 20.8498 | 2.50E-05 | 4.20E-04 |
| PDLIM1 | 10 | 9.5E+07 | 9.5E+07 | - | 2227 | PDZ and LIM d | -0.1411 | 7.55598 | 20.8166 | 2.53E-05 | 4.25E-04 |
| LARGE1 | 22 | 3.3E+07 | 3.4E+07 | - | 10687 | LARGE xylosyl- | -0.1601 | 6.70634 | 20.7969 | 2.55E-05 | 4.28E-04 |
| RPL36AL | 14 | 5E+07 | 5E+07 | - | 746 | ribosomal prot | -0.1607 | 6.6847 | 20.7868 | 2.56E-05 | 4.29E-04 |
| JAM2 | 21 | 2.6E+07 | 2.6E+07 | + | 5947 | junctional adhe | -0.3969 | 3.7538 | 20.7471 | 2.60E-05 | 4.35E-04 |
| ARL5B | 10 | 1.9E+07 | 1.9E+07 | + | 7170 | ADP ribosylatic | -0.1521 | 6.94333 | 20.7175 | 2.63E-05 | 4.39E-04 |
| SOX11 | 2 | 5692384 | 5701385 | + | 9002 | SRY-box 11 [So | -0.1519 | 6.90898 | 20.6897 | 2.66E-05 | 4.44E-04 |
| RENBP | X | 1.5E+08 | 1.5E+08 | - | 2047 | renin binding p | -0.4237 | 3.61731 | 20.5988 | 2.75E-05 | 4.59E-04 |
| OPA3 | 19 | 4.6E+07 | 4.6E+07 | - | 9999 | OPA3, outer m | -0.2517 | 5.14589 | 20.5425 | 2.82E-05 | 4.69E-04 |
| COL5A1 | 9 | 1.3E+08 | 1.3E+08 | + | 11189 | collagen type V | 0.30693 | 4.85023 | 21.2842 | 2.82E-05 | 4.69E-04 |
| CRYZL1 | 21 | 3.4E+07 | 3.4E+07 | - | 6177 | crystallin zeta l | -0.3627 | 4.11241 | 20.5235 | 2.84E-05 | 4.71E-04 |
| ATP1A1 | 1 | 1.2E+08 | 1.2E+08 | + | 7277 | ATPase Na+/K- | -0.1082 | 9.23368 | 20.5131 | 2.85E-05 | 4.72E-04 |

| | | | | | | | | | | | |
|---|---|---|---|---|---|---|---|---|---|---|---|
| NPC1 | 18 | 2.4E+07 | 2.4E+07 | - | 9705 | NPC intracellul | -0.1869 | 6.1332 | 20.4847 | 2.88E-05 | 4.77E-04 |
| SPTBN1 | 2 | 5.4E+07 | 5.5E+07 | + | 18473 | spectrin beta, | 0.11173 | 9.20268 | 20.4788 | 2.89E-05 | 4.77E-04 |
| TP53BP1 | 15 | 4.3E+07 | 4.4E+07 | - | 12991 | tumor protein | 0.18967 | 6.27579 | 20.6874 | 2.89E-05 | 4.77E-04 |
| EXTL3 | 8 | 2.9E+07 | 2.9E+07 | + | 13220 | exostosin like g | 0.13101 | 7.91291 | 20.4531 | 2.91E-05 | 4.80E-04 |
| HMG20B | 19 | 3572777 | 3579088 | + | 3857 | high mobility g | 0.15323 | 6.90828 | 20.4382 | 2.93E-05 | 4.83E-04 |
| LEPR | 1 | 6.5E+07 | 6.6E+07 | + | 11370 | leptin receptor | 0.537 | 2.80673 | 20.3875 | 2.99E-05 | 4.92E-04 |
| LRRC8C | 1 | 9E+07 | 9E+07 | + | 11724 | leucine rich rep | 0.5827 | 2.63228 | 20.3837 | 2.99E-05 | 4.92E-04 |
| CCDC141 | 2 | 1.8E+08 | 1.8E+08 | - | 15446 | coiled-coil dom | -0.2897 | 4.68218 | 20.37 | 3.01E-05 | 4.94E-04 |
| AACS | 12 | 1.3E+08 | 1.3E+08 | + | 16039 | acetoacetyl-Co | -0.1681 | 6.50284 | 20.366 | 3.02E-05 | 4.94E-04 |
| RFX5 | 1 | 1.5E+08 | 1.5E+08 | - | 4586 | regulatory fact | 0.20553 | 5.89586 | 20.356 | 3.03E-05 | 4.95E-04 |
| LPL | 8 | 2E+07 | 2E+07 | + | 4131 | lipoprotein lipa | 0.45838 | 3.32956 | 20.292 | 3.10E-05 | 5.07E-04 |
| RESF1 | 12 | 3.2E+07 | 3.2E+07 | + | 6646 | retroelement s | -0.1574 | 6.80412 | 20.2849 | 3.11E-05 | 5.08E-04 |
| NR3C1 | 5 | 1.4E+08 | 1.4E+08 | - | 9161 | nuclear recept | 0.39344 | 3.69748 | 20.2491 | 3.16E-05 | 5.15E-04 |
| SEPT10 | 2 | 1.1E+08 | 1.1E+08 | - | 4251 | septin 10 [Sour | 0.15655 | 6.8154 | 20.2374 | 3.17E-05 | 5.17E-04 |
| EDIL3 | 5 | 8.4E+07 | 8.4E+07 | - | 5825 | EGF like repeat | 0.17801 | 6.35905 | 20.2126 | 3.20E-05 | 5.21E-04 |
| TMEM151 | 6 | 4.4E+07 | 4.4E+07 | + | 5459 | transmembran | -0.307 | 4.58905 | 20.2001 | 3.22E-05 | 5.23E-04 |
| TCF4 | 18 | 5.5E+07 | 5.6E+07 | - | 27828 | transcription fa | -0.1616 | 6.81201 | 20.2007 | 3.24E-05 | 5.27E-04 |
| CDC5L | 6 | 4.4E+07 | 4.4E+07 | + | 6241 | cell division cyc | -0.1701 | 6.62498 | 20.2641 | 3.26E-05 | 5.28E-04 |
| SPTBN2 | 11 | 6.7E+07 | 6.7E+07 | - | 10072 | spectrin beta, | 0.17733 | 6.29974 | 20.1569 | 3.27E-05 | 5.30E-04 |
| IFITM3 | 11 | 319676 | 327537 | - | 1151 | interferon indu | -0.1356 | 7.47402 | 20.1418 | 3.29E-05 | 5.33E-04 |
| CDT1 | 16 | 8.9E+07 | 8.9E+07 | + | 2760 | chromatin licen | 0.14269 | 7.13528 | 20.1225 | 3.32E-05 | 5.36E-04 |
| MAP4 | 3 | 4.8E+07 | 4.8E+07 | - | 14943 | microtubule as | 0.11877 | 8.79919 | 20.2843 | 3.32E-05 | 5.36E-04 |
| DENND5B | 12 | 3.1E+07 | 3.2E+07 | - | 11346 | DENN domain | 0.27788 | 5.0339 | 20.3873 | 3.34E-05 | 5.39E-04 |
| GPRC5C | 17 | 7.4E+07 | 7.4E+07 | + | 8005 | G protein-coup | -0.2735 | 4.86191 | 20.0984 | 3.35E-05 | 5.39E-04 |
| KCTD8 | 4 | 4.4E+07 | 4.4E+07 | - | 2712 | potassium cha | 0.66334 | 2.21025 | 20.0836 | 3.37E-05 | 5.42E-04 |
| GRIA4 | 11 | 1.1E+08 | 1.1E+08 | + | 10083 | glutamate ionc | 0.35012 | 4.11283 | 20.0695 | 3.39E-05 | 5.44E-04 |
| GPC6 | 13 | 9.3E+07 | 9.4E+07 | + | 7122 | glypican 6 [Sou | 0.72149 | 5.99945 | 76.3201 | 3.40E-05 | 5.45E-04 |
| RAB15 | 14 | 6.5E+07 | 6.5E+07 | - | 5015 | RAB15, membe | -0.1651 | 6.72099 | 20.1041 | 3.41E-05 | 5.47E-04 |
| CCNL2 | 1 | 1385711 | 1399335 | - | 6857 | cyclin L2 [Sour | 0.16499 | 6.54224 | 20.0355 | 3.43E-05 | 5.50E-04 |
| GPAA1 | 8 | 1.4E+08 | 1.4E+08 | + | 3268 | glycosylphosph | 0.21278 | 5.65009 | 19.9999 | 3.48E-05 | 5.57E-04 |
| HS6ST2 | X | 1.3E+08 | 1.3E+08 | - | 5002 | heparan sulfat | 0.41164 | 3.57351 | 19.9883 | 3.50E-05 | 5.59E-04 |
| PRR14L | 22 | 3.2E+07 | 3.2E+07 | - | 11809 | proline rich 14 | -0.1746 | 6.35746 | 19.9661 | 3.53E-05 | 5.63E-04 |
| NELL2 | 12 | 4.5E+07 | 4.5E+07 | - | 6424 | neural EGFL lik | 0.19353 | 6.18343 | 20.2327 | 3.54E-05 | 5.65E-04 |
| RASAL3 | 19 | 1.5E+07 | 1.5E+07 | - | 4654 | RAS protein ac | -0.6082 | 2.53729 | 19.9388 | 3.56E-05 | 5.67E-04 |
| DIP2A | 21 | 4.6E+07 | 4.7E+07 | + | 13758 | disco interactir | -0.2045 | 5.79757 | 19.9136 | 3.60E-05 | 5.72E-04 |
| PALM3 | 19 | 1.4E+07 | 1.4E+07 | - | 2533 | paralemmin 3 | -0.225 | 5.47335 | 19.9036 | 3.61E-05 | 5.74E-04 |
| BEND3 | 6 | 1.1E+08 | 1.1E+08 | - | 6839 | BEN domain co | -0.1963 | 5.94737 | 19.9004 | 3.62E-05 | 5.74E-04 |
| RAB25 | 1 | 1.6E+08 | 1.6E+08 | + | 1689 | RAB25, membe | 0.27594 | 4.91211 | 19.8809 | 3.65E-05 | 5.78E-04 |
| PGP | 16 | 2211997 | 2214807 | - | 2727 | phosphoglycol | 0.20496 | 5.79587 | 19.8786 | 3.65E-05 | 5.78E-04 |
| CAVIN3 | 11 | 6318946 | 6320532 | - | 1557 | caveolae assoc | 0.4834 | 3.05764 | 19.8235 | 3.73E-05 | 5.90E-04 |
| BCL7A | 12 | 1.2E+08 | 1.2E+08 | + | 6565 | BCL7A, BAF co | 0.29417 | 4.62523 | 19.8108 | 3.75E-05 | 5.92E-04 |
| MOV10 | 1 | 1.1E+08 | 1.1E+08 | + | 6509 | Mov10 RISC co | -0.1623 | 6.58175 | 19.8081 | 3.75E-05 | 5.92E-04 |
| EHBP1 | 2 | 6.3E+07 | 6.3E+07 | + | 8219 | EH domain bin | 0.18776 | 6.20404 | 19.8657 | 3.77E-05 | 5.94E-04 |

| | | | | | | | | | | | | |
|---|---|---|---|---|---|---|---|---|---|---|---|---|
| FABP5 | 8 | 8.1E+07 | 8.1E+07 | + | | 1904 | fatty acid bind | -0.2018 | 5.97698 | 19.8389 | 3.80E-05 | 5.99E-04 |
| SEC24B | 4 | 1.1E+08 | 1.1E+08 | + | | 5219 | SEC24 homolo | 0.21441 | 5.64356 | 19.7421 | 3.85E-05 | 6.05E-04 |
| PRRC2A | R6_MHC_ | 3.2E+07 | 3.2E+07 | + | | 9115 | proline rich co | -0.1016 | 9.62348 | 19.7395 | 3.86E-05 | 6.05E-04 |
| CBLN1 | 16 | 4.9E+07 | 4.9E+07 | - | | 2601 | cerebellin 1 pr | -0.6233 | 2.37084 | 19.7295 | 3.87E-05 | 6.07E-04 |
| IMPDH1 | 7 | 1.3E+08 | 1.3E+08 | - | | 3662 | inosine monop | 0.2153 | 5.6312 | 19.7105 | 3.90E-05 | 6.11E-04 |
| RNF187 | 1 | 2.3E+08 | 2.3E+08 | + | | 3386 | ring finger prot | -0.1731 | 6.32858 | 19.6993 | 3.92E-05 | 6.13E-04 |
| CXXC5 | 5 | 1.4E+08 | 1.4E+08 | + | | 3795 | CXXC finger pr | 0.14412 | 7.09341 | 19.6585 | 3.98E-05 | 6.22E-04 |
| NOP53 | 19 | 4.8E+07 | 4.8E+07 | + | | 5317 | NOP53 ribosor | 0.11786 | 8.23905 | 19.6237 | 4.04E-05 | 6.30E-04 |
| TIMM17B | X | 4.9E+07 | 4.9E+07 | - | | 2562 | translocase of | -0.1922 | 6.00613 | 19.6102 | 4.06E-05 | 6.32E-04 |
| LGALS3BP | 17 | 7.9E+07 | 7.9E+07 | - | | 3422 | galectin 3 bind | 0.19316 | 5.93714 | 19.6054 | 4.06E-05 | 6.32E-04 |
| C6orf48 | R6_MHC_ | 3.2E+07 | 3.2E+07 | + | | 1801 | chromosome 6 | 0.21719 | 5.58598 | 19.6039 | 4.07E-05 | 6.32E-04 |
| KIAA1109 | 4 | 1.2E+08 | 1.2E+08 | + | | 20470 | KIAA1109 [Sou | 0.15633 | 6.66872 | 19.6021 | 4.07E-05 | 6.32E-04 |
| FLT4 | 5 | 1.8E+08 | 1.8E+08 | - | | 8852 | fms related tyr | 0.34166 | 4.09086 | 19.5994 | 4.07E-05 | 6.32E-04 |
| KBTBD8 | 3 | 6.7E+07 | 6.7E+07 | + | | 4736 | kelch repeat ar | -0.2879 | 4.73302 | 19.5678 | 4.13E-05 | 6.39E-04 |
| ASAP2 | 2 | 9206765 | 9405683 | + | | 7257 | ArfGAP with SH | 0.28284 | 4.67773 | 19.5652 | 4.13E-05 | 6.39E-04 |
| LTBP1 | 2 | 3.3E+07 | 3.3E+07 | + | | 7348 | latent transfor | 0.1667 | 6.47851 | 19.5364 | 4.18E-05 | 6.46E-04 |
| GARS | 7 | 3.1E+07 | 3.1E+07 | + | | 3995 | glycyl-tRNA syr | -0.1226 | 8.28203 | 19.5116 | 4.22E-05 | 6.52E-04 |
| NECAP2 | 1 | 1.6E+07 | 1.6E+07 | + | | 5510 | NECAP endocy | 0.23604 | 5.25972 | 19.4729 | 4.28E-05 | 6.61E-04 |
| RMND5A | 2 | 8.7E+07 | 8.7E+07 | + | | 6686 | required for m | -0.1486 | 6.96456 | 19.4407 | 4.34E-05 | 6.69E-04 |
| MAN2C1 | 15 | 7.5E+07 | 7.5E+07 | - | | 9124 | mannosidase a | 0.26199 | 4.92126 | 19.4099 | 4.39E-05 | 6.76E-04 |
| SRSF6 | 20 | 4.3E+07 | 4.3E+07 | + | | 4621 | serine and argi | -0.1384 | 7.59014 | 19.563 | 4.41E-05 | 6.79E-04 |
| ATP6V1E1 | 22 | 1.8E+07 | 1.8E+07 | - | | 3143 | ATPase H+ trar | -0.1706 | 6.31881 | 19.3883 | 4.43E-05 | 6.80E-04 |
| GNL2 | 1 | 3.8E+07 | 3.8E+07 | - | | 5747 | G protein nucle | -0.1415 | 7.12377 | 19.3763 | 4.45E-05 | 6.82E-04 |
| FAM217B | 20 | 6E+07 | 6E+07 | + | | 5373 | family with sec | -0.2682 | 4.92622 | 19.3498 | 4.50E-05 | 6.89E-04 |
| COX6B1 | 19 | 3.6E+07 | 3.6E+07 | + | | 1548 | cytochrome c c | -0.1472 | 7.00683 | 19.3481 | 4.50E-05 | 6.89E-04 |
| NDUFV3 | 21 | 4.3E+07 | 4.3E+07 | + | | 6342 | NADH:ubiquin | -0.2158 | 5.5627 | 19.3286 | 4.53E-05 | 6.93E-04 |
| IDH3G | X | 1.5E+08 | 1.5E+08 | - | | 2717 | isocitrate dehy | 0.31075 | 4.49461 | 19.3146 | 4.56E-05 | 6.96E-04 |
| RTL5 | X | 7.2E+07 | 7.2E+07 | - | | 4779 | retrotransposo | 0.65868 | 2.18166 | 19.3071 | 4.57E-05 | 6.98E-04 |
| CLDN19 | 1 | 4.3E+07 | 4.3E+07 | - | | 3602 | claudin 19 [Sou | -0.3399 | 4.14979 | 19.2766 | 4.63E-05 | 7.05E-04 |
| CHST6 | 16 | 7.5E+07 | 7.5E+07 | - | | 8620 | carbohydrate s | -0.2519 | 5.06733 | 19.2599 | 4.66E-05 | 7.09E-04 |
| CD151 | 11 | 832887 | 839831 | + | | 3894 | CD151 molecu | 0.1547 | 6.66046 | 19.2518 | 4.67E-05 | 7.11E-04 |
| OLMALIN | 10 | 1E+08 | 1E+08 | + | | 7763 | oligodendrocy | -0.2724 | 4.82879 | 19.2491 | 4.68E-05 | 7.11E-04 |
| ZNF850 | 19 | 3.7E+07 | 3.7E+07 | - | | 7850 | zinc finger prot | -0.2574 | 5.11049 | 19.2402 | 4.70E-05 | 7.13E-04 |
| JPH1 | 8 | 7.4E+07 | 7.4E+07 | - | | 4590 | junctophilin 1 | 0.24058 | 5.28582 | 19.2038 | 4.76E-05 | 7.22E-04 |
| TET3G | TET3G | 1 | 2517 | + | | 2517 | Transactivator | -0.1149 | 9.08996 | 19.5183 | 4.78E-05 | 7.23E-04 |
| NDNF | 4 | 1.2E+08 | 1.2E+08 | - | | 3573 | neuron derived | 1.11244 | 0.8172 | 19.1968 | 4.78E-05 | 7.23E-04 |
| TTK | 6 | 8E+07 | 8E+07 | + | | 4579 | TTK protein kir | -0.1817 | 6.23942 | 19.144 | 4.88E-05 | 7.37E-04 |
| L3MBTL2 | 22 | 4.1E+07 | 4.1E+07 | + | | 6374 | L3MBTL2, poly | 0.25404 | 5.02215 | 19.1319 | 4.90E-05 | 7.40E-04 |
| TEK | 9 | 2.7E+07 | 2.7E+07 | + | | 4804 | TEK receptor ty | 0.22122 | 5.51469 | 19.1251 | 4.92E-05 | 7.40E-04 |
| DNAJC2 | 7 | 1E+08 | 1E+08 | - | | 6706 | DnaJ heat shoc | -0.2127 | 5.58104 | 19.1219 | 4.92E-05 | 7.40E-04 |
| L3MBTL3 | 6 | 1.3E+08 | 1.3E+08 | + | | 5113 | L3MBTL3, histo | 0.2512 | 5.16057 | 19.1202 | 4.93E-05 | 7.40E-04 |
| RHPN1 | 8 | 1.4E+08 | 1.4E+08 | + | | 4184 | rhophilin Rho G | 0.2969 | 4.49568 | 19.1186 | 4.93E-05 | 7.40E-04 |
| SLC35C1 | 11 | 4.6E+07 | 4.6E+07 | + | | 4505 | solute carrier f | -0.2681 | 4.88784 | 19.1174 | 4.93E-05 | 7.40E-04 |

| | | | | | | | | | | | |
|---|---|---|---|---|---|---|---|---|---|---|---|
| SOCS3 | 17 | 7.8E+07 | 7.8E+07 | - | 2734 | suppressor of | 0.26563 | 5.27074 | 20.0678 | 4.97E-05 | 7.46E-04 |
| THOP1 | 19 | 2785503 | 2815807 | + | 7421 | thimet oligope | 0.15469 | 6.63929 | 19.0603 | 5.04E-05 | 7.56E-04 |
| SNX17 | 2 | 2.7E+07 | 2.7E+07 | + | 3767 | sorting nexin 1 | 0.17843 | 6.175 | 19.018 | 5.13E-05 | 7.68E-04 |
| TNFRSF10 | 8 | 2.3E+07 | 2.3E+07 | - | 5010 | TNF receptor s | 0.14597 | 7.0579 | 19.0083 | 5.15E-05 | 7.70E-04 |
| ZDHHC20 | 13 | 2.1E+07 | 2.1E+07 | - | 5574 | zinc finger DHH | -0.1497 | 6.88712 | 18.9957 | 5.18E-05 | 7.73E-04 |
| NTN1 | 17 | 9021510 | 9244000 | + | 6415 | netrin 1 [Sourc | -0.3969 | 3.64557 | 18.9724 | 5.22E-05 | 7.79E-04 |
| GRPR | X | 1.6E+07 | 1.6E+07 | + | 1929 | gastrin releasir | 0.34627 | 4.2033 | 19.0907 | 5.31E-05 | 7.91E-04 |
| ADK | 10 | 7.4E+07 | 7.5E+07 | + | 3184 | adenosine kina | 0.26397 | 4.83949 | 18.9034 | 5.37E-05 | 8.00E-04 |
| FNBP4 | 11 | 4.8E+07 | 4.8E+07 | - | 7596 | formin binding | 0.17544 | 6.2593 | 18.8123 | 5.57E-05 | 8.27E-04 |
| EBP | X | 4.9E+07 | 4.9E+07 | + | 1768 | EBP, cholesten | -0.2459 | 5.18563 | 18.8112 | 5.57E-05 | 8.27E-04 |
| RAI2 | X | 1.8E+07 | 1.8E+07 | - | 2622 | retinoic acid in | -0.9289 | 1.34823 | 18.8096 | 5.57E-05 | 8.27E-04 |
| FZD7 | 2 | 2E+08 | 2E+08 | + | 3859 | frizzled class re | -0.1059 | 9.21406 | 18.8086 | 5.58E-05 | 8.27E-04 |
| GPSM1 | 9 | 1.4E+08 | 1.4E+08 | + | 5138 | G protein signa | 0.26046 | 5.13933 | 19.0551 | 5.58E-05 | 8.27E-04 |
| EPHA1 | 7 | 1.4E+08 | 1.4E+08 | - | 4697 | EPH receptor A | 0.12772 | 7.76906 | 18.7745 | 5.65E-05 | 8.37E-04 |
| PAK4 | 19 | 3.9E+07 | 3.9E+07 | + | 7115 | p21 (RAC1) act | -0.1445 | 6.95921 | 18.7133 | 5.79E-05 | 8.56E-04 |
| RPL13A | 19 | 4.9E+07 | 4.9E+07 | + | 2369 | ribosomal prot | 0.09224 | 10.9076 | 18.6753 | 5.88E-05 | 8.68E-04 |
| AL157817 | 13 | 5.1E+07 | 5.1E+07 | + | 790 | novel transcrip | -0.3696 | 3.80799 | 18.6736 | 5.89E-05 | 8.68E-04 |
| TMEM263 | 12 | 1.1E+08 | 1.1E+08 | + | 4768 | transmembran | -0.3527 | 4.13415 | 18.8261 | 5.89E-05 | 8.68E-04 |
| CCDC50 | 3 | 1.9E+08 | 1.9E+08 | + | 9183 | coiled-coil dom | 0.23102 | 5.41023 | 18.6406 | 5.96E-05 | 8.77E-04 |
| TSPYL1 | 6 | 1.2E+08 | 1.2E+08 | - | 7342 | TSPY like 1 [So | 0.15531 | 6.57144 | 18.6135 | 6.03E-05 | 8.86E-04 |
| EDRF1 | 10 | 1.3E+08 | 1.3E+08 | + | 7741 | erythroid diffe | 0.22021 | 5.54411 | 18.6048 | 6.05E-05 | 8.88E-04 |
| CDC42SE1 | 1 | 1.5E+08 | 1.5E+08 | - | 4184 | CDC42 small ef | 0.17093 | 6.36192 | 18.5526 | 6.18E-05 | 9.06E-04 |
| SEPT11 | 4 | 7.7E+07 | 7.7E+07 | + | 9787 | septin 11 [Sour | 0.13218 | 7.37205 | 18.4611 | 6.41E-05 | 9.38E-04 |
| EDEM2 | 20 | 3.5E+07 | 3.5E+07 | - | 1884 | ER degradation | -0.2835 | 4.64669 | 18.455 | 6.42E-05 | 9.39E-04 |
| NDRG3 | 20 | 3.7E+07 | 3.7E+07 | - | 3260 | NDRG family m | -0.2369 | 5.19968 | 18.4472 | 6.44E-05 | 9.41E-04 |
| SLC25A1 | 22 | 1.9E+07 | 1.9E+07 | - | 1999 | solute carrier f | -0.1248 | 7.6847 | 18.4388 | 6.47E-05 | 9.43E-04 |
| FKBP10 | 17 | 4.2E+07 | 4.2E+07 | + | 4273 | FKBP prolyl iso | 0.12236 | 7.87065 | 18.4308 | 6.49E-05 | 9.45E-04 |
| TDGF1 | 3 | 4.7E+07 | 4.7E+07 | + | 2489 | teratocarcinon | -0.1035 | 9.04132 | 18.4261 | 6.50E-05 | 9.46E-04 |
| GINS1 | 20 | 2.5E+07 | 2.5E+07 | + | 4130 | GINS complex | -0.1596 | 6.47531 | 18.4215 | 6.51E-05 | 9.47E-04 |
| HINT1 | 5 | 1.3E+08 | 1.3E+08 | - | 1936 | histidine triad | 0.1304 | 7.39875 | 18.3853 | 6.61E-05 | 9.60E-04 |
| PELP1 | 17 | 4669774 | 4704337 | - | 6577 | proline, glutam | -0.1215 | 8.05184 | 18.3829 | 6.61E-05 | 9.60E-04 |
| MT-TY | MT | 5826 | 5891 | - | 66 | mitochondriall | -0.4997 | 2.93294 | 18.3705 | 6.65E-05 | 9.63E-04 |
| RPL14 | 3 | 4E+07 | 4E+07 | + | 8626 | ribosomal prot | -0.1012 | 9.27483 | 18.3693 | 6.65E-05 | 9.63E-04 |
| UNC13A | 19 | 1.8E+07 | 1.8E+07 | - | 11188 | unc-13 homolo | 0.17984 | 6.13826 | 18.3166 | 6.79E-05 | 9.82E-04 |
| GNAI2 | 3 | 5E+07 | 5E+07 | + | 7405 | G protein subu | 0.1082 | 8.61259 | 18.3039 | 6.83E-05 | 9.86E-04 |
| CERS6 | 2 | 1.7E+08 | 1.7E+08 | + | 7241 | ceramide synth | 0.22472 | 5.41914 | 18.2948 | 6.85E-05 | 9.89E-04 |
| GAK | 4 | 849276 | 932373 | - | 15155 | cyclin G associ | 0.16164 | 6.54096 | 18.2797 | 6.89E-05 | 9.94E-04 |
| CPE | 4 | 1.7E+08 | 1.7E+08 | + | 3119 | carboxypeptid | 0.36014 | 3.89894 | 18.2756 | 6.90E-05 | 9.94E-04 |
| POLQ | 3 | 1.2E+08 | 1.2E+08 | - | 9382 | DNA polymera | 0.20956 | 5.65455 | 18.2688 | 6.92E-05 | 9.95E-04 |
| FAT3 | 11 | 9.2E+07 | 9.3E+07 | + | 19280 | FAT atypical ca | 0.17126 | 6.38093 | 18.3142 | 6.92E-05 | 9.95E-04 |
| BTF3 | 5 | 7.3E+07 | 7.4E+07 | + | 2467 | basic transcrip | -0.1208 | 7.92635 | 18.26 | 6.95E-05 | 9.97E-04 |
| GCAT | 22 | 3.8E+07 | 3.8E+07 | + | 2104 | glycine C-acety | 0.24453 | 5.17621 | 18.23 | 7.03E-05 | 0.001009 |
| CTPS1 | 1 | 4.1E+07 | 4.1E+07 | + | 11517 | CTP synthase 1 | 0.14137 | 7.15145 | 18.1664 | 7.22E-05 | 0.001034 |

| | | | | | | | | | | | |
|---|---|---|---|---|---|---|---|---|---|---|---|
| SOCS1 | 16 | 1.1E+07 | 1.1E+07 | - | 1796 | suppressor of | -0.2651 | 5.46122 | 20.2696 | 7.27E-05 | 0.00104 |
| FGFBP3 | 10 | 9.2E+07 | 9.2E+07 | - | 2545 | fibroblast grow | 0.1433 | 7.16024 | 18.2492 | 7.30E-05 | 0.001043 |
| PC | 11 | 6.7E+07 | 6.7E+07 | - | 7355 | pyruvate carbo | 0.31902 | 4.25707 | 18.1021 | 7.41E-05 | 0.001058 |
| PHF8 | X | 5.4E+07 | 5.4E+07 | - | 9015 | PHD finger pro | -0.1984 | 5.74357 | 18.0995 | 7.41E-05 | 0.001058 |
| LINC0064 | 21 | 3.4E+07 | 3.4E+07 | + | 14336 | long intergenic | -0.3783 | 3.67875 | 18.0529 | 7.55E-05 | 0.001077 |
| KDM4A | 1 | 4.4E+07 | 4.4E+07 | + | 5184 | lysine demethy | -0.1586 | 6.58203 | 18.0408 | 7.59E-05 | 0.00108 |
| RGS19 | 20 | 6.4E+07 | 6.4E+07 | - | 2028 | regulator of G | 0.36775 | 3.74011 | 18.0101 | 7.69E-05 | 0.001092 |
| HK1 | 10 | 6.9E+07 | 6.9E+07 | + | 6296 | hexokinase 1 [ | -0.1082 | 8.83934 | 18 | 7.72E-05 | 0.001096 |
| GDF3 | 12 | 7689784 | 7695775 | - | 1236 | growth differe | -0.4673 | 3.11514 | 17.9959 | 7.73E-05 | 0.001096 |
| BRD4 | 19 | 1.5E+07 | 1.5E+07 | - | 12093 | bromodomain | -0.1421 | 6.89943 | 17.9897 | 7.75E-05 | 0.001098 |
| HAGHL | 16 | 726936 | 735525 | + | 5287 | hydroxyacylglu | 0.2945 | 4.65118 | 18.1792 | 7.79E-05 | 0.001103 |
| MCM3 | 6 | 5.2E+07 | 5.2E+07 | - | 3650 | minichromoso | 0.1042 | 8.84454 | 17.956 | 7.86E-05 | 0.001111 |
| USP34 | 2 | 6.1E+07 | 6.1E+07 | - | 18125 | ubiquitin speci | 0.12863 | 7.39301 | 17.8741 | 8.12E-05 | 0.001147 |
| RCBTB2 | 13 | 4.8E+07 | 4.9E+07 | - | 3545 | RCC1 and BTB | 0.3306 | 4.06157 | 17.8588 | 8.17E-05 | 0.001152 |
| AKR1A1 | 1 | 4.6E+07 | 4.6E+07 | + | 3121 | aldo-keto redu | -0.1229 | 7.65569 | 17.8503 | 8.20E-05 | 0.001155 |
| DDX1 | 2 | 1.6E+07 | 1.6E+07 | + | 3833 | DEAD-box heli | -0.127 | 7.61639 | 17.8183 | 8.31E-05 | 0.001169 |
| ISYNA1 | 19 | 1.8E+07 | 1.8E+07 | - | 3284 | inositol-3-phos | 0.12077 | 7.76483 | 17.8122 | 8.33E-05 | 0.001171 |
| LINC0123 | 12 | 1.1E+08 | 1.1E+08 | - | 4484 | long intergenic | 0.66316 | 2.07721 | 17.8047 | 8.35E-05 | 0.001172 |
| TCF12 | 15 | 5.7E+07 | 5.7E+07 | + | 9446 | transcription fa | 0.14944 | 6.6382 | 17.8028 | 8.36E-05 | 0.001172 |
| AP3B1 | 5 | 7.8E+07 | 7.8E+07 | - | 7310 | adaptor relate | 0.16983 | 6.40402 | 17.9416 | 8.36E-05 | 0.001172 |
| CLIP2 | 7 | 7.4E+07 | 7.4E+07 | + | 7315 | CAP-Gly domai | -0.152 | 6.66578 | 17.7772 | 8.45E-05 | 0.001183 |
| ARFGAP1 | 20 | 6.3E+07 | 6.3E+07 | + | 6610 | ADP ribosylatio | 0.15546 | 6.51173 | 17.7672 | 8.48E-05 | 0.001186 |
| PLEKHH1 | 14 | 6.8E+07 | 6.8E+07 | + | 10817 | pleckstrin hom | 0.21425 | 5.43782 | 17.762 | 8.50E-05 | 0.001188 |
| RPL13 | 16 | 9E+07 | 9E+07 | + | 6172 | ribosomal prot | 0.10008 | 9.97816 | 17.8899 | 8.54E-05 | 0.001192 |
| PTTG1 | 5 | 1.6E+08 | 1.6E+08 | + | 1975 | pituitary tumo | -0.137 | 7.1755 | 17.7494 | 8.54E-05 | 0.001192 |
| ATP5PD | 17 | 7.5E+07 | 7.5E+07 | - | 3231 | ATP synthase p | -0.1505 | 6.67768 | 17.7261 | 8.62E-05 | 0.001201 |
| RAB34 | 17 | 2.9E+07 | 2.9E+07 | - | 3303 | RAB34, membe | -0.118 | 7.92249 | 17.7251 | 8.63E-05 | 0.001201 |
| THSD7A | 7 | 1.1E+07 | 1.2E+07 | - | 10921 | thrombospond | 0.97032 | 1.05539 | 17.7155 | 8.66E-05 | 0.001204 |
| CHST9 | 18 | 2.7E+07 | 2.7E+07 | - | 11512 | carbohydrate s | -0.3716 | 3.74628 | 17.7133 | 8.67E-05 | 0.001204 |
| C20orf27 | 20 | 3753508 | 3768387 | - | 2415 | chromosome 2 | -0.1579 | 6.55267 | 17.7069 | 8.69E-05 | 0.001206 |
| PCDHGA1 | 5 | 1.4E+08 | 1.4E+08 | + | 6643 | protocadherin | 0.40372 | 3.43482 | 17.7034 | 8.70E-05 | 0.001207 |
| LFNG | 7 | 2512529 | 2529177 | + | 3486 | LFNG O-fucosy | 0.23728 | 5.15312 | 17.6914 | 8.74E-05 | 0.001212 |
| GPR176 | 15 | 4E+07 | 4E+07 | - | 5119 | G protein-coup | 0.13578 | 8.10271 | 19.1864 | 8.77E-05 | 0.001212 |
| CNTN1 | 12 | 4.1E+07 | 4.1E+07 | + | 7492 | contactin 1 [So | -0.3455 | 4.00771 | 17.6843 | 8.77E-05 | 0.001212 |
| ACAP3 | 1 | 1292390 | 1309609 | - | 8707 | ArfGAP with co | 0.26927 | 4.7341 | 17.6841 | 8.77E-05 | 0.001212 |
| ATXN10 | 22 | 4.6E+07 | 4.6E+07 | + | 5127 | ataxin 10 [Sou | 0.13151 | 7.32184 | 17.678 | 8.79E-05 | 0.001213 |
| PDE4DIP | 1 | 1.5E+08 | 1.5E+08 | + | 27623 | phosphodieste | -0.2458 | 5.09382 | 17.6762 | 8.80E-05 | 0.001213 |
| DIPK1B | 9 | 1.4E+08 | 1.4E+08 | + | 3394 | divergent prot | 0.22954 | 5.24177 | 17.6712 | 8.82E-05 | 0.001215 |
| SHC3 | 9 | 8.9E+07 | 8.9E+07 | - | 9980 | SHC adaptor p | 0.32747 | 4.6536 | 19.3997 | 8.83E-05 | 0.001216 |
| AARS | 16 | 7E+07 | 7E+07 | - | 4498 | alanyl-tRNA sy | -0.1078 | 8.70352 | 17.6427 | 8.92E-05 | 0.001226 |
| SOX9 | 17 | 7.2E+07 | 7.2E+07 | + | 3935 | SRY-box 9 [Sou | -0.5677 | 2.4777 | 17.6265 | 8.98E-05 | 0.001233 |
| TOMM7 | 7 | 2.3E+07 | 2.3E+07 | - | 1751 | translocase of | -0.1495 | 6.95083 | 17.7839 | 9.02E-05 | 0.001237 |
| INKA2 | 1 | 1.1E+08 | 1.1E+08 | - | 6393 | inka box actin | -0.252 | 4.98953 | 17.6104 | 9.04E-05 | 0.001239 |

| | | | | | | | | | | | |
|---|---|---|---|---|---|---|---|---|---|---|---|
| DCTPP1 | 16 | 3E+07 | 3E+07 | - | 1581 | dCTP pyrophos | 0.20517 | 5.56625 | 17.577 | 9.16E-05 | 0.001255 |
| NUP205 | 7 | 1.4E+08 | 1.4E+08 | + | 7621 | nucleoporin 20 | 0.10973 | 8.36666 | 17.537 | 9.31E-05 | 0.001274 |
| SYTL2 | 11 | 8.6E+07 | 8.6E+07 | - | 11258 | synaptotagmin | 0.38973 | 3.72107 | 17.6686 | 9.33E-05 | 0.001275 |
| CSMD2 | 1 | 3.4E+07 | 3.4E+07 | - | 18157 | CUB and Sushi | 0.2305 | 5.15817 | 17.5142 | 9.40E-05 | 0.001284 |
| MTAP | 9 | 2.2E+07 | 2.2E+07 | + | 14620 | methylthioade | -0.1451 | 6.75433 | 17.4839 | 9.51E-05 | 0.001297 |
| DCAF13 | 8 | 1E+08 | 1E+08 | + | 9199 | DDB1 and CUL | -0.1518 | 6.61221 | 17.4837 | 9.52E-05 | 0.001297 |
| ZNF672 | 1 | 2.5E+08 | 2.5E+08 | + | 3741 | zinc finger pro | 0.20088 | 5.74519 | 17.4753 | 9.55E-05 | 0.0013 |
| KCNK12 | 2 | 4.8E+07 | 4.8E+07 | - | 6685 | potassium two | -0.3458 | 3.96463 | 17.4713 | 9.56E-05 | 0.001301 |
| LINC0070 | 10 | 6779549 | 6879450 | + | 9833 | long intergenic | -1.2291 | 0.53983 | 17.4427 | 9.68E-05 | 0.001315 |
| KCNH2 | 7 | 1.5E+08 | 1.5E+08 | - | 5877 | potassium volt | -0.3192 | 4.35043 | 17.5789 | 9.75E-05 | 0.001325 |
| SORBS1 | 10 | 9.5E+07 | 9.6E+07 | - | 10951 | sorbin and SH3 | -0.2127 | 5.66121 | 17.589 | 9.78E-05 | 0.001327 |
| TIMP2 | 17 | 7.9E+07 | 7.9E+07 | - | 5633 | TIMP metallop | -0.1819 | 5.96257 | 17.3952 | 9.86E-05 | 0.001337 |
| TMEM94 | 17 | 7.5E+07 | 7.6E+07 | + | 9045 | transmembran | 0.18703 | 5.86872 | 17.3901 | 9.89E-05 | 0.001339 |
| SMC6 | 2 | 1.8E+07 | 1.8E+07 | - | 9824 | structural mair | -0.1527 | 6.62092 | 17.3811 | 9.92E-05 | 0.001341 |
| KIAA0040 | 1 | 1.8E+08 | 1.8E+08 | - | 4767 | KIAA0040 [Sou | -0.2894 | 4.4669 | 17.375 | 9.95E-05 | 0.001343 |
| MIR302CH | 4 | 1.1E+08 | 1.1E+08 | - | 743 | miR-302/367 c | -0.3584 | 3.86052 | 17.3625 | 1.00E-04 | 0.001349 |
| C8orf33 | 8 | 1.5E+08 | 1.5E+08 | + | 4353 | chromosome 8 | 0.18674 | 5.88569 | 17.3539 | 1.00E-04 | 0.001352 |
| PRICKLE1 | 12 | 4.2E+07 | 4.3E+07 | - | 8223 | prickle planar | 0.32869 | 4.08508 | 17.3458 | 1.01E-04 | 0.001355 |
| FSTL1 | 3 | 1.2E+08 | 1.2E+08 | - | 8467 | follistatin like 1 | 0.10832 | 9.24451 | 17.6626 | 1.03E-04 | 0.001384 |
| ALYREF | 17 | 8.2E+07 | 8.2E+07 | - | 1574 | Aly/REF export | 0.12933 | 7.32976 | 17.2785 | 1.03E-04 | 0.001391 |
| RRP7A | 22 | 4.3E+07 | 4.3E+07 | - | 5700 | ribosomal RNA | 0.2194 | 5.33814 | 17.2698 | 1.04E-04 | 0.001394 |
| CANX | 5 | 1.8E+08 | 1.8E+08 | + | 6951 | calnexin [Sour | -0.1019 | 9.12034 | 17.2648 | 1.04E-04 | 0.001396 |
| IQGAP1 | 15 | 9E+07 | 9.1E+07 | + | 11507 | IQ motif conta | -0.1066 | 8.57063 | 17.26 | 1.04E-04 | 0.001397 |
| CDCA7 | 2 | 1.7E+08 | 1.7E+08 | + | 3934 | cell division cy | 0.14052 | 6.86655 | 17.2503 | 1.05E-04 | 0.001402 |
| ITM2B | 13 | 4.8E+07 | 4.8E+07 | + | 10706 | integral memb | 0.13692 | 7.44681 | 17.6006 | 1.05E-04 | 0.001411 |
| ASPHD2 | 22 | 2.6E+07 | 2.6E+07 | + | 3370 | aspartate beta | -0.3886 | 3.57569 | 17.1762 | 1.08E-04 | 0.001441 |
| TMTC4 | 13 | 1E+08 | 1E+08 | - | 6272 | transmembran | 0.17887 | 5.9619 | 17.124 | 1.10E-04 | 0.00147 |
| LMO7 | 13 | 7.6E+07 | 7.6E+07 | + | 14367 | LIM domain 7 | 0.25606 | 4.9329 | 17.1058 | 1.11E-04 | 0.00148 |
| JAK1 | 1 | 6.5E+07 | 6.5E+07 | - | 6362 | Janus kinase 1 | 0.18516 | 5.89233 | 17.099 | 1.11E-04 | 0.001483 |
| TMX4 | 20 | 7977346 | 8019805 | - | 6557 | thioredoxin rel | -0.3065 | 4.22503 | 17.0864 | 1.12E-04 | 0.001489 |
| CREBBP | 16 | 3725054 | 3880726 | - | 15492 | CREB binding p | -0.1431 | 6.86123 | 17.0842 | 1.12E-04 | 0.001489 |
| DOP1A | 6 | 8.3E+07 | 8.3E+07 | + | 10045 | DOP1 leucine z | 0.27742 | 4.54518 | 17.0798 | 1.12E-04 | 0.00149 |
| ARPIN | 15 | 9E+07 | 9E+07 | - | 8590 | actin related p | 0.34573 | 3.84357 | 17.0483 | 1.14E-04 | 0.001507 |
| TBC1D4 | 13 | 7.5E+07 | 7.5E+07 | - | 7977 | TBC1 domain f | 0.15678 | 6.50843 | 17.0375 | 1.14E-04 | 0.001512 |
| HERC2P3 | 15 | 2E+07 | 2.1E+07 | - | 9413 | hect domain a | 0.33054 | 4.03566 | 17.0339 | 1.14E-04 | 0.001513 |
| GADD45G | 9 | 9E+07 | 9E+07 | + | 1323 | growth arrest a | 0.44842 | 3.15722 | 17.0303 | 1.15E-04 | 0.001514 |
| FAM20A | 17 | 6.9E+07 | 6.9E+07 | - | 6181 | FAM20A, golgi | -1.1678 | 2.38267 | 28.802 | 1.15E-04 | 0.001516 |
| HSD17B12 | 11 | 4.4E+07 | 4.4E+07 | + | 7716 | hydroxysteroid | -0.1504 | 6.52948 | 17.0228 | 1.15E-04 | 0.001516 |
| ATP2B4 | 1 | 2E+08 | 2E+08 | + | 9721 | ATPase plasma | -0.1803 | 5.93307 | 17.015 | 1.15E-04 | 0.001519 |
| SMC3 | 10 | 1.1E+08 | 1.1E+08 | + | 4275 | structural mair | -0.1339 | 7.37958 | 17.13 | 1.16E-04 | 0.001526 |
| CHAF1A | 19 | 4402640 | 4445018 | + | 4326 | chromatin asse | 0.1233 | 7.67942 | 16.9969 | 1.16E-04 | 0.001528 |
| USP31 | 16 | 2.3E+07 | 2.3E+07 | - | 12017 | ubiquitin speci | -0.2827 | 4.59346 | 16.9675 | 1.18E-04 | 0.001545 |
| MT1X | 16 | 5.7E+07 | 5.7E+07 | + | 1727 | metallothionei | -0.261 | 4.74836 | 16.9579 | 1.18E-04 | 0.00155 |

| | | | | | | | | | | | | |
|---|---|---|---|---|---|---|---|---|---|---|---|---|
| SLC52A3 | 20 | 760080 | 776015 | - | | 3776 | solute carrier f | -0.3354 | 4.06486 | 16.9188 | 1.20E-04 | 0.001572 |
| SLCO4C1 | 5 | 1E+08 | 1E+08 | - | | 5069 | solute carrier o | -0.408 | 3.44774 | 16.9115 | 1.20E-04 | 0.001575 |
| NEXN | 1 | 7.8E+07 | 7.8E+07 | + | | 4436 | nexilin F-actin | 0.47313 | 2.95504 | 16.9027 | 1.21E-04 | 0.001579 |
| PRKCQ-AS | 10 | 6580419 | 6616452 | + | | 3892 | PRKCQ antisen | -0.3179 | 4.07575 | 16.879 | 1.22E-04 | 0.001593 |
| C20orf96 | 20 | 270863 | 290778 | - | | 1730 | chromosome 2 | -0.3077 | 4.29463 | 16.866 | 1.23E-04 | 0.0016 |
| ATP1A2 | 1 | 1.6E+08 | 1.6E+08 | + | | 6298 | ATPase Na+/K- | 0.1611 | 6.42529 | 16.8585 | 1.23E-04 | 0.001604 |
| POLR2L | 11 | 837356 | 842529 | - | | 1126 | RNA polymera | -0.1689 | 6.24399 | 16.834 | 1.24E-04 | 0.001619 |
| RAB13 | 1 | 1.5E+08 | 1.5E+08 | - | | 2751 | RAB13, memb | 0.23876 | 5.03396 | 16.8175 | 1.25E-04 | 0.001628 |
| INHBE | 12 | 5.7E+07 | 5.7E+07 | + | | 3046 | inhibin subunit | -0.3614 | 3.74109 | 16.81 | 1.25E-04 | 0.001632 |
| PLPP1 | 5 | 5.5E+07 | 5.6E+07 | - | | 1935 | phospholipid p | -0.2731 | 4.90488 | 17.3094 | 1.26E-04 | 0.001637 |
| NRBP2 | 8 | 1.4E+08 | 1.4E+08 | - | | 4934 | nuclear recept | 0.39592 | 3.41501 | 16.7961 | 1.26E-04 | 0.001638 |
| PRKDC | 8 | 4.8E+07 | 4.8E+07 | - | | 15417 | protein kinase, | 0.09011 | 10.0544 | 16.7929 | 1.26E-04 | 0.001639 |
| PDCL3 | 2 | 1E+08 | 1E+08 | + | | 1532 | phosducin like | -0.2874 | 4.44664 | 16.7874 | 1.27E-04 | 0.001641 |
| OAZ1 | 19 | 2269509 | 2273490 | + | | 3436 | ornithine deca | 0.10196 | 8.80922 | 16.7848 | 1.27E-04 | 0.001641 |
| RAB3C | 5 | 5.9E+07 | 5.9E+07 | + | | 9621 | RAB3C, memb | 0.41353 | 3.43331 | 16.8105 | 1.27E-04 | 0.001649 |
| ADGRV1 | 5 | 9.1E+07 | 9.1E+07 | + | | 33822 | adhesion G pro | 0.16964 | 6.06883 | 16.76 | 1.28E-04 | 0.001655 |
| FADS1 | 11 | 6.2E+07 | 6.2E+07 | - | | 10230 | fatty acid desa | 0.11729 | 7.75194 | 16.718 | 1.30E-04 | 0.001683 |
| KIF5A | 12 | 5.8E+07 | 5.8E+07 | + | | 6216 | kinesin family | -0.2124 | 5.5892 | 16.8325 | 1.31E-04 | 0.001688 |
| TACC2 | 10 | 1.2E+08 | 1.2E+08 | + | | 18855 | transforming a | 0.19636 | 5.64125 | 16.7021 | 1.31E-04 | 0.001691 |
| MDC1 | R6_MHC_ | 3.1E+07 | 3.1E+07 | - | | 7622 | mediator of DN | 0.13514 | 6.95772 | 16.6546 | 1.34E-04 | 0.001723 |
| KIRREL2 | 19 | 3.6E+07 | 3.6E+07 | + | | 3382 | kirre like neph | -0.4438 | 3.23495 | 16.6898 | 1.34E-04 | 0.001723 |
| AC003975 | 7 | 1.3E+08 | 1.3E+08 | + | | 837 | novel transcrip | -0.4631 | 2.96175 | 16.6457 | 1.34E-04 | 0.001726 |
| DGKH | 13 | 4.2E+07 | 4.2E+07 | + | | 19637 | diacylglycerol | 0.33202 | 3.93473 | 16.6374 | 1.35E-04 | 0.00173 |
| AC131212 | 12 | 1.3E+08 | 1.3E+08 | + | | 4219 | novel transcrip | 0.61518 | 2.12291 | 16.6248 | 1.35E-04 | 0.001737 |
| BTBD6 | 14 | 1.1E+08 | 1.1E+08 | + | | 2392 | BTB domain co | 0.32646 | 3.95063 | 16.6241 | 1.35E-04 | 0.001737 |
| CPT1A | 11 | 6.9E+07 | 6.9E+07 | - | | 6571 | carnitine palm | -0.2512 | 4.8576 | 16.6052 | 1.36E-04 | 0.001749 |
| KAT6B | 10 | 7.5E+07 | 7.5E+07 | + | | 32495 | lysine acetyltra | -0.1821 | 5.8692 | 16.6019 | 1.37E-04 | 0.00175 |
| PLCB2 | 15 | 4E+07 | 4E+07 | - | | 7349 | phospholipase | 0.28015 | 4.52999 | 16.5958 | 1.37E-04 | 0.001752 |
| PHKA1 | X | 7.3E+07 | 7.3E+07 | - | | 6350 | phosphorylase | 0.30196 | 4.36913 | 16.6223 | 1.37E-04 | 0.001753 |
| PVT1 | 8 | 1.3E+08 | 1.3E+08 | + | | 8959 | Pvt1 oncogene | 0.3509 | 3.76844 | 16.5891 | 1.37E-04 | 0.001754 |
| IGF2BP1 | 17 | 4.9E+07 | 4.9E+07 | + | | 9631 | insulin like gro | -0.0944 | 9.46995 | 16.5772 | 1.38E-04 | 0.001761 |
| BBX | 3 | 1.1E+08 | 1.1E+08 | + | | 11892 | BBX, HMG-box | 0.15332 | 6.55437 | 16.5658 | 1.39E-04 | 0.001768 |
| AASS | 7 | 1.2E+08 | 1.2E+08 | - | | 6664 | aminoadipate- | -0.0946 | 10.0019 | 16.5562 | 1.39E-04 | 0.001773 |
| DAGLA | 11 | 6.2E+07 | 6.2E+07 | + | | 5799 | diacylglycerol | 0.28868 | 4.37758 | 16.5529 | 1.39E-04 | 0.001774 |
| MX2 | 21 | 4.1E+07 | 4.1E+07 | + | | 12994 | MX dynamin li | -0.3088 | 4.14396 | 16.5514 | 1.39E-04 | 0.001774 |
| GRB14 | 2 | 1.6E+08 | 1.6E+08 | - | | 2958 | growth factor | 0.2978 | 4.28992 | 16.542 | 1.40E-04 | 0.001779 |
| PIP5K1C | 19 | 3630183 | 3700479 | - | | 6963 | phosphatidylin | 0.19054 | 5.78091 | 16.5301 | 1.41E-04 | 0.001786 |
| PIK3R3 | 1 | 4.6E+07 | 4.6E+07 | - | | 7397 | phosphoinositi | 0.31356 | 4.12506 | 16.5184 | 1.41E-04 | 0.001793 |
| ABCA2 | 9 | 1.4E+08 | 1.4E+08 | - | | 11507 | ATP binding ca | 0.26595 | 4.8106 | 16.6369 | 1.42E-04 | 0.001794 |
| SPINT1 | 15 | 4.1E+07 | 4.1E+07 | + | | 3542 | serine peptida | -0.1222 | 7.56686 | 16.464 | 1.45E-04 | 0.00183 |
| FERMT1 | 20 | 6074845 | 6123544 | - | | 5359 | fermitin family | -0.2782 | 4.60852 | 16.4634 | 1.45E-04 | 0.00183 |
| MT-TC | MT | 5761 | 5826 | - | | 66 | mitochondriall | -0.5475 | 2.65339 | 16.5966 | 1.45E-04 | 0.001833 |
| SSR4 | X | 1.5E+08 | 1.5E+08 | + | | 4590 | signal sequenc | 0.18278 | 5.83656 | 16.4541 | 1.45E-04 | 0.001834 |

| | | | | | | | | | | | |
|---|---|---|---|---|---|---|---|---|---|---|---|
| IFNAR1 | 21 | 3.3E+07 | 3.3E+07 | + | 6879 | interferon alph | -0.2913 | 4.46381 | 16.4461 | 1.46E-04 | 0.001838 |
| GPC2 | 7 | 1E+08 | 1E+08 | - | 3837 | glypican 2 [Sou | 0.18911 | 5.8017 | 16.4185 | 1.47E-04 | 0.001858 |
| ANKRD27 | 19 | 3.3E+07 | 3.3E+07 | - | 7849 | ankyrin repeat | -0.1545 | 6.36949 | 16.4058 | 1.48E-04 | 0.001864 |
| AURKA | 20 | 5.6E+07 | 5.6E+07 | - | 2928 | aurora kinase A | -0.1344 | 6.92776 | 16.4056 | 1.48E-04 | 0.001864 |
| MOXD1 | 6 | 1.3E+08 | 1.3E+08 | - | 5223 | monooxygenas | 0.59135 | 2.20722 | 16.3801 | 1.50E-04 | 0.001881 |
| DPPA2 | 3 | 1.1E+08 | 1.1E+08 | - | 1383 | developmenta | -0.4226 | 3.23758 | 16.3754 | 1.50E-04 | 0.001883 |
| FBN2 | 5 | 1.3E+08 | 1.3E+08 | - | 13218 | fibrillin 2 [Sour | 0.68399 | 6.03236 | 61.0283 | 1.50E-04 | 0.001884 |
| PTPRD | 9 | 8314246 | 1.1E+07 | - | 11198 | protein tyrosin | 0.16655 | 6.13931 | 16.3643 | 1.51E-04 | 0.001887 |
| SNHG8 | 4 | 1.2E+08 | 1.2E+08 | + | 1115 | small nucleolar | 0.20266 | 5.51233 | 16.3389 | 1.52E-04 | 0.001905 |
| SLFN5 | 17 | 3.5E+07 | 3.5E+07 | + | 10674 | schlafen family | -0.3164 | 4.05499 | 16.3332 | 1.53E-04 | 0.001908 |
| SMOC2 | 6 | 1.7E+08 | 1.7E+08 | + | 4032 | SPARC related | -0.5633 | 2.53937 | 16.4351 | 1.54E-04 | 0.001917 |
| FAM120B | HSCHR6_1 | 1.7E+08 | 1.7E+08 | + | 6029 | family with seq | 0.26222 | 4.62524 | 16.3156 | 1.54E-04 | 0.001917 |
| ZFP64 | 20 | 5.2E+07 | 5.2E+07 | - | 6790 | ZFP64 zinc fing | -0.2442 | 4.96766 | 16.3152 | 1.54E-04 | 0.001917 |
| DSE | 6 | 1.2E+08 | 1.2E+08 | + | 19193 | dermatan sulfa | 0.18753 | 5.93233 | 16.4289 | 1.54E-04 | 0.00192 |
| RSL24D1 | 15 | 5.5E+07 | 5.5E+07 | - | 2965 | ribosomal L24 | -0.1243 | 7.30978 | 16.3056 | 1.54E-04 | 0.001921 |
| ZMPSTE24 | 1 | 4E+07 | 4E+07 | + | 3439 | zinc metallope | -0.205 | 5.85533 | 17.0613 | 1.55E-04 | 0.001927 |
| UBE2E2 | 3 | 2.3E+07 | 2.4E+07 | + | 3241 | ubiquitin conju | 0.24237 | 4.99086 | 16.2687 | 1.57E-04 | 0.001948 |
| ZFAND5 | 9 | 7.2E+07 | 7.2E+07 | - | 7065 | zinc finger AN1 | -0.1179 | 7.92324 | 16.2295 | 1.59E-04 | 0.001978 |
| RPL7A | HG2030_P | 1.3E+08 | 1.3E+08 | + | 2238 | ribosomal prot | 0.08956 | 10.1136 | 16.2248 | 1.60E-04 | 0.00198 |
| UQCR10 | 22 | 3E+07 | 3E+07 | + | 975 | ubiquinol-cyto | -0.1799 | 5.91172 | 16.2164 | 1.60E-04 | 0.001984 |
| CARMIL1 | 6 | 2.5E+07 | 2.6E+07 | + | 6853 | capping protei | 0.1603 | 6.22408 | 16.1967 | 1.62E-04 | 0.001998 |
| FZR1 | 19 | 3506273 | 3538330 | + | 6511 | fizzy and cell d | 0.2068 | 5.54077 | 16.1901 | 1.62E-04 | 0.002002 |
| MAP3K1 | 5 | 5.7E+07 | 5.7E+07 | + | 7741 | mitogen-activa | 0.25413 | 4.83472 | 16.1736 | 1.63E-04 | 0.002014 |
| NLRP12 | 19 | 5.4E+07 | 5.4E+07 | - | 5088 | NLR family pyr | -0.427 | 3.15512 | 16.1397 | 1.65E-04 | 0.002041 |
| CLIC6 | 21 | 3.5E+07 | 3.5E+07 | + | 3860 | chloride intrac | -0.7136 | 1.85256 | 16.1608 | 1.66E-04 | 0.002046 |
| CDC42BPA | 1 | 2.3E+08 | 2.3E+08 | - | 14055 | CDC42 binding | 0.15187 | 6.50212 | 16.1296 | 1.66E-04 | 0.002046 |
| IGFBPL1 | 9 | 3.8E+07 | 3.8E+07 | - | 3566 | insulin like gro | -0.1974 | 5.60271 | 16.1186 | 1.67E-04 | 0.002053 |
| PCSK7 | 11 | 1.2E+08 | 1.2E+08 | - | 22303 | proprotein con | 0.23493 | 4.99017 | 16.1147 | 1.67E-04 | 0.002053 |
| RAB31 | 18 | 9708275 | 9862551 | + | 5476 | RAB31, membe | -0.1817 | 5.84946 | 16.1047 | 1.68E-04 | 0.00206 |
| IFITM1 | 11 | 313506 | 315272 | + | 936 | interferon indu | -0.1331 | 6.98997 | 16.0788 | 1.70E-04 | 0.002081 |
| PAK3 | X | 1.1E+08 | 1.1E+08 | + | 10446 | p21 (RAC1) act | 0.46617 | 2.90742 | 16.0651 | 1.71E-04 | 0.002091 |
| SET | 9 | 1.3E+08 | 1.3E+08 | + | 4355 | SET nuclear pr | -0.087 | 10.2708 | 16.0458 | 1.72E-04 | 0.002106 |
| SAT2 | 17 | 7626234 | 7627876 | - | 1643 | spermidine/sp | -0.2501 | 4.81277 | 16.0366 | 1.73E-04 | 0.002111 |
| ELMOD2 | 4 | 1.4E+08 | 1.4E+08 | + | 6617 | ELMO domain | -0.22 | 5.21876 | 16.0347 | 1.73E-04 | 0.002111 |
| IGLON5 | 19 | 5.1E+07 | 5.1E+07 | + | 2606 | IgLON family m | -0.1837 | 5.83684 | 16.0319 | 1.73E-04 | 0.002111 |
| SNRPB2 | 20 | 1.7E+07 | 1.7E+07 | + | 2608 | small nuclear r | -0.1597 | 6.2195 | 16.0307 | 1.73E-04 | 0.002111 |
| LPCAT2 | 16 | 5.6E+07 | 5.6E+07 | + | 7361 | lysophosphatid | -0.4744 | 2.8541 | 16.0302 | 1.73E-04 | 0.002111 |
| PSMD11 | 17 | 3.2E+07 | 3.2E+07 | + | 7083 | proteasome 26 | -0.1253 | 7.25886 | 16.0149 | 1.74E-04 | 0.002122 |
| MRPS21 | 1 | 1.5E+08 | 1.5E+08 | + | 1641 | mitochondrial | -0.1421 | 6.83976 | 16.0069 | 1.75E-04 | 0.002127 |
| DEF8 | 16 | 9E+07 | 9E+07 | + | 6565 | differentially e | 0.15796 | 6.22973 | 16.0059 | 1.75E-04 | 0.002127 |
| TFE3 | X | 4.9E+07 | 4.9E+07 | - | 4207 | transcription fa | -0.1516 | 6.54831 | 16.0016 | 1.75E-04 | 0.002129 |
| ZNF217 | 20 | 5.4E+07 | 5.4E+07 | - | 6181 | zinc finger prot | -0.1387 | 6.87009 | 15.9967 | 1.76E-04 | 0.002131 |
| CCDC124 | 19 | 1.8E+07 | 1.8E+07 | + | 1661 | coiled-coil dom | 0.15706 | 6.24383 | 15.9673 | 1.78E-04 | 0.002156 |

| | | | | | | | | | | | |
|---|---|---|---|---|---|---|---|---|---|---|---|
| RNF145 | 5 | 1.6E+08 | 1.6E+08 | - | 4614 | ring finger prot | 0.14305 | 6.67466 | 15.9611 | 1.78E-04 | 0.002157 |
| PLXNB1 | 3 | 4.8E+07 | 4.8E+07 | - | 9768 | plexin B1 [Sou | 0.13627 | 6.89672 | 15.9595 | 1.78E-04 | 0.002157 |
| SYK | 9 | 9.1E+07 | 9.1E+07 | + | 5210 | spleen associat | 0.35155 | 3.69749 | 15.9595 | 1.78E-04 | 0.002157 |
| SCNN1G | 16 | 2.3E+07 | 2.3E+07 | + | 3477 | sodium channe | -1.1893 | 0.58399 | 15.9755 | 1.79E-04 | 0.00216 |
| HSPA4L | 4 | 1.3E+08 | 1.3E+08 | + | 11439 | heat shock pro | 0.37841 | 3.51877 | 15.9357 | 1.80E-04 | 0.002174 |
| SPART | 13 | 3.6E+07 | 3.6E+07 | - | 6534 | spartin [Source | -0.1578 | 6.26685 | 15.9348 | 1.80E-04 | 0.002174 |
| FAM210B | 20 | 5.6E+07 | 5.6E+07 | + | 3518 | family with sec | -0.2434 | 5.09508 | 16.1129 | 1.81E-04 | 0.002175 |
| EPHA2 | 1 | 1.6E+07 | 1.6E+07 | - | 4141 | EPH receptor A | 0.19665 | 6.03065 | 17.0203 | 1.81E-04 | 0.002175 |
| MYO5B | 18 | 5E+07 | 5E+07 | - | 10566 | myosin VB [Sou | -0.2841 | 4.36771 | 15.9139 | 1.82E-04 | 0.002188 |
| SDC2 | 8 | 9.6E+07 | 9.7E+07 | + | 4084 | syndecan 2 [So | 0.1314 | 7.2857 | 16.0186 | 1.82E-04 | 0.002193 |
| DNAJB5 | 9 | 3.5E+07 | 3.5E+07 | + | 5891 | DnaJ heat shoc | 0.15253 | 6.38113 | 15.9001 | 1.83E-04 | 0.002197 |
| CEP70 | 3 | 1.4E+08 | 1.4E+08 | - | 4409 | centrosomal p | -0.2734 | 4.51817 | 15.8919 | 1.83E-04 | 0.002202 |
| CCDC78 | 16 | 722582 | 726954 | - | 4147 | coiled-coil dom | 0.72184 | 1.64884 | 15.8849 | 1.84E-04 | 0.002207 |
| LRIG3 | 12 | 5.9E+07 | 5.9E+07 | - | 5893 | leucine rich rep | 0.3662 | 3.5615 | 15.8792 | 1.84E-04 | 0.00221 |
| GALNT3 | 2 | 1.7E+08 | 1.7E+08 | - | 4686 | polypeptide N- | -0.2479 | 4.85499 | 15.8723 | 1.85E-04 | 0.002215 |
| EXT1 | 8 | 1.2E+08 | 1.2E+08 | - | 8325 | exostosin glyco | 0.18033 | 5.87325 | 15.8675 | 1.85E-04 | 0.002217 |
| FAM184A | 6 | 1.2E+08 | 1.2E+08 | - | 7791 | family with sec | 0.35342 | 3.66507 | 15.8643 | 1.86E-04 | 0.002219 |
| CTCFL | 20 | 5.7E+07 | 5.8E+07 | - | 8928 | CCCTC-binding | -0.7343 | 1.64758 | 15.8622 | 1.86E-04 | 0.002219 |
| CREB3L1 | 11 | 4.6E+07 | 4.6E+07 | + | 4037 | cAMP responsi | -0.2796 | 4.48253 | 15.8519 | 1.87E-04 | 0.002226 |
| RHOBTB1 | 10 | 6.1E+07 | 6.1E+07 | - | 4779 | Rho related BT | 0.30703 | 4.3209 | 16.0297 | 1.87E-04 | 0.002228 |
| LRRK1 | 15 | 1E+08 | 1E+08 | + | 17491 | leucine rich rep | -0.3606 | 3.81243 | 15.958 | 1.87E-04 | 0.002228 |
| SENP2 | 3 | 1.9E+08 | 1.9E+08 | + | 7102 | SUMO specific | -0.1687 | 6.08715 | 15.8416 | 1.87E-04 | 0.00223 |
| SREBF1 | 17 | 1.8E+07 | 1.8E+07 | - | 8030 | sterol regulato | -0.1968 | 6.50597 | 18.7534 | 1.88E-04 | 0.002234 |
| FAM98A | 2 | 3.4E+07 | 3.4E+07 | - | 4919 | family with sec | -0.1384 | 6.83181 | 15.8155 | 1.89E-04 | 0.002249 |
| BOD1L1 | 4 | 1.4E+07 | 1.4E+07 | - | 15862 | biorientation o | -0.1296 | 7.12693 | 15.8151 | 1.89E-04 | 0.002249 |
| PWWP2A | 5 | 1.6E+08 | 1.6E+08 | - | 6537 | PWWP domain | -0.1564 | 6.3173 | 15.814 | 1.90E-04 | 0.002249 |
| MALT1 | 18 | 5.9E+07 | 5.9E+07 | + | 11527 | MALT1 paracas | -0.2531 | 4.91334 | 15.9186 | 1.90E-04 | 0.002254 |
| ECEL1P2 | 2 | 2.3E+08 | 2.3E+08 | - | 1708 | endothelin con | -0.7629 | 1.60109 | 15.7839 | 1.92E-04 | 0.002274 |
| ALDH1B1 | 9 | 3.8E+07 | 3.8E+07 | + | 3117 | aldehyde dehy | -0.1477 | 6.45675 | 15.7818 | 1.92E-04 | 0.002274 |
| TMEM158 | 3 | 4.5E+07 | 4.5E+07 | - | 1813 | transmembran | -0.3417 | 4.87468 | 19.521 | 1.93E-04 | 0.002285 |
| HSPA1B | R6_MHC_ | 3.2E+07 | 3.2E+07 | + | 2517 | heat shock pro | -0.2582 | 4.72789 | 15.7661 | 1.93E-04 | 0.002285 |
| MDM2 | 12 | 6.9E+07 | 6.9E+07 | + | 13283 | MDM2 proto-o | -0.1739 | 5.98346 | 15.7535 | 1.94E-04 | 0.002295 |
| BNIP3 | 10 | 1.3E+08 | 1.3E+08 | - | 5325 | BCL2 interactin | 0.45134 | 4.63837 | 23.4306 | 1.97E-04 | 0.002319 |
| ALDH16A | 19 | 4.9E+07 | 4.9E+07 | + | 4589 | aldehyde dehy | -0.1682 | 6.25781 | 15.8732 | 1.99E-04 | 0.002346 |
| VPS26A | 10 | 6.9E+07 | 6.9E+07 | + | 5116 | VPS26, retrom | -0.2114 | 5.35046 | 15.6842 | 2.00E-04 | 0.002353 |
| RBM7 | 11 | 1.1E+08 | 1.1E+08 | + | 7885 | RNA binding m | -0.2099 | 5.29548 | 15.6826 | 2.00E-04 | 0.002353 |
| COPS6 | 7 | 1E+08 | 1E+08 | + | 2779 | COP9 signaloso | 0.16331 | 6.18706 | 15.6819 | 2.00E-04 | 0.002353 |
| POGZ | 1 | 1.5E+08 | 1.5E+08 | - | 10301 | pogo transposa | -0.1345 | 6.84789 | 15.6479 | 2.03E-04 | 0.002385 |
| TP53INP2 | 20 | 3.5E+07 | 3.5E+07 | + | 4179 | tumor protein | -0.372 | 3.69747 | 15.7545 | 2.03E-04 | 0.002386 |
| SPPL2A | 15 | 5.1E+07 | 5.1E+07 | - | 8045 | signal peptide | -0.2431 | 4.84036 | 15.6198 | 2.06E-04 | 0.002408 |
| PLAGL2 | 20 | 3.2E+07 | 3.2E+07 | - | 5656 | PLAG1 like zinc | -0.1765 | 5.85435 | 15.6197 | 2.06E-04 | 0.002408 |
| WDR19 | 4 | 3.9E+07 | 3.9E+07 | + | 8773 | WD repeat dor | -0.2652 | 4.58484 | 15.6168 | 2.06E-04 | 0.002409 |
| MLEC | 12 | 1.2E+08 | 1.2E+08 | + | 6438 | malectin [Sour | -0.0987 | 8.7405 | 15.6017 | 2.07E-04 | 0.002422 |

| | | | | | | | | | | | |
|---|---|---|---|---|---|---|---|---|---|---|---|
| DISP1 | 1 | 2.2E+08 | 2.2E+08 | + | 5382 | dispatched RN | 0.41086 | 3.27555 | 15.6002 | 2.07E-04 | 0.002422 |
| DEPTOR | 8 | 1.2E+08 | 1.2E+08 | + | 2627 | DEP domain cc | -0.6636 | 1.8846 | 15.5929 | 2.08E-04 | 0.002427 |
| VIRMA | 8 | 9.4E+07 | 9.5E+07 | - | 10775 | vir like m6A m | 0.14721 | 6.50761 | 15.5844 | 2.09E-04 | 0.002434 |
| GDAP1L1 | 20 | 4.4E+07 | 4.4E+07 | + | 3385 | ganglioside ind | -0.4467 | 3.01872 | 15.5782 | 2.09E-04 | 0.002438 |
| PITPNM3 | 17 | 6451263 | 6556555 | - | 10671 | PITPNM family | -0.4565 | 3.12898 | 15.7778 | 2.10E-04 | 0.002447 |
| PRDX1 | 1 | 4.6E+07 | 4.6E+07 | - | 1504 | peroxiredoxin | -0.0942 | 9.18606 | 15.5597 | 2.11E-04 | 0.002453 |
| PDP2 | 16 | 6.7E+07 | 6.7E+07 | + | 8364 | pyruvate dehy | -0.1806 | 5.83241 | 15.5518 | 2.12E-04 | 0.002459 |
| TASOR | 3 | 5.7E+07 | 5.7E+07 | - | 9824 | transcription a | 0.17793 | 5.84474 | 15.5453 | 2.12E-04 | 0.002464 |
| CHMP2A | 19 | 5.9E+07 | 5.9E+07 | - | 2129 | charged multiv | 0.21485 | 5.31681 | 15.529 | 2.14E-04 | 0.002479 |
| BMPR1A | 10 | 8.7E+07 | 8.7E+07 | + | 11412 | bone morphog | -0.1534 | 6.29643 | 15.5061 | 2.16E-04 | 0.002501 |
| RBM14 | 11 | 6.7E+07 | 6.7E+07 | + | 3937 | RNA binding m | -0.1541 | 6.5155 | 15.6003 | 2.17E-04 | 0.002513 |
| NUAK2 | 1 | 2.1E+08 | 2.1E+08 | - | 3441 | NUAK family ki | -0.3282 | 4.06199 | 15.5804 | 2.19E-04 | 0.00253 |
| FHDC1 | 4 | 1.5E+08 | 1.5E+08 | + | 6593 | FH2 domain cc | 0.17503 | 5.88173 | 15.4726 | 2.19E-04 | 0.00253 |
| KIF20B | 10 | 9E+07 | 9E+07 | + | 6851 | kinesin family | -0.1418 | 6.69084 | 15.4664 | 2.19E-04 | 0.002535 |
| AP001453 | 11 | 6.4E+07 | 6.4E+07 | - | 853 | novel transcrip | 0.59808 | 2.10485 | 15.4554 | 2.20E-04 | 0.002545 |
| VASP | 19 | 4.6E+07 | 4.6E+07 | + | 3538 | vasodilator stir | -0.112 | 7.77271 | 15.4529 | 2.21E-04 | 0.002545 |
| MAFG | 17 | 8.2E+07 | 8.2E+07 | - | 5241 | MAF bZIP trans | 0.16926 | 6.2022 | 15.6172 | 2.21E-04 | 0.002553 |
| NOP10 | 15 | 3.4E+07 | 3.4E+07 | - | 554 | NOP10 ribonu | -0.1432 | 6.57429 | 15.4078 | 2.25E-04 | 0.00259 |
| CPNE2 | 16 | 5.7E+07 | 5.7E+07 | + | 7737 | copine 2 [Sour | 0.21827 | 5.22751 | 15.3619 | 2.29E-04 | 0.002636 |
| STAT1 | 2 | 1.9E+08 | 1.9E+08 | - | 6053 | signal transduc | 0.13772 | 6.72753 | 15.3609 | 2.29E-04 | 0.002636 |
| ALCAM | 3 | 1.1E+08 | 1.1E+08 | + | 7179 | activated leuk | 0.17395 | 5.89221 | 15.36 | 2.29E-04 | 0.002636 |
| TTC9 | 14 | 7.1E+07 | 7.1E+07 | + | 5094 | tetratricopepti | 0.18803 | 5.79186 | 15.3915 | 2.32E-04 | 0.002663 |
| HMGA2 | 12 | 6.6E+07 | 6.6E+07 | + | 15403 | high mobility g | 0.11283 | 8.1509 | 15.4198 | 2.34E-04 | 0.002683 |
| FUBP1 | 1 | 7.8E+07 | 7.8E+07 | - | 5666 | far upstream e | 0.10843 | 8.15853 | 15.3078 | 2.34E-04 | 0.002688 |
| METTL7A | 12 | 5.1E+07 | 5.1E+07 | + | 4076 | methyltransfer | 0.24304 | 4.94978 | 15.3351 | 2.35E-04 | 0.002692 |
| SCN1A | 2 | 1.7E+08 | 1.7E+08 | - | 16946 | sodium voltage | 1.00999 | 0.83176 | 15.3001 | 2.35E-04 | 0.002693 |
| GPR173 | X | 5.3E+07 | 5.3E+07 | + | 4608 | G protein-coup | -0.3496 | 3.64173 | 15.2479 | 2.40E-04 | 0.002748 |
| GPRASP1 | X | 1E+08 | 1E+08 | + | 6789 | G protein-coup | 0.58071 | 2.13185 | 15.2338 | 2.42E-04 | 0.002763 |
| NINL | 20 | 2.5E+07 | 2.6E+07 | - | 6030 | ninein like [Sou | -0.2043 | 5.8415 | 16.2273 | 2.43E-04 | 0.002769 |
| ADAMTS8 | 11 | 1.3E+08 | 1.3E+08 | - | 4141 | ADAM metallo | 0.30756 | 4.20766 | 15.3233 | 2.43E-04 | 0.00277 |
| SH2D3A | 19 | 6752160 | 6767463 | - | 5591 | SH2 domain cc | -0.2487 | 4.73893 | 15.2196 | 2.43E-04 | 0.002773 |
| HSPD1 | 2 | 2E+08 | 2E+08 | - | 4123 | heat shock pro | -0.0843 | 10.3687 | 15.2156 | 2.44E-04 | 0.002774 |
| PAK1IP1 | 6 | 1.1E+07 | 1.1E+07 | + | 1523 | PAK1 interacti | -0.1939 | 5.71702 | 15.318 | 2.44E-04 | 0.002774 |
| TNFRSF10 | 8 | 2.3E+07 | 2.3E+07 | + | 1868 | TNF receptor s | 0.54454 | 2.38246 | 15.2126 | 2.44E-04 | 0.002774 |
| INTS13 | 12 | 2.7E+07 | 2.7E+07 | - | 3059 | integrator com | -0.1728 | 5.94711 | 15.2041 | 2.45E-04 | 0.002782 |
| SERTM2 | X | 1.1E+08 | 1.1E+08 | + | 4758 | serine rich and | -0.8291 | 1.28544 | 15.2026 | 2.45E-04 | 0.002782 |
| AOX1 | 2 | 2E+08 | 2E+08 | + | 5410 | aldehyde oxida | 0.54221 | 2.39763 | 15.1971 | 2.46E-04 | 0.002786 |
| TMCO3 | 13 | 1.1E+08 | 1.1E+08 | + | 10381 | transmembran | 0.17798 | 5.77205 | 15.1901 | 2.46E-04 | 0.002792 |
| RND1 | 12 | 4.9E+07 | 4.9E+07 | - | 3948 | Rho family GTI | -0.3588 | 3.6154 | 15.1839 | 2.47E-04 | 0.002797 |
| CELF5 | 19 | 3224661 | 3297076 | + | 6066 | CUGBP Elav-lik | 0.40503 | 3.68658 | 16.2747 | 2.50E-04 | 0.002829 |
| TNXB | R6_MHC_ | 3.2E+07 | 3.2E+07 | - | 15909 | tenascin XB [Sc | -0.4296 | 3.10193 | 15.1203 | 2.54E-04 | 0.002869 |
| BTG2 | 1 | 2E+08 | 2E+08 | + | 2757 | BTG anti-prolif | 0.31448 | 4.03525 | 15.1079 | 2.55E-04 | 0.002882 |
| LIN7A | 12 | 8.1E+07 | 8.1E+07 | - | 6795 | lin-7 homolog | 0.29662 | 4.22857 | 15.0776 | 2.58E-04 | 0.002916 |

| | | | | | | | | | | | |
|---|---|---|---|---|---|---|---|---|---|---|---|
| HERC2 | SCHR15_1 | 2.8E+07 | 2.8E+07 | - | 20183 | HECT and RLD | 0.11371 | 7.58264 | 15.0714 | 2.59E-04 | 0.002922 |
| CNBP | 3 | 1.3E+08 | 1.3E+08 | - | 1927 | CCHC-type zinc | 0.09831 | 8.67101 | 15.0661 | 2.60E-04 | 0.002926 |
| TRIO | 5 | 1.4E+07 | 1.5E+07 | + | 17860 | trio Rho guanin | 0.14517 | 6.7042 | 15.1642 | 2.60E-04 | 0.002929 |
| LLGL1 | 17 | 1.8E+07 | 1.8E+07 | + | 6590 | LLGL scribble c | -0.1187 | 7.39519 | 15.0476 | 2.62E-04 | 0.002944 |
| ECEL1P1 | 2 | 2.3E+08 | 2.3E+08 | - | 1742 | endothelin con | -0.7802 | 1.70054 | 15.5572 | 2.62E-04 | 0.002944 |
| SYT11 | 1 | 1.6E+08 | 1.6E+08 | + | 5182 | synaptotagmin | -0.3106 | 4.06662 | 15.0335 | 2.63E-04 | 0.002957 |
| NSA2 | 5 | 7.5E+07 | 7.5E+07 | + | 4962 | NSA2, ribosom | -0.1538 | 6.24672 | 15.031 | 2.64E-04 | 0.002958 |
| FAM102B | 1 | 1.1E+08 | 1.1E+08 | + | 9350 | family with seq | -0.2329 | 4.9154 | 15.0058 | 2.66E-04 | 0.002988 |
| AKT3 | 1 | 2.4E+08 | 2.4E+08 | - | 9475 | AKT serine/thr | 0.21122 | 5.1894 | 14.9625 | 2.71E-04 | 0.00304 |
| EPS8L1 | 19 | 5.5E+07 | 5.5E+07 | + | 5341 | EPS8 like 1 [So | -0.2493 | 4.78641 | 14.9611 | 2.71E-04 | 0.00304 |
| SPTB | 14 | 6.5E+07 | 6.5E+07 | - | 14796 | spectrin beta, | -0.6362 | 2.20566 | 15.3621 | 2.72E-04 | 0.003049 |
| ADAM19 | 5 | 1.6E+08 | 1.6E+08 | - | 7704 | ADAM metallo | 0.43042 | 5.80399 | 30.7293 | 2.73E-04 | 0.003053 |
| LRP12 | 8 | 1E+08 | 1E+08 | - | 9777 | LDL receptor r | 0.30833 | 4.0009 | 14.9218 | 2.76E-04 | 0.003082 |
| FAT4 | 4 | 1.3E+08 | 1.3E+08 | + | 16439 | FAT atypical ca | -0.3444 | 3.68786 | 14.8268 | 2.87E-04 | 0.003203 |
| DPF1 | 19 | 3.8E+07 | 3.8E+07 | - | 4800 | double PHD fin | -0.3672 | 3.77969 | 15.2027 | 2.91E-04 | 0.003236 |
| PRICKLE3 | X | 4.9E+07 | 4.9E+07 | - | 3656 | prickle planar | -0.4626 | 2.84699 | 14.7881 | 2.92E-04 | 0.003252 |
| GRK3 | 22 | 2.6E+07 | 2.6E+07 | + | 9477 | G protein-coup | -0.1706 | 5.92667 | 14.775 | 2.94E-04 | 0.003267 |
| ARMCX4 | X | 1E+08 | 1E+08 | + | 13351 | armadillo repe | -0.2183 | 5.11199 | 14.7689 | 2.95E-04 | 0.003273 |
| RAB12 | 18 | 8609437 | 8639382 | + | 2476 | RAB12, membe | -0.1949 | 5.4332 | 14.7413 | 2.98E-04 | 0.003307 |
| STX8 | 17 | 9250471 | 9576591 | - | 3455 | syntaxin 8 [Sou | -0.309 | 3.98846 | 14.7287 | 3.00E-04 | 0.003322 |
| TP73-AS1 | 1 | 3735511 | 3747373 | - | 9701 | TP73 antisense | 0.17307 | 5.86112 | 14.7258 | 3.00E-04 | 0.003324 |
| IGSF21 | 1 | 1.8E+07 | 1.8E+07 | + | 3033 | immunoglobin | 0.24217 | 4.80688 | 14.7204 | 3.01E-04 | 0.003329 |
| GJC1 | 17 | 4.5E+07 | 4.5E+07 | - | 8716 | gap junction p | 0.12488 | 7.14685 | 14.7154 | 3.01E-04 | 0.003333 |
| ZSWIM8 | 10 | 7.4E+07 | 7.4E+07 | + | 7953 | zinc finger SWI | 0.23713 | 4.84865 | 14.7084 | 3.02E-04 | 0.00334 |
| TAPBP | R6_MHC_ | 3.3E+07 | 3.3E+07 | - | 5056 | TAP binding pr | -0.1964 | 5.43784 | 14.7009 | 3.03E-04 | 0.003349 |
| AC004890 | 7 | 1.5E+08 | 1.5E+08 | + | 4288 | zinc finger prot | 1.02736 | 0.6445 | 14.6977 | 3.04E-04 | 0.003351 |
| KLC1 | 14 | 1E+08 | 1E+08 | + | 19251 | kinesin light ch | 0.17606 | 5.82774 | 14.6869 | 3.05E-04 | 0.00336 |
| GSPT1 | 16 | 1.2E+07 | 1.2E+07 | - | 8054 | G1 to S phase | -0.1081 | 8.21091 | 14.7095 | 3.05E-04 | 0.00336 |
| PCLO | 7 | 8.3E+07 | 8.3E+07 | - | 22874 | piccolo presyn | 0.35381 | 3.6112 | 14.6655 | 3.08E-04 | 0.003381 |
| ECE1 | 1 | 2.1E+07 | 2.1E+07 | - | 7308 | endothelin con | 0.14416 | 6.60291 | 14.6601 | 3.09E-04 | 0.003386 |
| EIF1 | 17 | 4.2E+07 | 4.2E+07 | + | 3623 | eukaryotic tran | -0.1035 | 8.20539 | 14.6567 | 3.09E-04 | 0.003388 |
| TBCD | SCHR17_1 | 8.3E+07 | 8.3E+07 | + | 13537 | tubulin folding | 0.11383 | 7.56016 | 14.6556 | 3.09E-04 | 0.003388 |
| ZBTB21 | 21 | 4.2E+07 | 4.2E+07 | - | 8062 | zinc finger and | -0.2472 | 4.67401 | 14.6511 | 3.10E-04 | 0.003392 |
| MLST8 | 16 | 2204248 | 2209453 | + | 4360 | MTOR associat | 0.17845 | 5.72497 | 14.6472 | 3.10E-04 | 0.003395 |
| UBAP2L | 1 | 1.5E+08 | 1.5E+08 | + | 7503 | ubiquitin assoc | 0.1036 | 8.1079 | 14.6306 | 3.12E-04 | 0.003416 |
| SKA1 | 18 | 5E+07 | 5E+07 | + | 2900 | spindle and kin | -0.3106 | 3.94962 | 14.6275 | 3.13E-04 | 0.003418 |
| SHISAL2B | 5 | 6.5E+07 | 6.5E+07 | + | 1314 | shisa like 2B [S | 0.54672 | 2.32132 | 14.6098 | 3.15E-04 | 0.003442 |
| KCNK5 | 6 | 3.9E+07 | 3.9E+07 | - | 3783 | potassium two | -0.1531 | 6.24194 | 14.5878 | 3.18E-04 | 0.003471 |
| SPPL2B | 19 | 2328615 | 2355095 | + | 7419 | signal peptide | 0.20933 | 5.22865 | 14.5852 | 3.19E-04 | 0.003473 |
| PPA1 | 10 | 7E+07 | 7E+07 | - | 5166 | pyrophosphata | -0.1267 | 6.92642 | 14.562 | 3.22E-04 | 0.003504 |
| FASN | 17 | 8.2E+07 | 8.2E+07 | - | 9321 | fatty acid synth | -0.0829 | 10.6278 | 14.5606 | 3.22E-04 | 0.003504 |
| CD47 | 3 | 1.1E+08 | 1.1E+08 | - | 8228 | CD47 molecule | 0.29354 | 4.10193 | 14.5446 | 3.24E-04 | 0.003525 |
| HS6ST1 | 2 | 1.3E+08 | 1.3E+08 | - | 4974 | heparan sulfat | -0.1329 | 6.83072 | 14.4897 | 3.32E-04 | 0.003605 |

| | | | | | | | | | | | |
|---|---|---|---|---|---|---|---|---|---|---|---|
| COG6 | 13 | 4E+07 | 4E+07 | + | 8047 | component of | -0.263 | 4.52479 | 14.489 | 3.32E-04 | 0.003605 |
| AGPAT1 | R6_MHC_ | 3.2E+07 | 3.2E+07 | - | 3593 | 1-acylglycerol- | -0.1442 | 6.4444 | 14.4738 | 3.34E-04 | 0.003625 |
| RARS | 5 | 1.7E+08 | 1.7E+08 | + | 3417 | arginyl-tRNA sy | -0.1272 | 6.99554 | 14.4625 | 3.36E-04 | 0.003639 |
| AGPAT4-IT | 6 | 1.6E+08 | 1.6E+08 | - | 1878 | AGPAT4 intron | 0.97446 | 0.83919 | 14.4613 | 3.36E-04 | 0.003639 |
| CDC123 | 10 | 1.2E+07 | 1.2E+07 | + | 2298 | cell division cy | -0.1262 | 7.22221 | 14.5067 | 3.37E-04 | 0.003645 |
| PRKCI | 3 | 1.7E+08 | 1.7E+08 | + | 5778 | protein kinase | 0.12219 | 7.19159 | 14.4541 | 3.37E-04 | 0.003645 |
| DVL1 | 1 | 1335276 | 1349418 | - | 3882 | dishevelled seg | 0.15432 | 6.31791 | 14.4455 | 3.41E-04 | 0.003683 |
| SPG11 | 15 | 4.5E+07 | 4.5E+07 | - | 11853 | SPG11, spatacs | 0.17316 | 5.84651 | 14.4172 | 3.42E-04 | 0.003697 |
| CSNK1D | 17 | 8.2E+07 | 8.2E+07 | - | 11024 | casein kinase 1 | 0.12444 | 7.02284 | 14.3752 | 3.48E-04 | 0.003762 |
| ZNF516 | 18 | 7.6E+07 | 7.6E+07 | - | 8777 | zinc finger prot | 0.31832 | 3.91714 | 14.367 | 3.50E-04 | 0.003772 |
| ADCY8 | 8 | 1.3E+08 | 1.3E+08 | - | 4444 | adenylate cycla | -0.3101 | 3.90929 | 14.3327 | 3.55E-04 | 0.003825 |
| FCHO1 | 19 | 1.8E+07 | 1.8E+07 | + | 5694 | FCH domain or | -0.2067 | 5.2755 | 14.3118 | 3.58E-04 | 0.003853 |
| ANKRD33 | 5 | 1.1E+07 | 1.1E+07 | + | 10043 | ankyrin repeat | -0.2497 | 4.60657 | 14.311 | 3.58E-04 | 0.003853 |
| RPS15A | 16 | 1.9E+07 | 1.9E+07 | - | 6679 | ribosomal prot | 0.10094 | 8.21255 | 14.3102 | 3.58E-04 | 0.003853 |
| TNFAIP8 | 5 | 1.2E+08 | 1.2E+08 | + | 8506 | TNF alpha indu | 0.30392 | 4.00108 | 14.3048 | 3.59E-04 | 0.00386 |
| AIMP1 | 4 | 1.1E+08 | 1.1E+08 | + | 3328 | aminoacyl tRN | -0.1874 | 5.64633 | 14.2783 | 3.63E-04 | 0.003901 |
| MIB1 | 18 | 2.2E+07 | 2.2E+07 | + | 10033 | mindbomb E3 | -0.1179 | 7.31721 | 14.2764 | 3.64E-04 | 0.003901 |
| SP8 | 7 | 2.1E+07 | 2.1E+07 | - | 3718 | Sp8 transcripti | 0.32064 | 3.87549 | 14.266 | 3.65E-04 | 0.003915 |
| FA2H | 16 | 7.5E+07 | 7.5E+07 | - | 3279 | fatty acid 2-hy | -0.5426 | 2.32489 | 14.2364 | 3.70E-04 | 0.003963 |
| MYO9A | 15 | 7.2E+07 | 7.2E+07 | - | 20320 | myosin IXA [So | -0.1859 | 5.549 | 14.2346 | 3.70E-04 | 0.003963 |
| PEAK1 | 15 | 7.7E+07 | 7.7E+07 | - | 24184 | pseudopodium | 0.19567 | 5.97066 | 15.5099 | 3.71E-04 | 0.00397 |
| GSE1 | 16 | 8.5E+07 | 8.6E+07 | + | 13376 | Gse1 coiled-co | 0.11438 | 7.41115 | 14.1891 | 3.77E-04 | 0.004035 |
| GAN | 16 | 8.1E+07 | 8.1E+07 | + | 15411 | gigaxonin [Sou | -0.2398 | 4.80412 | 14.1868 | 3.78E-04 | 0.004036 |
| TMUB2 | 17 | 4.4E+07 | 4.4E+07 | + | 3709 | transmembran | 0.31747 | 3.92789 | 14.1845 | 3.78E-04 | 0.004037 |
| TPRG1L | 1 | 3625015 | 3630127 | + | 2401 | tumor protein | -0.1953 | 5.42662 | 14.1809 | 3.79E-04 | 0.00404 |
| TSPEAR | 21 | 4.4E+07 | 4.5E+07 | - | 4463 | thrombospond | -0.5743 | 2.28835 | 14.2551 | 3.81E-04 | 0.004066 |
| WNK3 | X | 5.4E+07 | 5.4E+07 | - | 11472 | WNK lysine de | -0.2076 | 5.2614 | 14.1592 | 3.82E-04 | 0.004072 |
| AC125807 | 12 | 3041437 | 3044950 | + | 3514 | novel transcrip | -0.1631 | 5.93048 | 14.1511 | 3.84E-04 | 0.004081 |
| IGDCC4 | 15 | 6.5E+07 | 6.5E+07 | - | 7619 | immunoglobul | 0.24651 | 4.59989 | 14.1308 | 3.87E-04 | 0.004105 |
| TSPAN12 | 7 | 1.2E+08 | 1.2E+08 | - | 3368 | tetraspanin 12 | 0.30273 | 3.97297 | 14.1288 | 3.87E-04 | 0.004105 |
| HYAL2 | 3 | 5E+07 | 5E+07 | - | 4421 | hyaluronidase | 0.15919 | 5.98914 | 14.1288 | 3.87E-04 | 0.004105 |
| ST6GALNA | 1 | 7.6E+07 | 7.7E+07 | + | 7212 | ST6 N-acetylga | 0.237 | 4.71647 | 14.1285 | 3.87E-04 | 0.004105 |
| HELZ | 17 | 6.7E+07 | 6.7E+07 | - | 15680 | helicase with z | -0.1752 | 5.71711 | 14.1284 | 3.87E-04 | 0.004105 |
| LAMC3 | 9 | 1.3E+08 | 1.3E+08 | + | 7502 | laminin subuni | 0.20489 | 5.2267 | 14.1233 | 3.88E-04 | 0.004111 |
| MTR | 1 | 2.4E+08 | 2.4E+08 | + | 12357 | 5-methyltetrah | 0.16671 | 5.91077 | 14.1183 | 3.89E-04 | 0.004116 |
| RALGAPA | 20 | 2E+07 | 2.1E+07 | - | 11649 | Ral GTPase act | -0.1966 | 5.36159 | 14.1169 | 3.89E-04 | 0.004116 |
| SUB1 | 5 | 3.3E+07 | 3.3E+07 | + | 9266 | SUB1 homolog | -0.1179 | 7.37522 | 14.0994 | 3.92E-04 | 0.004144 |
| PLCB1 | 20 | 8077251 | 8968360 | + | 17412 | phospholipase | 0.33039 | 3.79616 | 14.0696 | 3.97E-04 | 0.004195 |
| CYLD | 16 | 5.1E+07 | 5.1E+07 | + | 12954 | CYLD lysine 63 | -0.211 | 5.10584 | 14.0621 | 3.99E-04 | 0.004205 |
| MAPK8IP1 | 11 | 4.6E+07 | 4.6E+07 | + | 3498 | mitogen-activa | -0.1591 | 6.02274 | 14.0544 | 4.00E-04 | 0.004216 |
| SERPINB9 | 6 | 2887270 | 2903309 | - | 4143 | serpin family B | -0.2644 | 5.11688 | 15.7008 | 4.01E-04 | 0.00422 |
| GRIPAP1 | X | 4.9E+07 | 4.9E+07 | - | 6841 | GRIP1 associat | -0.1658 | 5.87581 | 14.0476 | 4.01E-04 | 0.004222 |
| CIAO3 | 16 | 729760 | 741329 | - | 8094 | cytosolic iron-s | 0.2944 | 4.02863 | 14.038 | 4.03E-04 | 0.004234 |

| | | | | | | | | | | | |
|---|---|---|---|---|---|---|---|---|---|---|---|
| TWSG1 | 18 | 9334767 | 9402420 | + | 4435 | twisted gastru | -0.196 | 5.35826 | 14.0336 | 4.04E-04 | 0.004238 |
| XRN1 | 3 | 1.4E+08 | 1.4E+08 | - | 11982 | 5'-3' exoribonu | 0.25008 | 4.74453 | 14.1091 | 4.06E-04 | 0.004259 |
| NOP14-AS | 4 | 2934899 | 2961738 | + | 7195 | NOP14 antisen | 0.40022 | 3.0822 | 14.0191 | 4.06E-04 | 0.004259 |
| PASK | 2 | 2.4E+08 | 2.4E+08 | - | 10686 | PAS domain co | 0.15445 | 6.11521 | 13.9986 | 4.10E-04 | 0.004293 |
| PWP2 | 21 | 4.4E+07 | 4.4E+07 | + | 6839 | PWP2, small su | -0.3283 | 3.81149 | 13.9941 | 4.10E-04 | 0.004296 |
| MPHOSPH | 2 | 7.1E+07 | 7.1E+07 | + | 4355 | M-phase phos | -0.1996 | 5.29269 | 13.9938 | 4.10E-04 | 0.004296 |
| ARPP19 | 15 | 5.3E+07 | 5.3E+07 | - | 6917 | cAMP regulate | -0.1104 | 7.68453 | 13.9796 | 4.13E-04 | 0.004319 |
| ANO8 | 19 | 1.7E+07 | 1.7E+07 | - | 5005 | anoctamin 8 [S | 0.20637 | 5.38606 | 14.0726 | 4.14E-04 | 0.004324 |
| RIPOR3 | 20 | 5.1E+07 | 5.1E+07 | - | 6008 | RIPOR family n | -0.5077 | 2.42529 | 13.9541 | 4.18E-04 | 0.004361 |
| HSP90AB1 | 6 | 4.4E+07 | 4.4E+07 | + | 2980 | heat shock pro | -0.0736 | 11.8492 | 13.9401 | 4.20E-04 | 0.004384 |
| RADX | X | 1.1E+08 | 1.1E+08 | + | 4850 | RPA1 related s | 0.52 | 2.35981 | 13.9038 | 4.27E-04 | 0.00445 |
| CDK5RAP1 | 9 | 1.2E+08 | 1.2E+08 | - | 10080 | CDK5 regulato | 0.13184 | 6.66163 | 13.8987 | 4.28E-04 | 0.004457 |
| HERPUD2 | 7 | 3.6E+07 | 3.6E+07 | - | 3246 | HERPUD family | 0.25382 | 4.53481 | 13.8943 | 4.29E-04 | 0.004462 |
| RGL2 | 6 | 3.3E+07 | 3.3E+07 | - | 4159 | ral guanine nu | 0.18301 | 5.64067 | 13.889 | 4.29E-04 | 0.004468 |
| NANS | 9 | 9.8E+07 | 9.8E+07 | + | 4931 | N-acetylneura | -0.1844 | 5.62177 | 13.8878 | 4.30E-04 | 0.004468 |
| C9orf135 | 9 | 7E+07 | 7E+07 | + | 1250 | chromosome 9 | -0.5824 | 2.19712 | 13.9076 | 4.31E-04 | 0.004477 |
| IMPACT | 18 | 2.4E+07 | 2.4E+07 | + | 4896 | impact RWD d | -0.2109 | 5.82824 | 15.5469 | 4.32E-04 | 0.004482 |
| GNG5 | 1 | 8.4E+07 | 8.5E+07 | - | 1128 | G protein subu | 0.20145 | 5.23393 | 13.8718 | 4.33E-04 | 0.004486 |
| AFF4 | 5 | 1.3E+08 | 1.3E+08 | - | 11281 | AF4/FMR2 fam | 0.1162 | 7.30746 | 13.8606 | 4.35E-04 | 0.004504 |
| AXIN2 | 17 | 6.6E+07 | 6.6E+07 | - | 5227 | axin 2 [Source: | -0.2535 | 4.53909 | 13.8573 | 4.35E-04 | 0.004507 |
| ATXN3 | 14 | 9.2E+07 | 9.2E+07 | - | 9665 | ataxin 3 [Sourc | -0.2645 | 4.34288 | 13.8462 | 4.37E-04 | 0.004523 |
| ABCC4 | 13 | 9.5E+07 | 9.5E+07 | - | 8629 | ATP binding ca | 0.26695 | 4.37547 | 13.8428 | 4.38E-04 | 0.004524 |
| PPM1F | 22 | 2.2E+07 | 2.2E+07 | - | 10089 | protein phosp | 0.18409 | 5.56931 | 13.8423 | 4.38E-04 | 0.004524 |
| NIN | 14 | 5.1E+07 | 5.1E+07 | - | 13683 | ninein [Source | 0.19796 | 5.26351 | 13.8128 | 4.44E-04 | 0.004579 |
| SETX | 9 | 1.3E+08 | 1.3E+08 | - | 12141 | senataxin [Sou | 0.1271 | 6.96908 | 13.7781 | 4.51E-04 | 0.004644 |
| SFXN1 | 5 | 1.8E+08 | 1.8E+08 | + | 7407 | sideroflexin 1 [ | 0.14784 | 6.29368 | 13.7633 | 4.53E-04 | 0.004671 |
| MYRF | 11 | 6.2E+07 | 6.2E+07 | + | 10680 | myelin regulat | 0.30832 | 3.86427 | 13.7568 | 4.55E-04 | 0.004681 |
| PDK3 | X | 2.4E+07 | 2.5E+07 | + | 13555 | pyruvate dehy | -0.1948 | 5.40433 | 13.7547 | 4.55E-04 | 0.004682 |
| FRAT2 | 10 | 9.7E+07 | 9.7E+07 | - | 2213 | FRAT2, WNT si | -0.1402 | 6.60235 | 13.7573 | 4.58E-04 | 0.004709 |
| CIRBP | 19 | 1259384 | 1274880 | + | 6368 | cold inducible | 0.13132 | 6.71478 | 13.7382 | 4.58E-04 | 0.004709 |
| AC091820 | 5 | 1.7E+08 | 1.7E+08 | - | 648 | novel transcrip | -0.5721 | 2.07287 | 13.7307 | 4.60E-04 | 0.004719 |
| SYT17 | 16 | 1.9E+07 | 1.9E+07 | + | 7545 | synaptotagmin | -0.481 | 2.66175 | 13.7298 | 4.60E-04 | 0.004719 |
| TBL1XR1 | 3 | 1.8E+08 | 1.8E+08 | - | 11603 | transducin bet | -0.1565 | 6.01561 | 13.7228 | 4.61E-04 | 0.00473 |
| CTDSPL2 | 15 | 4.4E+07 | 4.5E+07 | + | 9277 | CTD small pho | 0.16394 | 5.87387 | 13.6994 | 4.66E-04 | 0.004775 |
| SLC19A1 | 21 | 4.5E+07 | 4.6E+07 | - | 11235 | solute carrier f | -0.1771 | 5.84855 | 13.7932 | 4.67E-04 | 0.004781 |
| DHCR24 | 1 | 5.5E+07 | 5.5E+07 | - | 6214 | 24-dehydrocho | -0.0849 | 9.75234 | 13.6929 | 4.68E-04 | 0.004781 |
| NET1 | 10 | 5412557 | 5459056 | + | 4762 | neuroepithelia | -0.1321 | 6.75453 | 13.6816 | 4.70E-04 | 0.004801 |
| WDFY1 | 2 | 2.2E+08 | 2.2E+08 | - | 5485 | WD repeat and | -0.1184 | 7.15684 | 13.674 | 4.71E-04 | 0.004814 |
| MAN2A1 | 5 | 1.1E+08 | 1.1E+08 | + | 11815 | mannosidase a | 0.11708 | 7.27815 | 13.671 | 4.72E-04 | 0.004815 |
| SIRT1 | 10 | 6.8E+07 | 6.8E+07 | + | 4564 | sirtuin 1 [Sourc | -0.2076 | 6.43636 | 17.3637 | 4.72E-04 | 0.004815 |
| CTNNA3 | 10 | 6.6E+07 | 6.8E+07 | - | 11157 | catenin alpha 3 | 0.44825 | 2.88448 | 13.6686 | 4.72E-04 | 0.004815 |
| RRM2B | 8 | 1E+08 | 1E+08 | - | 5548 | ribonucleotide | -0.1248 | 6.97625 | 13.6585 | 4.75E-04 | 0.004833 |
| GABPB1-A | 15 | 5E+07 | 5E+07 | + | 17179 | GABPB1 antise | -0.2045 | 5.28661 | 13.6312 | 4.80E-04 | 0.004887 |

| | | | | | | | | | | | |
|---|---|---|---|---|---|---|---|---|---|---|---|
| ZNF92 | 7 | 6.5E+07 | 6.5E+07 | + | 3512 | zinc finger pro | -0.259 | 4.59632 | 13.6954 | 4.84E-04 | 0.004924 |
| AC012085 | 12 | 9.4E+07 | 9.4E+07 | - | 765 | phosphoglycer | 0.20297 | 5.19709 | 13.6101 | 4.85E-04 | 0.004924 |
| DOK1 | 2 | 7.5E+07 | 7.5E+07 | + | 3049 | docking protei | 0.49172 | 2.46048 | 13.6046 | 4.86E-04 | 0.00493 |
| FAM136A | 2 | 7E+07 | 7E+07 | - | 3086 | family with seq | -0.1301 | 6.84566 | 13.6041 | 4.86E-04 | 0.00493 |
| UBE2S | 19 | 5.5E+07 | 5.5E+07 | - | 2799 | ubiquitin conju | 0.10391 | 8.01872 | 13.6022 | 4.86E-04 | 0.004931 |
| PEX26 | 22 | 1.8E+07 | 1.8E+07 | + | 20045 | peroxisomal bi | -0.1418 | 6.36859 | 13.5989 | 4.87E-04 | 0.004931 |
| ACAN | 15 | 8.9E+07 | 8.9E+07 | + | 10909 | aggrecan [Sour | -0.8289 | 1.1335 | 13.5814 | 4.91E-04 | 0.004962 |
| FKBP1B | 2 | 2.4E+07 | 2.4E+07 | + | 1677 | FKBP prolyl iso | 0.25525 | 4.55563 | 13.5602 | 4.95E-04 | 0.005004 |
| SUCLA2-A | 13 | 4.8E+07 | 4.8E+07 | + | 552 | SUCLA2 antise | -1.0065 | 0.74801 | 13.5474 | 4.98E-04 | 0.005029 |
| SGCB | 4 | 5.2E+07 | 5.2E+07 | - | 4541 | sarcoglycan be | -0.203 | 5.18996 | 13.5407 | 4.99E-04 | 0.00504 |
| CLN8 | HSCHR8_7 | 1755340 | 1786132 | + | 9459 | CLN8, transme | -0.1859 | 5.57976 | 13.5125 | 5.06E-04 | 0.005098 |
| CDC6 | 17 | 4E+07 | 4E+07 | + | 5311 | cell division cy | 0.12919 | 6.82918 | 13.5046 | 5.07E-04 | 0.005112 |
| BAHCC1 | 17 | 8.1E+07 | 8.1E+07 | + | 13473 | BAH domain ar | 0.60656 | 1.88594 | 13.4953 | 5.09E-04 | 0.005129 |
| ERCC6 | 10 | 4.9E+07 | 5E+07 | - | 17716 | ERCC excision | -0.3097 | 3.86086 | 13.474 | 5.14E-04 | 0.005173 |
| G3BP1 | 5 | 1.5E+08 | 1.5E+08 | + | 12406 | G3BP stress gr | -0.0912 | 9.02785 | 13.4671 | 5.16E-04 | 0.005183 |
| CLNS1A | 11 | 7.8E+07 | 7.8E+07 | - | 4592 | chloride nucle | -0.1254 | 6.81083 | 13.4664 | 5.16E-04 | 0.005183 |
| SCCPDH | 1 | 2.5E+08 | 2.5E+08 | + | 2141 | saccharopine d | 0.21868 | 5.00869 | 13.4619 | 5.17E-04 | 0.005188 |
| NME7 | 1 | 1.7E+08 | 1.7E+08 | - | 5718 | NME/NM23 fa | -0.3483 | 3.71126 | 13.6231 | 5.17E-04 | 0.005188 |
| SEZ6L2 | 16 | 3E+07 | 3E+07 | - | 4110 | seizure related | -0.1285 | 6.74675 | 13.4544 | 5.19E-04 | 0.0052 |
| KLHL5 | 4 | 3.9E+07 | 3.9E+07 | + | 5144 | kelch like fami | -0.1965 | 5.26521 | 13.4402 | 5.22E-04 | 0.005224 |
| SDC4 | 20 | 4.5E+07 | 4.5E+07 | - | 2613 | syndecan 4 [So | -0.1576 | 5.98419 | 13.437 | 5.22E-04 | 0.005228 |
| JPH4 | 14 | 2.4E+07 | 2.4E+07 | - | 5297 | junctophilin 4 | -0.199 | 5.36362 | 13.4228 | 5.26E-04 | 0.005254 |
| CHD3 | 17 | 7884806 | 7912760 | + | 9758 | chromodomair | -0.1679 | 5.79874 | 13.4218 | 5.26E-04 | 0.005254 |
| CDS2 | 20 | 5126879 | 5197887 | + | 11800 | CDP-diacylglyc | -0.135 | 6.62294 | 13.421 | 5.26E-04 | 0.005254 |
| ACADVL | 17 | 7217125 | 7225266 | + | 4820 | acyl-CoA dehy | 0.13994 | 6.43753 | 13.4133 | 5.28E-04 | 0.005266 |
| KCNJ6 | 21 | 3.8E+07 | 3.8E+07 | - | 19979 | potassium volt | -0.5455 | 2.18892 | 13.4105 | 5.29E-04 | 0.005266 |
| SEC14L1 | 17 | 7.7E+07 | 7.7E+07 | + | 10356 | SEC14 like lipid | -0.1237 | 6.90538 | 13.4102 | 5.29E-04 | 0.005266 |
| EFHD2 | 1 | 1.5E+07 | 1.5E+07 | + | 2534 | EF-hand doma | -0.207 | 5.15575 | 13.4093 | 5.29E-04 | 0.005266 |
| ASPM | 1 | 2E+08 | 2E+08 | - | 10888 | abnormal spin | 0.11022 | 7.48965 | 13.3803 | 5.36E-04 | 0.005329 |
| CLIP1 | 12 | 1.2E+08 | 1.2E+08 | - | 18559 | CAP-Gly domai | 0.18308 | 5.58695 | 13.3689 | 5.38E-04 | 0.005352 |
| ADARB1 | 21 | 4.5E+07 | 4.5E+07 | + | 10049 | adenosine dea | -0.2524 | 4.69088 | 13.4963 | 5.39E-04 | 0.005358 |
| SEC13 | 3 | 1E+07 | 1E+07 | - | 5380 | SEC13 homolo | -0.1239 | 6.97029 | 13.3601 | 5.40E-04 | 0.005365 |
| MYSM1 | 1 | 5.9E+07 | 5.9E+07 | - | 8925 | Myb like, SWIR | 0.12757 | 6.8103 | 13.3559 | 5.41E-04 | 0.005368 |
| CNKSR2 | X | 2.1E+07 | 2.2E+07 | + | 31865 | connector enh | -0.3435 | 3.4914 | 13.3493 | 5.43E-04 | 0.005379 |
| ZNF90 | 19 | 2E+07 | 2E+07 | + | 6088 | zinc finger pro | -0.1664 | 5.78426 | 13.3478 | 5.43E-04 | 0.005379 |
| ITM2C | 2 | 2.3E+08 | 2.3E+08 | + | 2569 | integral memb | 0.09082 | 9.25502 | 13.3338 | 5.47E-04 | 0.005407 |
| AMMECR | X | 1.1E+08 | 1.1E+08 | - | 6362 | Alport syndron | 0.18518 | 5.463 | 13.3321 | 5.47E-04 | 0.005407 |
| CFDP1 | 16 | 7.5E+07 | 7.5E+07 | - | 3335 | craniofacial de | -0.1684 | 5.81591 | 13.3308 | 5.47E-04 | 0.005407 |
| DHX32 | 10 | 1.3E+08 | 1.3E+08 | - | 3948 | DEAH-box heli | 0.24107 | 4.5943 | 13.3296 | 5.48E-04 | 0.005407 |
| FP565260 | 21 | 5130871 | 5154734 | - | 7665 | periodic trypto | -0.3347 | 3.59509 | 13.325 | 5.49E-04 | 0.00541 |
| TCF7L2 | 10 | 1.1E+08 | 1.1E+08 | + | 6739 | transcription fa | -0.157 | 5.99645 | 13.325 | 5.49E-04 | 0.00541 |
| MT-RNR1 | MT | 648 | 1601 | + | 954 | mitochondriall | 0.0916 | 9.03773 | 13.3086 | 5.53E-04 | 0.005446 |
| RRM2 | 2 | 1E+07 | 1E+07 | + | 9155 | ribonucleotide | 0.12035 | 7.14283 | 13.3067 | 5.53E-04 | 0.005446 |

| | | | | | | | | | | | |
|---|---|---|---|---|---|---|---|---|---|---|---|
| OR2A9P | 7 | 1.4E+08 | 1.4E+08 | + | 3337 | olfactory recep | -0.6778 | 1.64736 | 13.3012 | 5.54E-04 | 0.005456 |
| KIRREL1 | 1 | 1.6E+08 | 1.6E+08 | + | 7872 | kirre like neph | 0.13637 | 6.4816 | 13.2838 | 5.59E-04 | 0.005493 |
| ATP5F1E | 20 | 5.9E+07 | 5.9E+07 | - | 5081 | ATP synthase F | 0.18044 | 5.54233 | 13.2788 | 5.60E-04 | 0.005502 |
| ONECUT2 | 18 | 5.7E+07 | 5.7E+07 | + | 16546 | one cut homeo | -0.4486 | 2.77274 | 13.2736 | 5.61E-04 | 0.00551 |
| UBXN1 | 11 | 6.3E+07 | 6.3E+07 | - | 2532 | UBX domain pr | 0.13713 | 6.45602 | 13.2612 | 5.64E-04 | 0.005537 |
| LINC0196 | 2 | 2.2E+08 | 2.2E+08 | + | 3148 | long intergenic | 0.55135 | 2.14948 | 13.2347 | 5.71E-04 | 0.005597 |
| ELK3 | 12 | 9.6E+07 | 9.6E+07 | + | 4644 | ELK3, ETS trans | 0.22142 | 5.13533 | 13.4305 | 5.78E-04 | 0.005662 |
| ZMYND8 | 20 | 4.7E+07 | 4.7E+07 | - | 7625 | zinc finger MYI | -0.1054 | 7.78473 | 13.2038 | 5.78E-04 | 0.005666 |
| MX1 | 21 | 4.1E+07 | 4.1E+07 | + | 8616 | MX dynamin li | -0.455 | 2.69021 | 13.1982 | 5.80E-04 | 0.005676 |
| ACOT7 | 1 | 6264269 | 6394391 | - | 4124 | acyl-CoA thioe | 0.11217 | 7.49905 | 13.1899 | 5.82E-04 | 0.005692 |
| BCCIP | 10 | 1.3E+08 | 1.3E+08 | + | 4496 | BRCA2 and CD | -0.1272 | 6.77749 | 13.1849 | 5.83E-04 | 0.005701 |
| OGT | X | 7.2E+07 | 7.2E+07 | + | 9786 | O-linked N-ace | 0.10235 | 7.96268 | 13.1817 | 5.84E-04 | 0.005701 |
| TRAPPC12 | HSCHR2_1 | 3401777 | 3486461 | + | 9193 | trafficking prot | 0.17498 | 5.63109 | 13.1816 | 5.84E-04 | 0.005701 |
| FLYWCH1 | 16 | 2911937 | 2951208 | + | 8419 | FLYWCH-type z | 0.1999 | 5.24962 | 13.1728 | 5.86E-04 | 0.005719 |
| ZNF808 | 19 | 5.3E+07 | 5.3E+07 | + | 5008 | zinc finger prot | -0.4622 | 2.73757 | 13.1665 | 5.88E-04 | 0.005728 |
| E2F5 | 8 | 8.5E+07 | 8.5E+07 | + | 5999 | E2F transcripti | 0.19626 | 5.26601 | 13.1664 | 5.88E-04 | 0.005728 |
| DNAJC5 | 20 | 6.4E+07 | 6.4E+07 | + | 5343 | DnaJ heat shoc | 0.13465 | 6.51108 | 13.1646 | 5.88E-04 | 0.005728 |
| THY1 | 11 | 1.2E+08 | 1.2E+08 | - | 5925 | Thy-1 cell surfa | 0.09574 | 8.67068 | 13.16 | 5.90E-04 | 0.005736 |
| MTHFD2 | 2 | 7.4E+07 | 7.4E+07 | + | 4822 | methylenetetr | -0.1068 | 7.80949 | 13.1561 | 5.91E-04 | 0.005742 |
| B3GALNT | 3 | 1.6E+08 | 1.6E+08 | - | 7816 | beta-1,3-N-ace | 0.38767 | 3.12013 | 13.1462 | 5.93E-04 | 0.005763 |
| VPS52 | R6_MHC_ | 3.3E+07 | 3.3E+07 | - | 4493 | VPS52, GARP c | 0.19541 | 5.36282 | 13.1413 | 5.94E-04 | 0.005771 |
| U2AF2 | 19 | 5.6E+07 | 5.6E+07 | + | 4115 | U2 small nucle | 0.09742 | 8.31563 | 13.1363 | 5.96E-04 | 0.00578 |
| IGF2BP2 | 3 | 1.9E+08 | 1.9E+08 | - | 4735 | insulin like gro | 0.10782 | 7.48261 | 13.1295 | 5.98E-04 | 0.005793 |
| KIAA0895 | 16 | 6.7E+07 | 6.7E+07 | - | 4783 | KIAA0895 like | 0.28178 | 4.08331 | 13.1282 | 5.98E-04 | 0.005793 |
| WIPF2 | 17 | 4E+07 | 4E+07 | + | 8123 | WAS/WASL int | -0.1753 | 5.59096 | 13.1196 | 6.00E-04 | 0.005811 |
| SLCO1A2 | 12 | 2.1E+07 | 2.1E+07 | - | 11520 | solute carrier o | 0.33817 | 3.49471 | 13.1121 | 6.02E-04 | 0.005826 |
| NOTCH2 | 1 | 1.2E+08 | 1.2E+08 | - | 19930 | notch 2 [Sourc | -0.1174 | 7.02981 | 13.1103 | 6.03E-04 | 0.005827 |
| SOX12 | 20 | 325595 | 330224 | + | 4630 | SRY-box 12 [So | -0.1101 | 7.45397 | 13.1036 | 6.04E-04 | 0.00584 |
| SDK2 | 17 | 7.3E+07 | 7.4E+07 | - | 11963 | sidekick cell ad | -0.3236 | 3.76672 | 13.0971 | 6.06E-04 | 0.005853 |
| IRS2 | 13 | 1.1E+08 | 1.1E+08 | - | 8156 | insulin recepto | -0.1813 | 5.52424 | 13.0885 | 6.08E-04 | 0.005867 |
| ADA2 | 22 | 1.7E+07 | 1.7E+07 | - | 8704 | adenosine dea | -0.2718 | 4.2816 | 13.0883 | 6.08E-04 | 0.005867 |
| BLCAP | 20 | 3.7E+07 | 3.8E+07 | - | 7869 | BLCAP, apopto | 0.15483 | 5.97896 | 13.0766 | 6.12E-04 | 0.005894 |
| PLEKHG5 | 1 | 6466092 | 6520061 | - | 7053 | pleckstrin hom | 0.19259 | 5.2692 | 13.0711 | 6.13E-04 | 0.005904 |
| NOMO1 | 16 | 1.5E+07 | 1.5E+07 | + | 5776 | NODAL modula | -0.1584 | 5.92125 | 13.0646 | 6.15E-04 | 0.005917 |
| FAM53B | 10 | 1.2E+08 | 1.2E+08 | - | 6533 | family with seq | 0.21737 | 4.89493 | 13.0607 | 6.16E-04 | 0.005919 |
| EPHB2 | 1 | 2.3E+07 | 2.3E+07 | + | 12516 | EPH receptor B | -0.2349 | 4.72936 | 13.0451 | 6.20E-04 | 0.005955 |
| PLCB3 | 11 | 6.4E+07 | 6.4E+07 | + | 4964 | phospholipase | 0.1106 | 7.34001 | 13.0361 | 6.22E-04 | 0.005975 |
| EEFSEC | 3 | 1.3E+08 | 1.3E+08 | + | 2394 | eukaryotic elor | -0.2608 | 4.36955 | 13.0247 | 6.26E-04 | 0.006001 |
| FOXO3 | 6 | 1.1E+08 | 1.1E+08 | + | 7670 | forkhead box O | 0.21629 | 4.92711 | 13.0072 | 6.30E-04 | 0.006043 |
| NDUFAB1 | 16 | 2.4E+07 | 2.4E+07 | - | 1548 | NADH:ubiquin | -0.1374 | 6.35681 | 12.9538 | 0.0006454 | 0.006182 |
| ACTB | 7 | 5527148 | 5563784 | - | 3413 | actin beta [Sou | -0.074 | 12.1561 | 12.9554 | 6.51E-04 | 0.006229 |
| NMI | 2 | 1.5E+08 | 1.5E+08 | - | 3480 | N-myc and STA | -0.4808 | 2.48701 | 12.916 | 6.56E-04 | 0.006277 |
| TMBIM1 | 2 | 2.2E+08 | 2.2E+08 | - | 5093 | transmembran | 0.16811 | 5.7538 | 12.9107 | 6.58E-04 | 0.006288 |

| | | | | | | | | | | | |
|---|---|---|---|---|---|---|---|---|---|---|---|
| CDH4 | SCHR20_1 | 6.2E+07 | 6.2E+07 | + | 7516 | cadherin 4 [So | -0.2653 | 4.23387 | 12.9026 | 6.60E-04 | 0.006305 |
| APBB1 | 11 | 6395124 | 6419414 | - | 5244 | amyloid beta p | -0.1767 | 5.53823 | 12.9013 | 6.60E-04 | 0.006305 |
| LINC0177 | 1 | 1430539 | 1434573 | - | 851 | long intergenic | 0.93209 | 0.75663 | 12.8979 | 6.61E-04 | 0.006308 |
| AC106038 | 8 | 9.1E+07 | 9.1E+07 | + | 3808 | novel transcrip | -0.5407 | 2.245 | 12.8972 | 6.62E-04 | 0.006308 |
| CCDC150 | 2 | 2E+08 | 2E+08 | + | 9678 | coiled-coil dom | 0.25716 | 4.3263 | 12.8944 | 6.62E-04 | 0.006312 |
| TAF10 | 11 | 6606294 | 6612539 | - | 5820 | TATA-box bind | 0.2587 | 4.29678 | 12.8904 | 6.64E-04 | 0.006319 |
| PEG13 | 8 | 1.4E+08 | 1.4E+08 | - | 5650 | paternally exp | -0.6051 | 1.93337 | 12.886 | 6.65E-04 | 0.006327 |
| INF2 | 14 | 1E+08 | 1E+08 | + | 9488 | inverted formi | 0.13416 | 6.51361 | 12.8817 | 6.66E-04 | 0.006335 |
| RHOB | 2 | 2E+07 | 2E+07 | + | 2372 | ras homolog fa | 0.30105 | 4.60482 | 14.4941 | 6.68E-04 | 0.006348 |
| DCP2 | 5 | 1.1E+08 | 1.1E+08 | + | 11176 | decapping mRI | -0.1114 | 7.31277 | 12.8738 | 6.68E-04 | 0.006348 |
| UNC13B | 9 | 3.5E+07 | 3.5E+07 | + | 17916 | unc-13 homolo | -0.1201 | 7.04077 | 12.8613 | 6.72E-04 | 0.006379 |
| SLC25A23 | 19 | 6436079 | 6465203 | - | 4881 | solute carrier f | -0.1446 | 6.19081 | 12.8516 | 6.75E-04 | 0.006402 |
| SERINC3 | 20 | 4.4E+07 | 4.5E+07 | - | 4649 | serine incorpo | -0.147 | 6.17873 | 12.8442 | 6.77E-04 | 0.006416 |
| CD55 | 1 | 2.1E+08 | 2.1E+08 | + | 5908 | CD55 molecule | -0.2466 | 4.51816 | 12.8436 | 6.77E-04 | 0.006416 |
| PAPPA2 | 1 | 1.8E+08 | 1.8E+08 | + | 10926 | pappalysin 2 [S | 0.85346 | 1.00431 | 12.8372 | 6.79E-04 | 0.00643 |
| GOLGA2P | SCHR15_5 | 8.3E+07 | 8.3E+07 | - | 1758 | GOLGA2 pseud | 0.3141 | 3.75368 | 12.8311 | 6.81E-04 | 0.00644 |
| SYNJ1 | 21 | 3.3E+07 | 3.3E+07 | - | 8070 | synaptojanin 1 | -0.4013 | 3.09416 | 12.8309 | 6.81E-04 | 0.00644 |
| ZNF532 | 18 | 5.9E+07 | 5.9E+07 | + | 11951 | zinc finger pro | -0.0995 | 7.93587 | 12.8278 | 6.82E-04 | 0.006444 |
| B3GALT5- | 21 | 4E+07 | 4E+07 | - | 2516 | B3GALT5 antis | -0.2316 | 4.71201 | 12.823 | 6.84E-04 | 0.006453 |
| NOX4 | 11 | 8.9E+07 | 8.9E+07 | - | 6099 | NADPH oxidase | 0.62231 | 1.74108 | 12.8139 | 6.86E-04 | 0.006472 |
| DENND1A | 9 | 1.2E+08 | 1.2E+08 | - | 10256 | DENN domain | 0.18263 | 5.44719 | 12.8135 | 6.86E-04 | 0.006472 |
| NDUFB7 | 19 | 1.5E+07 | 1.5E+07 | - | 535 | NADH:ubiquin | -0.1384 | 6.32617 | 12.8047 | 6.89E-04 | 0.006493 |
| DVL3 | 3 | 1.8E+08 | 1.8E+08 | + | 6368 | dishevelled seg | 0.12094 | 6.88931 | 12.802 | 6.90E-04 | 0.006496 |
| ZNF613 | 19 | 5.2E+07 | 5.2E+07 | + | 3853 | zinc finger pro | -0.3073 | 3.75226 | 12.7946 | 6.92E-04 | 0.006513 |
| IGSF9 | 1 | 1.6E+08 | 1.6E+08 | - | 6637 | immunoglobul | -0.1649 | 5.87399 | 12.7886 | 6.94E-04 | 0.006525 |
| NF2 | 22 | 3E+07 | 3E+07 | + | 7170 | neurofibromin | -0.1975 | 5.27339 | 12.7876 | 6.94E-04 | 0.006525 |
| ZNF580 | 19 | 5.6E+07 | 5.6E+07 | + | 2604 | zinc finger pro | 0.26238 | 4.29999 | 12.7814 | 6.96E-04 | 0.006539 |
| TXNDC17 | 17 | 6640758 | 6644541 | + | 3784 | thioredoxin do | -0.1871 | 5.48396 | 12.7748 | 6.98E-04 | 0.006553 |
| AGAP1 | 2 | 2.4E+08 | 2.4E+08 | + | 13956 | ArfGAP with G | 0.14048 | 6.41786 | 12.77 | 7.00E-04 | 0.006563 |
| ATRX | X | 7.8E+07 | 7.8E+07 | - | 19266 | ATRX, chromat | -0.1094 | 7.49963 | 12.7683 | 7.00E-04 | 0.006563 |
| HS3ST3A1 | 17 | 1.3E+07 | 1.4E+07 | - | 4287 | heparan sulfat | -0.4187 | 2.98292 | 12.764 | 7.02E-04 | 0.006568 |
| RPSAP45 | 6 | 1.1E+08 | 1.1E+08 | - | 874 | ribosomal prot | -0.3276 | 3.65819 | 12.7639 | 7.02E-04 | 0.006568 |
| KIT | 4 | 5.5E+07 | 5.5E+07 | + | 5563 | KIT proto-onco | -0.2225 | 4.83598 | 12.7567 | 7.04E-04 | 0.006578 |
| SLC27A3 | SCHR1_1_ | 1.5E+08 | 1.5E+08 | + | 5470 | solute carrier f | -0.1907 | 5.28007 | 12.7561 | 7.04E-04 | 0.006578 |
| SNX12 | X | 7.1E+07 | 7.1E+07 | - | 2595 | sorting nexin 1 | -0.1306 | 6.53903 | 12.7559 | 7.04E-04 | 0.006578 |
| TRDN | 6 | 1.2E+08 | 1.2E+08 | - | 6700 | triadin [Source | -0.1871 | 5.36149 | 12.7535 | 7.05E-04 | 0.006581 |
| NAP1L4 | 11 | 2944431 | 2992377 | - | 5200 | nucleosome as | 0.10241 | 7.73698 | 12.7491 | 7.06E-04 | 0.006589 |
| ADH5 | 4 | 9.9E+07 | 9.9E+07 | - | 4866 | alcohol dehydr | 0.09329 | 8.51887 | 12.7453 | 7.07E-04 | 0.006596 |
| DDX46 | 5 | 1.3E+08 | 1.3E+08 | + | 9205 | DEAD-box heli | -0.105 | 7.71041 | 12.7435 | 7.08E-04 | 0.006597 |
| MESD | 15 | 8.1E+07 | 8.1E+07 | - | 7455 | mesoderm dev | 0.14347 | 6.21942 | 12.7282 | 7.13E-04 | 0.006637 |
| SEMA4A | 1 | 1.6E+08 | 1.6E+08 | + | 6771 | semaphorin 4A | -0.2963 | 3.87527 | 12.7122 | 7.18E-04 | 0.006679 |
| MDH2 | 7 | 7.6E+07 | 7.6E+07 | + | 3641 | malate dehydr | -0.098 | 8.03654 | 12.711 | 7.18E-04 | 0.006679 |
| MDFIC | 7 | 1.1E+08 | 1.2E+08 | + | 5252 | MyoD family ir | 0.27371 | 4.14753 | 12.706 | 7.20E-04 | 0.006682 |

| | | | | | | | | | | | |
|---|---|---|---|---|---|---|---|---|---|---|---|
| THOC1 | 18 | 214520 | 268050 | - | 8102 | THO complex 1 | -0.1758 | 5.6248 | 12.706 | 7.20E-04 | 0.006682 |
| RPL22 | 1 | 6185020 | 6209389 | - | 4665 | ribosomal prot | 0.11019 | 7.3214 | 12.7055 | 7.20E-04 | 0.006682 |
| SESN1 | 6 | 1.1E+08 | 1.1E+08 | - | 5121 | sestrin 1 [Sour | 0.19016 | 5.38356 | 12.6947 | 7.23E-04 | 0.00671 |
| PFKP | 10 | 3066333 | 3137718 | + | 5554 | phosphofructo | -0.1098 | 7.61153 | 12.6705 | 7.37E-04 | 0.006831 |
| NLRP2 | 19LRC_LR | 5.5E+07 | 5.5E+07 | + | 6313 | NLR family pyr | -0.7197 | 1.37368 | 12.6447 | 7.39E-04 | 0.006851 |
| BCHE | 3 | 1.7E+08 | 1.7E+08 | - | 3072 | butyrylcholine | 0.77792 | 1.42905 | 12.9722 | 7.41E-04 | 0.006859 |
| PREP | 6 | 1.1E+08 | 1.1E+08 | - | 8917 | prolyl endopep | 0.12708 | 6.65948 | 12.6373 | 7.42E-04 | 0.006864 |
| SSH3 | 11 | 6.7E+07 | 6.7E+07 | + | 3611 | slingshot prote | 0.23305 | 4.63933 | 12.6288 | 7.45E-04 | 0.006886 |
| PARK7 | 1 | 7954291 | 7985505 | + | 2397 | Parkinsonism a | -0.1035 | 7.67554 | 12.6255 | 7.46E-04 | 0.006891 |
| ANKRD50 | 4 | 1.2E+08 | 1.2E+08 | - | 8798 | ankyrin repeat | 0.14645 | 6.09717 | 12.6238 | 7.46E-04 | 0.006892 |
| ST6GAL2 | 2 | 1.1E+08 | 1.1E+08 | - | 7708 | ST6 beta-galac | 0.25881 | 4.28996 | 12.6197 | 7.48E-04 | 0.0069 |
| DNAJB6 | 7 | 1.6E+08 | 1.6E+08 | + | 14981 | DnaJ heat shoc | -0.0899 | 8.74202 | 12.617 | 7.49E-04 | 0.006902 |
| COPG2 | 7 | 1.3E+08 | 1.3E+08 | - | 4139 | coatomer prot | 0.12094 | 6.82413 | 12.6155 | 7.49E-04 | 0.006902 |
| RECQL4 | 8 | 1.4E+08 | 1.4E+08 | - | 4236 | RecQ like helic | 0.14186 | 6.27268 | 12.6146 | 7.49E-04 | 0.006902 |
| THADA | 2 | 4.3E+07 | 4.4E+07 | - | 14220 | THADA, armad | 0.1822 | 5.41225 | 12.6094 | 7.51E-04 | 0.006913 |
| SFN | 1 | 2.7E+07 | 2.7E+07 | + | 1308 | stratifin [Sourc | 0.33691 | 3.47916 | 12.6082 | 7.51E-04 | 0.006913 |
| PTPN9 | 15 | 7.5E+07 | 7.6E+07 | - | 8632 | protein tyrosin | 0.13393 | 6.4459 | 12.6056 | 7.52E-04 | 0.006917 |
| SNCG | 10 | 8.7E+07 | 8.7E+07 | + | 1264 | synuclein gam | 0.48014 | 2.41224 | 12.583 | 7.60E-04 | 0.006981 |
| ADGRG2 | X | 1.9E+07 | 1.9E+07 | - | 10157 | adhesion G pro | -0.2723 | 4.31309 | 12.6538 | 7.61E-04 | 0.006983 |
| PFAS | 17 | 8247618 | 8270491 | + | 6471 | phosphoribosy | -0.099 | 7.92351 | 12.5785 | 7.61E-04 | 0.006985 |
| LINC0260 | 4 | 3758748 | 3763390 | - | 4525 | long intergenic | -0.5353 | 2.19723 | 12.5775 | 7.62E-04 | 0.006985 |
| IFT140 | 16 | 1510427 | 1612072 | - | 8130 | intraflagellar t | 0.22509 | 4.78287 | 12.5761 | 7.62E-04 | 0.006985 |
| KLK10 | 19 | 5.1E+07 | 5.1E+07 | - | 3798 | kallikrein relat | -0.6555 | 1.60965 | 12.5584 | 7.68E-04 | 0.007031 |
| NDUFAF3 | 3 | 4.9E+07 | 4.9E+07 | + | 2434 | NADH:ubiquin | 0.33239 | 3.56389 | 12.5583 | 7.68E-04 | 0.007031 |
| GGT1 | 22 | 2.5E+07 | 2.5E+07 | + | 5050 | gamma-glutam | -0.2639 | 4.19653 | 12.5341 | 7.77E-04 | 0.007102 |
| ERC1 | 12 | 990509 | 1495933 | + | 13484 | ELKS/RAB6-int | 0.15255 | 6.07249 | 12.532 | 7.77E-04 | 0.007104 |
| RBM38 | 20 | 5.7E+07 | 5.7E+07 | + | 2697 | RNA binding m | -0.1269 | 6.62065 | 12.5285 | 7.78E-04 | 0.007111 |
| MAN1B1 | 9 | 1.4E+08 | 1.4E+08 | + | 8269 | mannosidase a | 0.18354 | 5.37901 | 12.5162 | 7.83E-04 | 0.007145 |
| TFDP1 | 13 | 1.1E+08 | 1.1E+08 | + | 3744 | transcription fa | 0.11135 | 7.20717 | 12.5147 | 7.83E-04 | 0.007145 |
| ABTB2 | 11 | 3.4E+07 | 3.4E+07 | - | 4976 | ankyrin repeat | 0.24167 | 4.5156 | 12.4874 | 7.93E-04 | 0.007228 |
| POLR2G | 11 | 6.3E+07 | 6.3E+07 | + | 1902 | RNA polymera | 0.14627 | 6.12542 | 12.4848 | 7.94E-04 | 0.007231 |
| OSBPL1A | 18 | 2.4E+07 | 2.4E+07 | - | 8029 | oxysterol bindi | -0.1384 | 6.28998 | 12.4825 | 7.94E-04 | 0.007234 |
| EIF3K | 19 | 3.9E+07 | 3.9E+07 | + | 2798 | eukaryotic tran | -0.1077 | 7.40526 | 12.4766 | 7.97E-04 | 0.007248 |
| C6orf106 | 6 | 3.5E+07 | 3.5E+07 | - | 4551 | chromosome 6 | -0.1065 | 7.57726 | 12.4733 | 7.98E-04 | 0.007255 |
| NR6A1 | 9 | 1.2E+08 | 1.2E+08 | - | 7125 | nuclear recept | 0.11819 | 8.33804 | 13.9669 | 8.01E-04 | 0.007279 |
| MT-ND1 | MT | 3307 | 4262 | + | 956 | mitochondriall | -0.0738 | 11.5128 | 12.4594 | 8.03E-04 | 0.007286 |
| HTT | 4 | 3041422 | 3243960 | + | 19796 | huntingtin [Sou | 0.10155 | 7.82884 | 12.441 | 8.09E-04 | 0.00734 |
| ECSIT | 19 | 1.2E+07 | 1.2E+07 | - | 2355 | ECSIT signalling | -0.1644 | 5.71549 | 12.4338 | 8.12E-04 | 0.007359 |
| KCNQ2 | SCHR20_1 | 6.3E+07 | 6.3E+07 | - | 12775 | potassium volt | 0.1389 | 6.28218 | 12.43 | 8.13E-04 | 0.007367 |
| OPTN | 10 | 1.3E+07 | 1.3E+07 | + | 5138 | optineurin [Sou | -0.2641 | 4.20543 | 12.428 | 8.14E-04 | 0.007367 |
| ASPH | 8 | 6.2E+07 | 6.2E+07 | - | 20047 | aspartate beta | 0.15138 | 6.20408 | 12.5416 | 8.14E-04 | 0.007367 |
| DYNC2H1 | 11 | 1E+08 | 1E+08 | + | 15012 | dynein cytopla | 0.17334 | 5.70421 | 12.4445 | 8.15E-04 | 0.007367 |
| CD46 | 1 | 2.1E+08 | 2.1E+08 | + | 8708 | CD46 molecule | 0.10693 | 7.55254 | 12.4211 | 8.16E-04 | 0.007378 |

| | | | | | | | | | | | |
|---|---|---|---|---|---|---|---|---|---|---|---|
| CMPK1 | 1 | 4.7E+07 | 4.7E+07 | + | 3844 | cytidine/uridin | 0.16292 | 5.76231 | 12.4108 | 8.20E-04 | 0.007405 |
| CST3 | 20 | 2.4E+07 | 2.4E+07 | - | 3615 | cystatin C [Sou | -0.2291 | 4.69755 | 12.4099 | 8.20E-04 | 0.007405 |
| PLCD1 | 3 | 3.8E+07 | 3.8E+07 | - | 4314 | phospholipase | -0.2285 | 4.8597 | 12.4807 | 8.23E-04 | 0.007423 |
| PTGES2 | 9 | 1.3E+08 | 1.3E+08 | - | 2655 | prostaglandin | 0.13374 | 6.47629 | 12.4016 | 8.23E-04 | 0.007423 |
| N4BP3 | 5 | 1.8E+08 | 1.8E+08 | + | 5985 | NEDD4 binding | 0.1153 | 7.05675 | 12.3956 | 8.26E-04 | 0.007438 |
| SMARCD3 | 7 | 1.5E+08 | 1.5E+08 | - | 5198 | SWI/SNF relate | 0.58396 | 1.93784 | 12.3801 | 8.31E-04 | 0.007482 |
| UTP15 | 5 | 7.4E+07 | 7.4E+07 | + | 6324 | UTP15, small s | -0.2223 | 4.77104 | 12.3792 | 8.32E-04 | 0.007482 |
| SMAP1 | 6 | 7.1E+07 | 7.1E+07 | + | 4161 | small ArfGAP 1 | 0.25141 | 4.3299 | 12.3781 | 8.32E-04 | 0.007482 |
| INSIG1 | 7 | 1.6E+08 | 1.6E+08 | + | 3650 | insulin induced | -0.1378 | 6.44264 | 12.3629 | 8.38E-04 | 0.007527 |
| DGKQ | 4 | 958887 | 986895 | - | 4983 | diacylglycerol | 0.2552 | 4.26655 | 12.3619 | 8.38E-04 | 0.007527 |
| BRINP1 | 9 | 1.2E+08 | 1.2E+08 | - | 3874 | BMP/retinoic a | -0.2025 | 5.02445 | 12.36 | 8.39E-04 | 0.007528 |
| OBSCN | 1 | 2.3E+08 | 2.3E+08 | + | 32290 | obscurin, cytos | 0.19401 | 5.15768 | 12.3539 | 8.41E-04 | 0.007544 |
| WSCD1 | 17 | 6057807 | 6124427 | + | 7136 | WSC domain c | 0.16276 | 5.76288 | 12.3384 | 8.47E-04 | 0.007591 |
| PRKCG | 19 | 5.4E+07 | 5.4E+07 | + | 3397 | protein kinase | -0.3163 | 3.75515 | 12.3346 | 8.48E-04 | 0.0076 |
| CDK18 | 1 | 2.1E+08 | 2.1E+08 | + | 6831 | cyclin depende | -0.1694 | 5.85137 | 12.4579 | 8.54E-04 | 0.007642 |
| C11orf74 | 11 | 3.7E+07 | 3.7E+07 | + | 1474 | chromosome 1 | -0.359 | 3.25846 | 12.3117 | 8.57E-04 | 0.007667 |
| IQCA1 | 2 | 2.4E+08 | 2.4E+08 | - | 5968 | IQ motif conta | -0.3432 | 3.43595 | 12.3056 | 8.59E-04 | 0.007684 |
| MARCH6 | 5 | 1E+07 | 1E+07 | + | 11344 | membrane ass | 0.11969 | 6.85804 | 12.3024 | 8.61E-04 | 0.007689 |
| SDAD1 | 4 | 7.6E+07 | 7.6E+07 | - | 3491 | SDA1 domain c | -0.1399 | 6.26659 | 12.3013 | 8.61E-04 | 0.007689 |
| KANTR | X | 5.3E+07 | 5.3E+07 | + | 9927 | KDM5C adjace | -0.3757 | 3.14788 | 12.2971 | 8.63E-04 | 0.007698 |
| HESX1 | 3 | 5.7E+07 | 5.7E+07 | - | 1493 | HESX homeob | -0.4286 | 2.87918 | 12.3171 | 8.63E-04 | 0.007698 |
| LRRC61 | 7 | 1.5E+08 | 1.5E+08 | + | 1968 | leucine rich re | -0.307 | 3.99408 | 12.437 | 8.64E-04 | 0.007698 |
| PHYKPL | 5 | 1.8E+08 | 1.8E+08 | - | 8439 | 5-phosphohyd | 0.33135 | 3.49569 | 12.2921 | 8.64E-04 | 0.007701 |
| ATP13A2 | 1 | 1.7E+07 | 1.7E+07 | - | 5253 | ATPase cation | -0.1161 | 7.00698 | 12.2904 | 8.65E-04 | 0.007702 |
| CEP55 | 10 | 9.3E+07 | 9.4E+07 | + | 3271 | centrosomal p | -0.2151 | 4.85787 | 12.2846 | 8.67E-04 | 0.007712 |
| BBS4 | 15 | 7.3E+07 | 7.3E+07 | + | 4148 | Bardet-Biedl sy | 0.20439 | 4.95664 | 12.2468 | 8.82E-04 | 0.007838 |
| CUEDC2 | 10 | 1E+08 | 1E+08 | - | 1335 | CUE domain co | 0.21815 | 4.80917 | 12.2409 | 8.84E-04 | 0.007852 |
| STARD13 | 13 | 3.3E+07 | 3.3E+07 | - | 8386 | StAR related li | 0.45165 | 2.51601 | 12.24 | 8.85E-04 | 0.007852 |
| SLC29A1 | 6 | 4.4E+07 | 4.4E+07 | + | 4410 | solute carrier f | -0.1017 | 7.76934 | 12.2373 | 8.86E-04 | 0.007857 |
| OSMR | 5 | 3.9E+07 | 3.9E+07 | + | 7174 | oncostatin M r | 0.53067 | 2.11385 | 12.2321 | 8.88E-04 | 0.00787 |
| CIB1 | 15 | 9E+07 | 9E+07 | - | 1472 | calcium and in | -0.2635 | 4.28165 | 12.1976 | 9.02E-04 | 0.007987 |
| GDPD2 | X | 7E+07 | 7E+07 | + | 2638 | glycerophosph | 0.32536 | 3.87772 | 12.6114 | 9.03E-04 | 0.007992 |
| GON4L | 1 | 1.6E+08 | 1.6E+08 | - | 11967 | gon-4 like [Sou | 0.1666 | 5.63954 | 12.1806 | 9.08E-04 | 0.008038 |
| RYR3 | 15 | 3.3E+07 | 3.4E+07 | + | 20370 | ryanodine rece | 0.48285 | 2.34713 | 12.1774 | 9.10E-04 | 0.008042 |
| TACC1 | 8 | 3.9E+07 | 3.9E+07 | + | 12577 | transforming a | -0.2202 | 4.73132 | 12.1766 | 9.10E-04 | 0.008042 |
| AC007325 | KI270734.1 | 138082 | 161852 | - | 2405 | proline dehydr | -0.6904 | 1.43757 | 12.174 | 9.11E-04 | 0.008047 |
| RNF125 | 18 | 3.2E+07 | 3.2E+07 | + | 6061 | ring finger pro | -0.2283 | 4.62537 | 12.1698 | 9.13E-04 | 0.008057 |
| FBRSL1 | 12 | 1.3E+08 | 1.3E+08 | + | 9378 | fibrosin like 1 [ | 0.12025 | 6.94996 | 12.1667 | 9.14E-04 | 0.008063 |
| ANKRD40 | 17 | 5.1E+07 | 5.1E+07 | - | 4831 | ankyrin repeat | -0.1323 | 6.44813 | 12.1613 | 9.16E-04 | 0.008077 |
| CBL | 11 | 1.2E+08 | 1.2E+08 | + | 11740 | Cbl proto-onco | -0.1112 | 7.36706 | 12.1558 | 9.19E-04 | 0.008089 |
| DIAPH2 | X | 9.7E+07 | 9.8E+07 | + | 9550 | diaphanous re | -0.1536 | 5.93581 | 12.1552 | 9.19E-04 | 0.008089 |
| CAMK4 | 5 | 1.1E+08 | 1.1E+08 | + | 13614 | calcium/calmo | 0.3451 | 3.31344 | 12.15 | 9.21E-04 | 0.008103 |
| CBR3 | 21 | 3.6E+07 | 3.6E+07 | + | 998 | carbonyl reduc | -0.709 | 1.37888 | 12.1436 | 9.24E-04 | 0.008121 |

| | | | | | | | | | | | |
|---|---|---|---|---|---|---|---|---|---|---|---|
| ABHD2 | 15 | 8.9E+07 | 8.9E+07 | + | 10068 | abhydrolase do | 0.16435 | 5.66737 | 12.1408 | 9.25E-04 | 0.008127 |
| RTRAF | 14 | 5.2E+07 | 5.2E+07 | + | 8287 | RNA transcript | -0.1073 | 7.304 | 12.1359 | 9.27E-04 | 0.008139 |
| HNRNPLL | 2 | 3.9E+07 | 3.9E+07 | - | 5932 | heterogeneous | 0.13317 | 6.33815 | 12.1212 | 9.33E-04 | 0.008188 |
| TIMELESS | 12 | 5.6E+07 | 5.6E+07 | - | 5851 | timeless circad | 0.11365 | 7.14212 | 12.112 | 9.37E-04 | 0.008216 |
| THUMPD1 | 16 | 2.1E+07 | 2.1E+07 | - | 6501 | THUMP domai | -0.145 | 6.2487 | 12.1237 | 9.41E-04 | 0.008246 |
| SELENOH | 11 | 5.8E+07 | 5.8E+07 | + | 2305 | selenoprotein | 0.13459 | 6.35418 | 12.0869 | 9.47E-04 | 0.008299 |
| CASP3 | 4 | 1.8E+08 | 1.8E+08 | - | 2899 | caspase 3 [Sou | 0.1414 | 6.6047 | 12.5239 | 9.48E-04 | 0.0083 |
| E2F8 | 11 | 1.9E+07 | 1.9E+07 | - | 4134 | E2F transcripti | -0.3104 | 3.67875 | 12.0613 | 9.58E-04 | 0.008384 |
| TRIM24 | 7 | 1.4E+08 | 1.4E+08 | + | 9301 | tripartite motif | -0.1042 | 7.70034 | 12.058 | 9.59E-04 | 0.008391 |
| RPAP3 | 12 | 4.8E+07 | 4.8E+07 | - | 5350 | RNA polymera | -0.1681 | 5.63387 | 12.053 | 9.62E-04 | 0.008405 |
| TJP3 | 19 | 3708362 | 3750813 | + | 3964 | tight junction p | -0.2309 | 4.63628 | 12.0474 | 9.64E-04 | 0.008421 |
| UCP2 | 11 | 7.4E+07 | 7.4E+07 | - | 2768 | uncoupling pro | 0.23711 | 4.55142 | 12.0426 | 9.66E-04 | 0.008434 |
| RNPEPL1 | 2 | 2.4E+08 | 2.4E+08 | + | 7970 | arginyl aminop | 0.24252 | 4.48206 | 12.0411 | 9.67E-04 | 0.008434 |
| TBC1D8 | 2 | 1E+08 | 1E+08 | - | 6030 | TBC1 domain f | 0.2151 | 4.84519 | 12.0277 | 9.73E-04 | 0.00847 |
| BX322639 | 10 | 4.2E+07 | 4.2E+07 | - | 7208 | zinc finger prot | -0.1583 | 5.92628 | 12.0276 | 9.73E-04 | 0.00847 |
| CAPN1 | 11 | 6.5E+07 | 6.5E+07 | + | 7553 | calpain 1 [Sour | -0.0998 | 7.82287 | 12.0101 | 9.80E-04 | 0.008531 |
| PLPPR3 | SCHR19_5 | 821485 | 821977 | - | 2741 | phospholipid p | 0.1566 | 6.05186 | 12.1269 | 9.86E-04 | 0.008579 |
| MRPL42 | 12 | 9.3E+07 | 9.4E+07 | + | 16394 | mitochondrial | -0.1206 | 6.78959 | 11.9886 | 9.90E-04 | 0.008603 |
| POR | 7 | 7.6E+07 | 7.6E+07 | + | 6042 | cytochrome p4 | -0.1114 | 7.15844 | 11.981 | 9.93E-04 | 0.008627 |
| HGS | 17 | 8.2E+07 | 8.2E+07 | + | 8205 | hepatocyte gro | 0.13554 | 6.26962 | 11.9788 | 9.94E-04 | 0.00863 |
| RANGAP1 | 22 | 4.1E+07 | 4.1E+07 | - | 5348 | Ran GTPase ac | -0.0979 | 8.39951 | 12.1078 | 9.97E-04 | 0.008649 |
| SPAG5 | 17 | 2.9E+07 | 2.9E+07 | - | 5218 | sperm associat | 0.10609 | 7.32582 | 11.9453 | 0.001009 | 0.00875 |
| PTPRK | 6 | 1.3E+08 | 1.3E+08 | - | 9000 | protein tyrosin | 0.1358 | 6.37901 | 11.9418 | 0.0010106 | 0.008759 |
| CLDN7 | 17 | 7259903 | 7263983 | - | 2492 | claudin 7 [Sour | -0.1195 | 6.83328 | 11.9379 | 0.0010124 | 0.008765 |
| LINC0113 | 1 | 2.4E+08 | 2.4E+08 | - | 5959 | long intergenic | 0.84358 | 1.0738 | 12.0549 | 0.0010126 | 0.008765 |
| PSD | 10 | 1E+08 | 1E+08 | - | 5979 | pleckstrin and | -0.3858 | 2.98673 | 11.9357 | 0.0010134 | 0.008767 |
| SCAF1 | 19 | 5E+07 | 5E+07 | + | 4464 | SR-related CTD | 0.11091 | 7.11759 | 11.9337 | 0.0010143 | 0.008769 |
| LIG1 | 19 | 4.8E+07 | 4.8E+07 | - | 6576 | DNA ligase 1 [S | 0.12273 | 6.72141 | 11.9214 | 0.0010199 | 0.008807 |
| PKNOX1 | 21 | 4.3E+07 | 4.3E+07 | + | 7621 | PBX/knotted 1 | -0.1821 | 5.32465 | 11.9214 | 0.0010199 | 0.008807 |
| ITPRIP | 10 | 1E+08 | 1E+08 | - | 7296 | inositol 1,4,5-t | 0.20853 | 4.84645 | 11.9113 | 0.0010245 | 0.008836 |
| ORMDL3 | 17 | 4E+07 | 4E+07 | - | 3752 | ORMDL sphing | -0.1561 | 5.83986 | 11.9015 | 0.001029 | 0.008868 |
| SERP1 | 3 | 1.5E+08 | 1.5E+08 | - | 5743 | stress associat | 0.10581 | 7.41541 | 11.9007 | 0.0010294 | 0.008868 |
| DGAT1 | 8 | 1.4E+08 | 1.4E+08 | - | 4131 | diacylglycerol O | 0.2266 | 4.65565 | 11.8856 | 0.0010364 | 0.008922 |
| GPM6A | 4 | 1.8E+08 | 1.8E+08 | - | 7040 | glycoprotein M | 0.57405 | 1.86091 | 11.8746 | 0.0010415 | 0.008961 |
| LZTS1 | 8 | 2E+07 | 2E+07 | - | 5706 | leucine zipper | -0.3701 | 3.23213 | 11.8712 | 0.0010431 | 0.00897 |
| GPR107 | 9 | 1.3E+08 | 1.3E+08 | + | 9139 | G protein-coup | 0.1198 | 6.74579 | 11.8661 | 0.0010454 | 0.008984 |
| CTNNBIP1 | 1 | 9848276 | 9910336 | - | 3339 | catenin beta in | 0.14391 | 6.12266 | 11.8643 | 0.0010463 | 0.008986 |
| RPL22L1 | 3 | 1.7E+08 | 1.7E+08 | - | 2051 | ribosomal prot | -0.152 | 5.8977 | 11.8617 | 0.0010475 | 0.00899 |
| ACOT9 | X | 2.4E+07 | 2.4E+07 | - | 6244 | acyl-CoA thioe | -0.1664 | 5.62196 | 11.8608 | 0.001048 | 0.00899 |
| LINC0059 | 8 | 9900064 | 9905366 | - | 4136 | long intergenic | -0.2173 | 5.14493 | 12.2549 | 0.0010565 | 0.009058 |
| NME2 | 17 | 5.1E+07 | 5.1E+07 | + | 2632 | NME/NM23 nu | 0.1669 | 5.71802 | 11.8408 | 0.0010574 | 0.00906 |
| SLTM | 15 | 5.9E+07 | 5.9E+07 | - | 8178 | SAFB like trans | -0.1025 | 7.80157 | 11.8364 | 0.0010595 | 0.009069 |
| GMFB | 14 | 5.4E+07 | 5.4E+07 | - | 7796 | glia maturation | -0.124 | 6.98555 | 12.0179 | 0.0010597 | 0.009069 |

| | | | | | | | | | | | | |
|---|---|---|---|---|---|---|---|---|---|---|---|---|
| KCTD17 | 22 | 3.7E+07 | 3.7E+07 | + | | 2786 | potassium cha | 0.21002 | 4.82139 | 11.8329 | 0.0010611 | 0.009076 |
| PFKFB2 | 1 | 2.1E+08 | 2.1E+08 | + | | 10176 | 6-phosphofruc | -0.1892 | 5.1799 | 11.8232 | 0.0010658 | 0.00911 |
| AKIRIN1 | 1 | 3.9E+07 | 3.9E+07 | + | | 2688 | akirin 1 [Sourc | -0.0914 | 8.53098 | 11.8151 | 0.0010697 | 0.009132 |
| AC093297 | 5 | 4.5E+07 | 4.5E+07 | + | | 2517 | novel transcrip | -0.4659 | 2.48243 | 11.8139 | 0.0010702 | 0.009132 |
| PRCC | 1 | 1.6E+08 | 1.6E+08 | + | | 4071 | proline rich mi | -0.1177 | 6.80632 | 11.8139 | 0.0010702 | 0.009132 |
| EME2 | 16 | 1773207 | 1781708 | + | | 6909 | essential meio | 0.35714 | 3.29543 | 11.8104 | 0.0010719 | 0.009141 |
| PTK2 | 8 | 1.4E+08 | 1.4E+08 | - | | 13742 | protein tyrosin | 0.10072 | 7.67175 | 11.8058 | 0.0010741 | 0.009148 |
| PPP1R9B | 17 | 5E+07 | 5E+07 | - | | 4343 | protein phosph | 0.20591 | 4.89489 | 11.8045 | 0.0010747 | 0.009149 |
| ATXN7L1 | 7 | 1.1E+08 | 1.1E+08 | - | | 8295 | ataxin 7 like 1 | 0.21479 | 4.87857 | 11.7974 | 0.0010782 | 0.009172 |
| GRAMD2I | 5 | 1.3E+08 | 1.3E+08 | + | | 5051 | GRAM domain | 0.30472 | 3.69223 | 11.7683 | 0.0010923 | 0.009287 |
| ST6GALNA | 1 | 7.7E+07 | 7.7E+07 | + | | 6059 | ST6 N-acetylga | 0.40145 | 2.8948 | 11.7671 | 0.0010929 | 0.009287 |
| TNRC6B | 22 | 4E+07 | 4E+07 | + | | 19998 | trinucleotide r | -0.1334 | 6.34036 | 11.7572 | 0.0010978 | 0.009323 |
| AC009831 | 18 | 3.2E+07 | 3.2E+07 | + | | 1582 | novel transcrip | -0.501 | 2.28791 | 11.7267 | 0.0011129 | 0.009444 |
| RASA3 | 13 | 1.1E+08 | 1.1E+08 | - | | 4775 | RAS p21 prote | 0.15542 | 5.82787 | 11.7258 | 0.0011134 | 0.009444 |
| GATD3A | 21 | 4.4E+07 | 4.4E+07 | + | | 7947 | glutamine ami | -0.385 | 2.98787 | 11.7243 | 0.0011141 | 0.009445 |
| DNAJA1 | 9 | 3.3E+07 | 3.3E+07 | + | | 2668 | DnaJ heat shoc | -0.0972 | 7.93806 | 11.7139 | 0.0011194 | 0.009483 |
| VENTX | 10 | 1.3E+08 | 1.3E+08 | + | | 2459 | VENT homeob | -0.4304 | 2.73224 | 11.7048 | 0.0011239 | 0.009516 |
| HHLA1 | 8 | 1.3E+08 | 1.3E+08 | - | | 6143 | HERV-H LTR-as | -0.5524 | 1.99332 | 11.7002 | 0.0011263 | 0.009531 |
| FAM210A | 18 | 1.4E+07 | 1.4E+07 | - | | 4681 | family with seq | -0.2674 | 4.25651 | 11.7613 | 0.0011276 | 0.009537 |
| THRA | 17 | 4E+07 | 4E+07 | + | | 6959 | thyroid hormo | -0.2175 | 4.71475 | 11.6921 | 0.0011303 | 0.009554 |
| GLI4 | 8 | 1.4E+08 | 1.4E+08 | + | | 3057 | GLI family zinc | 0.40024 | 2.81908 | 11.6832 | 0.0011349 | 0.009585 |
| LRRC59 | 17 | 5E+07 | 5E+07 | - | | 4540 | leucine rich re | -0.1059 | 7.39737 | 11.6822 | 0.0011354 | 0.009585 |
| MFSD12 | 19 | 3538261 | 3574290 | - | | 5290 | major facilitato | 0.17339 | 5.47328 | 11.6704 | 0.0011414 | 0.009631 |
| FBXW8 | 12 | 1.2E+08 | 1.2E+08 | + | | 5209 | F-box and WD | 0.21209 | 4.92171 | 11.6509 | 0.0011515 | 0.009702 |
| CD63 | 12 | 5.6E+07 | 5.6E+07 | - | | 2633 | CD63 molecule | -0.0918 | 8.36535 | 11.6501 | 0.0011519 | 0.009702 |
| CAMKV | 3 | 5E+07 | 5E+07 | - | | 3613 | CaM kinase lik | -0.1434 | 6.16469 | 11.65 | 0.0011519 | 0.009702 |
| IGFBP3 | 7 | 4.6E+07 | 4.6E+07 | - | | 4571 | insulin like gro | -0.3552 | 3.21693 | 11.6433 | 0.0011554 | 0.009726 |
| TUBA1C | 12 | 4.9E+07 | 4.9E+07 | + | | 4631 | tubulin alpha 1 | -0.0929 | 8.12526 | 11.6416 | 0.0011563 | 0.009727 |
| LIMCH1 | 4 | 4.1E+07 | 4.2E+07 | + | | 10862 | LIM and calpo | -0.4067 | 2.9124 | 11.6404 | 0.001157 | 0.009727 |
| CDC42BPI | 14 | 1E+08 | 1E+08 | - | | 8095 | CDC42 binding | 0.10158 | 7.69852 | 11.6391 | 0.0011576 | 0.009727 |
| COX7C | 5 | 8.7E+07 | 8.7E+07 | + | | 1969 | cytochrome c | -0.0984 | 8.09523 | 11.6647 | 0.0011621 | 0.00976 |
| NUAK1 | 12 | 1.1E+08 | 1.1E+08 | - | | 7008 | NUAK family ki | -0.1643 | 5.70846 | 11.6271 | 0.0011639 | 0.009769 |
| ANAPC4 | 4 | 2.5E+07 | 2.5E+07 | + | | 7494 | anaphase pror | -0.2096 | 4.83845 | 11.6241 | 0.0011655 | 0.009772 |
| LARP1B | 4 | 1.3E+08 | 1.3E+08 | + | | 9480 | La ribonucleop | -0.1683 | 5.71217 | 11.6386 | 0.0011758 | 0.00985 |
| KIDINS22( | 2 | 8721081 | 8837630 | - | | 15362 | kinase D intera | 0.12382 | 6.54153 | 11.6023 | 0.001177 | 0.00985 |
| RPS3 | 11 | 7.5E+07 | 7.5E+07 | + | | 5859 | ribosomal prot | -0.0728 | 10.6157 | 11.602 | 0.0011771 | 0.00985 |
| ZNF334 | 20 | 4.6E+07 | 4.7E+07 | - | | 6865 | zinc finger pro | -0.2811 | 3.91479 | 11.6008 | 0.0011777 | 0.00985 |
| CNIH4 | 1 | 2.2E+08 | 2.2E+08 | + | | 5090 | cornichon fam | -0.1562 | 5.79558 | 11.5981 | 0.0011792 | 0.009857 |
| SLC44A1 | 9 | 1.1E+08 | 1.1E+08 | + | | 12172 | solute carrier f | -0.1273 | 7.09424 | 12.1936 | 0.0011864 | 0.009912 |
| AKR1B1 | 7 | 1.3E+08 | 1.3E+08 | - | | 3654 | aldo-keto redu | 0.10168 | 7.51736 | 11.58 | 0.0011888 | 0.009926 |
| ZNF563 | 19 | 1.2E+07 | 1.2E+07 | - | | 2847 | zinc finger pro | -0.653 | 1.5353 | 11.5718 | 0.0011932 | 0.009946 |
| RIC1 | 9 | 5629025 | 5776557 | + | | 8591 | RIC1 homolog, | 0.16018 | 5.75748 | 11.5717 | 0.0011933 | 0.009946 |
| OSCAR | 19LRC_LR | 5.4E+07 | 5.4E+07 | - | | 2084 | osteoclast ass | 0.58765 | 1.83192 | 11.5443 | 0.0012081 | 0.010064 |

| | | | | | | | | | | | | |
|---|---|---|---|---|---|---|---|---|---|---|---|---|
| CSPP1 | 8 | 6.7E+07 | 6.7E+07 | + | | 7096 | centrosome ar | -0.2399 | 4.36731 | 11.5423 | 0.0012092 | 0.010067 |
| JADE2 | 5 | 1.3E+08 | 1.3E+08 | + | | 9877 | jade family PH | -0.1317 | 6.38501 | 11.5389 | 0.0012111 | 0.010073 |
| HERC2P2 | SCHR15_3 | 2.2E+07 | 2.3E+07 | + | | 6965 | hect domain ar | 0.3663 | 3.18494 | 11.5384 | 0.0012113 | 0.010073 |
| ACAD8 | 11 | 1.3E+08 | 1.3E+08 | + | | 7174 | acyl-CoA dehy | 0.13844 | 6.15922 | 11.5296 | 0.0012161 | 0.010107 |
| BSDC1 | 1 | 3.2E+07 | 3.2E+07 | - | | 6271 | BSD domain co | 0.14319 | 6.03903 | 11.5256 | 0.0012183 | 0.010119 |
| NECAB3 | 20 | 3.4E+07 | 3.4E+07 | - | | 4342 | N-terminal EF- | 0.3665 | 3.58451 | 12.2808 | 0.001224 | 0.01016 |
| PAIP2B | 2 | 7.1E+07 | 7.1E+07 | - | | 6300 | poly(A) binding | -0.1581 | 5.7844 | 11.5135 | 0.001225 | 0.010163 |
| SEMA6B | 19 | 4542593 | 4559684 | - | | 4288 | semaphorin 6E | -0.1127 | 7.06687 | 11.5118 | 0.001226 | 0.010165 |
| SMARCC2 | 12 | 5.6E+07 | 5.6E+07 | - | | 9179 | SWI/SNF relate | 0.12883 | 6.49632 | 11.5101 | 0.0012269 | 0.010167 |
| NECTIN1 | 11 | 1.2E+08 | 1.2E+08 | - | | 7817 | nectin cell adh | -0.0952 | 8.01772 | 11.4865 | 0.00124 | 0.010264 |
| UQCRB | 8 | 9.6E+07 | 9.6E+07 | - | | 9715 | ubiquinol-cyto | -0.1164 | 6.89788 | 11.4827 | 0.0012422 | 0.010276 |
| RACGAP1 | 12 | 5E+07 | 5E+07 | - | | 4487 | Rac GTPase act | 0.13263 | 6.38045 | 11.4707 | 0.0012489 | 0.010325 |
| MYL5 | 4 | 673580 | 682033 | + | | 4305 | myosin light ch | 0.59743 | 1.6693 | 11.4606 | 0.0012546 | 0.010363 |
| NDRG2 | 14 | 2.1E+07 | 2.1E+07 | - | | 7550 | NDRG family m | -0.3339 | 4.35094 | 13.4604 | 0.0012549 | 0.010363 |
| ALPL | 1 | 2.2E+07 | 2.2E+07 | + | | 2980 | alkaline phosp | -0.0997 | 7.70973 | 11.443 | 0.0012646 | 0.010431 |
| CDX4 | X | 7.3E+07 | 7.3E+07 | + | | 1515 | caudal type ho | 0.83492 | 0.9102 | 11.435 | 0.0012692 | 0.010461 |
| RIN1 | 11 | 6.6E+07 | 6.6E+07 | - | | 5377 | Ras and Rab in | 0.4173 | 2.66957 | 11.4341 | 0.0012697 | 0.010461 |
| SELENOV | 19 | 4E+07 | 4E+07 | + | | 1731 | selenoprotein | -0.4336 | 2.64726 | 11.4207 | 0.0012774 | 0.010516 |
| TUBA4A | 2 | 2.2E+08 | 2.2E+08 | - | | 3966 | tubulin alpha 4 | 0.14607 | 5.95129 | 11.4201 | 0.0012778 | 0.010516 |
| JUND | 19 | 1.8E+07 | 1.8E+07 | - | | 1863 | JunD proto-on | 0.11628 | 6.87211 | 11.4141 | 0.0012812 | 0.010538 |
| RCAN1 | 21 | 3.5E+07 | 3.5E+07 | - | | 6964 | regulator of ca | -0.2729 | 3.99475 | 11.4125 | 0.0012821 | 0.010539 |
| PTGR1 | 9 | 1.1E+08 | 1.1E+08 | - | | 3770 | prostaglandin | 0.17397 | 5.40673 | 11.4027 | 0.0012878 | 0.01058 |
| CA2 | 8 | 8.5E+07 | 8.5E+07 | + | | 2007 | carbonic anhyd | -0.449 | 2.46602 | 11.401 | 0.0012888 | 0.010583 |
| SEMA5A | 5 | 9035033 | 9546075 | - | | 12308 | semaphorin 5A | 0.2404 | 4.33739 | 11.3962 | 0.0012916 | 0.010599 |
| U91319.1 | 16 | 1.3E+07 | 1.4E+07 | + | | 826 | novel transcrip | -1.0803 | 0.46624 | 11.5262 | 0.0012946 | 0.010618 |
| LRG1 | 19 | 4536402 | 4540036 | - | | 2702 | leucine rich alp | 0.66724 | 1.39959 | 11.3828 | 0.0012995 | 0.010652 |
| EIF3M | 11 | 3.3E+07 | 3.3E+07 | + | | 7430 | eukaryotic tran | -0.0995 | 7.64639 | 11.3797 | 0.0013013 | 0.010661 |
| SYNE2 | 14 | 6.4E+07 | 6.4E+07 | + | | 31374 | spectrin repeat | 0.10383 | 7.46583 | 11.3732 | 0.0013051 | 0.010686 |
| ADCY1 | 7 | 4.6E+07 | 4.6E+07 | + | | 13966 | adenylate cycla | -0.1668 | 5.60281 | 11.364 | 0.0013106 | 0.010718 |
| ACSM3 | 16 | 2.1E+07 | 2.1E+07 | + | | 7510 | acyl-CoA synth | -0.54 | 1.96031 | 11.3599 | 0.001313 | 0.010732 |
| GOT1 | 10 | 9.9E+07 | 9.9E+07 | - | | 2424 | glutamic-oxalo | -0.1376 | 6.18695 | 11.3579 | 0.0013142 | 0.010736 |
| RBM48 | 7 | 9.3E+07 | 9.3E+07 | + | | 6829 | RNA binding m | 0.3815 | 2.92293 | 11.3555 | 0.0013157 | 0.010741 |
| CTSZ | 20 | 5.9E+07 | 5.9E+07 | - | | 2045 | cathepsin Z [Sc | 0.2086 | 4.81267 | 11.3517 | 0.0013179 | 0.010752 |
| EEF1A2 | 20 | 6.3E+07 | 6.3E+07 | - | | 4771 | eukaryotic tran | 0.19323 | 5.17211 | 11.3492 | 0.0013194 | 0.010752 |
| SOWAHC | 2 | 1.1E+08 | 1.1E+08 | + | | 4627 | sosondowah a | 0.20503 | 4.93315 | 11.3484 | 0.0013198 | 0.010752 |
| LHFPL2 | 5 | 7.8E+07 | 7.9E+07 | - | | 7977 | LHFPL tetraspa | 0.14684 | 5.97067 | 11.3477 | 0.0013203 | 0.010752 |
| UBA52 | 19 | 1.9E+07 | 1.9E+07 | + | | 4306 | ubiquitin A-52 | -0.0915 | 8.33475 | 11.3462 | 0.0013212 | 0.010752 |
| TM7SF3 | 12 | 2.7E+07 | 2.7E+07 | - | | 5504 | transmembran | 0.13263 | 6.56638 | 11.48 | 0.0013214 | 0.010752 |
| CHM | X | 8.6E+07 | 8.6E+07 | - | | 8644 | CHM, Rab esco | 0.21045 | 4.851 | 11.3439 | 0.0013225 | 0.010755 |
| RNASEH2 | 11 | 6.6E+07 | 6.6E+07 | - | | 5101 | ribonuclease H | 0.21699 | 4.70021 | 11.341 | 0.0013243 | 0.010763 |
| EXT2 | 11 | 4.4E+07 | 4.4E+07 | + | | 5286 | exostosin glyco | -0.1246 | 6.46325 | 11.3342 | 0.0013284 | 0.010784 |
| THRB | 3 | 2.4E+07 | 2.4E+07 | - | | 10968 | thyroid hormo | -0.6091 | 1.67232 | 11.3327 | 0.0013293 | 0.010785 |
| C11orf96 | 11 | 4.4E+07 | 4.4E+07 | + | | 1918 | chromosome 1 | -0.9762 | 0.47586 | 11.3301 | 0.0013308 | 0.010792 |

| | | | | | | | | | | | |
|---|---|---|---|---|---|---|---|---|---|---|---|
| DCBLD1 | 6 | 1.2E+08 | 1.2E+08 | + | | 6819 | discoidin, CUB | 0.2701 | 3.98774 | 11.3258 | 0.0013335 | 0.010807 |
| COCH | 14 | 3.1E+07 | 3.1E+07 | + | | 4031 | cochlin [Source | 0.40753 | 2.72781 | 11.3234 | 0.0013349 | 0.010812 |
| BCAP31 | X | 1.5E+08 | 1.5E+08 | - | | 3533 | B cell receptor | -0.1041 | 7.33754 | 11.3164 | 0.0013391 | 0.010835 |
| YY1AP1 | 1 | 1.6E+08 | 1.6E+08 | - | | 5001 | YY1 associated | -0.1887 | 5.18084 | 11.3163 | 0.0013392 | 0.010835 |
| YME1L1 | 10 | 2.7E+07 | 2.7E+07 | - | | 7335 | YME1 like 1 AT | -0.1 | 7.55248 | 11.3085 | 0.0013439 | 0.010867 |
| SHARPIN | 8 | 1.4E+08 | 1.4E+08 | - | | 2783 | SHANK associa | 0.18893 | 5.08054 | 11.2941 | 0.0013527 | 0.010926 |
| MINK1 | 17 | 4833340 | 4898061 | + | | 7865 | misshapen like | 0.11871 | 6.69475 | 11.2933 | 0.0013532 | 0.010926 |
| NUP153 | 6 | 1.8E+07 | 1.8E+07 | - | | 6030 | nucleoporin 15 | 0.09949 | 7.59734 | 11.2929 | 0.0013535 | 0.010926 |
| SMAD3 | 15 | 6.7E+07 | 6.7E+07 | + | | 9330 | SMAD family m | 0.1967 | 5.0815 | 11.2904 | 0.001355 | 0.010931 |
| STK17B | 2 | 2E+08 | 2E+08 | - | | 6566 | serine/threoni | 0.30334 | 3.68638 | 11.2893 | 0.0013556 | 0.010931 |
| MCM4 | 8 | 4.8E+07 | 4.8E+07 | + | | 6973 | minichromoso | 0.08213 | 9.01094 | 11.2833 | 0.0013594 | 0.010955 |
| NEPRO | 3 | 1.1E+08 | 1.1E+08 | - | | 6490 | nucleolus and | -0.1523 | 5.86938 | 11.2771 | 0.0013632 | 0.010979 |
| ACTR5 | 20 | 3.9E+07 | 3.9E+07 | + | | 2547 | ARP5 actin rela | -0.2117 | 4.75723 | 11.2759 | 0.0013639 | 0.010979 |
| LEO1 | 15 | 5.2E+07 | 5.2E+07 | - | | 2651 | LEO1 homolog | -0.1826 | 5.31072 | 11.2721 | 0.0013662 | 0.010992 |
| CALM1 | 14 | 9E+07 | 9E+07 | + | | 7666 | calmodulin 1 [S | -0.0885 | 8.35791 | 11.2691 | 0.0013681 | 0.011001 |
| ELAVL2 | 9 | 2.4E+07 | 2.4E+07 | - | | 5140 | ELAV like RNA | 0.23002 | 4.45932 | 11.2617 | 0.0013727 | 0.011028 |
| NMT2 | 10 | 1.5E+07 | 1.5E+07 | - | | 6470 | N-myristoyltra | -0.2493 | 4.21997 | 11.2612 | 0.001373 | 0.011028 |
| TRPT1 | 11 | 6.4E+07 | 6.4E+07 | - | | 2210 | tRNA phospho | 0.26524 | 4.0108 | 11.2505 | 0.0013797 | 0.011075 |
| ABHD11 | 7 | 7.4E+07 | 7.4E+07 | - | | 2289 | abhydrolase d | 0.33631 | 3.29793 | 11.2466 | 0.0013822 | 0.011089 |
| USP33 | 1 | 7.8E+07 | 7.8E+07 | - | | 6918 | ubiquitin speci | 0.13509 | 6.19403 | 11.2375 | 0.0013879 | 0.011129 |
| UBE2V2 | 8 | 4.8E+07 | 4.8E+07 | + | | 6591 | ubiquitin conju | -0.1252 | 6.4253 | 11.2359 | 0.0013889 | 0.011129 |
| ARL2 | 11 | 6.5E+07 | 6.5E+07 | + | | 2037 | ADP ribosylatio | 0.13014 | 6.32004 | 11.235 | 0.0013894 | 0.011129 |
| GNPDA1 | 5 | 1.4E+08 | 1.4E+08 | - | | 3959 | glucosamine-6 | -0.109 | 7.03744 | 11.2332 | 0.0013906 | 0.011132 |
| DDX18 | 2 | 1.2E+08 | 1.2E+08 | + | | 7808 | DEAD-box helic | -0.1055 | 7.5507 | 11.2741 | 0.0014019 | 0.011216 |
| CARHSP1 | 16 | 8852942 | 8869012 | - | | 5577 | calcium regula | -0.104 | 7.30733 | 11.2135 | 0.001403 | 0.011219 |
| AURKB | 17 | 8204733 | 8210600 | - | | 2241 | aurora kinase | -0.1161 | 6.74546 | 11.203 | 0.0014097 | 0.01126 |
| B4GALT2 | 1 | 4.4E+07 | 4.4E+07 | + | | 3772 | beta-1,4-galact | 0.12407 | 6.45649 | 11.1969 | 0.0014137 | 0.011281 |
| SPIRE2 | 16 | 9E+07 | 9E+07 | + | | 7999 | spire type actin | 0.36487 | 3.3027 | 11.3596 | 0.0014145 | 0.011281 |
| PSMC1 | 14 | 9E+07 | 9E+07 | + | | 6129 | proteasome 26 | -0.15 | 5.82364 | 11.1952 | 0.0014147 | 0.011281 |
| SYNPO | 5 | 1.5E+08 | 1.5E+08 | + | | 10243 | synaptopodin | 0.44972 | 2.41213 | 11.1927 | 0.0014163 | 0.011287 |
| ZNF271P | 18 | 3.5E+07 | 3.5E+07 | + | | 7450 | zinc finger prot | -0.2103 | 4.86279 | 11.1913 | 0.0014172 | 0.011288 |
| PIP4K2B | 17 | 3.9E+07 | 3.9E+07 | - | | 6135 | phosphatidylin | 0.10839 | 7.02694 | 11.182 | 0.0014233 | 0.01133 |
| LDB2 | 4 | 1.7E+07 | 1.7E+07 | - | | 5455 | LIM domain bi | -0.1787 | 5.32556 | 11.1779 | 0.0014259 | 0.011345 |
| LTA4H | 12 | 9.6E+07 | 9.6E+07 | - | | 7405 | leukotriene A4 | 0.10045 | 7.42995 | 11.1761 | 0.001427 | 0.011348 |
| SLC6A15 | 12 | 8.5E+07 | 8.5E+07 | - | | 8727 | solute carrier f | 0.22634 | 4.48878 | 11.1728 | 0.0014292 | 0.011359 |
| SYF2 | 1 | 2.5E+07 | 2.5E+07 | - | | 3773 | SYF2 pre-mRNA | -0.2066 | 5.00245 | 11.222 | 0.0014351 | 0.011399 |
| CELSR2 | 1 | 1.1E+08 | 1.1E+08 | + | | 12830 | cadherin EGF L | -0.1388 | 6.05959 | 11.149 | 0.0014447 | 0.011463 |
| DHDDS | 1 | 2.6E+07 | 2.6E+07 | + | | 5608 | dehydrodolich | -0.1797 | 5.36756 | 11.1489 | 0.0014448 | 0.011463 |
| TOMM34 | 20 | 4.5E+07 | 4.5E+07 | - | | 1973 | translocase of | -0.1485 | 5.91156 | 11.1466 | 0.0014463 | 0.011469 |
| UHRF1BP | 12 | 1E+08 | 1E+08 | - | | 6646 | UHRF1 binding | -0.18 | 5.34142 | 11.1437 | 0.0014482 | 0.011478 |
| SPEG | 2 | 2.2E+08 | 2.2E+08 | + | | 17066 | striated muscle | 0.16554 | 5.63895 | 11.1214 | 0.001463 | 0.011584 |
| SLC25A24 | 1 | 1.1E+08 | 1.1E+08 | - | | 6726 | solute carrier f | 0.12527 | 6.40913 | 11.121 | 0.0014633 | 0.011584 |
| UBA1 | X | 4.7E+07 | 4.7E+07 | + | | 6437 | ubiquitin like m | -0.0843 | 8.96773 | 11.1185 | 0.0014649 | 0.011591 |

| | | | | | | | | | | | |
|---|---|---|---|---|---|---|---|---|---|---|---|
| ARC | 8 | 1.4E+08 | 1.4E+08 | - | 2993 | activity regulat | 0.25145 | 4.15148 | 11.1068 | 0.0014727 | 0.011646 |
| GNB4 | 3 | 1.8E+08 | 1.8E+08 | - | 6922 | G protein subu | 0.16023 | 5.68686 | 11.1056 | 0.0014735 | 0.011646 |
| NASP | 1 | 4.6E+07 | 4.6E+07 | + | 8304 | nuclear autoar | -0.075 | 9.8094 | 11.1033 | 0.001475 | 0.011652 |
| NDUFS5 | 1 | 3.9E+07 | 3.9E+07 | + | 548 | NADH:ubiquin | -0.1001 | 7.42601 | 11.0964 | 0.0014797 | 0.011682 |
| CNR1 | 6 | 8.8E+07 | 8.8E+07 | - | 6357 | cannabinoid re | -0.594 | 1.77472 | 11.0852 | 0.0014873 | 0.011736 |
| NEFH | 22 | 2.9E+07 | 2.9E+07 | + | 3783 | neurofilament | -0.3686 | 3.13175 | 11.0769 | 0.0014929 | 0.011773 |
| KLHL13 | X | 1.2E+08 | 1.2E+08 | - | 6983 | kelch like famil | 0.31775 | 3.62844 | 11.1279 | 0.0014974 | 0.011802 |
| AKAP17A | X | 1591604 | 1602520 | + | 4070 | A-kinase ancho | 0.13002 | 6.31152 | 11.0631 | 0.0015023 | 0.011835 |
| CNTROB | 17 | 7932101 | 7949918 | + | 6234 | centrobin, cen | -0.1698 | 5.44702 | 11.0492 | 0.0015118 | 0.011903 |
| MAT2B | 5 | 1.6E+08 | 1.6E+08 | + | 5135 | methionine ad | -0.1374 | 6.20097 | 11.0448 | 0.0015148 | 0.011921 |
| SFXN4 | 10 | 1.2E+08 | 1.2E+08 | - | 1868 | sideroflexin 4 [ | 0.23891 | 4.32955 | 11.0357 | 0.0015211 | 0.011964 |
| SLC25A36 | 3 | 1.4E+08 | 1.4E+08 | + | 10305 | solute carrier f | 0.11343 | 6.77729 | 10.9829 | 0.0015581 | 0.012248 |
| TMEM98 | 17 | 3.3E+07 | 3.3E+07 | + | 5187 | transmembran | -0.1323 | 6.19418 | 10.9774 | 0.001562 | 0.012272 |
| STK25 | 2 | 2.4E+08 | 2.4E+08 | - | 7588 | serine/threoni | 0.10279 | 7.23956 | 10.9617 | 0.0015733 | 0.012354 |
| RALGDS | 9 | 1.3E+08 | 1.3E+08 | - | 7648 | ral guanine nu | 0.4269 | 2.55015 | 10.9587 | 0.0015754 | 0.012363 |
| TPM2 | 9 | 3.6E+07 | 3.6E+07 | - | 3248 | tropomyosin 2 | 0.10976 | 7.15295 | 10.9539 | 0.0015789 | 0.012384 |
| COX7B | X | 7.8E+07 | 7.8E+07 | + | 3027 | cytochrome c c | -0.1399 | 6.04024 | 10.9384 | 0.0015901 | 0.012451 |
| VWA8 | 13 | 4.2E+07 | 4.2E+07 | - | 7635 | von Willebrand | 0.1512 | 5.75122 | 10.9306 | 0.0015957 | 0.012489 |
| ATP6V1B2 | 8 | 2E+07 | 2E+07 | + | 7797 | ATPase H+ tran | -0.119 | 6.67457 | 10.9272 | 0.0015983 | 0.012497 |
| AC010503 | 19 | 6469465 | 6470152 | + | 688 | novel transcrip | -0.4931 | 2.18593 | 10.9268 | 0.0015986 | 0.012497 |
| SAP18 | 13 | 2.1E+07 | 2.1E+07 | + | 3396 | Sin3A associate | -0.1048 | 7.32843 | 10.9211 | 0.0016027 | 0.012523 |
| CABYR | 18 | 2.4E+07 | 2.4E+07 | + | 3518 | calcium bindin | -0.3144 | 3.50187 | 10.9102 | 0.0016107 | 0.012573 |
| SLC39A11 | 17 | 7.3E+07 | 7.3E+07 | - | 5583 | solute carrier f | -0.2867 | 3.83169 | 10.9099 | 0.0016109 | 0.012573 |
| FAAP100 | 17 | 8.2E+07 | 8.2E+07 | - | 5353 | FA core comple | 0.16629 | 5.50115 | 10.9055 | 0.0016141 | 0.012591 |
| RSRC2 | 12 | 1.2E+08 | 1.2E+08 | - | 8745 | arginine and se | -0.1205 | 6.55156 | 10.9028 | 0.0016162 | 0.012595 |
| PEX7 | 6 | 1.4E+08 | 1.4E+08 | + | 1832 | peroxisomal bi | 0.34794 | 3.18052 | 10.9025 | 0.0016164 | 0.012595 |
| DAPK1 | 9 | 8.7E+07 | 8.8E+07 | + | 8908 | death associat | 0.10161 | 7.33906 | 10.8965 | 0.0016208 | 0.012619 |
| NOP9 | 14 | 2.4E+07 | 2.4E+07 | + | 6045 | NOP9 nucleola | 0.12986 | 6.23328 | 10.896 | 0.0016211 | 0.012619 |
| PLEKHN1 | 1 | 966497 | 975865 | + | 2833 | pleckstrin hom | 0.81098 | 0.81707 | 10.8945 | 0.0016223 | 0.012621 |
| MAGT1 | X | 7.8E+07 | 7.8E+07 | - | 4862 | magnesium tra | -0.1295 | 6.2634 | 10.8933 | 0.0016232 | 0.012621 |
| SCAMP5 | 15 | 7.5E+07 | 7.5E+07 | + | 4862 | secretory carri | -0.1255 | 6.39506 | 10.8904 | 0.0016253 | 0.012631 |
| N6AMT1 | 21 | 2.9E+07 | 2.9E+07 | - | 4865 | N-6 adenine-sp | -0.2089 | 4.72024 | 10.8841 | 0.00163 | 0.01266 |
| ALMS1 | 2 | 7.3E+07 | 7.4E+07 | + | 14783 | ALMS1, centro | 0.12022 | 6.67567 | 10.8816 | 0.0016319 | 0.012667 |
| TAF1B | 2 | 9843443 | 9934416 | + | 3975 | TATA-box bind | -0.2217 | 4.58054 | 10.8805 | 0.0016327 | 0.012667 |
| ZNF813 | 19 | 5.3E+07 | 5.3E+07 | + | 6246 | zinc finger prot | -0.1649 | 5.52172 | 10.8751 | 0.0016367 | 0.012692 |
| ANXA11 | 10 | 8E+07 | 8E+07 | - | 8513 | annexin A11 [S | -0.132 | 6.46417 | 10.9561 | 0.0016431 | 0.012735 |
| RYBP | HG126_P | 7.2E+07 | 7.2E+07 | - | 7215 | RING1 and YY1 | -0.1185 | 6.59376 | 10.8521 | 0.001654 | 0.012812 |
| BMI1 | 10 | 2.2E+07 | 2.2E+07 | + | 4395 | BMI1 proto-on | 0.43311 | 2.47443 | 10.8407 | 0.0016626 | 0.012872 |
| INPP5F | 10 | 1.2E+08 | 1.2E+08 | + | 7610 | inositol polyph | 0.09737 | 7.57158 | 10.8346 | 0.0016673 | 0.012883 |
| CDCA7L | 7 | 2.2E+07 | 2.2E+07 | - | 3894 | cell division cy | 0.11462 | 6.71762 | 10.834 | 0.0016677 | 0.012883 |
| MAN1C1 | 1 | 2.6E+07 | 2.6E+07 | + | 9026 | mannosidase a | -0.2675 | 4.03632 | 10.8335 | 0.0016681 | 0.012883 |
| CCDC18-A | 1 | 9.3E+07 | 9.3E+07 | - | 4786 | CCDC18 antise | 0.56349 | 1.87898 | 10.8332 | 0.0016683 | 0.012883 |
| CENPB | 20 | 3783851 | 3786740 | - | 2890 | centromere pr | 0.14022 | 5.99707 | 10.8321 | 0.0016691 | 0.012883 |

| | | | | | | | | | | | |
|---|---|---|---|---|---|---|---|---|---|---|---|
| NDUFS6 | 5 | 1801400 | 1816605 | + | 1451 | NADH:ubiquin | -0.1122 | 6.85586 | 10.8318 | 0.0016694 | 0.012883 |
| DBNL | 7 | 4.4E+07 | 4.4E+07 | + | 12587 | drebrin like [Sc | 0.17315 | 5.43512 | 10.8248 | 0.0016748 | 0.012917 |
| AC011498 | 19 | 4448810 | 4450836 | + | 2027 | TEC | 0.85 | 0.74895 | 10.8232 | 0.001676 | 0.01292 |
| ARMC4 | 10 | 2.8E+07 | 2.8E+07 | - | 4773 | armadillo repe | 0.4549 | 2.35969 | 10.8032 | 0.0016914 | 0.013026 |
| WDR4 | 21 | 4.3E+07 | 4.3E+07 | - | 2738 | WD repeat dor | -0.1543 | 5.681 | 10.803 | 0.0016916 | 0.013026 |
| PLK4 | 4 | 1.3E+08 | 1.3E+08 | + | 4479 | polo like kinas | -0.1539 | 5.74065 | 10.7997 | 0.0016941 | 0.013038 |
| AJAP1 | 1 | 4654732 | 4792534 | + | 12405 | adherens junct | -0.5695 | 1.87424 | 10.7968 | 0.0016963 | 0.013044 |
| SLC16A1-/ | 1 | 1.1E+08 | 1.1E+08 | + | 10138 | SLC16A1 antise | -0.318 | 3.49697 | 10.7964 | 0.0016966 | 0.013044 |
| CTSL | 9 | 8.8E+07 | 8.8E+07 | + | 1852 | cathepsin L [Sc | -0.1762 | 5.24435 | 10.7884 | 0.0017028 | 0.013085 |
| SETD7 | 4 | 1.4E+08 | 1.4E+08 | - | 13916 | SET domain co | 0.29935 | 3.65747 | 10.7852 | 0.0017053 | 0.013097 |
| DSG2 | 18 | 3.1E+07 | 3.2E+07 | + | 6056 | desmoglein 2 [ | -0.0925 | 8.37428 | 10.8675 | 0.0017099 | 0.013125 |
| IFT52 | 20 | 4.4E+07 | 4.4E+07 | + | 2542 | intraflagellar tr | -0.193 | 4.98416 | 10.7677 | 0.0017191 | 0.013188 |
| SMAD2 | 18 | 4.8E+07 | 4.8E+07 | - | 36426 | SMAD family n | -0.1254 | 6.36816 | 10.7648 | 0.0017214 | 0.013199 |
| SCAMP1 | 5 | 7.8E+07 | 7.8E+07 | + | 6746 | secretory carri | -0.147 | 5.91892 | 10.7577 | 0.001727 | 0.013235 |
| SLC7A1 | 13 | 3E+07 | 3E+07 | - | 7688 | solute carrier f | -0.098 | 7.6102 | 10.7449 | 0.0017371 | 0.013305 |
| INO80E | 16 | 3E+07 | 3E+07 | + | 5655 | INO80 complex | 0.14927 | 5.78566 | 10.7413 | 0.0017399 | 0.01332 |
| MSN | X | 6.6E+07 | 6.6E+07 | + | 5987 | moesin [Source | 0.08372 | 8.75967 | 10.7399 | 0.0017411 | 0.013321 |
| RAD17 | SCHR5_2_ | 6.9E+07 | 6.9E+07 | + | 4159 | RAD17 checkpo | -0.1877 | 5.11061 | 10.7368 | 0.0017436 | 0.013333 |
| PCDH18 | 4 | 1.4E+08 | 1.4E+08 | - | 6686 | protocadherin | -0.1344 | 6.16342 | 10.7356 | 0.0017445 | 0.013333 |
| PLBD1 | 12 | 1.5E+07 | 1.5E+07 | - | 2375 | phospholipase | 0.21142 | 4.74245 | 10.7257 | 0.0017524 | 0.013385 |
| PPP4R2 | 3 | 7.3E+07 | 7.3E+07 | + | 5964 | protein phosph | -0.1231 | 6.46462 | 10.7249 | 0.0017531 | 0.013385 |
| SCN9A | 2 | 1.7E+08 | 1.7E+08 | - | 13741 | sodium voltage | 0.37277 | 2.89847 | 10.7189 | 0.0017579 | 0.013414 |
| MPP1 | X | 1.5E+08 | 1.5E+08 | - | 5396 | membrane pal | -0.224 | 4.47989 | 10.7172 | 0.0017593 | 0.013414 |
| GCA | 2 | 1.6E+08 | 1.6E+08 | + | 5323 | grancalcin [Sou | 0.19241 | 5.00854 | 10.7167 | 0.0017597 | 0.013414 |
| RFX7 | 15 | 5.6E+07 | 5.6E+07 | - | 10588 | regulatory fact | 0.12992 | 6.32541 | 10.707 | 0.0017676 | 0.013467 |
| TOR1AIP2 | 1 | 1.8E+08 | 1.8E+08 | - | 18876 | torsin 1A inter | -0.1679 | 5.41386 | 10.7041 | 0.0017699 | 0.013477 |
| NNT | 5 | 4.4E+07 | 4.4E+07 | + | 8192 | nicotinamide n | 0.17351 | 5.30622 | 10.7012 | 0.0017722 | 0.013488 |
| LDHD | 16 | 7.5E+07 | 7.5E+07 | - | 2359 | lactate dehydr | -0.4005 | 2.87035 | 10.7123 | 0.0017747 | 0.0135 |
| CSRNP3 | 2 | 1.7E+08 | 1.7E+08 | + | 12565 | cysteine and se | 0.30671 | 3.57556 | 10.6875 | 0.0017834 | 0.013558 |
| NFE2L1 | 17 | 4.8E+07 | 4.8E+07 | + | 5883 | nuclear factor, | -0.0864 | 8.52414 | 10.6768 | 0.0017921 | 0.013618 |
| ZNF496 | 1 | 2.5E+08 | 2.5E+08 | - | 10531 | zinc finger prot | 0.10893 | 6.95637 | 10.6712 | 0.0017967 | 0.013639 |
| TPMT | 6 | 1.8E+07 | 1.8E+07 | - | 3184 | thiopurine S-m | -0.2279 | 4.46333 | 10.6711 | 0.0017968 | 0.013639 |
| RASGEF1A | 10 | 4.3E+07 | 4.3E+07 | - | 6073 | RasGEF domain | -0.2012 | 4.91143 | 10.6561 | 0.0018093 | 0.013719 |
| DOCK11 | X | 1.2E+08 | 1.2E+08 | + | 6721 | dedicator of cy | -0.2286 | 4.48657 | 10.649 | 0.0018152 | 0.013756 |
| KIAA1671 | 22 | 2.5E+07 | 2.5E+07 | + | 16439 | KIAA1671 [Sou | -0.1571 | 5.84228 | 10.7187 | 0.0018288 | 0.013838 |
| PRDM15 | 21 | 4.2E+07 | 4.2E+07 | - | 13004 | PR/SET domain | -0.2151 | 4.59355 | 10.6308 | 0.0018304 | 0.013842 |
| CAP2 | 6 | 1.7E+07 | 1.8E+07 | + | 3346 | cyclase associa | 0.20354 | 4.85688 | 10.629 | 0.0018319 | 0.013847 |
| NEK9 | 14 | 7.5E+07 | 7.5E+07 | - | 10760 | NIMA related k | 0.13085 | 6.2039 | 10.6253 | 0.001835 | 0.013857 |
| GID8 | 20 | 6.3E+07 | 6.3E+07 | + | 4594 | GID complex su | -0.1063 | 7.04804 | 10.625 | 0.0018352 | 0.013857 |
| CEP170 | 1 | 2.4E+08 | 2.4E+08 | - | 10939 | centrosomal p | 0.13116 | 6.22203 | 10.6215 | 0.0018382 | 0.013866 |
| PEPD | 19 | 3.3E+07 | 3.4E+07 | - | 4581 | peptidase D [S | -0.1234 | 6.3998 | 10.6214 | 0.0018383 | 0.013866 |
| TMCC2 | 1 | 2.1E+08 | 2.1E+08 | + | 5822 | transmembran | -0.2571 | 4.10087 | 10.6176 | 0.0018415 | 0.013877 |
| KLF13 | 15 | 3.1E+07 | 3.1E+07 | + | 9429 | Kruppel like fa | -0.1257 | 6.44714 | 10.6174 | 0.0018417 | 0.013877 |

| | | | | | | | | | | |
|---|---|---|---|---|---|---|---|---|---|---|
| DTX3 | 12 | 5.8E+07 | 5.8E+07 | + | 4087 | deltex E3 ubiq | 0.18203 | 5.22357 | 10.6094 | 0.0018485 | 0.013921 |
| RAB11FIP | 16 | 425649 | 523011 | + | 6287 | RAB11 family i | 0.14271 | 5.93288 | 10.5979 | 0.0018583 | 0.013987 |
| ELP2 | 18 | 3.6E+07 | 3.6E+07 | + | 11060 | elongator acet | -0.1068 | 7.02588 | 10.5943 | 0.0018613 | 0.014003 |
| PCED1A | 20 | 2835314 | 2841190 | - | 2320 | PC-esterase do | 0.34991 | 3.30211 | 10.6823 | 0.0018664 | 0.014034 |
| GBGT1 | 9 | 1.3E+08 | 1.3E+08 | - | 4020 | globoside alph | 0.43386 | 2.486 | 10.5872 | 0.0018674 | 0.014034 |
| CCDC3 | 10 | 1.3E+07 | 1.3E+07 | - | 3699 | coiled-coil dom | -0.4816 | 2.27502 | 10.5848 | 0.0018694 | 0.014042 |
| KIF3A | 5 | 1.3E+08 | 1.3E+08 | - | 6860 | kinesin family | -0.2454 | 4.2662 | 10.5803 | 0.0018734 | 0.014059 |
| CALCRL | 2 | 1.9E+08 | 1.9E+08 | - | 7309 | calcitonin rece | 0.53392 | 1.95161 | 10.5795 | 0.001874 | 0.014059 |
| RRBP1 | 20 | 1.8E+07 | 1.8E+07 | - | 6980 | ribosome bind | -0.0871 | 8.20201 | 10.5788 | 0.0018747 | 0.014059 |
| HMMR | 5 | 1.6E+08 | 1.6E+08 | + | 3932 | hyaluronan me | -0.1205 | 6.48678 | 10.5637 | 0.0018876 | 0.014149 |
| ADCYAP1 | 7 | 3.1E+07 | 3.1E+07 | + | 6832 | ADCYAP recept | -0.2127 | 4.60749 | 10.5604 | 0.0018905 | 0.014163 |
| NOL11 | 17 | 6.8E+07 | 6.8E+07 | + | 4615 | nucleolar prot | -0.1173 | 6.72088 | 10.5502 | 0.0018994 | 0.014223 |
| IPO9 | 1 | 2E+08 | 2E+08 | + | 11904 | importin 9 [So | 0.09344 | 7.83778 | 10.5461 | 0.001903 | 0.014237 |
| TSBP1-AS | 6 | 3.2E+07 | 3.2E+07 | + | 8816 | TSBP1 and BTN | -0.6264 | 1.5126 | 10.5457 | 0.0019034 | 0.014237 |
| PIK3AP1 | 10 | 9.7E+07 | 9.7E+07 | - | 5688 | phosphoinositi | 0.44208 | 2.40961 | 10.5426 | 0.001906 | 0.014249 |
| ITPR3 | 6 | 3.4E+07 | 3.4E+07 | + | 9870 | inositol 1,4,5-t | 0.14896 | 6.91461 | 12.2409 | 0.001907 | 0.014249 |
| SPRY1 | 4 | 1.2E+08 | 1.2E+08 | + | 3963 | sprouty RTK sig | 0.18762 | 5.02565 | 10.5375 | 0.0019105 | 0.014268 |
| CDHR1 | 10 | 8.4E+07 | 8.4E+07 | + | 7729 | cadherin relate | -0.3259 | 3.44935 | 10.5328 | 0.0019147 | 0.014292 |
| ATP6AP1L | 5 | 8.2E+07 | 8.2E+07 | + | 7229 | ATPase H+ tran | 0.73064 | 1.08005 | 10.5297 | 0.0019174 | 0.014295 |
| CCBE1 | 18 | 5.9E+07 | 6E+07 | - | 6753 | collagen and ca | -0.3822 | 2.91929 | 10.5291 | 0.0019179 | 0.014295 |
| NAAA | 4 | 7.6E+07 | 7.6E+07 | - | 4418 | N-acylethanola | -0.3821 | 2.90718 | 10.5286 | 0.0019184 | 0.014295 |
| PSMA5 | 1 | 1.1E+08 | 1.1E+08 | - | 5734 | proteasome su | -0.101 | 7.25371 | 10.5278 | 0.0019191 | 0.014295 |
| FAM49A | 2 | 1.7E+07 | 1.7E+07 | - | 5448 | family with seq | -0.3297 | 3.29599 | 10.5256 | 0.001921 | 0.014302 |
| LIN7C | 11 | 2.7E+07 | 2.8E+07 | - | 4842 | lin-7 homolog | -0.1908 | 4.96729 | 10.5208 | 0.0019253 | 0.014327 |
| EFEMP2 | 11 | 6.6E+07 | 6.6E+07 | - | 5397 | EGF containing | 0.3507 | 3.07577 | 10.5163 | 0.0019292 | 0.014349 |
| TAX1BP1 | 7 | 2.8E+07 | 2.8E+07 | + | 9165 | Tax1 binding p | -0.1138 | 6.78436 | 10.5142 | 0.0019311 | 0.014355 |
| NPIPB5 | 16 | 2.2E+07 | 2.3E+07 | + | 10005 | nuclear pore c | -0.2324 | 4.4332 | 10.5118 | 0.0019333 | 0.014358 |
| AC022893 | 8 | 7.3E+07 | 7.3E+07 | + | 3549 | novel transcrip | -0.3375 | 3.48905 | 10.6923 | 0.0019352 | 0.014358 |
| CDIP1 | 16 | 4510669 | 4538828 | - | 4274 | cell death indu | -0.1207 | 6.61328 | 10.5092 | 0.0019355 | 0.014358 |
| PORCN | X | 4.9E+07 | 4.9E+07 | + | 3011 | porcupine O-ac | -0.2595 | 3.99876 | 10.5078 | 0.0019368 | 0.01436 |
| GPX7 | 1 | 5.3E+07 | 5.3E+07 | + | 1692 | glutathione pe | 0.32899 | 3.25583 | 10.5057 | 0.0019387 | 0.01436 |
| OGFRL1 | 6 | 7.1E+07 | 7.1E+07 | + | 9176 | opioid growth | 0.24172 | 4.57642 | 10.8342 | 0.0019392 | 0.01436 |
| PRODH | 22 | 1.9E+07 | 1.9E+07 | - | 8776 | proline dehydr | -0.5186 | 2.02233 | 10.5045 | 0.0019397 | 0.01436 |
| FANCD2 | 3 | 1E+07 | 1E+07 | + | 8378 | FA complemen | 0.11247 | 6.82955 | 10.4975 | 0.001946 | 0.014398 |
| P3H3 | 12 | 6828407 | 6839847 | + | 4600 | prolyl 3-hydrox | 0.15498 | 5.69561 | 10.4965 | 0.001947 | 0.014398 |
| NEU1 | R6_MHC_ | 3.2E+07 | 3.2E+07 | - | 4822 | neuraminidase | -0.1579 | 5.6358 | 10.4824 | 0.0019596 | 0.014484 |
| NUF2 | 1 | 1.6E+08 | 1.6E+08 | + | 3362 | NDC80 kinetoc | -0.2324 | 4.39264 | 10.4784 | 0.0019632 | 0.014504 |
| ZNF195 | 11 | 3339261 | 3379222 | - | 11711 | zinc finger prot | -0.1604 | 5.54495 | 10.466 | 0.0019744 | 0.014569 |
| VAV1 | 19 | 6772708 | 6857366 | + | 4311 | vav guanine nu | -0.2923 | 3.66031 | 10.4657 | 0.0019747 | 0.014569 |
| CALR | 19 | 1.3E+07 | 1.3E+07 | + | 2714 | calreticulin [So | -0.0724 | 9.92821 | 10.4652 | 0.0019751 | 0.014569 |
| CDKN1B | 12 | 1.3E+07 | 1.3E+07 | + | 2565 | cyclin depende | 0.17143 | 5.38589 | 10.4642 | 0.0019761 | 0.014569 |
| DNM1 | 9 | 1.3E+08 | 1.3E+08 | + | 10775 | dynamin 1 [Sou | 0.18927 | 5.10445 | 10.4612 | 0.0019788 | 0.014582 |
| TAF3 | 10 | 7818505 | 8016631 | + | 4875 | TATA-box bind | -0.2037 | 4.7488 | 10.4568 | 0.0019829 | 0.014595 |

| | | | | | | | | | | | |
|---|---|---|---|---|---|---|---|---|---|---|---|
| HCCS | X | 1.1E+07 | 1.1E+07 | + | 2343 | holocytochrom | -0.1842 | 5.07934 | 10.4561 | 0.0019834 | 0.014595 |
| TRIM25 | 17 | 5.7E+07 | 5.7E+07 | - | 9501 | tripartite motif | -0.1422 | 5.94634 | 10.4558 | 0.0019837 | 0.014595 |
| FGD5-AS1 | 3 | 1.5E+07 | 1.5E+07 | - | 5131 | FGD5 antisense | 0.10715 | 6.99107 | 10.4449 | 0.0019937 | 0.014661 |
| EBNA1BP | 1 | 4.3E+07 | 4.3E+07 | - | 3561 | EBNA1 binding | -0.1178 | 6.65644 | 10.4426 | 0.0019959 | 0.014669 |
| SLC16A3 | 17 | 8.2E+07 | 8.2E+07 | + | 8995 | solute carrier f | 0.44437 | 2.64462 | 10.6477 | 0.0019989 | 0.014684 |
| CNTNAP3 | 9 | 3.9E+07 | 3.9E+07 | - | 9521 | contactin asso | -0.476 | 2.20019 | 10.4361 | 0.0020018 | 0.014698 |
| GRAMD1A | 19 | 3.5E+07 | 3.5E+07 | + | 5942 | GRAM domain | -0.1001 | 7.30787 | 10.4345 | 0.0020033 | 0.014701 |
| SEMA6C | 1 | 1.5E+08 | 1.5E+08 | - | 5809 | semaphorin 6C | 0.20176 | 4.87455 | 10.4267 | 0.0020105 | 0.014747 |
| MAP7D2 | X | 2E+07 | 2E+07 | - | 4366 | MAP7 domain | 0.65306 | 1.32352 | 10.4223 | 0.0020146 | 0.014765 |
| NLGN1 | 3 | 1.7E+08 | 1.7E+08 | + | 10148 | neuroligin 1 [S | 0.22754 | 4.4388 | 10.4218 | 0.002015 | 0.014765 |
| FGFR4 | 5 | 1.8E+08 | 1.8E+08 | + | 4810 | fibroblast grow | -0.1066 | 7.0086 | 10.4104 | 0.0020257 | 0.014832 |
| POP5 | 12 | 1.2E+08 | 1.2E+08 | - | 2173 | POP5 homolog | 0.22829 | 4.37226 | 10.4098 | 0.0020263 | 0.014832 |
| ZNF267 | 16 | 3.2E+07 | 3.2E+07 | + | 6284 | zinc finger prot | -0.2491 | 4.14469 | 10.4074 | 0.0020285 | 0.014841 |
| PIPOX | 17 | 2.9E+07 | 2.9E+07 | + | 3814 | pipecolic acid a | 0.16012 | 5.62387 | 10.4002 | 0.0020352 | 0.014882 |
| BCYRN1 | 2 | 4.7E+07 | 4.7E+07 | + | 200 | brain cytoplasm | 0.5843 | 1.64434 | 10.3985 | 0.0020368 | 0.014887 |
| ACACB | 12 | 1.1E+08 | 1.1E+08 | + | 14505 | acetyl-CoA car | -0.2353 | 4.35926 | 10.3915 | 0.0020434 | 0.014927 |
| TMEM256 | 17 | 7402975 | 7404097 | - | 667 | transmembran | 0.29706 | 3.60366 | 10.3866 | 0.002048 | 0.014953 |
| FAM120A | 9 | 9.3E+07 | 9.3E+07 | - | 10144 | family with sec | 0.17753 | 5.16841 | 10.385 | 0.0020496 | 0.014956 |
| NHSL2 | X | 7.2E+07 | 7.2E+07 | + | 17706 | NHS like 2 [Sou | 0.18557 | 5.03176 | 10.3841 | 0.0020504 | 0.014956 |
| CYHR1 | 8 | 1.4E+08 | 1.4E+08 | - | 9787 | cysteine and h | 0.14912 | 5.90493 | 10.4014 | 0.0020564 | 0.014984 |
| TMEM178 | 2 | 4E+07 | 4E+07 | + | 2472 | transmembran | 0.43432 | 2.40724 | 10.3777 | 0.0020564 | 0.014984 |
| JUP | 17 | 4.2E+07 | 4.2E+07 | - | 4942 | junction plako | -0.0926 | 7.72226 | 10.3758 | 0.0020582 | 0.01499 |
| UTP3 | 4 | 7.1E+07 | 7.1E+07 | + | 2020 | UTP3, small su | -0.1471 | 5.82732 | 10.3718 | 0.0020621 | 0.01501 |
| LRRN1 | 3 | 3799437 | 3847703 | + | 4264 | leucine rich re | 0.0867 | 8.257 | 10.3653 | 0.0020682 | 0.015047 |
| PLXND1 | 3 | 1.3E+08 | 1.3E+08 | - | 9738 | plexin D1 [Sou | 0.15392 | 7.10686 | 12.688 | 0.0020715 | 0.015063 |
| KLK8 | 19 | 5.1E+07 | 5.1E+07 | - | 1865 | kallikrein relat | -0.7802 | 1.02256 | 10.3543 | 0.0020788 | 0.015109 |
| KCNMA1 | 10 | 7.7E+07 | 7.8E+07 | - | 35644 | potassium calc | -0.4282 | 2.58783 | 10.3474 | 0.0020854 | 0.015143 |
| SMC2 | 9 | 1E+08 | 1E+08 | + | 6470 | structural mair | -0.1168 | 6.67031 | 10.3472 | 0.0020856 | 0.015143 |
| FAM189A | SCHR15_4 | 2.9E+07 | 3E+07 | - | 5545 | family with sec | 0.27793 | 3.92648 | 10.3965 | 0.0020868 | 0.015144 |
| ARSA | 22 | 5.1E+07 | 5.1E+07 | - | 4501 | arylsulfatase A | 0.31983 | 3.39157 | 10.3445 | 0.0020882 | 0.015147 |
| GPS1 | 17 | 8.2E+07 | 8.2E+07 | + | 6155 | G protein path | 0.10235 | 7.20455 | 10.3356 | 0.0020968 | 0.015201 |
| PPIE | 1 | 4E+07 | 4E+07 | + | 8071 | peptidylprolyl | 0.19007 | 4.99118 | 10.3309 | 0.0021013 | 0.015227 |
| FXN | 9 | 6.9E+07 | 6.9E+07 | + | 8163 | frataxin [Sourc | -0.2343 | 4.28762 | 10.3187 | 0.0021132 | 0.015305 |
| TASOR2 | 10 | 5684838 | 5763740 | + | 10960 | transcription a | 0.11292 | 6.91739 | 10.3296 | 0.002116 | 0.015317 |
| SMC1A | X | 5.3E+07 | 5.3E+07 | - | 10484 | structural mair | 0.08908 | 8.10845 | 10.307 | 0.0021247 | 0.015365 |
| RAB24 | 5 | 1.8E+08 | 1.8E+08 | - | 2065 | RAB24, memb | 0.45761 | 2.26265 | 10.3045 | 0.0021272 | 0.015375 |
| ZNF254 | 19 | 2.4E+07 | 2.4E+07 | + | 7602 | zinc finger prot | -0.2105 | 4.88027 | 10.4112 | 0.0021323 | 0.015404 |
| FGF12 | 3 | 1.9E+08 | 1.9E+08 | - | 7355 | fibroblast grow | 0.28107 | 3.70818 | 10.2963 | 0.0021352 | 0.015418 |
| TUBA1B | 12 | 4.9E+07 | 4.9E+07 | - | 3574 | tubulin alpha 1 | 0.07897 | 8.9747 | 10.2881 | 0.0021433 | 0.015465 |
| TRMT9B | 8 | 1.3E+07 | 1.3E+07 | + | 14669 | tRNA methyltr | 0.82647 | 0.74901 | 10.2875 | 0.0021439 | 0.015465 |
| MRPS35 | 12 | 2.8E+07 | 2.8E+07 | + | 1930 | mitochondrial | -0.1365 | 6.09443 | 10.2812 | 0.0021501 | 0.015502 |
| CYP20A1 | 2 | 2E+08 | 2E+08 | + | 3079 | cytochrome P4 | -0.2953 | 3.67408 | 10.2791 | 0.0021523 | 0.01551 |
| FBXO44 | 1 | 1.2E+07 | 1.2E+07 | + | 3527 | F-box protein 4 | 0.20706 | 4.74682 | 10.271 | 0.0021603 | 0.01556 |

| | | | | | | | | | | | |
|---|---|---|---|---|---|---|---|---|---|---|---|
| CCDC117 | 22 | 2.9E+07 | 2.9E+07 | + | 4326 | coiled-coil dom | -0.1434 | 5.96347 | 10.265 | 0.0021664 | 0.015591 |
| PIK3R5 | 17 | 8878911 | 8965712 | - | 7506 | phosphoinositi | -0.3242 | 3.27995 | 10.2645 | 0.0021669 | 0.015591 |
| CCL26 | 7 | 7.6E+07 | 7.6E+07 | - | 706 | C-C motif chen | -0.5053 | 2.03855 | 10.2513 | 0.0021801 | 0.015672 |
| DLC1 | 8 | 1.3E+07 | 1.4E+07 | - | 12794 | DLC1 Rho GTPa | -0.422 | 2.56169 | 10.2503 | 0.0021811 | 0.015672 |
| LINC0257 | 21 | 4.4E+07 | 4.4E+07 | + | 912 | long intergenic | -0.2907 | 3.62647 | 10.2493 | 0.0021821 | 0.015672 |
| E4F1 | 16 | 2223580 | 2235742 | + | 6388 | E4F transcripti | 0.18306 | 5.03525 | 10.249 | 0.0021825 | 0.015672 |
| HSD11B2 | 16 | 6.7E+07 | 6.7E+07 | + | 2670 | hydroxysteroid | 0.17149 | 5.26739 | 10.2467 | 0.0021847 | 0.015681 |
| APELA | 4 | 1.6E+08 | 1.6E+08 | + | 2919 | apelin recepto | -0.2495 | 5.75402 | 13.7081 | 0.0021881 | 0.015697 |
| SNRPF | 12 | 9.6E+07 | 9.6E+07 | + | 3043 | small nuclear r | -0.1067 | 6.91669 | 10.237 | 0.0021946 | 0.015735 |
| NOL9 | 1 | 6521347 | 6554513 | - | 7271 | nucleolar prote | -0.184 | 5.0179 | 10.2311 | 0.0022006 | 0.015771 |
| MASTL | 10 | 2.7E+07 | 2.7E+07 | + | 3631 | microtubule as | 0.13783 | 5.96334 | 10.2185 | 0.0022135 | 0.015855 |
| ANKLE2 | 12 | 1.3E+08 | 1.3E+08 | - | 17461 | ankyrin repeat | 0.12778 | 6.24638 | 10.2141 | 0.0022179 | 0.015879 |
| EEA1 | 12 | 9.3E+07 | 9.3E+07 | - | 10150 | early endosom | -0.1711 | 5.29525 | 10.2108 | 0.0022213 | 0.015896 |
| SLC25A13 | 7 | 9.6E+07 | 9.6E+07 | - | 3710 | solute carrier f | -0.1043 | 7.16369 | 10.2048 | 0.0022275 | 0.015931 |
| ZKSCAN8 | 6 | 2.8E+07 | 2.8E+07 | + | 7580 | zinc finger with | 0.15039 | 5.8 | 10.2039 | 0.0022285 | 0.015931 |
| RPL29 | 3 | 5.2E+07 | 5.2E+07 | - | 2058 | ribosomal prot | 0.07575 | 9.31358 | 10.1935 | 0.0022392 | 0.015999 |
| RHOT2 | 16 | 668105 | 674174 | + | 5155 | ras homolog fa | 0.0993 | 7.23328 | 10.1894 | 0.0022434 | 0.016011 |
| TRAF4 | 17 | 2.9E+07 | 2.9E+07 | + | 4573 | TNF receptor a | 0.09571 | 7.47856 | 10.188 | 0.0022449 | 0.016011 |
| SLC1A2 | 11 | 3.5E+07 | 3.5E+07 | - | 22800 | solute carrier f | -0.8586 | 0.7325 | 10.1879 | 0.002245 | 0.016011 |
| ANKZF1 | 2 | 2.2E+08 | 2.2E+08 | + | 6119 | ankyrin repeat | 0.1565 | 5.5565 | 10.1876 | 0.0022453 | 0.016011 |
| DBNDD1 | 16 | 9E+07 | 9E+07 | - | 4791 | dysbindin dom | 0.11808 | 6.55286 | 10.1857 | 0.0022474 | 0.016018 |
| FN3KRP | 17 | 8.3E+07 | 8.3E+07 | + | 2944 | fructosamine 3 | 0.19708 | 4.81703 | 10.1798 | 0.0022535 | 0.016054 |
| INTS3 | SCHR1_1_ | 1.5E+08 | 1.5E+08 | + | 10467 | integrator com | 0.11277 | 6.68411 | 10.1699 | 0.0022639 | 0.01612 |
| USP54 | 10 | 7.3E+07 | 7.4E+07 | - | 9498 | ubiquitin speci | 0.14789 | 5.79629 | 10.1548 | 0.0022798 | 0.016222 |
| PTPA | 9 | 1.3E+08 | 1.3E+08 | + | 5624 | protein phosph | -0.0883 | 8.03711 | 10.153 | 0.0022816 | 0.016222 |
| GPR143 | X | 9725346 | 9786297 | - | 2369 | G protein-coup | -0.1602 | 5.47017 | 10.153 | 0.0022816 | 0.016222 |
| DIRAS1 | 19 | 2714567 | 2721372 | - | 3439 | DIRAS family G | 0.16783 | 5.40793 | 10.1502 | 0.0022845 | 0.016235 |
| RAB5IF | 20 | 3.7E+07 | 3.7E+07 | + | 1504 | RAB5 interacti | 0.18625 | 5.08643 | 10.1489 | 0.0022859 | 0.016237 |
| KCNT2 | 1 | 2E+08 | 2E+08 | - | 7100 | potassium sod | 0.28805 | 3.6059 | 10.1385 | 0.0022971 | 0.016307 |
| TLN2 | 15 | 6.2E+07 | 6.3E+07 | + | 16209 | talin 2 [Source | 0.12489 | 6.36415 | 10.1339 | 0.0023019 | 0.016334 |
| EPCAM | 2 | 4.7E+07 | 4.7E+07 | + | 2519 | epithelial cell a | -0.0949 | 7.55374 | 10.1247 | 0.0023118 | 0.016396 |
| MBNL1 | 3 | 1.5E+08 | 1.5E+08 | + | 9989 | muscleblind lik | 0.27491 | 3.81299 | 10.1165 | 0.0023206 | 0.01645 |
| ANP32A | 15 | 6.9E+07 | 6.9E+07 | - | 7544 | acidic nuclear p | -0.0881 | 8.20144 | 10.1124 | 0.0023249 | 0.01647 |
| KIF2C | 1 | 4.5E+07 | 4.5E+07 | + | 3646 | kinesin family n | 0.10594 | 7.10649 | 10.1117 | 0.0023257 | 0.01647 |
| LINC0123 | 13 | 9.9E+07 | 9.9E+07 | - | 6199 | long intergenic | 0.69615 | 1.12275 | 10.1063 | 0.0023315 | 0.016503 |
| DKK3 | 11 | 1.2E+07 | 1.2E+07 | - | 5059 | dickkopf WNT | -0.1916 | 5.07068 | 10.1289 | 0.0023359 | 0.016526 |
| CSTF1 | 20 | 5.6E+07 | 5.6E+07 | + | 4800 | cleavage stimu | -0.152 | 5.7631 | 10.0953 | 0.0023435 | 0.016572 |
| ZNF587B | 19 | 5.8E+07 | 5.8E+07 | + | 5312 | zinc finger prot | -0.1752 | 5.1997 | 10.0938 | 0.0023451 | 0.016575 |
| CASC15 | 6 | 2.2E+07 | 2.3E+07 | + | 27177 | cancer suscept | 0.35827 | 3.06836 | 10.0878 | 0.0023517 | 0.016613 |
| SQSTM1 | 5 | 1.8E+08 | 1.8E+08 | + | 7535 | sequestosome | 0.1192 | 6.57633 | 10.0718 | 0.0023692 | 0.016729 |
| TMEM99 | 17 | 4.1E+07 | 4.1E+07 | + | 2362 | transmembran | -0.4068 | 2.70848 | 10.0655 | 0.0023762 | 0.01677 |
| HRH3 | 20 | 6.2E+07 | 6.2E+07 | - | 2692 | histamine rece | -0.4476 | 2.29897 | 10.062 | 0.0023801 | 0.016789 |
| AL050341 | 1 | 4E+07 | 4E+07 | - | 1541 | novel transcrip | -0.4247 | 2.47558 | 10.0595 | 0.0023827 | 0.0168 |

| | | | | | | | | | | | |
|---|---|---|---|---|---|---|---|---|---|---|---|
| MAPK1IP1 | 14 | 5.5E+07 | 5.5E+07 | + | 6974 | mitogen-activa | -0.1108 | 6.9142 | 10.054 | 0.0023889 | 0.016834 |
| PRDM5 | 4 | 1.2E+08 | 1.2E+08 | - | 9517 | PR/SET domair | 0.17979 | 5.09233 | 10.0488 | 0.0023946 | 0.016867 |
| FAM219B | 15 | 7.5E+07 | 7.5E+07 | - | 5015 | family with sec | 0.23321 | 4.21871 | 10.0472 | 0.0023965 | 0.016871 |
| BTN3A2 | 6 | 2.6E+07 | 2.6E+07 | + | 7042 | butyrophilin su | 0.43928 | 2.39733 | 10.0455 | 0.0023984 | 0.016877 |
| PDLIM5 | 4 | 9.4E+07 | 9.5E+07 | + | 11090 | PDZ and LIM d | 0.12104 | 6.46994 | 10.0389 | 0.0024057 | 0.016911 |
| CNNM4 | 2 | 9.7E+07 | 9.7E+07 | + | 5088 | cyclin and CBS | -0.2095 | 4.60444 | 10.0338 | 0.0024114 | 0.016944 |
| INSYN1 | 15 | 7.4E+07 | 7.4E+07 | - | 6573 | inhibitory syna | -0.3328 | 3.75808 | 10.772 | 0.0024435 | 0.017161 |
| CAPN6 | X | 1.1E+08 | 1.1E+08 | - | 3532 | calpain 6 [Sour | 0.21186 | 4.60485 | 9.9988 | 0.002451 | 0.017203 |
| PCDHB15 | 5 | 1.4E+08 | 1.4E+08 | + | 3971 | protocadherin | 0.27242 | 3.81857 | 9.99797 | 0.0024519 | 0.017203 |
| TRAIP | 3 | 5E+07 | 5E+07 | - | 3261 | TRAF interactir | 0.22295 | 4.36232 | 9.99536 | 0.0024549 | 0.017215 |
| RPL38 | 17 | 7.4E+07 | 7.4E+07 | + | 3829 | ribosomal prot | -0.1051 | 8.2097 | 10.8825 | 0.0024569 | 0.017221 |
| HMGB3 | X | 1.5E+08 | 1.5E+08 | + | 3914 | high mobility g | -0.0862 | 8.30936 | 9.99006 | 0.0024609 | 0.017241 |
| GAA | 17 | 8E+07 | 8E+07 | + | 5236 | glucosidase alp | 0.13779 | 6.05398 | 9.98712 | 0.0024643 | 0.017256 |
| NORAD | 20 | 3.6E+07 | 3.6E+07 | - | 5401 | non-coding RN | -0.0857 | 8.10379 | 9.97624 | 0.0024768 | 0.017331 |
| UGT8 | 4 | 1.1E+08 | 1.1E+08 | + | 4385 | UDP glycosyltr | 0.1644 | 5.58342 | 10.0337 | 0.0024774 | 0.017331 |
| EGLN3 | 14 | 3.4E+07 | 3.4E+07 | - | 7359 | egl-9 family hy | -0.2172 | 4.5669 | 9.9747 | 0.0024786 | 0.017331 |
| RIC8A | 11 | 207511 | 215113 | + | 6685 | RIC8 guanine n | 0.1138 | 6.58049 | 9.97025 | 0.0024837 | 0.017358 |
| TECPR1 | 7 | 9.8E+07 | 9.8E+07 | - | 9066 | tectonin beta-| | 0.22326 | 4.36796 | 9.95854 | 0.0024973 | 0.017445 |
| KCTD20 | 6 | 3.6E+07 | 3.6E+07 | + | 6239 | potassium cha | 0.12728 | 6.17146 | 9.95393 | 0.0025026 | 0.017474 |
| CUL4B | X | 1.2E+08 | 1.2E+08 | - | 7031 | cullin 4B [Sour | -0.127 | 6.17398 | 9.94752 | 0.0025101 | 0.017517 |
| GRB10 | 7 | 5.1E+07 | 5.1E+07 | - | 9795 | growth factor | 0.11768 | 6.86806 | 10.1287 | 0.0025212 | 0.017586 |
| S1PR2 | 19 | 1E+07 | 1E+07 | - | 3643 | sphingosine-1- | -0.32 | 3.37098 | 9.93686 | 0.0025226 | 0.017587 |
| OGA | 10 | 1E+08 | 1E+08 | - | 7247 | O-GlcNAcase [ | 0.08995 | 7.81771 | 9.9289 | 0.002532 | 0.017644 |
| RIOX2 | 3 | 9.8E+07 | 9.8E+07 | - | 6880 | ribosomal oxyg | 0.25232 | 3.9984 | 9.92708 | 0.0025341 | 0.01765 |
| RHOBTB2 | 8 | 2.3E+07 | 2.3E+07 | + | 6580 | Rho related BT | 0.27264 | 3.7765 | 9.92297 | 0.002539 | 0.017676 |
| DERA | 12 | 1.6E+07 | 1.6E+07 | + | 2813 | deoxyribose-pl | 0.26456 | 3.93072 | 9.9196 | 0.002543 | 0.017695 |
| ECI1 | 16 | 2239402 | 2252300 | - | 3204 | enoyl-CoA delt | 0.13962 | 5.98206 | 9.90968 | 0.0025547 | 0.017768 |
| KCNH6 | 17 | 6.4E+07 | 6.4E+07 | + | 4211 | potassium volt | -0.8048 | 0.81622 | 9.89943 | 0.002567 | 0.017845 |
| PLCB4 | 20 | 9068763 | 9481242 | + | 6900 | phospholipase | -0.2393 | 4.22145 | 9.8855 | 0.0025837 | 0.017943 |
| TONSL | 8 | 1.4E+08 | 1.4E+08 | - | 7107 | tonsoku like, D | 0.10832 | 6.74888 | 9.884 | 0.0025855 | 0.017943 |
| DMWD | 19 | 4.6E+07 | 4.6E+07 | - | 5701 | DM1 locus, WD | 0.19917 | 4.76291 | 9.8827 | 0.0025871 | 0.017943 |
| BCL9L | 11 | 1.2E+08 | 1.2E+08 | - | 10470 | BCL9 like [Sour | -0.1007 | 7.15646 | 9.88263 | 0.0025871 | 0.017943 |
| BANF1 | 11 | 6.6E+07 | 6.6E+07 | + | 1496 | barrier to auto | 0.09772 | 7.35913 | 9.88242 | 0.0025874 | 0.017943 |
| MAP3K8 | 10 | 3E+07 | 3E+07 | + | 3517 | mitogen-activa | -0.5003 | 1.96691 | 9.87726 | 0.0025936 | 0.017978 |
| ABCC5 | 3 | 1.8E+08 | 1.8E+08 | - | 11365 | ATP binding ca | 0.11636 | 6.51241 | 9.87592 | 0.0025953 | 0.01798 |
| ZNF703 | 8 | 3.8E+07 | 3.8E+07 | + | 3316 | zinc finger prot | -0.6971 | 1.33803 | 9.94351 | 0.0026041 | 0.018033 |
| CALN1 | 7 | 7.2E+07 | 7.2E+07 | - | 9997 | calneuron 1 [S | 0.20982 | 4.56107 | 9.86411 | 0.0026096 | 0.018059 |
| MSRB3 | 12 | 6.5E+07 | 6.5E+07 | + | 7405 | methionine su | 0.68012 | 1.29721 | 9.87372 | 0.0026116 | 0.018059 |
| PEBP1 | 12 | 1.2E+08 | 1.2E+08 | + | 1514 | phosphatidylet | -0.0765 | 9.064 | 9.86242 | 0.0026116 | 0.018059 |
| HOMER3 | 19 | 1.9E+07 | 1.9E+07 | - | 3422 | homer scaffold | 0.16901 | 5.30219 | 9.86098 | 0.0026134 | 0.018062 |
| TENT5C | 1 | 1.2E+08 | 1.2E+08 | + | 5654 | terminal nucle | -0.831 | 0.69277 | 9.8544 | 0.0026214 | 0.018109 |
| ZSWIM3 | 20 | 4.6E+07 | 4.6E+07 | + | 2776 | zinc finger SWI | -0.3808 | 2.77476 | 9.84987 | 0.002627 | 0.018139 |
| AL591030 | 6 | 7.7E+07 | 7.7E+07 | + | 627 | novel transcrip | -0.2063 | 4.82152 | 9.89845 | 0.0026367 | 0.018198 |

| | | | | | | | | | | | |
|---|---|---|---|---|---|---|---|---|---|---|---|
| CABLES1 | 18 | 2.3E+07 | 2.3E+07 | + | 6654 | Cdk5 and Abl e | -0.2051 | 4.85713 | 9.91055 | 0.0026439 | 0.018238 |
| AFF2 | X | 1.5E+08 | 1.5E+08 | + | 14006 | AF4/FMR2 fam | 0.26867 | 3.90189 | 9.83362 | 0.002647 | 0.018251 |
| ZNF880 | 19 | 5.2E+07 | 5.2E+07 | + | 3719 | zinc finger prot | -0.3589 | 2.98622 | 9.82205 | 0.0026613 | 0.018341 |
| C14orf132 | 14 | 9.6E+07 | 9.6E+07 | + | 7775 | chromosome 1 | -0.1567 | 5.53521 | 9.8143 | 0.0026709 | 0.018398 |
| MICB | R6_MHC_ | 3.1E+07 | 3.2E+07 | + | 2798 | MHC class I po | -0.2807 | 3.64082 | 9.8113 | 0.0026747 | 0.018415 |
| SETDB1 | 1 | 1.5E+08 | 1.5E+08 | + | 7093 | SET domain bif | 0.15437 | 5.57837 | 9.80641 | 0.0026808 | 0.018442 |
| OS9 | 12 | 5.8E+07 | 5.8E+07 | + | 5169 | OS9, endoplas | 0.11145 | 6.63216 | 9.80613 | 0.0026811 | 0.018442 |
| AC080038 | 17 | 6.3E+07 | 6.3E+07 | + | 6591 | mannose recep | 0.18243 | 5.10694 | 9.80245 | 0.0026858 | 0.01845 |
| AFDN | 6 | 1.7E+08 | 1.7E+08 | + | 13720 | afadin, adhere | 0.08335 | 8.24316 | 9.80217 | 0.0026861 | 0.01845 |
| ACTG1 | 17 | 8.2E+07 | 8.2E+07 | - | 2993 | actin gamma 1 | 0.06511 | 11.7966 | 9.80211 | 0.0026862 | 0.01845 |
| WASF2 | 1 | 2.7E+07 | 2.7E+07 | - | 5681 | WAS protein fa | 0.09109 | 7.70616 | 9.799 | 0.0026901 | 0.018468 |
| SAT1 | X | 2.4E+07 | 2.4E+07 | + | 2498 | spermidine/sp | -0.1878 | 4.87761 | 9.79608 | 0.0026938 | 0.018485 |
| STX3 | 11 | 6E+07 | 6E+07 | + | 7308 | syntaxin 3 [Sou | -0.1189 | 6.59256 | 9.80718 | 0.0026978 | 0.018496 |
| ZFPM1 | 16 | 8.8E+07 | 8.9E+07 | + | 6162 | zinc finger prot | 0.31526 | 3.33697 | 9.79276 | 0.0026979 | 0.018496 |
| COPE | 19 | 1.9E+07 | 1.9E+07 | - | 2875 | coatomer prot | 0.11124 | 6.63856 | 9.78973 | 0.0027018 | 0.018513 |
| RNF138 | 18 | 3.2E+07 | 3.2E+07 | + | 3776 | ring finger prot | -0.1135 | 6.60883 | 9.78563 | 0.0027069 | 0.018536 |
| MGAT1 | 5 | 1.8E+08 | 1.8E+08 | - | 11576 | mannosyl (alph | 0.10398 | 6.9509 | 9.78497 | 0.0027078 | 0.018536 |
| SUGP2 | 19 | 1.9E+07 | 1.9E+07 | - | 8468 | SURP and G-pa | 0.09469 | 7.38887 | 9.78386 | 0.0027092 | 0.018537 |
| C7orf50 | 7 | 996986 | 1138260 | - | 4961 | chromosome 7 | 0.133 | 6.01263 | 9.77662 | 0.0027184 | 0.018591 |
| NPM3 | 10 | 1E+08 | 1E+08 | - | 928 | nucleophosmin | -0.1122 | 6.60057 | 9.77542 | 0.0027199 | 0.018593 |
| TMEM147 | 19 | 3.6E+07 | 3.6E+07 | + | 1927 | transmembran | -0.1209 | 6.36502 | 9.77418 | 0.0027215 | 0.018595 |
| BCR | 22 | 2.3E+07 | 2.3E+07 | + | 12389 | BCR, RhoGEF a | -0.0915 | 7.65285 | 9.77218 | 0.002724 | 0.018603 |
| TPGS1 | 19 | 507497 | 519654 | + | 3642 | tubulin polyglu | 0.44027 | 2.46493 | 9.80921 | 0.0027349 | 0.018668 |
| BPTF | 17 | 6.8E+07 | 6.8E+07 | + | 15119 | bromodomain | -0.0813 | 8.5121 | 9.76234 | 0.0027366 | 0.018668 |
| SLC35C2 | 20 | 4.6E+07 | 4.6E+07 | - | 7708 | solute carrier f | -0.1518 | 5.6992 | 9.76166 | 0.0027374 | 0.018668 |
| ZRSR2 | X | 1.6E+07 | 1.6E+07 | + | 2077 | zinc finger CCC | -0.3153 | 3.31288 | 9.75789 | 0.0027422 | 0.018692 |
| ARHGEF5 | 7 | 1.4E+08 | 1.4E+08 | + | 5818 | Rho guanine n | -0.2435 | 4.09291 | 9.74868 | 0.0027541 | 0.018759 |
| DLG5 | 10 | 7.8E+07 | 7.8E+07 | - | 10500 | discs large MA | 0.10631 | 7.04872 | 9.74899 | 0.0027546 | 0.018759 |
| EZR | 6 | 1.6E+08 | 1.6E+08 | - | 3298 | ezrin [Source:H | -0.0815 | 8.80772 | 9.74738 | 0.0027567 | 0.018764 |
| UBE3C | 7 | 1.6E+08 | 1.6E+08 | + | 9215 | ubiquitin prote | 0.10224 | 7.01932 | 9.74427 | 0.0027598 | 0.018776 |
| C12orf45 | 12 | 1E+08 | 1.1E+08 | + | 25882 | chromosome 1 | -0.189 | 4.86068 | 9.73931 | 0.0027662 | 0.018811 |
| REEP2 | 5 | 1.4E+08 | 1.4E+08 | + | 3244 | receptor acces | -0.1608 | 5.49243 | 9.73598 | 0.0027705 | 0.01883 |
| SLC25A3 | 12 | 9.9E+07 | 9.9E+07 | + | 8896 | solute carrier f | -0.074 | 9.28667 | 9.73516 | 0.0027716 | 0.01883 |
| SETBP1 | 18 | 4.5E+07 | 4.5E+07 | + | 14025 | SET binding pr | -0.3764 | 2.89162 | 9.72999 | 0.0027783 | 0.01885 |
| KRT7 | 12 | 5.2E+07 | 5.2E+07 | + | 3999 | keratin 7 [Sour | -0.3906 | 2.63903 | 9.72999 | 0.0027783 | 0.01885 |
| CPSF4 | 7 | 9.9E+07 | 9.9E+07 | + | 4252 | cleavage and p | 0.12678 | 6.13383 | 9.72889 | 0.0027797 | 0.01885 |
| FAM160A | 4 | 1.5E+08 | 1.5E+08 | + | 11843 | family with seq | -0.1236 | 6.36051 | 9.7288 | 0.0027798 | 0.01885 |
| THSD4 | 15 | 7.1E+07 | 7.2E+07 | + | 15837 | thrombospond | 0.33124 | 3.12919 | 9.72621 | 0.0027832 | 0.018861 |
| PUM3 | 9 | 2720469 | 2844095 | - | 2573 | pumilio RNA b | -0.1414 | 5.92521 | 9.72556 | 0.002784 | 0.018861 |
| RPS8 | 1 | 4.5E+07 | 4.5E+07 | + | 2781 | ribosomal prot | 0.06919 | 10.0378 | 9.71962 | 0.0027918 | 0.018904 |
| SOX2 | 3 | 1.8E+08 | 1.8E+08 | + | 2512 | SRY-box 2 [Sou | -0.0894 | 8.46567 | 9.90649 | 0.0027948 | 0.018916 |
| TNFRSF8 | 1 | 1.2E+07 | 1.2E+07 | + | 3812 | TNF receptor s | -0.161 | 5.38305 | 9.71151 | 0.0028024 | 0.018956 |
| RAB1A | 2 | 6.5E+07 | 6.5E+07 | - | 3313 | RAB1A, memb | -0.1031 | 7.06577 | 9.7108 | 0.0028033 | 0.018956 |

| | | | | | | | | | | | |
|---|---|---|---|---|---|---|---|---|---|---|---|
| PMPCA | 9 | 1.4E+08 | 1.4E+08 | + | 6519 | peptidase, mit | 0.10749 | 6.95607 | 9.70619 | 0.0028094 | 0.018988 |
| SMAD1 | 4 | 1.5E+08 | 1.5E+08 | + | 7094 | SMAD family n | 0.20163 | 4.63763 | 9.69198 | 0.0028281 | 0.019105 |
| SH3BGRL | X | 8.1E+07 | 8.1E+07 | + | 2145 | SH3 domain bi | 0.13687 | 5.88026 | 9.68898 | 0.0028321 | 0.019123 |
| NECAB1 | 8 | 9.1E+07 | 9.1E+07 | + | 6202 | N-terminal EF- | -0.2971 | 3.58125 | 9.68739 | 0.0028342 | 0.019128 |
| RBM17 | 10 | 6089034 | 6117457 | + | 6372 | RNA binding m | -0.1065 | 6.88348 | 9.68571 | 0.0028364 | 0.019134 |
| MYADM | 19 | 5.4E+07 | 5.4E+07 | + | 4298 | myeloid associ | -0.1237 | 6.27697 | 9.67681 | 0.0028483 | 0.019205 |
| LIMS1 | 2 | 1.1E+08 | 1.1E+08 | + | 7771 | LIM zinc finger | 0.15082 | 5.59733 | 9.66072 | 0.0028698 | 0.019342 |
| CASK | X | 4.2E+07 | 4.2E+07 | - | 19939 | calcium/calmo | 0.13003 | 6.07901 | 9.65943 | 0.0028715 | 0.019344 |
| PDE8A | 15 | 8.5E+07 | 8.5E+07 | + | 8004 | phosphodieste | 0.33934 | 3.03038 | 9.64926 | 0.0028852 | 0.019427 |
| PBXIP1 | 1 | 1.5E+08 | 1.5E+08 | - | 3851 | PBX homeobox | 0.17772 | 5.34084 | 9.76645 | 0.0028946 | 0.019481 |
| EPHB6 | HSCHR7_2 | 1.4E+08 | 1.4E+08 | + | 5325 | EPH receptor E | -0.1934 | 5.01587 | 9.707 | 0.002904 | 0.019536 |
| UCK1 | 9 | 1.3E+08 | 1.3E+08 | - | 2684 | uridine-cytidin | 0.16308 | 5.33243 | 9.62977 | 0.0029117 | 0.019578 |
| SAMD14 | 17 | 5E+07 | 5E+07 | - | 7016 | sterile alpha m | 0.35182 | 2.90014 | 9.61666 | 0.0029297 | 0.01968 |
| ACSL1 | 4 | 1.8E+08 | 1.8E+08 | - | 6284 | acyl-CoA synth | -0.2525 | 3.97419 | 9.61506 | 0.0029319 | 0.019686 |
| XYLT2 | 17 | 5E+07 | 5E+07 | + | 5472 | xylosyltransfer | -0.1303 | 6.03554 | 9.61325 | 0.0029343 | 0.019693 |
| HTR3A | 11 | 1.1E+08 | 1.1E+08 | + | 2772 | 5-hydroxytrypt | -0.3046 | 3.39331 | 9.60015 | 0.0029524 | 0.019806 |
| TSPAN18 | 11 | 4.5E+07 | 4.5E+07 | + | 5871 | tetraspanin 18 | -0.1858 | 4.98152 | 9.59421 | 0.0029607 | 0.019847 |
| PRR12 | 19 | 5E+07 | 5E+07 | + | 7554 | proline rich 12 | -0.0966 | 7.23455 | 9.59373 | 0.0029613 | 0.019847 |
| VANGL2 | 1 | 1.6E+08 | 1.6E+08 | + | 5354 | VANGL planar | -0.1029 | 6.91142 | 9.5898 | 0.0029668 | 0.019865 |
| PLA2G6 | 22 | 3.8E+07 | 3.8E+07 | - | 6758 | phospholipase | 0.43993 | 2.31339 | 9.58975 | 0.0029669 | 0.019865 |
| IRF2 | 4 | 1.8E+08 | 1.8E+08 | - | 3070 | interferon regu | 0.2665 | 3.77153 | 9.58625 | 0.0029717 | 0.019889 |
| JMJD1C | 10 | 6.3E+07 | 6.4E+07 | - | 11162 | jumonji domai | 0.08632 | 7.97634 | 9.57208 | 0.0029916 | 0.020012 |
| ZNF559 | 19 | 9323772 | 9351162 | + | 6783 | zinc finger prot | -0.3165 | 3.92904 | 10.4504 | 0.0030019 | 0.020072 |
| TRIM21 | 11 | 4384897 | 4393702 | - | 1935 | tripartite motif | -0.2502 | 3.99579 | 9.55802 | 0.0030114 | 0.020126 |
| ABCG1 | 21 | 4.2E+07 | 4.2E+07 | + | 6121 | ATP binding ca | -0.3755 | 2.75575 | 9.5541 | 0.0030169 | 0.020152 |
| RAI1 | 17 | 1.8E+07 | 1.8E+07 | + | 9708 | retinoic acid in | 0.3626 | 2.90354 | 9.55325 | 0.0030181 | 0.020152 |
| SZT2 | 1 | 4.3E+07 | 4.3E+07 | + | 26354 | SZT2, KICSTOR | 0.23017 | 4.23239 | 9.55192 | 0.00302 | 0.020155 |
| ITGA7 | 12 | 5.6E+07 | 5.6E+07 | - | 6373 | integrin subun | 0.15188 | 5.69792 | 9.55058 | 0.0030294 | 0.020209 |
| WDHD1 | 14 | 5.5E+07 | 5.5E+07 | - | 6386 | WD repeat and | -0.1244 | 6.89729 | 10.1726 | 0.0030348 | 0.020222 |
| BUD13 | 11 | 1.2E+08 | 1.2E+08 | - | 2195 | BUD13 homolo | 0.17769 | 5.09845 | 9.5409 | 0.0030357 | 0.020222 |
| GRK6 | 5 | 1.8E+08 | 1.8E+08 | + | 4357 | G protein-coup | 0.11848 | 6.32053 | 9.53434 | 0.003045 | 0.020275 |
| RPUSD1 | 16 | 784974 | 788397 | - | 2769 | RNA pseudour | 0.16385 | 5.31391 | 9.53008 | 0.0030511 | 0.020301 |
| AC074143 | 16 | 6.7E+07 | 6.7E+07 | + | 2992 | nucleolar prot | 0.77553 | 0.7946 | 9.52838 | 0.0030536 | 0.020301 |
| L1CAM | X | 1.5E+08 | 1.5E+08 | - | 7270 | L1 cell adhesio | -0.1446 | 5.75859 | 9.52782 | 0.0030544 | 0.020301 |
| LTBP3 | 11 | 6.6E+07 | 6.6E+07 | - | 10298 | latent transfor | 0.24146 | 4.16524 | 9.52769 | 0.0030546 | 0.020301 |
| LDHA | 11 | 1.8E+07 | 1.8E+07 | + | 6038 | lactate dehydr | -0.0823 | 8.4852 | 9.52595 | 0.0030571 | 0.020308 |
| PCSK1N | X | 4.9E+07 | 4.9E+07 | - | 1338 | proprotein cor | -0.411 | 2.45421 | 9.52016 | 0.0030654 | 0.020349 |
| ADGRA3 | 4 | 2.2E+07 | 2.3E+07 | - | 9261 | adhesion G pro | 0.10141 | 6.98837 | 9.5189 | 0.0030672 | 0.020349 |
| LPIN1 | 2 | 1.2E+07 | 1.2E+07 | + | 11861 | lipin 1 [Source | 0.19394 | 4.74694 | 9.51879 | 0.0030674 | 0.020349 |
| SREBF2 | 22 | 4.2E+07 | 4.2E+07 | + | 7556 | sterol regulato | 0.07739 | 8.77741 | 9.51131 | 0.0030782 | 0.020411 |
| ATP6V1G1 | 9 | 1.1E+08 | 1.1E+08 | + | 1788 | ATPase H+ tran | -0.1186 | 6.5454 | 9.51341 | 0.0030947 | 0.020511 |
| VAMP8 | 2 | 8.6E+07 | 8.6E+07 | + | 895 | vesicle associa | -0.1945 | 4.75569 | 9.49614 | 0.0031002 | 0.020538 |
| C2orf76 | 2 | 1.2E+08 | 1.2E+08 | - | 1554 | chromosome 2 | -0.5844 | 1.58812 | 9.49437 | 0.0031028 | 0.020545 |

| | | | | | | | | | | | |
|---|---|---|---|---|---|---|---|---|---|---|---|
| KIAA1549 | 7 | 1.4E+08 | 1.4E+08 | - | 12427 | KIAA1549 [Sou | 0.15485 | 5.47493 | 9.48621 | 0.0031147 | 0.020615 |
| WBP1 | 2 | 7.4E+07 | 7.4E+07 | + | 2367 | WW domain b | 0.29857 | 3.38254 | 9.4829 | 0.0031195 | 0.020636 |
| PSMA3 | 14 | 5.8E+07 | 5.8E+07 | + | 3107 | proteasome su | -0.1118 | 6.94622 | 9.60234 | 0.0031208 | 0.020636 |
| STON2 | 14 | 8.1E+07 | 8.1E+07 | - | 7812 | stonin 2 [Sour | 0.26835 | 3.83849 | 9.46848 | 0.0031408 | 0.020759 |
| EPB41L5 | 2 | 1.2E+08 | 1.2E+08 | + | 9209 | erythrocyte me | -0.1543 | 5.48533 | 9.46555 | 0.0031451 | 0.020778 |
| GOT2 | 16 | 5.9E+07 | 5.9E+07 | - | 3751 | glutamic-oxalo | -0.0804 | 8.35213 | 9.44881 | 0.00317 | 0.020933 |
| LDLRAD4 | 18 | 1.3E+07 | 1.4E+07 | + | 16759 | low density lip | 0.70525 | 1.06777 | 9.4351 | 0.0031905 | 0.021058 |
| TRIM13 | 13 | 5E+07 | 5E+07 | + | 8096 | tripartite motif | 0.19047 | 4.80321 | 9.42852 | 0.0032004 | 0.021111 |
| MICALL1 | 22 | 3.8E+07 | 3.8E+07 | + | 5968 | MICAL like 1 [S | 0.11622 | 6.41412 | 9.42787 | 0.0032014 | 0.021111 |
| AP000866 | 11 | 1.2E+08 | 1.2E+08 | - | 3579 | novel transcrip | 0.59639 | 1.51364 | 9.42636 | 0.0032036 | 0.021116 |
| RET | 10 | 4.3E+07 | 4.3E+07 | + | 6869 | ret proto-onco | -0.2763 | 3.76319 | 9.42469 | 0.0032061 | 0.021123 |
| PPA2 | 4 | 1.1E+08 | 1.1E+08 | - | 4640 | pyrophosphata | 0.15754 | 5.39424 | 9.42142 | 0.0032111 | 0.021146 |
| TNK2 | 3 | 2E+08 | 2E+08 | - | 5153 | tyrosine kinase | 0.16026 | 5.39554 | 9.4175 | 0.003217 | 0.021175 |
| HAS2 | 8 | 1.2E+08 | 1.2E+08 | - | 4240 | hyaluronan syn | 0.49468 | 1.99772 | 9.4128 | 0.0032241 | 0.021212 |
| PLEKHH2 | 2 | 4.4E+07 | 4.4E+07 | + | 9848 | pleckstrin hom | 0.30838 | 3.29769 | 9.39482 | 0.0032516 | 0.021383 |
| AIG1 | 6 | 1.4E+08 | 1.4E+08 | + | 7677 | androgen indu | 0.19976 | 4.66464 | 9.37169 | 0.0032872 | 0.021594 |
| STUB1 | 16 | 680224 | 682870 | + | 2318 | STIP1 homolog | 0.1513 | 5.55028 | 9.37104 | 0.0032882 | 0.021594 |
| ELK1 | X | 4.8E+07 | 4.8E+07 | - | 3398 | ELK1, ETS trans | -0.136 | 5.8605 | 9.36689 | 0.0032946 | 0.021627 |
| LINC0113 | 1 | 1.5E+08 | 1.5E+08 | - | 7737 | long intergenic | -0.4778 | 2.03215 | 9.3644 | 0.0032985 | 0.021642 |
| GAMT | 19 | 1397026 | 1401570 | - | 2403 | guanidinoaceta | 0.27676 | 3.90386 | 9.53322 | 0.0033005 | 0.021645 |
| JMJD4 | 1 | 2.3E+08 | 2.3E+08 | - | 4315 | jumonji domai | 0.39145 | 2.53203 | 9.34855 | 0.0033232 | 0.021783 |
| BICD1 | 12 | 3.2E+07 | 3.2E+07 | + | 12868 | BICD cargo ada | -0.1388 | 5.93964 | 9.34776 | 0.0033245 | 0.021783 |
| FAM160B | 8 | 2.2E+07 | 2.2E+07 | + | 8300 | family with seq | 0.26311 | 3.76607 | 9.3435 | 0.0033312 | 0.021817 |
| KIAA1147 | HSCHR7_1 | 1.4E+08 | 1.4E+08 | - | 7369 | KIAA1147 [Sou | 0.11558 | 6.37051 | 9.3416 | 0.0033342 | 0.021824 |
| TTC17 | 11 | 4.3E+07 | 4.3E+07 | + | 11296 | tetratricopepti | 0.12678 | 6.05489 | 9.34083 | 0.0033354 | 0.021824 |
| SLC7A6 | 16 | 6.8E+07 | 6.8E+07 | + | 7588 | solute carrier f | 0.20281 | 4.58035 | 9.33631 | 0.0033425 | 0.021861 |
| RPL21P16 | 10 | 1.2E+08 | 1.2E+08 | - | 483 | ribosomal prot | -0.1705 | 5.1579 | 9.33097 | 0.0033509 | 0.021906 |
| NAPG | 18 | 1.1E+07 | 1.1E+07 | + | 5589 | NSF attachmer | -0.1962 | 4.77307 | 9.32909 | 0.0033539 | 0.021915 |
| TTC30A | 2 | 1.8E+08 | 1.8E+08 | - | 5744 | tetratricopepti | 0.53191 | 1.73334 | 9.32659 | 0.0033579 | 0.021931 |
| OBSL1 | 2 | 2.2E+08 | 2.2E+08 | - | 9716 | obscurin like 1 | 0.09094 | 7.76957 | 9.31551 | 0.0033755 | 0.022036 |
| ERGIC2 | 12 | 2.9E+07 | 2.9E+07 | - | 5826 | ERGIC and golg | -0.1996 | 4.68723 | 9.31079 | 0.003383 | 0.022075 |
| ADGRG6 | 6 | 1.4E+08 | 1.4E+08 | + | 8473 | adhesion G pro | 0.34495 | 2.99801 | 9.29228 | 0.0034127 | 0.022251 |
| DCUN1D5 | 11 | 1E+08 | 1E+08 | - | 13069 | defective in cu | -0.1095 | 6.62783 | 9.29206 | 0.003413 | 0.022251 |
| ZNF205 | 16 | 3112560 | 3120517 | + | 2330 | zinc finger prot | 0.199 | 4.64521 | 9.28784 | 0.0034198 | 0.022285 |
| ZNF430 | 19 | 2.1E+07 | 2.1E+07 | + | 6831 | zinc finger prot | -0.2909 | 3.58015 | 9.28159 | 0.0034299 | 0.022341 |
| GRK2 | 11 | 6.7E+07 | 6.7E+07 | + | 6602 | G protein-coup | 0.10812 | 6.63373 | 9.27689 | 0.0034376 | 0.022381 |
| ZNF28 | 19 | 5.3E+07 | 5.3E+07 | - | 5975 | zinc finger prot | -0.246 | 3.94645 | 9.27543 | 0.0034399 | 0.022386 |
| FCGBP | 19 | 4E+07 | 4E+07 | - | 12961 | Fc fragment of | -0.448 | 2.20016 | 9.2663 | 0.0034548 | 0.022473 |
| DANCR | 4 | 5.3E+07 | 5.3E+07 | + | 2285 | differentiation | -0.0965 | 7.35582 | 9.26229 | 0.0034613 | 0.022505 |
| PDP1 | 8 | 9.4E+07 | 9.4E+07 | + | 5604 | pyruvate dehy | 0.20211 | 4.78004 | 9.30543 | 0.0034733 | 0.022572 |
| DHX29 | 5 | 5.5E+07 | 5.5E+07 | - | 4896 | DExH-box helic | -0.1173 | 6.3904 | 9.25377 | 0.0034753 | 0.022575 |
| CLDN9 | 16 | 3012923 | 3014505 | + | 1583 | claudin 9 [Sour | -0.7027 | 1.029 | 9.25241 | 0.0034776 | 0.02258 |
| MET | 7 | 1.2E+08 | 1.2E+08 | + | 7233 | MET proto-onc | 0.21054 | 4.54407 | 9.24253 | 0.0034938 | 0.022675 |

| | | | | | | | | | | | |
|---|---|---|---|---|---|---|---|---|---|---|---|
| DERL2 | 17 | 5471254 | 5486811 | - | 6383 | derlin 2 [Sourc | -0.2035 | 4.74658 | 9.27582 | 0.0035065 | 0.022747 |
| ERV3-1 | 7 | 6.5E+07 | 6.5E+07 | - | 5027 | endogenous re | 0.2938 | 3.39141 | 9.23327 | 0.0035091 | 0.022754 |
| PCGF3 | 4 | 705748 | 770640 | + | 7425 | polycomb grou | 0.11478 | 6.44056 | 9.23194 | 0.0035114 | 0.022758 |
| CYP1B1 | 2 | 3.8E+07 | 3.8E+07 | - | 5985 | cytochrome P4 | -0.3868 | 2.66192 | 9.22913 | 0.003516 | 0.022778 |
| MVK | 12 | 1.1E+08 | 1.1E+08 | + | 6488 | mevalonate kir | -0.1977 | 4.77959 | 9.22151 | 0.0035287 | 0.02285 |
| CUTA | 6 | 3.3E+07 | 3.3E+07 | - | 1876 | cutA divalent c | -0.1065 | 6.85117 | 9.21159 | 0.0035453 | 0.022941 |
| ZNF518A | 10 | 9.6E+07 | 9.6E+07 | + | 10891 | zinc finger prot | -0.1386 | 5.91251 | 9.21123 | 0.0035459 | 0.022941 |
| ENY2 | 8 | 1.1E+08 | 1.1E+08 | + | 6784 | ENY2, transcrip | -0.1248 | 6.21535 | 9.20582 | 0.003555 | 0.022981 |
| WDFY3 | 4 | 8.5E+07 | 8.5E+07 | - | 16856 | WD repeat and | 0.10975 | 6.57142 | 9.20563 | 0.0035553 | 0.022981 |
| MAP3K20 | 2 | 1.7E+08 | 1.7E+08 | + | 13804 | mitogen-activa | 0.16475 | 5.36682 | 9.2024 | 0.0035608 | 0.023002 |
| AMBP | 9 | 1.1E+08 | 1.1E+08 | - | 1897 | alpha-1-microg | 0.58575 | 1.49986 | 9.20182 | 0.0035617 | 0.023002 |
| RANBP9 | 6 | 1.4E+07 | 1.4E+07 | - | 3618 | RAN binding pr | 0.13403 | 5.90911 | 9.19992 | 0.0035649 | 0.023012 |
| SUV39H1 | X | 4.9E+07 | 4.9E+07 | + | 3368 | suppressor of v | -0.2191 | 4.39165 | 9.18939 | 0.0035827 | 0.023116 |
| OTX2 | 14 | 5.7E+07 | 5.7E+07 | - | 3272 | orthodenticle h | -0.1757 | 5.25213 | 9.25334 | 0.0035854 | 0.023124 |
| JAK3 | 19 | 1.8E+07 | 1.8E+07 | - | 6813 | Janus kinase 3 | -0.346 | 3.0257 | 9.18175 | 0.0035957 | 0.023179 |
| ZNF616 | 19 | 5.2E+07 | 5.2E+07 | - | 4907 | zinc finger prot | -0.1559 | 5.444 | 9.18065 | 0.0035976 | 0.023181 |
| MPP6 | 7 | 2.5E+07 | 2.5E+07 | + | 9369 | membrane pal | 0.11194 | 6.62382 | 9.17948 | 0.0035996 | 0.023184 |
| EVA1B | 1 | 3.6E+07 | 3.6E+07 | - | 1211 | eva-1 homolog | 0.24641 | 4.05044 | 9.17716 | 0.0036035 | 0.023199 |
| PARD3B | 2 | 2E+08 | 2.1E+08 | + | 10700 | par-3 family ce | 0.2857 | 3.48263 | 9.17515 | 0.003607 | 0.02321 |
| ARHGAP1 | SCHR15_4 | 3.3E+07 | 3.3E+07 | + | 6406 | Rho GTPase ac | -0.1119 | 6.585 | 9.16955 | 0.0036165 | 0.023261 |
| F11R | 1 | 1.6E+08 | 1.6E+08 | - | 5061 | F11 receptor [S | -0.0919 | 7.59897 | 9.16814 | 0.003619 | 0.023267 |
| RAB11FIP | 17 | 3.1E+07 | 3.2E+07 | + | 15326 | RAB11 family i | -0.1035 | 6.81938 | 9.16687 | 0.0036211 | 0.02327 |
| MTMR9LP | 1 | 3.2E+07 | 3.2E+07 | - | 2961 | myotubularin r | 0.32869 | 3.10632 | 9.16199 | 0.0036295 | 0.023314 |
| SIN3B | 19 | 1.7E+07 | 1.7E+07 | + | 8777 | SIN3 transcript | -0.1023 | 6.92331 | 9.16056 | 0.003632 | 0.023319 |
| DMKN | 19 | 3.5E+07 | 3.6E+07 | - | 7444 | dermokine [So | -0.1054 | 6.74268 | 9.15624 | 0.0036394 | 0.023353 |
| FUT1 | 19 | 4.9E+07 | 4.9E+07 | - | 4343 | fucosyltransfer | -0.2998 | 3.34773 | 9.15559 | 0.0036405 | 0.023353 |
| GFPT1 | 2 | 6.9E+07 | 6.9E+07 | - | 8866 | glutamine--fru | -0.0953 | 7.20899 | 9.12761 | 0.0036891 | 0.023654 |
| NSRP1 | 17 | 3E+07 | 3E+07 | + | 4806 | nuclear speckle | -0.2106 | 4.51732 | 9.12646 | 0.0036911 | 0.023656 |
| TRPS1 | 8 | 1.2E+08 | 1.2E+08 | - | 11072 | transcriptional | 0.33187 | 3.14969 | 9.12003 | 0.0037024 | 0.023708 |
| NCDN | 1 | 3.6E+07 | 3.6E+07 | + | 4917 | neurochondrin | 0.14861 | 5.60049 | 9.11999 | 0.0037025 | 0.023708 |
| MT-CO3 | MT | 9207 | 9990 | + | 784 | mitochondriall | -0.0595 | 12.2082 | 9.10859 | 0.0037225 | 0.023826 |
| SNRPG | 2 | 7E+07 | 7E+07 | - | 1764 | small nuclear r | -0.1062 | 6.67864 | 9.10739 | 0.0037246 | 0.023829 |
| OTUD3 | 1 | 2E+07 | 2E+07 | + | 6885 | OTU deubiquit | 0.19126 | 4.71865 | 9.1032 | 0.003732 | 0.02386 |
| EFNA2 | 19 | 1285873 | 1301431 | + | 2424 | ephrin A2 [Sou | 0.31821 | 3.14535 | 9.10214 | 0.0037339 | 0.02386 |
| CD40 | 20 | 4.6E+07 | 4.6E+07 | + | 2527 | CD40 molecule | -0.266 | 3.70726 | 9.10127 | 0.0037354 | 0.02386 |
| TFAM | 10 | 5.8E+07 | 5.8E+07 | + | 5282 | transcription fa | -0.1034 | 7.06911 | 9.1325 | 0.0037362 | 0.02386 |
| METTL8 | 2 | 1.7E+08 | 1.7E+08 | - | 10074 | methyltransfer | -0.1348 | 5.87544 | 9.09332 | 0.0037496 | 0.023935 |
| COPS3 | 17 | 1.7E+07 | 1.7E+07 | - | 2972 | COP9 signaloso | -0.1079 | 6.64259 | 9.08482 | 0.0037647 | 0.024021 |
| TRIM44 | 11 | 3.6E+07 | 3.6E+07 | + | 13107 | tripartite motif | 0.10187 | 6.8647 | 9.08016 | 0.003773 | 0.024063 |
| AKNA | 9 | 1.1E+08 | 1.1E+08 | - | 10601 | AT-hook transc | 0.15207 | 5.52076 | 9.07202 | 0.0037876 | 0.024144 |
| IFI6 | 1 | 2.8E+07 | 2.8E+07 | - | 842 | interferon alph | 0.50326 | 1.91705 | 9.07124 | 0.003789 | 0.024144 |
| RPLP0 | 12 | 1.2E+08 | 1.2E+08 | - | 3988 | ribosomal prot | 0.06381 | 10.6616 | 9.07013 | 0.003791 | 0.024146 |
| ZNF845 | 19 | 5.3E+07 | 5.3E+07 | + | 6453 | zinc finger prot | -0.2318 | 4.30626 | 9.10891 | 0.0037927 | 0.024146 |

| | | | | | | | | | | | |
|---|---|---|---|---|---|---|---|---|---|---|---|
| ANKRD52 | 12 | 5.6E+07 | 5.6E+07 | - | 8813 | ankyrin repeat | -0.1085 | 6.81433 | 9.07665 | 0.0038016 | 0.024192 |
| CMBL | 5 | 1E+07 | 1E+07 | - | 4882 | carboxymethy | 0.18333 | 4.93037 | 9.05915 | 0.0038108 | 0.02424 |
| DTNB | 2 | 2.5E+07 | 2.6E+07 | - | 7146 | dystrobrevin b | 0.16006 | 5.27669 | 9.05192 | 0.0038239 | 0.024313 |
| STXBP6 | 14 | 2.5E+07 | 2.5E+07 | - | 8067 | syntaxin bindir | -0.3125 | 3.20552 | 9.04727 | 0.0038324 | 0.024355 |
| SAPCD2 | 9 | 1.4E+08 | 1.4E+08 | - | 3814 | suppressor AP( | 0.1102 | 6.56699 | 9.0452 | 0.0038362 | 0.024358 |
| LHFPL4 | 3 | 9498361 | 9553822 | - | 5349 | LHFPL tetraspa | 0.19005 | 4.7077 | 9.04508 | 0.0038364 | 0.024358 |
| FAM214A | 15 | 5.3E+07 | 5.3E+07 | - | 9832 | family with sec | -0.2761 | 3.59559 | 9.04422 | 0.0038379 | 0.024358 |
| PTPN23 | 3 | 4.7E+07 | 4.7E+07 | + | 5975 | protein tyrosin | -0.108 | 6.63794 | 9.02726 | 0.003869 | 0.024539 |
| NUTM2B- | 10 | 8E+07 | 8E+07 | - | 14755 | NUTM2B antis | -0.2082 | 4.51668 | 9.02679 | 0.0038699 | 0.024539 |
| DARS2 | 1 | 1.7E+08 | 1.7E+08 | + | 7438 | aspartyl-tRNA | 0.14036 | 5.69903 | 9.02179 | 0.003879 | 0.024587 |
| KLHL23 | 2 | 1.7E+08 | 1.7E+08 | + | 5513 | kelch like fami | 0.12023 | 6.45915 | 9.08281 | 0.0038833 | 0.024603 |
| TMEM50E | 21 | 3.3E+07 | 3.3E+07 | - | 4458 | transmembran | -0.2506 | 3.97189 | 9.0094 | 0.003902 | 0.02471 |
| U2SURP | 3 | 1.4E+08 | 1.4E+08 | + | 11118 | U2 snRNP asso | 0.08983 | 7.76598 | 9.00769 | 0.0039051 | 0.024718 |
| NDUFAF8 | 17 | 8.1E+07 | 8.1E+07 | + | 1339 | NADH:ubiquin | 0.24009 | 4.39163 | 9.3268 | 0.0039067 | 0.024718 |
| DGAT2 | 11 | 7.6E+07 | 7.6E+07 | + | 7904 | diacylglycerol ( | -0.5495 | 1.63519 | 8.99241 | 0.0039336 | 0.024877 |
| PHGDH | 1 | 1.2E+08 | 1.2E+08 | + | 9217 | phosphoglycer | -0.0715 | 9.33506 | 8.99148 | 0.0039353 | 0.024877 |
| NUDT15 | 13 | 4.8E+07 | 4.8E+07 | + | 1938 | nudix hydrolas | -0.1158 | 6.33095 | 8.98854 | 0.0039408 | 0.024901 |
| AVPI1 | 10 | 9.8E+07 | 9.8E+07 | - | 1375 | arginine vasop | -0.3176 | 3.16695 | 8.98716 | 0.0039434 | 0.024903 |
| PHF14 | 7 | 1.1E+07 | 1.1E+07 | + | 14051 | PHD finger pro | -0.1084 | 6.83686 | 9.02544 | 0.0039445 | 0.024903 |
| TARBP1 | 1 | 2.3E+08 | 2.3E+08 | - | 7819 | TAR (HIV-1) RN | 0.15859 | 5.32771 | 8.98388 | 0.0039496 | 0.024924 |
| LRRC49 | 15 | 7.1E+07 | 7.1E+07 | + | 9991 | leucine rich re | -0.317 | 3.21543 | 8.96241 | 0.0039901 | 0.025168 |
| FGFR2 | 10 | 1.2E+08 | 1.2E+08 | - | 7578 | fibroblast grow | -0.0999 | 6.95353 | 8.95323 | 0.0040076 | 0.025267 |
| CHRDL1 | X | 1.1E+08 | 1.1E+08 | - | 3920 | chordin like 1 [ | 0.20364 | 4.5271 | 8.95126 | 0.0040113 | 0.02528 |
| CLDN23 | 8 | 8701938 | 8704106 | + | 2169 | claudin 23 [Sou | -0.46 | 2.06673 | 8.94773 | 0.0040181 | 0.025311 |
| INTS6 | 13 | 5.1E+07 | 5.1E+07 | - | 25509 | integrator com | 0.15753 | 5.3636 | 8.94397 | 0.0040252 | 0.025335 |
| ATP6V0E1 | 5 | 1.7E+08 | 1.7E+08 | + | 1609 | ATPase H+ trar | -0.1412 | 5.7028 | 8.9409 | 0.0040311 | 0.025347 |
| DTD1 | 20 | 1.9E+07 | 1.9E+07 | + | 6465 | D-tyrosyl-tRNA | -0.1564 | 5.37681 | 8.94032 | 0.0040323 | 0.025347 |
| LAYN | 11 | 1.1E+08 | 1.1E+08 | + | 3144 | layilin [Source: | -0.4066 | 2.40349 | 8.94021 | 0.0040324 | 0.025347 |
| CHDH | 3 | 5.4E+07 | 5.4E+07 | - | 7787 | choline dehydr | 0.23319 | 4.09531 | 8.93691 | 0.0040388 | 0.025375 |
| NLGN4Y | Y | 1.5E+07 | 1.5E+07 | + | 8589 | neuroligin 4 Y- | 0.1272 | 6.05447 | 8.92665 | 0.0040586 | 0.025488 |
| RPARP-AS | 10 | 1E+08 | 1E+08 | + | 5660 | RPARP antisen | -0.1947 | 4.67332 | 8.92562 | 0.0040606 | 0.02549 |
| AC107871 | 15 | 6.8E+07 | 6.8E+07 | - | 7276 | novel protein | 0.46664 | 2.01736 | 8.92053 | 0.0040704 | 0.025541 |
| COMMD4 | 15 | 7.5E+07 | 7.5E+07 | + | 6779 | COMM domair | 0.12569 | 6.06411 | 8.91796 | 0.0040754 | 0.02555 |
| COQ8B | 19 | 4.1E+07 | 4.1E+07 | - | 5487 | coenzyme Q8E | -0.1459 | 5.57318 | 8.91787 | 0.0040756 | 0.02555 |
| AL390719 | 1 | 1059708 | 1069355 | + | 2203 | protein tyrosin | 0.66444 | 1.13466 | 8.91526 | 0.0040806 | 0.025565 |
| ELOVL5 | 6 | 5.3E+07 | 5.3E+07 | - | 3813 | ELOVL fatty ac | 0.08823 | 7.51915 | 8.91483 | 0.0040815 | 0.025565 |
| NDUFB11 | X | 4.7E+07 | 4.7E+07 | - | 1418 | NADH:ubiquin | -0.1185 | 6.25101 | 8.91256 | 0.0040859 | 0.025582 |
| MACROD1 | 11 | 6.4E+07 | 6.4E+07 | - | 2096 | MACRO domai | 0.22458 | 4.16171 | 8.90528 | 0.0041001 | 0.025659 |
| HRK | 12 | 1.2E+08 | 1.2E+08 | - | 6853 | harakiri, BCL2 i | -0.4832 | 1.92986 | 8.89778 | 0.0041148 | 0.025737 |
| AC009005 | 19 | 567210 | 572228 | - | 865 | novel transcrip | 0.51168 | 1.74125 | 8.8971 | 0.0041161 | 0.025737 |
| MAPRE1 | 20 | 3.3E+07 | 3.3E+07 | + | 2562 | microtubule as | -0.084 | 7.91449 | 8.8867 | 0.0041365 | 0.025842 |
| DCUN1D4 | 4 | 5.2E+07 | 5.2E+07 | + | 7625 | defective in cu | -0.1581 | 5.36299 | 8.88386 | 0.0041421 | 0.025866 |
| EML3 | 11 | 6.3E+07 | 6.3E+07 | - | 5490 | EMAP like 3 [S | 0.20484 | 4.42718 | 8.88026 | 0.0041492 | 0.025894 |

| | | | | | | | | | | | |
|---|---|---|---|---|---|---|---|---|---|---|---|
| ACSS2 | 20 | 3.5E+07 | 3.5E+07 | + | 5248 | acyl-CoA synth | -0.1636 | 5.19637 | 8.87981 | 0.0041501 | 0.025894 |
| RPRD1A | 18 | 3.6E+07 | 3.6E+07 | - | 8442 | regulation of n | -0.091 | 7.38958 | 8.87335 | 0.0041629 | 0.025962 |
| TTN | 2 | 1.8E+08 | 1.8E+08 | - | 118976 | titin [Source:H | -0.1515 | 5.44908 | 8.86686 | 0.0041758 | 0.026031 |
| RPL12 | 9 | 1.3E+08 | 1.3E+08 | - | 2453 | ribosomal prot | 0.07087 | 9.38764 | 8.85694 | 0.0041956 | 0.026138 |
| PTP4A2 | 1 | 3.2E+07 | 3.2E+07 | - | 6708 | protein tyrosin | -0.0882 | 7.542 | 8.85651 | 0.0041965 | 0.026138 |
| PIGV | 1 | 2.7E+07 | 2.7E+07 | + | 3382 | phosphatidylin | 0.28535 | 3.4437 | 8.85244 | 0.0042046 | 0.026172 |
| NCS1 | 9 | 1.3E+08 | 1.3E+08 | + | 5462 | neuronal calciu | -0.123 | 6.12037 | 8.85193 | 0.0042057 | 0.026172 |
| PCDHGA1 | 5 | 1.4E+08 | 1.4E+08 | + | 5810 | protocadherin | 0.36238 | 2.72371 | 8.84811 | 0.0042133 | 0.026208 |
| ZNF710 | 15 | 9E+07 | 9E+07 | + | 5004 | zinc finger prot | 0.16348 | 5.20092 | 8.83671 | 0.0042363 | 0.02634 |
| ADAMTS1 | 5 | 5140330 | 5320304 | + | 5863 | ADAM metallo | -0.2412 | 4.0158 | 8.83364 | 0.0042425 | 0.026367 |
| GPR161 | 1 | 1.7E+08 | 1.7E+08 | - | 11224 | G protein-coup | 0.19074 | 4.64798 | 8.83174 | 0.0042464 | 0.02638 |
| BRK1 | 3 | 1E+07 | 1E+07 | + | 1150 | BRICK1, SCAR/ | -0.1037 | 6.77978 | 8.82698 | 0.004256 | 0.026427 |
| TBC1D25 | X | 4.9E+07 | 4.9E+07 | + | 3966 | TBC1 domain f | -0.3119 | 3.44941 | 8.9464 | 0.0042576 | 0.026427 |
| AP4M1 | 7 | 1E+08 | 1E+08 | + | 4722 | adaptor relate | 0.2623 | 3.7504 | 8.82512 | 0.0042598 | 0.026429 |
| TEX261 | 2 | 7.1E+07 | 7.1E+07 | - | 4184 | testis expresse | -0.1724 | 5.1739 | 8.82953 | 0.004265 | 0.02645 |
| SUMO1 | 2 | 2E+08 | 2E+08 | - | 2164 | small ubiquitin | -0.0869 | 7.64157 | 8.81903 | 0.0042722 | 0.026483 |
| MYLIP | 6 | 1.6E+07 | 1.6E+07 | + | 3072 | myosin regulat | 0.16927 | 5.10912 | 8.81797 | 0.0042743 | 0.026485 |
| BGN | X | 1.5E+08 | 1.5E+08 | + | 2404 | biglycan [Sourc | 0.42308 | 2.2865 | 8.81263 | 0.0042852 | 0.026541 |
| MST1R | 3 | 5E+07 | 5E+07 | - | 5252 | macrophage st | 0.47201 | 2.00431 | 8.79645 | 0.0043185 | 0.026735 |
| RIMKLB | 12 | 8681600 | 8783095 | + | 8597 | ribosomal mod | 0.11567 | 6.4932 | 8.81335 | 0.0043255 | 0.026767 |
| TRMT2B | X | 1E+08 | 1E+08 | - | 4055 | tRNA methyltr | -0.2128 | 4.37127 | 8.79189 | 0.0043279 | 0.02677 |
| CALHM5 | 6 | 1.2E+08 | 1.2E+08 | + | 9787 | calcium homed | 0.87119 | 0.38961 | 8.78539 | 0.0043413 | 0.02684 |
| STYX | 14 | 5.3E+07 | 5.3E+07 | + | 4983 | serine/threoni | -0.1527 | 5.49293 | 8.78413 | 0.0043439 | 0.02684 |
| SNHG4 | 5 | 1.4E+08 | 1.4E+08 | + | 4469 | small nucleolar | 0.10315 | 6.75968 | 8.7831 | 0.0043461 | 0.02684 |
| EPHX3 | 19 | 1.5E+07 | 1.5E+07 | - | 2050 | epoxide hydro | -0.239 | 4.17961 | 8.84163 | 0.0043485 | 0.02684 |
| SPCS3 | 4 | 1.8E+08 | 1.8E+08 | + | 6829 | signal peptidas | 0.10491 | 6.73882 | 8.78183 | 0.0043487 | 0.02684 |
| SLC16A9 | 10 | 6E+07 | 6E+07 | - | 4429 | solute carrier f | 0.29713 | 3.32921 | 8.78107 | 0.0043503 | 0.02684 |
| GNB1 | 1 | 1785285 | 1891117 | - | 6417 | G protein subu | 0.07472 | 9.08329 | 8.78016 | 0.0043522 | 0.02684 |
| KDELR2 | 7 | 6445953 | 6484190 | - | 3930 | KDEL endoplas | -0.1088 | 6.49702 | 8.7688 | 0.0043759 | 0.026974 |
| FNDC10 | 1 | 1598012 | 1600135 | - | 2124 | fibronectin typ | 0.24101 | 4.08378 | 8.78128 | 0.0043807 | 0.026992 |
| UBB | 17 | 1.6E+07 | 1.6E+07 | + | 1621 | ubiquitin B [So | -0.0776 | 8.42065 | 8.76437 | 0.0043852 | 0.027008 |
| LINC0088 | 3 | 1.9E+08 | 1.9E+08 | + | 2922 | long intergenic | 0.88434 | 0.36946 | 8.76264 | 0.0043888 | 0.027019 |
| ASCC3 | 6 | 1E+08 | 1E+08 | - | 11731 | activating signa | 0.10316 | 7.86807 | 9.46234 | 0.0043993 | 0.027072 |
| KLHDC2 | 14 | 5E+07 | 5E+07 | + | 6319 | kelch domain c | 0.15337 | 5.40092 | 8.75002 | 0.0044153 | 0.027159 |
| PFDN4 | 20 | 5.4E+07 | 5.4E+07 | + | 1743 | prefoldin subu | -0.1333 | 5.81484 | 8.7476 | 0.0044204 | 0.027179 |
| SMIM4 | 3 | 5.3E+07 | 5.3E+07 | + | 2481 | small integral r | -0.2822 | 3.50664 | 8.74512 | 0.0044257 | 0.0272 |
| FMNL3 | 12 | 5E+07 | 5E+07 | - | 13415 | formin like 3 [S | 0.14567 | 5.59336 | 8.74287 | 0.0044304 | 0.027217 |
| LARP7 | 4 | 1.1E+08 | 1.1E+08 | + | 3996 | La ribonucleop | -0.1228 | 7.20506 | 9.76272 | 0.0044372 | 0.027247 |
| SLC5A6 | 2 | 2.7E+07 | 2.7E+07 | - | 5588 | solute carrier f | -0.0931 | 7.30599 | 8.73016 | 0.0044575 | 0.027351 |
| AUTS2 | 7 | 7E+07 | 7.1E+07 | + | 24197 | AUTS2, activat | -0.1013 | 7.08986 | 8.76692 | 0.0044579 | 0.027351 |
| ZDBF2 | 2 | 2.1E+08 | 2.1E+08 | + | 10901 | zinc finger DBF | 0.19173 | 4.64282 | 8.72523 | 0.004468 | 0.027401 |
| SF1 | 11 | 6.5E+07 | 6.5E+07 | - | 5972 | splicing factor | -0.0789 | 8.32945 | 8.72299 | 0.0044728 | 0.027419 |
| WRN | 8 | 3.1E+07 | 3.1E+07 | + | 8000 | Werner syndro | 0.12076 | 6.18595 | 8.71703 | 0.0044855 | 0.027485 |

| | | | | | | | | | | | |
|---|---|---|---|---|---|---|---|---|---|---|---|
| WASH8P | 12 | 14522 | 32015 | - | 1817 | WAS protein fa | 0.36537 | 2.74929 | 8.71465 | 0.0044906 | 0.027497 |
| CADPS | 3 | 6.2E+07 | 6.3E+07 | - | 14665 | calcium depen | -0.2341 | 4.07712 | 8.71432 | 0.0044913 | 0.027497 |
| EGLN1 | 1 | 2.3E+08 | 2.3E+08 | - | 7265 | egl-9 family hy | 0.26482 | 3.81887 | 8.74675 | 0.0045008 | 0.027535 |
| RLF | 1 | 4E+07 | 4E+07 | + | 6232 | rearranged L-n | 0.15379 | 5.33726 | 8.70971 | 0.0045013 | 0.027535 |
| PLCE1 | 10 | 9.4E+07 | 9.4E+07 | + | 12930 | phospholipase | -0.4459 | 2.20606 | 8.70503 | 0.0045113 | 0.027579 |
| ARHGAP1 | 10 | 9.7E+07 | 9.7E+07 | - | 6167 | Rho GTPase ac | -0.1254 | 6.06462 | 8.70437 | 0.0045128 | 0.027579 |
| LEFTY1 | 1 | 2.3E+08 | 2.3E+08 | - | 1891 | left-right deter | 1.13063 | 5.8001 | 45.9278 | 0.0045143 | 0.027579 |
| UXS1 | 2 | 1.1E+08 | 1.1E+08 | - | 7023 | UDP-glucurona | -0.1256 | 6.04118 | 8.70124 | 0.0045195 | 0.027589 |
| TSPAN14 | 10 | 8E+07 | 8.1E+07 | + | 17889 | tetraspanin 14 | -0.1293 | 6.00977 | 8.70092 | 0.0045202 | 0.027589 |
| SMAD4 | 18 | 5.1E+07 | 5.1E+07 | + | 13123 | SMAD family n | -0.1179 | 6.38521 | 8.70992 | 0.0045216 | 0.027589 |
| CAMSAP1 | 9 | 1.4E+08 | 1.4E+08 | - | 8960 | calmodulin reg | 0.10817 | 6.52326 | 8.69081 | 0.0045421 | 0.027689 |
| TCTEX1D2 | 3 | 2E+08 | 2E+08 | - | 1246 | Tctex1 domain | 0.69932 | 0.94433 | 8.69011 | 0.0045437 | 0.027689 |
| RAP1B | 12 | 6.9E+07 | 6.9E+07 | + | 16926 | RAP1B, membe | -0.1465 | 5.56621 | 8.69003 | 0.0045439 | 0.027689 |
| LEFTY2 | 1 | 2.3E+08 | 2.3E+08 | - | 2286 | left-right deter | 1.52837 | 5.23447 | 46.459 | 0.0045498 | 0.027712 |
| TAZ | X | 1.5E+08 | 1.5E+08 | + | 5977 | tafazzin [Sourc | 0.20172 | 4.44477 | 8.68385 | 0.0045573 | 0.027736 |
| ZNF682 | 19 | 2E+07 | 2E+07 | - | 4967 | zinc finger pro | -0.2174 | 4.28959 | 8.68272 | 0.0045598 | 0.027739 |
| EME1 | 17 | 5E+07 | 5E+07 | + | 3294 | essential meio | 0.27443 | 3.75431 | 8.75823 | 0.0045683 | 0.027779 |
| GSTK1 | 7 | 1.4E+08 | 1.4E+08 | + | 4214 | glutathione S-t | 0.19701 | 4.51907 | 8.67661 | 0.0045731 | 0.027797 |
| PCYT1B | X | 2.5E+07 | 2.5E+07 | - | 5555 | phosphate cyti | -0.1651 | 5.13083 | 8.6742 | 0.0045784 | 0.027817 |
| CS | 12 | 5.6E+07 | 5.6E+07 | - | 6401 | citrate synthas | -0.0842 | 7.82516 | 8.66704 | 0.0045941 | 0.0279 |
| GALNT11 | 7 | 1.5E+08 | 1.5E+08 | + | 8288 | polypeptide N- | 0.15575 | 5.29957 | 8.66625 | 0.0045959 | 0.0279 |
| RNF216P1 | 7 | 4973988 | 5040675 | + | 5510 | ring finger pro | -0.194 | 4.66572 | 8.66396 | 0.0046009 | 0.027918 |
| GNAI1 | 7 | 8E+07 | 8E+07 | + | 24192 | G protein subu | 0.20379 | 4.42365 | 8.66129 | 0.0046068 | 0.027942 |
| RTN4RL2 | 11 | 5.7E+07 | 5.7E+07 | + | 2408 | reticulon 4 rec | -0.1972 | 4.62095 | 8.66014 | 0.0046093 | 0.027946 |
| PLD3 | 19 | 4E+07 | 4E+07 | + | 5186 | phospholipase | -0.0846 | 7.87447 | 8.65669 | 0.004617 | 0.02798 |
| PTPN5 | 11 | 1.9E+07 | 1.9E+07 | - | 5497 | protein tyrosin | -0.3213 | 3.16693 | 8.65316 | 0.0046248 | 0.028016 |
| SH3D21 | 1 | 3.6E+07 | 3.6E+07 | + | 4474 | SH3 domain co | 0.38692 | 2.53185 | 8.65104 | 0.0046295 | 0.028032 |
| TDRP | 8 | 489792 | 545781 | - | 3613 | testis developr | -0.107 | 6.59075 | 8.64576 | 0.0046412 | 0.028083 |
| GTF2H1 | 11 | 1.8E+07 | 1.8E+07 | + | 6940 | general transc | -0.1726 | 5.10063 | 8.64553 | 0.0046417 | 0.028083 |
| UBE2E1 | 3 | 2.4E+07 | 2.4E+07 | + | 3074 | ubiquitin conju | 0.10364 | 6.76752 | 8.6362 | 0.0046625 | 0.028197 |
| SLC39A10 | 2 | 2E+08 | 2E+08 | + | 6917 | solute carrier f | -0.0901 | 7.71411 | 8.67008 | 0.0046674 | 0.02821 |
| GCNT2 | 6 | 1E+07 | 1.1E+07 | + | 9072 | glucosaminyl ( | -0.1083 | 6.56396 | 8.6335 | 0.0046685 | 0.02821 |
| SLC35A3 | 1 | 1E+08 | 1E+08 | + | 18235 | solute carrier f | -0.2277 | 4.18731 | 8.62953 | 0.0046775 | 0.028251 |
| MUS81 | 11 | 6.6E+07 | 6.6E+07 | + | 5448 | MUS81 structu | 0.19333 | 4.57122 | 8.62843 | 0.0046799 | 0.028254 |
| ELF3 | 1 | 2E+08 | 2E+08 | + | 7186 | E74 like ETS tra | -0.3036 | 3.25831 | 8.61641 | 0.0047069 | 0.028394 |
| NALT1 | 9 | 1.4E+08 | 1.4E+08 | + | 1088 | NOTCH1 assoc | 0.89604 | 0.40932 | 8.60998 | 0.0047215 | 0.02847 |
| INHBA | 7 | 4.2E+07 | 4.2E+07 | - | 8284 | inhibin subunit | -0.9272 | 0.32799 | 8.60514 | 0.0047325 | 0.028512 |
| SLX4IP | 20 | 1E+07 | 1.1E+07 | + | 6486 | SLX4 interactin | -0.309 | 3.24478 | 8.60509 | 0.0047326 | 0.028512 |
| THAP2 | 12 | 7.2E+07 | 7.2E+07 | + | 5240 | THAP domain c | 0.35219 | 2.7609 | 8.59449 | 0.0047567 | 0.028646 |
| KIAA1211 | 4 | 5.6E+07 | 5.6E+07 | + | 6745 | KIAA1211 [Sou | 0.16287 | 5.28018 | 8.59214 | 0.0047621 | 0.028666 |
| TCERG1 | 5 | 1.5E+08 | 1.5E+08 | + | 8714 | transcription e | 0.08892 | 7.41694 | 8.58567 | 0.0047769 | 0.028743 |
| AC005224 | 17 | 1.4E+07 | 1.4E+07 | + | 1652 | novel transcrip | -0.4323 | 2.24475 | 8.58332 | 0.0047823 | 0.028764 |
| VPS39 | 15 | 4.2E+07 | 4.2E+07 | - | 8354 | VPS39, HOPS c | 0.11834 | 6.2173 | 8.57985 | 0.0047902 | 0.028796 |

| | | | | | | | | | | | |
|---|---|---|---|---|---|---|---|---|---|---|---|
| FGD1 | X | 5.4E+07 | 5.4E+07 | - | 4343 | FYVE, RhoGEF | -0.1509 | 5.41793 | 8.57789 | 0.0047947 | 0.028802 |
| PLEKHJ1 | 19 | 2230084 | 2237704 | - | 4655 | pleckstrin hom | 0.09349 | 7.13706 | 8.57638 | 0.0047982 | 0.028811 |
| SOCS4 | 14 | 5.5E+07 | 5.5E+07 | + | 7044 | suppressor of | -0.1634 | 5.37652 | 8.63216 | 0.0048025 | 0.028825 |
| CD2AP | 6 | 4.7E+07 | 4.8E+07 | + | 6751 | CD2 associated | -0.0918 | 7.38105 | 8.56486 | 0.0048248 | 0.028947 |
| GBE1 | 3 | 8.1E+07 | 8.2E+07 | - | 4651 | 1,4-alpha-gluca | 0.22705 | 4.08288 | 8.54968 | 0.0048601 | 0.029142 |
| CDH3 | 16 | 6.9E+07 | 6.9E+07 | + | 6076 | cadherin 3 [So | -0.0816 | 8.0523 | 8.54914 | 0.0048614 | 0.029142 |
| ZDHHC14 | 6 | 1.6E+08 | 1.6E+08 | + | 11132 | zinc finger DHH | 0.30337 | 3.19606 | 8.54571 | 0.0048694 | 0.029178 |
| HP1BP3 | 1 | 2.1E+07 | 2.1E+07 | - | 7402 | heterochromat | 0.10466 | 6.61337 | 8.53867 | 0.0048859 | 0.029264 |
| MRPL54 | 19 | 3762682 | 3768575 | + | 985 | mitochondrial | -0.175 | 4.9266 | 8.53301 | 0.0048992 | 0.029332 |
| SNX13 | 7 | 1.8E+07 | 1.8E+07 | - | 16418 | sorting nexin 1 | 0.1531 | 5.31232 | 8.51469 | 0.0049425 | 0.029566 |
| TMUB1 | 7 | 1.5E+08 | 1.5E+08 | - | 1947 | transmembran | 0.15542 | 5.27794 | 8.50801 | 0.0049584 | 0.029649 |
| AGA | 4 | 1.8E+08 | 1.8E+08 | - | 2605 | aspartylglucos | 0.29792 | 3.26273 | 8.50575 | 0.0049638 | 0.029669 |
| SCRN2 | 17 | 4.8E+07 | 4.8E+07 | - | 2999 | secernin 2 [Sou | 0.17255 | 5.00386 | 8.50069 | 0.0049758 | 0.029729 |
| SLC25A11 | 17 | 4937130 | 4940053 | - | 2471 | solute carrier f | 0.12774 | 5.91326 | 8.49771 | 0.004983 | 0.029759 |
| NHSL1 | 6 | 1.4E+08 | 1.4E+08 | - | 8741 | NHS like 1 [Sou | 0.16941 | 5.03402 | 8.48106 | 0.005023 | 0.029986 |
| NAPEPLD | CHR7_1_ | 1E+08 | 1E+08 | - | 7561 | N-acyl phospha | 0.26217 | 3.6667 | 8.47531 | 0.005037 | 0.030056 |
| BLOC1S6 | 15 | 4.6E+07 | 4.6E+07 | + | 6812 | biogenesis of ly | -0.1577 | 5.42321 | 8.49272 | 0.0050413 | 0.03007 |
| SGSH | 17 | 8E+07 | 8E+07 | - | 6661 | N-sulfoglucosa | 0.23083 | 3.98319 | 8.46977 | 0.0050504 | 0.030112 |
| PACSIN1 | 6 | 3.4E+07 | 3.5E+07 | + | 4805 | protein kinase | -0.196 | 4.54165 | 8.46431 | 0.0050637 | 0.030178 |
| ZDHHC21 | 9 | 1.5E+07 | 1.5E+07 | - | 9426 | zinc finger DHH | 0.19 | 4.72481 | 8.44662 | 0.005107 | 0.030424 |
| AEN | 15 | 8.9E+07 | 8.9E+07 | + | 5976 | apoptosis enha | -0.0874 | 7.67326 | 8.4441 | 0.0051132 | 0.030448 |
| ATG3 | 3 | 1.1E+08 | 1.1E+08 | - | 6111 | autophagy rela | -0.1318 | 5.83916 | 8.44177 | 0.0051189 | 0.03047 |
| VSIG10L | 19 | 5.1E+07 | 5.1E+07 | - | 3771 | V-set and imm | -0.1919 | 4.67563 | 8.43592 | 0.0051334 | 0.030543 |
| PATL1 | 11 | 6E+07 | 6E+07 | - | 4335 | PAT1 homolog | -0.1212 | 6.07469 | 8.43306 | 0.0051404 | 0.030551 |
| WNT8B | 10 | 1E+08 | 1E+08 | + | 2112 | Wnt family me | -0.9453 | 0.3689 | 8.48894 | 0.0051405 | 0.030551 |
| NEDD4 | 15 | 5.6E+07 | 5.6E+07 | - | 9122 | neural precurs | 0.21802 | 4.26242 | 8.43279 | 0.0051411 | 0.030551 |
| RNF11 | 1 | 5.1E+07 | 5.1E+07 | + | 3112 | ring finger pro | 0.11385 | 6.25957 | 8.42513 | 0.0051601 | 0.030651 |
| RASSF3 | 12 | 6.5E+07 | 6.5E+07 | + | 4241 | Ras associatior | -0.1427 | 5.65131 | 8.4226 | 0.0051664 | 0.03067 |
| CBARP | 19 | 1228287 | 1238027 | - | 4591 | CACN subunit | 0.21478 | 4.20856 | 8.42212 | 0.0051676 | 0.03067 |
| AL160006 | 1 | 1.1E+08 | 1.1E+08 | + | 4216 | novel transcrip | -0.2406 | 3.97756 | 8.42068 | 0.0051712 | 0.030671 |
| ZNRF3 | 22 | 2.9E+07 | 2.9E+07 | + | 6968 | zinc and ring fi | -0.1669 | 5.09819 | 8.42036 | 0.005172 | 0.030671 |
| KPNA6 | 1 | 3.2E+07 | 3.2E+07 | + | 7457 | karyopherin su | -0.0948 | 7.16291 | 8.41631 | 0.0051821 | 0.030706 |
| SMARCD2 | 17 | 6.4E+07 | 6.4E+07 | - | 3734 | SWI/SNF relate | 0.09278 | 7.1342 | 8.41628 | 0.0051821 | 0.030706 |
| ZNF219 | 14 | 2.1E+07 | 2.1E+07 | - | 6663 | zinc finger pro | 0.08647 | 7.57625 | 8.41112 | 0.005195 | 0.030762 |
| FREM2 | 13 | 3.9E+07 | 3.9E+07 | + | 16275 | FRAS1 related | 0.11062 | 6.38812 | 8.41078 | 0.0051959 | 0.030762 |
| RNF10 | 12 | 1.2E+08 | 1.2E+08 | + | 5900 | ring finger pro | -0.0841 | 7.73985 | 8.4088 | 0.0052009 | 0.030779 |
| ST7 | 7 | 1.2E+08 | 1.2E+08 | + | 6742 | suppression of | 0.20823 | 4.36098 | 8.3973 | 0.0052298 | 0.030925 |
| VMA21 | X | 1.5E+08 | 1.5E+08 | + | 5109 | VMA21, vacuo | -0.1551 | 5.48194 | 8.43157 | 0.0052299 | 0.030925 |
| MEX3D | 19 | 1554669 | 1568058 | - | 2850 | mex-3 RNA bin | 0.09994 | 6.86 | 8.39463 | 0.0052365 | 0.030952 |
| PRMT7 | 16 | 6.8E+07 | 6.8E+07 | + | 9458 | protein argini | 0.14789 | 5.41124 | 8.39238 | 0.0052422 | 0.030972 |
| E2F2 | 1 | 2.4E+07 | 2.4E+07 | - | 5462 | E2F transcripti | 0.37725 | 2.70636 | 8.43955 | 0.0052505 | 0.031009 |
| ITGA4 | 2 | 1.8E+08 | 1.8E+08 | + | 10654 | integrin subun | 0.44764 | 2.02685 | 8.38567 | 0.0052592 | 0.031047 |
| FOS | 14 | 7.5E+07 | 7.5E+07 | + | 3186 | Fos proto-onc | -0.4156 | 2.3483 | 8.38296 | 0.005266 | 0.031067 |

| | | | | | | | | | | | |
|---|---|---|---|---|---|---|---|---|---|---|---|
| RILPL2 | 12 | 1.2E+08 | 1.2E+08 | - | 6141 | Rab interacting | -0.2933 | 3.28665 | 8.38267 | 0.0052668 | 0.031067 |
| SP1 | 12 | 5.3E+07 | 5.3E+07 | + | 8083 | Sp1 transcripti | -0.0904 | 7.23353 | 8.37587 | 0.0052841 | 0.031156 |
| CDC34 | 19 | 531760 | 542092 | + | 1894 | cell division cy | 0.09198 | 7.13739 | 8.37267 | 0.0052922 | 0.031191 |
| SERTAD3 | 19 | 4E+07 | 4E+07 | - | 1848 | SERTA domain | -0.2337 | 4.075 | 8.36772 | 0.0053049 | 0.031242 |
| ZSCAN12 | 6 | 2.8E+07 | 2.8E+07 | - | 5495 | zinc finger and | 0.24523 | 3.93551 | 8.36679 | 0.0053073 | 0.031242 |
| AKR7A2 | 1 | 1.9E+07 | 1.9E+07 | - | 1781 | aldo-keto redu | 0.16287 | 5.16444 | 8.36674 | 0.0053074 | 0.031242 |
| RNF217 | 6 | 1.2E+08 | 1.3E+08 | + | 12779 | ring finger pro | 0.25545 | 3.78844 | 8.35928 | 0.0053265 | 0.031342 |
| NRIP3 | 11 | 8980576 | 9004049 | - | 4123 | nuclear recept | -0.3366 | 2.85445 | 8.3526 | 0.0053437 | 0.03143 |
| TAP2 | R6_MHC_ | 3.3E+07 | 3.3E+07 | - | 6380 | transporter 2, | -0.2573 | 3.66357 | 8.34885 | 0.0053534 | 0.031474 |
| PPP1R12C | 19 | 5.5E+07 | 5.5E+07 | - | 7803 | protein phosph | -0.1632 | 5.19881 | 8.34469 | 0.0053641 | 0.031524 |
| COMMD7 | 20 | 3.3E+07 | 3.3E+07 | - | 2584 | COMM domain | -0.1698 | 5.04645 | 8.34183 | 0.0053716 | 0.031555 |
| SIMC1 | 5 | 1.8E+08 | 1.8E+08 | + | 5536 | SUMO interact | 0.26075 | 3.60571 | 8.33836 | 0.0053806 | 0.031595 |
| PARVB | 22 | 4.4E+07 | 4.4E+07 | + | 7403 | parvin beta [So | -0.2003 | 4.48846 | 8.33694 | 0.0053842 | 0.031604 |
| PRRC2C | 1 | 1.7E+08 | 1.7E+08 | + | 14173 | proline rich co | -0.0699 | 8.9758 | 8.33208 | 0.0053969 | 0.031665 |
| CNDP2 | 18 | 7.4E+07 | 7.5E+07 | + | 11822 | carnosine dipe | -0.0838 | 7.81315 | 8.32716 | 0.0054097 | 0.031727 |
| GGA1 | 22 | 3.8E+07 | 3.8E+07 | + | 9119 | golgi associate | 0.13196 | 5.74772 | 8.319 | 0.0054311 | 0.031829 |
| SYT6 | 1 | 1.1E+08 | 1.1E+08 | - | 5104 | synaptotagmin | -0.1503 | 5.35447 | 8.31882 | 0.0054315 | 0.031829 |
| LMNTD2 | 11 | 554850 | 560738 | - | 2898 | lamin tail dom | 0.59263 | 1.3184 | 8.3167 | 0.0054371 | 0.031849 |
| MIA2 | 14 | 3.9E+07 | 3.9E+07 | + | 10539 | MIA SH3 doma | 0.21923 | 4.12087 | 8.3134 | 0.0054458 | 0.031874 |
| PIGQ | 16 | 566995 | 584136 | + | 7904 | phosphatidylin | 0.20265 | 4.44368 | 8.31224 | 0.0054488 | 0.031879 |
| CEL | 9 | 1.3E+08 | 1.3E+08 | + | 2384 | carboxyl ester | 0.68931 | 0.9511 | 8.31049 | 0.0054534 | 0.031893 |
| ARFGAP3 | 22 | 4.3E+07 | 4.3E+07 | - | 3273 | ADP ribosylatio | -0.1837 | 4.74083 | 8.30804 | 0.0054599 | 0.031917 |
| NECAP1 | 12 | 8076939 | 8097881 | + | 8785 | NECAP endocy | -0.1871 | 4.62794 | 8.30304 | 0.0054731 | 0.031982 |
| MCMBP | 10 | 1.2E+08 | 1.2E+08 | - | 6000 | minichromoso | 0.09261 | 7.06901 | 8.29946 | 0.0054826 | 0.032024 |
| LINC0225 | 15 | 9.7E+07 | 9.7E+07 | + | 989 | long intergenic | 0.62434 | 1.22088 | 8.28856 | 0.0055115 | 0.03218 |
| SHQ1 | 3 | 7.3E+07 | 7.3E+07 | - | 5833 | SHQ1, H/ACA r | -0.1705 | 5.15925 | 8.32647 | 0.0055144 | 0.032183 |
| PBDC1 | X | 7.6E+07 | 7.6E+07 | + | 1201 | polysaccharide | -0.1889 | 4.8733 | 8.36138 | 0.0055165 | 0.032183 |
| EPHB3 | 3 | 1.8E+08 | 1.8E+08 | + | 4530 | EPH receptor B | -0.6635 | 1.26414 | 8.37917 | 0.0055195 | 0.032187 |
| SLC12A2 | 5 | 1.3E+08 | 1.3E+08 | + | 10653 | solute carrier f | 0.1875 | 4.68831 | 8.28089 | 0.005532 | 0.032247 |
| MEF2D | 1 | 1.6E+08 | 1.6E+08 | - | 6351 | myocyte enhar | -0.1502 | 5.38465 | 8.27979 | 0.0055349 | 0.032251 |
| SCNM1 | 1 | 1.5E+08 | 1.5E+08 | + | 3265 | sodium channe | -0.1979 | 4.48899 | 8.27712 | 0.0055421 | 0.032266 |
| IGIP | 5 | 1.4E+08 | 1.4E+08 | + | 3458 | IgA inducing pr | 0.9505 | 0.14042 | 8.27561 | 0.0055461 | 0.032266 |
| FNBP1 | 9 | 1.3E+08 | 1.3E+08 | - | 6600 | formin binding | 0.15212 | 5.3971 | 8.27502 | 0.0055477 | 0.032266 |
| ADAMTS2 | 5 | 1.8E+08 | 1.8E+08 | - | 8755 | ADAM metallo | -0.25 | 3.79713 | 8.27307 | 0.0055529 | 0.032266 |
| TBC1D23 | 3 | 1E+08 | 1E+08 | + | 5176 | TBC1 domain f | -0.1826 | 4.83718 | 8.27305 | 0.005553 | 0.032266 |
| RHBDL1 | 16 | 675666 | 678268 | + | 1884 | rhomboid like | 0.47833 | 2.0741 | 8.36575 | 0.0055546 | 0.032266 |
| NSFL1C | 20 | 1442162 | 1473842 | - | 6531 | NSFL1 cofactor | -0.1029 | 6.67366 | 8.27229 | 0.005555 | 0.032266 |
| ZKSCAN1 | 7 | 1E+08 | 1E+08 | + | 10319 | zinc finger with | 0.15544 | 5.26805 | 8.27165 | 0.0055567 | 0.032266 |
| RAB42 | 1 | 2.9E+07 | 2.9E+07 | + | 2200 | RAB42, membe | 0.32864 | 3.06333 | 8.28037 | 0.0055577 | 0.032266 |
| MARVELD | 16 | 7.2E+07 | 7.2E+07 | + | 6446 | MARVEL doma | -0.1487 | 5.45929 | 8.2672 | 0.0055687 | 0.032317 |
| TXLNGY | Y | 2E+07 | 2E+07 | + | 11377 | taxilin gamma | 0.12697 | 5.95541 | 8.26256 | 0.0055812 | 0.032376 |
| EXOSC9 | 4 | 1.2E+08 | 1.2E+08 | + | 4711 | exosome comp | -0.1166 | 6.18949 | 8.25556 | 0.0056001 | 0.032473 |
| THOC5 | 22 | 3E+07 | 3E+07 | - | 7878 | THO complex 5 | -0.1195 | 6.09705 | 8.24466 | 0.0056297 | 0.032632 |

| | | | | | | | | | | | |
|---|---|---|---|---|---|---|---|---|---|---|---|
| PRMT5 | 14 | 2.3E+07 | 2.3E+07 | - | 3276 | protein arginin | -0.0923 | 7.19094 | 8.24183 | 0.0056374 | 0.032655 |
| STK38L | 12 | 2.7E+07 | 2.7E+07 | + | 6863 | serine/threoni | 0.11259 | 6.31771 | 8.24108 | 0.0056395 | 0.032655 |
| PTER | 10 | 1.6E+07 | 1.7E+07 | + | 4003 | phosphotrieste | -0.2752 | 3.47057 | 8.24066 | 0.0056406 | 0.032655 |
| FAM86JP | 3 | 1.3E+08 | 1.3E+08 | + | 2519 | family with seq | 0.81419 | 0.53149 | 8.2391 | 0.0056449 | 0.032667 |
| FBN3 | 19 | 8065402 | 8149592 | - | 9985 | fibrillin 3 [Sour | -0.1017 | 6.68608 | 8.23328 | 0.0056608 | 0.032746 |
| C5orf38 | 5 | 2752131 | 2755397 | + | 1925 | chromosome 5 | -0.6195 | 1.26413 | 8.22949 | 0.0056712 | 0.032793 |
| DUSP3 | 17 | 4.4E+07 | 4.4E+07 | - | 4997 | dual specificity | -0.134 | 5.80903 | 8.22636 | 0.0056798 | 0.032804 |
| ALAS1 | 3 | 5.2E+07 | 5.2E+07 | + | 2536 | 5'-aminolevulin | -0.1271 | 5.89178 | 8.2259 | 0.0056811 | 0.032804 |
| METTL5 | 2 | 1.7E+08 | 1.7E+08 | - | 2865 | methyltransfer | -0.147 | 5.44826 | 8.22564 | 0.0056818 | 0.032804 |
| HTR7P1 | 12 | 1.3E+07 | 1.3E+07 | + | 4411 | 5-hydroxytrypt | -0.4223 | 2.32772 | 8.2261 | 0.0056824 | 0.032804 |
| RBBP8 | 18 | 2.3E+07 | 2.3E+07 | + | 5078 | RB binding pro | -0.1429 | 5.62019 | 8.22377 | 0.0056869 | 0.032817 |
| TRIM5 | 11 | 5663557 | 5938619 | - | 4804 | tripartite motif | -0.1765 | 4.88988 | 8.2227 | 0.0056899 | 0.032821 |
| GTPBP8 | 3 | 1.1E+08 | 1.1E+08 | + | 3826 | GTP binding pr | -0.2181 | 4.20314 | 8.21774 | 0.0057036 | 0.032887 |
| RTN2 | 19 | 4.5E+07 | 4.5E+07 | - | 3227 | reticulon 2 [So | -0.1885 | 4.69844 | 8.21026 | 0.0057242 | 0.032993 |
| GGNBP2 | SCHR17_7 | 3.7E+07 | 3.7E+07 | + | 9705 | gametogenetin | -0.1356 | 5.71854 | 8.20714 | 0.0057329 | 0.03303 |
| LRP8 | 1 | 5.3E+07 | 5.3E+07 | - | 10263 | LDL receptor re | 0.09504 | 6.95158 | 8.20562 | 0.0057371 | 0.033041 |
| PCDHB13 | 5 | 1.4E+08 | 1.4E+08 | + | 5061 | protocadherin | 0.50136 | 1.69777 | 8.2026 | 0.0057455 | 0.033076 |
| TOR3A | 1 | 1.8E+08 | 1.8E+08 | + | 3224 | torsin family 3 | -0.1607 | 5.17164 | 8.18759 | 0.0057874 | 0.033304 |
| ADAMTSL | 1 | 1.5E+08 | 1.5E+08 | + | 6432 | ADAMTS like 4 | 0.36417 | 2.58708 | 8.18225 | 0.0058024 | 0.033377 |
| ANKRD35 | 1 | 1.5E+08 | 1.5E+08 | - | 3363 | ankyrin repeat | -0.2829 | 3.36955 | 8.18113 | 0.0058055 | 0.033381 |
| SLC25A15 | 13 | 4.1E+07 | 4.1E+07 | + | 4186 | solute carrier f | 0.15349 | 5.30225 | 8.17821 | 0.0058138 | 0.033415 |
| SLC37A4 | 11 | 1.2E+08 | 1.2E+08 | - | 6186 | solute carrier f | 0.15053 | 5.31999 | 8.17693 | 0.0058174 | 0.033423 |
| GM2A | 5 | 1.5E+08 | 1.5E+08 | + | 4479 | GM2 gangliosi | -0.1352 | 5.655 | 8.1737 | 0.0058265 | 0.033445 |
| FZD6 | 8 | 1E+08 | 1E+08 | + | 4065 | frizzled class re | 0.20362 | 4.38326 | 8.17317 | 0.005828 | 0.033445 |
| E2F1 | 20 | 3.4E+07 | 3.4E+07 | - | 2690 | E2F transcripti | 0.14215 | 5.58127 | 8.17309 | 0.0058282 | 0.033445 |
| GDAP1 | 8 | 7.4E+07 | 7.4E+07 | + | 4052 | ganglioside ind | 0.15656 | 5.19058 | 8.16042 | 0.0058641 | 0.033637 |
| KIAA0753 | 17 | 6578147 | 6640711 | - | 6084 | KIAA0753 [Sou | -0.1673 | 4.97521 | 8.15678 | 0.0058745 | 0.033683 |
| AFG3L2 | 18 | 1.2E+07 | 1.2E+07 | - | 6624 | AFG3 like matr | -0.0942 | 6.9749 | 8.15223 | 0.0058874 | 0.033739 |
| CYTH2 | 19 | 4.8E+07 | 4.8E+07 | + | 9333 | cytohesin 2 [So | 0.11729 | 6.1724 | 8.15173 | 0.0058889 | 0.033739 |
| KXD1 | 19 | 1.9E+07 | 1.9E+07 | + | 4112 | KxDL motif con | 0.10481 | 6.55019 | 8.1445 | 0.0059095 | 0.033844 |
| DCBLD2 | 3 | 9.9E+07 | 9.9E+07 | - | 11280 | discoidin, CUB | 0.1058 | 6.45968 | 8.13823 | 0.0059275 | 0.03391 |
| GDI2 | 10 | 5765223 | 5842132 | - | 3918 | GDP dissociatio | -0.0702 | 8.87843 | 8.13801 | 0.0059282 | 0.03391 |
| ALKBH5 | 17 | 1.8E+07 | 1.8E+07 | + | 3601 | alkB homolog 5 | -0.1097 | 6.33115 | 8.13354 | 0.005941 | 0.03397 |
| SEMG1 | 20 | 4.5E+07 | 4.5E+07 | + | 1622 | semenogelin 1 | -0.7448 | 0.70883 | 8.12956 | 0.0059525 | 0.034022 |
| NT5C | 17 | 7.5E+07 | 7.5E+07 | - | 1533 | 5', 3'-nucleotid | 0.20627 | 4.2952 | 8.12269 | 0.0059724 | 0.034122 |
| ZBTB38 | 3 | 1.4E+08 | 1.4E+08 | + | 14243 | zinc finger and | 0.18014 | 4.82042 | 8.11324 | 0.0059998 | 0.034265 |
| MYCBP2 | 13 | 7.7E+07 | 7.7E+07 | - | 15987 | MYC binding p | 0.10437 | 6.55925 | 8.10962 | 0.0060104 | 0.034312 |
| ARFRP1 | 20 | 6.4E+07 | 6.4E+07 | - | 7627 | ADP ribosylatio | 0.20398 | 4.31401 | 8.10856 | 0.0060135 | 0.034316 |
| SMYD3 | 1 | 2.5E+08 | 2.5E+08 | - | 5866 | SET and MYND | 0.22739 | 4.05649 | 8.10695 | 0.0060182 | 0.034329 |
| TMEM64 | 8 | 9.1E+07 | 9.1E+07 | - | 5870 | transmembran | -0.5633 | 6.28285 | 27.2055 | 0.00603 | 0.034383 |
| TMEM159 | 16 | 2.1E+07 | 2.1E+07 | + | 2558 | transmembran | -0.1802 | 4.88119 | 8.10403 | 0.0060381 | 0.034406 |
| TIGAR | 12 | 4307763 | 4360028 | + | 9444 | TP53 induced g | -0.2696 | 3.49156 | 8.09987 | 0.0060389 | 0.034406 |
| RFWD3 | 16 | 7.5E+07 | 7.5E+07 | - | 6456 | ring finger and | 0.08388 | 7.56431 | 8.09851 | 0.0060429 | 0.034415 |

| | | | | | | | | | | | | |
|---|---|---|---|---|---|---|---|---|---|---|---|---|
| ZNF204P | 6 | 2.7E+07 | 2.7E+07 | - | | 2397 | zinc finger prot | -0.2053 | 4.50819 | 8.12587 | 0.0060569 | 0.034474 |
| IFIH1 | 2 | 1.6E+08 | 1.6E+08 | - | | 5113 | interferon indu | 0.64076 | 1.15809 | 8.09335 | 0.006058 | 0.034474 |
| MT-ATP6 | MT | 8527 | 9207 | + | | 681 | mitochondriall | -0.0571 | 11.6428 | 8.09019 | 0.0060673 | 0.034511 |
| TRAF5 | 1 | 2.1E+08 | 2.1E+08 | + | | 9022 | TNF receptor a | 0.1841 | 4.65764 | 8.08953 | 0.0060693 | 0.034511 |
| FTL | 19 | 4.9E+07 | 4.9E+07 | + | | 871 | ferritin light ch | 0.06871 | 9.88281 | 8.14247 | 0.0060808 | 0.03455 |
| SLC13A3 | 20 | 4.7E+07 | 4.7E+07 | - | | 6055 | solute carrier f | -0.2156 | 4.251 | 8.08558 | 0.0060809 | 0.03455 |
| UHRF1 | 19 | 4903080 | 4962154 | + | | 5145 | ubiquitin like w | 0.10899 | 6.37392 | 8.08287 | 0.006089 | 0.034578 |
| CLASP2 | 3 | 3.3E+07 | 3.4E+07 | - | | 15214 | cytoplasmic lin | 0.1077 | 6.4188 | 8.0823 | 0.0060906 | 0.034578 |
| KDELC2 | 11 | 1.1E+08 | 1.1E+08 | - | | 4938 | KDEL motif con | 0.15816 | 5.13267 | 8.08069 | 0.0060954 | 0.034591 |
| NBAS | 2 | 1.5E+07 | 1.6E+07 | - | | 8947 | neuroblastoma | 0.1096 | 6.35291 | 8.07785 | 0.0061038 | 0.034615 |
| NISCH | 3 | 5.2E+07 | 5.2E+07 | + | | 9062 | nischarin [Sour | 0.09457 | 7.11588 | 8.07763 | 0.0061044 | 0.034615 |
| CHP2 | 16 | 2.4E+07 | 2.4E+07 | + | | 1967 | calcineurin like | -0.5789 | 1.38388 | 8.07029 | 0.0061263 | 0.034702 |
| C1orf226 | 1 | 1.6E+08 | 1.6E+08 | + | | 4491 | chromosome 1 | 0.18673 | 4.60632 | 8.07002 | 0.0061271 | 0.034702 |
| CNIH2 | 11 | 6.6E+07 | 6.6E+07 | + | | 2078 | cornichon fam | 0.23387 | 3.97038 | 8.06712 | 0.0061357 | 0.034731 |
| TNRC6C | 17 | 7.8E+07 | 7.8E+07 | + | | 11111 | trinucleotide r | -0.1327 | 5.70974 | 8.06672 | 0.0061369 | 0.034731 |
| C10orf88 | 10 | 1.2E+08 | 1.2E+08 | - | | 3083 | chromosome 1 | 0.28069 | 3.43782 | 8.06197 | 0.0061511 | 0.034792 |
| HDGF | 1 | 1.6E+08 | 1.6E+08 | - | | 3674 | heparin bindin | 0.07006 | 8.84129 | 8.06144 | 0.0061527 | 0.034792 |
| HIBADH | 7 | 2.8E+07 | 2.8E+07 | - | | 2211 | 3-hydroxyisobu | 0.14207 | 5.52569 | 8.05769 | 0.0061639 | 0.034842 |
| GSTA4 | 6 | 5.3E+07 | 5.3E+07 | - | | 2101 | glutathione S-t | 0.20604 | 4.30569 | 8.05683 | 0.0061664 | 0.034843 |
| GGT7 | 20 | 3.5E+07 | 3.5E+07 | - | | 2962 | gamma-glutam | 0.16209 | 5.06914 | 8.05466 | 0.006173 | 0.034866 |
| ZNF525 | 19 | 5.3E+07 | 5.3E+07 | + | | 7193 | zinc finger prot | -0.1413 | 5.58062 | 8.04717 | 0.0061955 | 0.034979 |
| TGFBR2 | 3 | 3.1E+07 | 3.1E+07 | + | | 4704 | transforming g | 0.21376 | 4.18534 | 8.04531 | 0.0062011 | 0.034997 |
| TMEM59 | 1 | 5.4E+07 | 5.4E+07 | - | | 9149 | transmembran | -0.1091 | 6.45799 | 8.04299 | 0.0062081 | 0.035023 |
| DNMT3A | 2 | 2.5E+07 | 2.5E+07 | - | | 12020 | DNA methyltra | 0.08516 | 7.46557 | 8.04004 | 0.006217 | 0.03506 |
| B3GNTL1 | SCHR17_1 | 8.3E+07 | 8.3E+07 | - | | 6891 | UDP-GlcNAc:be | 0.35596 | 2.61744 | 8.03547 | 0.0062308 | 0.035124 |
| ZIC3 | X | 1.4E+08 | 1.4E+08 | + | | 4668 | Zic family mem | -0.1042 | 6.91695 | 8.15041 | 0.0062496 | 0.035216 |
| ARHGAP4 | 17 | 1.3E+07 | 1.3E+07 | + | | 9322 | Rho GTPase ac | -0.1571 | 5.15504 | 8.02609 | 0.0062593 | 0.035257 |
| PARP4 | 13 | 2.4E+07 | 2.5E+07 | - | | 6043 | poly(ADP-ribos | 0.10856 | 6.42624 | 8.02447 | 0.0062642 | 0.035271 |
| FAM19A5 | 22 | 4.8E+07 | 4.9E+07 | + | | 3173 | family with seq | -0.1679 | 5.00554 | 8.02217 | 0.0062712 | 0.035296 |
| TXLNG | X | 1.7E+07 | 1.7E+07 | + | | 4455 | taxilin gamma | -0.1109 | 6.43393 | 8.02029 | 0.006277 | 0.035315 |
| TAC3 | 12 | 5.7E+07 | 5.7E+07 | - | | 1571 | tachykinin 3 [S | 0.38973 | 2.38739 | 8.01391 | 0.0062965 | 0.035411 |
| PHB2 | 12 | 6965327 | 6970825 | - | | 2499 | prohibitin 2 [So | 0.07361 | 8.42873 | 8.01105 | 0.0063052 | 0.035446 |
| DGCR5 | 22 | 1.9E+07 | 1.9E+07 | + | | 5676 | DiGeorge synd | -0.4578 | 1.92651 | 8.00578 | 0.0063214 | 0.035519 |
| CDK7 | SCHR5_2_ | 6.9E+07 | 6.9E+07 | + | | 2416 | cyclin depende | -0.1355 | 5.64704 | 8.0052 | 0.0063232 | 0.035519 |
| PHACTR1 | 6 | 1.3E+07 | 1.3E+07 | + | | 11198 | phosphatase a | -0.3121 | 3.10774 | 8.00389 | 0.0063272 | 0.035528 |
| AC124319 | 17 | 8E+07 | 8E+07 | + | | 28570 | ring finger prot | 0.10189 | 6.87482 | 8.03878 | 0.0063323 | 0.035543 |
| LENG8 | R19LRC_CO | 5.4E+07 | 5.4E+07 | + | | 6469 | leukocyte rece | 0.10247 | 6.64538 | 7.99376 | 0.0063585 | 0.035672 |
| SNRPD2 | 19 | 4.6E+07 | 4.6E+07 | - | | 1608 | small nuclear r | -0.0828 | 7.7646 | 7.99287 | 0.0063613 | 0.035672 |
| TOP3A | 17 | 1.8E+07 | 1.8E+07 | - | | 8224 | DNA topoisom | 0.12834 | 5.77796 | 7.99212 | 0.0063636 | 0.035672 |
| CLPTM1 | 19 | 4.5E+07 | 4.5E+07 | + | | 5095 | CLPTM1, trans | -0.0943 | 6.91551 | 7.99156 | 0.0063653 | 0.035672 |
| PALM | 19 | 708935 | 748329 | + | | 4912 | paralemmin [S | 0.18543 | 4.63681 | 7.97081 | 0.00643 | 0.03602 |
| ZNF608 | 5 | 1.2E+08 | 1.2E+08 | - | | 6922 | zinc finger prot | 0.13923 | 5.63381 | 7.96977 | 0.0064332 | 0.036024 |
| HLA-DOA | R6_MHC_ | 3.3E+07 | 3.3E+07 | - | | 3987 | major histocom | 0.15971 | 5.15639 | 7.96877 | 0.0064363 | 0.036028 |

| | | | | | | | | | | | |
|---|---|---|---|---|---|---|---|---|---|---|---|
| TCP1 | 6 | 1.6E+08 | 1.6E+08 | - | 4883 | t-complex 1 [S | -0.0723 | 8.56675 | 7.96295 | 0.0064546 | 0.036116 |
| COL4A3 | 2 | 2.3E+08 | 2.3E+08 | + | 9124 | collagen type I | 0.66337 | 0.95114 | 7.95263 | 0.0064871 | 0.036262 |
| TMEM179 | SCHR14_2 | 1E+08 | 1E+08 | - | 5356 | transmembran | 0.41677 | 2.23513 | 7.95209 | 0.0064889 | 0.036262 |
| CHMP4C | 8 | 8.2E+07 | 8.2E+07 | + | 1852 | charged multiv | 0.37273 | 2.4411 | 7.9515 | 0.0064907 | 0.036262 |
| RNF146 | 6 | 1.3E+08 | 1.3E+08 | + | 3508 | ring finger pro | 0.15539 | 5.27738 | 7.94985 | 0.0064959 | 0.036277 |
| ARID1B | 6 | 1.6E+08 | 1.6E+08 | + | 44677 | AT-rich interac | 0.0935 | 6.95611 | 7.949 | 0.0064986 | 0.036278 |
| HIST1H2B | 6 | 2.7E+07 | 2.7E+07 | - | 950 | histone cluster | -0.509 | 1.73264 | 7.94759 | 0.0065031 | 0.036289 |
| CYB5A | 18 | 7.4E+07 | 7.4E+07 | - | 9271 | cytochrome b5 | -0.1242 | 6.00026 | 7.94235 | 0.0065197 | 0.03636 |
| ITGA3 | 17 | 5E+07 | 5E+07 | + | 9802 | integrin subun | 0.18438 | 5.83399 | 9.32225 | 0.0065228 | 0.03636 |
| ATP5MG | 11 | 1.2E+08 | 1.2E+08 | + | 2926 | ATP synthase r | 0.12349 | 6.0375 | 7.94064 | 0.0065252 | 0.03636 |
| NPR1 | 1 | 1.5E+08 | 1.5E+08 | + | 4952 | natriuretic pep | 0.18506 | 4.61619 | 7.94037 | 0.006526 | 0.03636 |
| ASPHD1 | 16 | 3E+07 | 3E+07 | + | 2312 | aspartate beta | -0.2033 | 4.3725 | 7.93629 | 0.006539 | 0.036418 |
| ITGB5 | 3 | 1.2E+08 | 1.2E+08 | - | 6515 | integrin subun | 0.27914 | 8.85019 | 20.9994 | 0.0066165 | 0.036835 |
| AEBP2 | 12 | 1.9E+07 | 2E+07 | + | 6754 | AE binding pro | -0.1225 | 5.92703 | 7.91012 | 0.0066229 | 0.036857 |
| GLUD1 | 10 | 8.7E+07 | 8.7E+07 | - | 3780 | glutamate deh | 0.08619 | 7.33155 | 7.90059 | 0.0066538 | 0.037015 |
| S100PBP | 1 | 3.3E+07 | 3.3E+07 | + | 9190 | S100P binding | -0.1399 | 5.54506 | 7.89609 | 0.0066685 | 0.037082 |
| DGCR8 | 22 | 2E+07 | 2E+07 | + | 7315 | DGCR8, microp | 0.11807 | 6.06018 | 7.89164 | 0.0066829 | 0.037148 |
| CCDC162I | 6 | 1.1E+08 | 1.1E+08 | + | 10012 | coiled-coil dom | 0.47134 | 1.80233 | 7.88644 | 0.0066999 | 0.037228 |
| FAM98B | 15 | 3.8E+07 | 3.8E+07 | + | 6851 | family with sec | -0.1089 | 6.40301 | 7.88538 | 0.0067034 | 0.037233 |
| KIAA0100 | 17 | 2.9E+07 | 2.9E+07 | - | 9232 | KIAA0100 [Sou | 0.07724 | 8.03947 | 7.87806 | 0.0067274 | 0.03734 |
| IL17RD | 3 | 5.7E+07 | 5.7E+07 | - | 9561 | interleukin 17 | -0.0821 | 7.69974 | 7.87789 | 0.0067279 | 0.03734 |
| UPF1 | 19 | 1.9E+07 | 1.9E+07 | + | 8600 | UPF1, RNA hel | 0.07984 | 7.78018 | 7.87682 | 0.0067314 | 0.03734 |
| PDCL3P4 | 3 | 1E+08 | 1E+08 | + | 720 | phosducin-like | -0.7245 | 0.74026 | 7.87622 | 0.0067334 | 0.03734 |
| AC073111 | 7 | 1.5E+08 | 1.5E+08 | + | 4333 | novel zinc fing | -0.232 | 3.9305 | 7.87531 | 0.0067364 | 0.03734 |
| NLGN3 | X | 7.1E+07 | 7.1E+07 | + | 4006 | neuroligin 3 [S | -0.196 | 4.3987 | 7.87473 | 0.0067383 | 0.03734 |
| EVC2 | 4 | 5542772 | 5709548 | - | 5501 | EvC ciliary com | -0.2436 | 3.798 | 7.8725 | 0.0067456 | 0.037363 |
| CPT2 | 1 | 5.3E+07 | 5.3E+07 | + | 10089 | carnitine palm | 0.17355 | 4.82167 | 7.87192 | 0.0067476 | 0.037363 |
| TMEM132 | 12 | 1.3E+08 | 1.3E+08 | + | 15703 | transmembran | 0.08868 | 7.40467 | 7.87071 | 0.0067516 | 0.03737 |
| ZNF862 | 7 | 1.5E+08 | 1.5E+08 | + | 8422 | zinc finger pro | 0.43103 | 2.03943 | 7.86913 | 0.0067567 | 0.037385 |
| SLC12A4 | 16 | 6.8E+07 | 6.8E+07 | - | 8298 | solute carrier f | 0.14476 | 5.83074 | 8.15146 | 0.0067637 | 0.037408 |
| NKRF | X | 1.2E+08 | 1.2E+08 | - | 4552 | NFKB repressin | 0.23151 | 3.88443 | 7.85874 | 0.0067911 | 0.037546 |
| TRIP6 | 7 | 1E+08 | 1E+08 | + | 2553 | thyroid hormo | 0.0917 | 7.22729 | 7.85173 | 0.0068144 | 0.03766 |
| MAST3 | 19 | 1.8E+07 | 1.8E+07 | + | 6772 | microtubule as | -0.2339 | 3.88213 | 7.85043 | 0.0068187 | 0.03767 |
| RNF182 | 6 | 1.4E+07 | 1.4E+07 | + | 4098 | ring finger pro | 0.26302 | 3.52622 | 7.8487 | 0.0068245 | 0.037687 |
| PFKFB4 | 3 | 4.9E+07 | 4.9E+07 | - | 6107 | 6-phosphofruc | 0.26871 | 3.55004 | 7.83376 | 0.0068745 | 0.037948 |
| CCSER2 | 10 | 8.4E+07 | 8.5E+07 | + | 8424 | coiled-coil seri | -0.1573 | 5.18317 | 7.82623 | 0.0068999 | 0.038074 |
| CAPRIN2 | 12 | 3.1E+07 | 3.1E+07 | - | 8392 | caprin family n | 0.14994 | 5.39717 | 7.82523 | 0.0069032 | 0.038078 |
| WASF1 | 6 | 1.1E+08 | 1.1E+08 | - | 3260 | WAS protein fa | 0.13447 | 5.68278 | 7.81967 | 0.006922 | 0.038152 |
| ILK | 11 | 6603708 | 6610874 | + | 3484 | integrin linked | 0.24524 | 3.82899 | 7.81966 | 0.006922 | 0.038152 |
| SEMA5B | 3 | 1.2E+08 | 1.2E+08 | - | 8517 | semaphorin 5E | -0.179 | 4.97239 | 7.90225 | 0.0069269 | 0.038164 |
| RNF168 | 3 | 2E+08 | 2E+08 | - | 5347 | ring finger pro | -0.1167 | 6.11767 | 7.81297 | 0.0069447 | 0.038248 |
| MRPL27 | 17 | 5E+07 | 5E+07 | - | 3020 | mitochondrial | -0.1455 | 5.4177 | 7.81001 | 0.0069548 | 0.038288 |
| EVC | 4 | 5711201 | 5814305 | + | 7144 | EvC ciliary com | 0.23024 | 3.86577 | 7.80659 | 0.0069664 | 0.038326 |

| | | | | | | | | | | | |
|---|---|---|---|---|---|---|---|---|---|---|---|
| MARS | 12 | 5.7E+07 | 5.8E+07 | + | 7028 | methionyl-tRN | -0.0708 | 8.60971 | 7.80645 | 0.0069669 | 0.038326 |
| PPP3CA | 4 | 1E+08 | 1E+08 | - | 5213 | protein phosph | 0.11064 | 6.28931 | 7.80458 | 0.0069733 | 0.038343 |
| FAR2 | 12 | 2.9E+07 | 2.9E+07 | + | 5411 | fatty acyl-CoA | 0.1527 | 5.33285 | 7.80397 | 0.0069753 | 0.038343 |
| ZNF415 | 19 | 5.3E+07 | 5.3E+07 | - | 3780 | zinc finger prot | -0.3101 | 3.08346 | 7.80239 | 0.0069807 | 0.038358 |
| NREP | 5 | 1.1E+08 | 1.1E+08 | - | 9337 | neuronal regen | 0.0821 | 7.74828 | 7.79955 | 0.0069904 | 0.038397 |
| CENPO | 2 | 2.5E+07 | 2.5E+07 | + | 5518 | centromere pr | 0.14469 | 5.49625 | 7.79836 | 0.0069945 | 0.038404 |
| MRPL47 | 3 | 1.8E+08 | 1.8E+08 | - | 1370 | mitochondrial | -0.1291 | 5.76537 | 7.79592 | 0.0070028 | 0.038435 |
| TMEM33 | 4 | 4.2E+07 | 4.2E+07 | + | 8462 | transmembran | -0.1088 | 6.30035 | 7.79278 | 0.0070136 | 0.03848 |
| WDR1 | 4 | 1E+07 | 1E+07 | - | 7827 | WD repeat dor | -0.0694 | 9.04703 | 7.78231 | 0.0070496 | 0.038648 |
| TYRO3 | 15 | 4.2E+07 | 4.2E+07 | + | 10770 | TYRO3 protein | 0.08468 | 7.42627 | 7.78104 | 0.007054 | 0.038657 |
| MYH2 | 17 | 1.1E+07 | 1.1E+07 | - | 6366 | myosin heavy c | -0.3317 | 2.77852 | 7.77993 | 0.0070578 | 0.038663 |
| FADS3 | 11 | 6.2E+07 | 6.2E+07 | - | 6294 | fatty acid desa | -0.1443 | 5.44246 | 7.77835 | 0.0070633 | 0.038678 |
| HSF1 | 8 | 1.4E+08 | 1.4E+08 | + | 5305 | heat shock trai | 0.10433 | 6.54475 | 7.7732 | 0.0070811 | 0.038753 |
| WLS | 1 | 6.8E+07 | 6.8E+07 | - | 4459 | Wnt ligand sec | 0.51465 | 1.57618 | 7.77286 | 0.0070823 | 0.038753 |
| MEX3A | 1 | 1.6E+08 | 1.6E+08 | - | 6591 | mex-3 RNA bin | 0.09065 | 7.1275 | 7.76702 | 0.0071026 | 0.038849 |
| NHLRC3 | 13 | 3.9E+07 | 3.9E+07 | + | 6409 | NHL repeat co | -0.2036 | 4.29843 | 7.76618 | 0.0071055 | 0.03885 |
| ARSJ | 4 | 1.1E+08 | 1.1E+08 | - | 5146 | arylsulfatase fa | 0.29786 | 3.17758 | 7.76339 | 0.0071152 | 0.038889 |
| UBXN7 | 3 | 2E+08 | 2E+08 | - | 11247 | UBX domain pr | -0.0953 | 7.05024 | 7.75609 | 0.0071731 | 0.039175 |
| FKBP11 | 12 | 4.9E+07 | 4.9E+07 | - | 4957 | FKBP prolyl iso | -0.1721 | 4.78278 | 7.74099 | 0.0071937 | 0.039273 |
| HSPH1 | 13 | 3.1E+07 | 3.1E+07 | - | 8390 | heat shock pro | -0.0835 | 7.7242 | 7.73584 | 0.0072118 | 0.039357 |
| TSPAN15 | 10 | 6.9E+07 | 7E+07 | + | 2651 | tetraspanin 15 | 0.28698 | 3.30186 | 7.73458 | 0.0072163 | 0.039366 |
| OSBP2 | 22 | 3.1E+07 | 3.1E+07 | + | 6632 | oxysterol bindi | -0.2774 | 3.36429 | 7.73241 | 0.007224 | 0.039393 |
| TRIM16 | 17 | 1.6E+07 | 1.6E+07 | - | 5640 | tripartite motif | 0.30277 | 3.12282 | 7.72771 | 0.0072406 | 0.039469 |
| AL731571 | 10 | 1.2E+08 | 1.3E+08 | + | 5428 | novel transcrip | 0.30119 | 3.11988 | 7.72379 | 0.0072545 | 0.03953 |
| TMCO6 | 5 | 1.4E+08 | 1.4E+08 | + | 3502 | transmembran | 0.2433 | 3.77062 | 7.72133 | 0.0072633 | 0.039563 |
| KREMEN2 | 16 | 2964216 | 2968383 | + | 2342 | kringle contain | -0.2151 | 4.08927 | 7.71522 | 0.0072851 | 0.039665 |
| TRIM59 | 3 | 1.6E+08 | 1.6E+08 | - | 7822 | tripartite motif | -0.1436 | 5.38045 | 7.71452 | 0.0072876 | 0.039665 |
| HACE1 | 6 | 1E+08 | 1E+08 | - | 9237 | HECT domain a | 0.18336 | 4.56277 | 7.71362 | 0.0072908 | 0.039667 |
| WNT3 | SCHR17_2 | 4.7E+07 | 4.7E+07 | - | 4266 | Wnt family me | 0.311 | 2.936 | 7.71171 | 0.0072976 | 0.039689 |
| AC092490 | 12 | 8788257 | 8795789 | + | 2279 | novel transcrip | -0.093 | 6.90894 | 7.7106 | 0.0073016 | 0.039696 |
| USP21 | 1 | 1.6E+08 | 1.6E+08 | + | 3764 | ubiquitin speci | 0.18536 | 4.57605 | 7.70189 | 0.0073328 | 0.03984 |
| CITED4 | 1 | 4.1E+07 | 4.1E+07 | - | 1316 | Cbp/p300 inte | 0.23216 | 3.87844 | 7.70168 | 0.0073336 | 0.03984 |
| RAB4A | 1 | 2.3E+08 | 2.3E+08 | + | 3310 | RAB4A, memb | 0.14486 | 5.39257 | 7.69924 | 0.0073424 | 0.039872 |
| RPL37 | 5 | 4.1E+07 | 4.1E+07 | - | 8040 | ribosomal prot | -0.0698 | 9.1195 | 7.6927 | 0.0073659 | 0.039985 |
| QARS | 3 | 4.9E+07 | 4.9E+07 | - | 4703 | glutaminyl-tRN | -0.0729 | 8.35585 | 7.69078 | 0.0073729 | 0.040008 |
| NDUFA9 | 12 | 4649095 | 4694317 | + | 10391 | NADH:ubiquin | -0.2395 | 3.80747 | 7.68643 | 0.0073886 | 0.040071 |
| PEX16 | 11 | 4.6E+07 | 4.6E+07 | - | 3019 | peroxisomal bi | -0.1708 | 4.81605 | 7.68603 | 0.0073901 | 0.040071 |
| TRIM38 | 6 | 2.6E+07 | 2.6E+07 | + | 9418 | tripartite motif | -0.4428 | 2.15986 | 7.71418 | 0.0073939 | 0.040077 |
| MAGED1 | X | 5.2E+07 | 5.2E+07 | + | 6016 | MAGE family n | -0.0823 | 7.76081 | 7.67766 | 0.0074205 | 0.040205 |
| SPG7 | 16 | 8.9E+07 | 9E+07 | + | 36008 | SPG7, parapleg | 0.10167 | 6.58301 | 7.67132 | 0.0074436 | 0.040316 |
| ITGA1 | 5 | 5.3E+07 | 5.3E+07 | + | 18566 | integrin subun | 0.21696 | 4.09988 | 7.66828 | 0.0074547 | 0.04036 |
| EHD4 | 15 | 4.2E+07 | 4.2E+07 | - | 7864 | EH domain cor | 0.11469 | 6.09402 | 7.66497 | 0.0074668 | 0.040411 |
| SCPEP1 | 17 | 5.7E+07 | 5.7E+07 | + | 3853 | serine carboxy | -0.1417 | 5.4398 | 7.66173 | 0.0074787 | 0.04046 |

| | | | | | | | | | | | |
|---|---|---|---|---|---|---|---|---|---|---|---|
| MTMR10 | 15 | 3.1E+07 | 3.1E+07 | - | 8838 | myotubularin | -0.1883 | 4.59976 | 7.64862 | 0.007527 | 0.040706 |
| AC021078 | 5 | 1.5E+08 | 1.5E+08 | - | 8636 | novel transcrip | 0.17861 | 4.6536 | 7.64572 | 0.0075377 | 0.040748 |
| AP001107 | 11 | 6.6E+07 | 6.6E+07 | + | 2552 | novel transcrip | 0.64584 | 0.93762 | 7.6399 | 0.0075593 | 0.04085 |
| PNPLA8 | 7 | 1.1E+08 | 1.1E+08 | - | 5673 | patatin like ph | -0.1837 | 4.59261 | 7.63292 | 0.0075852 | 0.040975 |
| CMTM6 | 3 | 3.2E+07 | 3.3E+07 | - | 3857 | CKLF like MAR | -0.1013 | 6.50408 | 7.63052 | 0.0075942 | 0.041007 |
| MEST | 7 | 1.3E+08 | 1.3E+08 | + | 4794 | mesoderm spe | 0.0982 | 6.78411 | 7.6296 | 0.0075976 | 0.041011 |
| RTN4IP1 | 6 | 1.1E+08 | 1.1E+08 | - | 1974 | reticulon 4 inte | -0.222 | 4.08401 | 7.62377 | 0.0076194 | 0.041113 |
| CEBPZOS | 2 | 3.7E+07 | 3.7E+07 | + | 4700 | CEBPZ opposit | -0.1304 | 5.68971 | 7.6226 | 0.0076238 | 0.041121 |
| TP53 | 17 | 7661779 | 7687550 | - | 5688 | tumor protein | -0.0833 | 7.45198 | 7.6174 | 0.0076433 | 0.041211 |
| SRP54 | 14 | 3.5E+07 | 3.5E+07 | + | 3903 | signal recognit | -0.1234 | 6.03081 | 7.63236 | 0.0076823 | 0.041391 |
| JUN | 1 | 5.9E+07 | 5.9E+07 | - | 3540 | Jun proto-onco | 0.21203 | 4.52799 | 7.85634 | 0.0076833 | 0.041391 |
| HACD4 | 9 | 2.1E+07 | 2.1E+07 | - | 9162 | 3-hydroxyacyl- | 0.46778 | 1.78718 | 7.60487 | 0.0076905 | 0.041391 |
| RPL10 | X | 1.5E+08 | 1.5E+08 | + | 4825 | ribosomal prot | 0.06419 | 9.50328 | 7.60482 | 0.0076907 | 0.041391 |
| CENPU | 4 | 1.8E+08 | 1.8E+08 | - | 3483 | centromere pr | -0.1167 | 6.06154 | 7.6047 | 0.0076911 | 0.041391 |
| DHX37 | 12 | 1.2E+08 | 1.2E+08 | - | 6627 | DEAH-box heli | -0.1013 | 6.50142 | 7.59454 | 0.0077296 | 0.041583 |
| GRM4 | 6 | 3.4E+07 | 3.4E+07 | - | 16300 | glutamate met | -0.1229 | 5.87756 | 7.59348 | 0.0077336 | 0.041589 |
| S100A16 | 1 | 1.5E+08 | 1.5E+08 | - | 1701 | S100 calcium b | 0.19253 | 4.41085 | 7.58577 | 0.007763 | 0.041729 |
| RPL17 | 18 | 4.9E+07 | 4.9E+07 | - | 2466 | ribosomal prot | -0.1162 | 6.12102 | 7.5851 | 0.0077656 | 0.041729 |
| LIPG | 18 | 5E+07 | 5E+07 | + | 13089 | lipase G, endot | -0.2439 | 3.71843 | 7.58047 | 0.0077833 | 0.041809 |
| ZNF135 | 19 | 5.8E+07 | 5.8E+07 | + | 6418 | zinc finger prot | 0.34983 | 2.68326 | 7.57902 | 0.0077888 | 0.041823 |
| C5orf24 | 5 | 1.3E+08 | 1.3E+08 | + | 5759 | chromosome 5 | 0.16101 | 5.04215 | 7.57718 | 0.0077959 | 0.041828 |
| ZFAND4 | 10 | 4.6E+07 | 4.6E+07 | - | 6232 | zinc finger AN1 | -0.2803 | 3.29729 | 7.57703 | 0.0077965 | 0.041828 |
| GRHPR | 9 | 3.7E+07 | 3.7E+07 | + | 7226 | glyoxylate and | 0.10409 | 6.46336 | 7.57595 | 0.0078006 | 0.041828 |
| SEMA4F | 2 | 7.5E+07 | 7.5E+07 | + | 6227 | ssemaphorin 4 | 0.2635 | 3.42663 | 7.57573 | 0.0078015 | 0.041828 |
| RPL6 | 12 | 1.1E+08 | 1.1E+08 | - | 4447 | ribosomal prot | -0.0633 | 9.58063 | 7.56999 | 0.0078235 | 0.041931 |
| FAM45A | 10 | 1.2E+08 | 1.2E+08 | + | 6291 | family with sec | -0.1474 | 5.4472 | 7.57752 | 0.0078286 | 0.041942 |
| HIP1R | 12 | 1.2E+08 | 1.2E+08 | + | 7072 | huntingtin inte | -0.2315 | 3.93567 | 7.56663 | 0.0078365 | 0.041969 |
| HNRNPH3 | 10 | 6.8E+07 | 6.8E+07 | + | 5373 | heterogeneous | -0.0728 | 8.32628 | 7.56305 | 0.0078503 | 0.042016 |
| RASSF9 | 12 | 8.6E+07 | 8.6E+07 | - | 5515 | Ras associatior | 0.80761 | 0.38951 | 7.56282 | 0.0078511 | 0.042016 |
| CHTF18 | 16 | 788046 | 800737 | + | 5916 | chromosome t | 0.12994 | 5.73192 | 7.55812 | 0.0078693 | 0.042098 |
| GPX1 | 3 | 4.9E+07 | 4.9E+07 | - | 1183 | glutathione pe | 0.08983 | 7.17713 | 7.55627 | 0.0078765 | 0.04212 |
| TUBB3 | 16 | 9E+07 | 9E+07 | + | 4430 | tubulin beta 3 | 0.26714 | 3.46038 | 7.55471 | 0.0078826 | 0.042137 |
| NXT2 | X | 1.1E+08 | 1.1E+08 | + | 3064 | nuclear transp | -0.2116 | 4.16324 | 7.53166 | 0.0079725 | 0.042602 |
| POLR3E | 16 | 2.2E+07 | 2.2E+07 | + | 9486 | RNA polymera | -0.0979 | 6.6236 | 7.52041 | 0.0080168 | 0.042823 |
| PNISR | 6 | 9.9E+07 | 9.9E+07 | - | 10871 | PNN interactin | 0.08046 | 7.98513 | 7.54056 | 0.0080254 | 0.042853 |
| VPS37C | 11 | 6.1E+07 | 6.1E+07 | - | 3957 | VPS37C, ESCRT | -0.1608 | 5.04273 | 7.51325 | 0.0080451 | 0.042942 |
| AL139158 | 1 | 3.8E+07 | 3.8E+07 | + | 684 | novel transcrip | 0.55384 | 1.42429 | 7.51167 | 0.0080514 | 0.042956 |
| OAT | 10 | 1.2E+08 | 1.2E+08 | - | 3388 | ornithine amin | -0.0955 | 6.75923 | 7.51107 | 0.0080538 | 0.042956 |
| GPC3 | X | 1.3E+08 | 1.3E+08 | - | 2711 | glypican 3 [Sou | -0.1436 | 5.56836 | 7.54424 | 0.0080579 | 0.042963 |
| ARHGEF6 | X | 1.4E+08 | 1.4E+08 | - | 5278 | Rac/Cdc42 gua | -0.2542 | 3.52588 | 7.50875 | 0.008063 | 0.042963 |
| CDC42EP5 | 19 | 5.4E+07 | 5.4E+07 | - | 935 | CDC42 effecto | -0.3103 | 3.06866 | 7.50852 | 0.0080639 | 0.042963 |
| SERPINH1 | 11 | 7.6E+07 | 7.6E+07 | + | 6281 | serpin family H | -0.064 | 9.57467 | 7.50696 | 0.0080701 | 0.04298 |
| LRCH1 | 13 | 4.7E+07 | 4.7E+07 | + | 8610 | leucine rich re | 0.21388 | 4.10359 | 7.5039 | 0.0080823 | 0.043029 |

| | | | | | | | | | | | |
|---|---|---|---|---|---|---|---|---|---|---|---|
| PMS2CL | 7 | 6710128 | 6753862 | + | 3879 | PMS2 C-termir | 0.33757 | 2.76483 | 7.49944 | 0.0081001 | 0.043094 |
| ARMC1 | 8 | 6.6E+07 | 6.6E+07 | - | 3419 | armadillo repe | -0.1177 | 5.97724 | 7.4993 | 0.0081006 | 0.043094 |
| CKS2 | 9 | 8.9E+07 | 8.9E+07 | + | 616 | CDC28 protein | -0.0901 | 7.01879 | 7.49857 | 0.0081035 | 0.043094 |
| SDC1 | 2 | 2E+07 | 2E+07 | - | 3613 | syndecan 1 [Sc | 0.10344 | 6.37847 | 7.49285 | 0.0081264 | 0.043199 |
| GAR1 | 4 | 1.1E+08 | 1.1E+08 | + | 1964 | GAR1 ribonucl | -0.1605 | 5.00696 | 7.49078 | 0.0081347 | 0.043225 |
| GTF2A1 | 14 | 8.1E+07 | 8.1E+07 | - | 6826 | general transcı | -0.1278 | 5.70613 | 7.49014 | 0.0081373 | 0.043225 |
| MCM2 | 3 | 1.3E+08 | 1.3E+08 | + | 4715 | minichromoso | 0.08203 | 7.48274 | 7.47809 | 0.0081858 | 0.043467 |
| ENDOD1 | 11 | 9.5E+07 | 9.5E+07 | + | 4651 | endonuclease | 0.31723 | 2.84664 | 7.47504 | 0.0081981 | 0.043516 |
| CAND2 | 3 | 1.3E+07 | 1.3E+07 | + | 5587 | cullin associate | 0.20314 | 4.2806 | 7.47424 | 0.0082013 | 0.043517 |
| GMPPB | 3 | 5E+07 | 5E+07 | - | 6742 | GDP-mannose | 0.19207 | 4.36383 | 7.47226 | 0.0082093 | 0.043544 |
| CNST | 1 | 2.5E+08 | 2.5E+08 | + | 5933 | consortin, con | 0.16285 | 4.92637 | 7.46761 | 0.0082281 | 0.043627 |
| SLC12A9 | 7 | 1E+08 | 1E+08 | + | 8901 | solute carrier f | -0.1186 | 5.98381 | 7.45827 | 0.0082662 | 0.043813 |
| MSANTD2 | 11 | 1.2E+08 | 1.2E+08 | - | 4501 | Myb/SANT DN | 0.20773 | 4.11874 | 7.45357 | 0.0082854 | 0.043898 |
| DDA1 | 19 | 1.7E+07 | 1.7E+07 | + | 4593 | DET1 and DDB | 0.13457 | 5.53754 | 7.45039 | 0.0082983 | 0.043951 |
| SGO2 | 2 | 2E+08 | 2E+08 | + | 6336 | shugoshin 2 [S | -0.1432 | 5.47421 | 7.43724 | 0.0083524 | 0.044221 |
| SDR39U1 | 14 | 2.4E+07 | 2.4E+07 | - | 3099 | short chain de | 0.31508 | 2.88421 | 7.43105 | 0.008378 | 0.04434 |
| ARMT1 | 6 | 1.5E+08 | 1.5E+08 | + | 2819 | acidic residue | -0.1073 | 6.29751 | 7.42659 | 0.0083964 | 0.044421 |
| EGR1 | 5 | 1.4E+08 | 1.4E+08 | + | 3137 | early growth re | -0.6469 | 4.62878 | 17.2148 | 0.0084037 | 0.044444 |
| ZMAT4 | 8 | 4.1E+07 | 4.1E+07 | - | 3691 | zinc finger mat | -0.2179 | 3.99135 | 7.42208 | 0.0084151 | 0.044487 |
| SERPINB6 | 6 | 2948159 | 2972165 | - | 13201 | serpin family B | 0.19314 | 4.55819 | 7.45056 | 0.0084211 | 0.044487 |
| MAD2L1 | 4 | 1.2E+08 | 1.2E+08 | - | 5517 | mitotic arrest | 0.09071 | 6.98487 | 7.42062 | 0.0084212 | 0.044487 |
| METTL9 | 16 | 2.2E+07 | 2.2E+07 | + | 6953 | methyltransfer | -0.0872 | 7.091 | 7.41599 | 0.0084405 | 0.044572 |
| SGSM1 | 22 | 2.5E+07 | 2.5E+07 | + | 8607 | small G protein | -0.2979 | 3.09763 | 7.40219 | 0.0084982 | 0.044861 |
| TBC1D24 | 16 | 2475051 | 2509560 | + | 12213 | TBC1 domain f | -0.2102 | 4.19423 | 7.38783 | 0.0085587 | 0.045147 |
| DLG3 | X | 7E+07 | 7.1E+07 | + | 7870 | discs large MA | -0.1132 | 6.20305 | 7.3861 | 0.008566 | 0.045169 |
| KYAT3 | 1 | 8.9E+07 | 8.9E+07 | - | 2196 | kynurenine am | -0.1318 | 5.60311 | 7.38494 | 0.008571 | 0.045178 |
| DCAF11 | 14 | 2.4E+07 | 2.4E+07 | + | 6350 | DDB1 and CUL | -0.1011 | 6.48906 | 7.37925 | 0.0085951 | 0.045289 |
| BEST2 | 19 | 1.3E+07 | 1.3E+07 | + | 2646 | bestrophin 2 [ | 0.44126 | 1.96143 | 7.37185 | 0.0086266 | 0.045438 |
| GK | X | 3.1E+07 | 3.1E+07 | + | 5359 | glycerol kinase | -0.2936 | 3.20698 | 7.37082 | 0.008631 | 0.045445 |
| NFATC2 | 20 | 5.1E+07 | 5.2E+07 | - | 8286 | nuclear factor | -0.6687 | 0.87441 | 7.36883 | 0.0086395 | 0.045461 |
| ARHGEF39 | 9 | 3.6E+07 | 3.6E+07 | - | 6405 | Rho guanine n | 0.21804 | 3.9406 | 7.36796 | 0.0086432 | 0.045461 |
| SEC24C | 10 | 7.4E+07 | 7.4E+07 | + | 5353 | SEC24 homolo | 0.10947 | 6.39135 | 7.3947 | 0.0086445 | 0.045461 |
| MFSD13A | 10 | 1E+08 | 1E+08 | + | 3526 | major facilitato | 0.29916 | 3.03836 | 7.36657 | 0.0086492 | 0.045461 |
| RAD51AP | 12 | 4538798 | 4560048 | + | 2558 | RAD51 associa | -0.1343 | 5.5445 | 7.3664 | 0.0086499 | 0.045461 |
| AL121885 | 22 | 2.8E+07 | 2.8E+07 | - | 819 | novel transcrip | -0.4336 | 1.97371 | 7.36296 | 0.0086646 | 0.045522 |
| PSKH1 | 16 | 6.8E+07 | 6.8E+07 | + | 3993 | protein serine | 0.14127 | 5.54621 | 7.3702 | 0.0086963 | 0.045671 |
| BPHL | 6 | 3118374 | 3153578 | + | 5780 | biphenyl hydro | 0.20849 | 4.08681 | 7.35088 | 0.0087166 | 0.045761 |
| GRM8 | 7 | 1.3E+08 | 1.3E+08 | - | 6457 | glutamate met | 0.56054 | 1.28668 | 7.34867 | 0.0087261 | 0.045779 |
| RAB11FIP | 2 | 7.3E+07 | 7.3E+07 | - | 7058 | RAB11 family i | 0.15022 | 5.20041 | 7.34863 | 0.0087263 | 0.045779 |
| ATG7 | 3 | 1.1E+07 | 1.2E+07 | + | 8833 | autophagy rela | 0.23287 | 3.87292 | 7.34433 | 0.0087448 | 0.045843 |
| ZNRD1AS | R6_MHC_ | 3E+07 | 3E+07 | - | 17538 | zinc ribbon do | -0.3298 | 2.83608 | 7.33874 | 0.0087691 | 0.045953 |
| TRPM6 | 9 | 7.5E+07 | 7.5E+07 | - | 9193 | transient recep | -0.4629 | 1.83819 | 7.33277 | 0.008795 | 0.046072 |
| TMEM131 | 4 | 1.5E+08 | 1.5E+08 | + | 6393 | transmembran | 0.12144 | 5.85027 | 7.33182 | 0.0087992 | 0.046077 |

| | | | | | | | | | | | |
|---|---|---|---|---|---|---|---|---|---|---|---|
| MRPL18 | 6 | 1.6E+08 | 1.6E+08 | + | 1374 | mitochondrial | -0.1238 | 5.84484 | 7.31614 | 0.0088678 | 0.046419 |
| IFFO1 | 12 | 6538375 | 6556083 | - | 5203 | intermediate f | 0.23396 | 3.72492 | 7.30862 | 0.0089008 | 0.046575 |
| MAST4 | 5 | 6.7E+07 | 6.7E+07 | + | 14675 | microtubule as | 0.29233 | 3.20581 | 7.30614 | 0.0089136 | 0.046611 |
| SSC4D | 7 | 7.6E+07 | 7.6E+07 | - | 2821 | scavenger rece | 0.24456 | 3.58195 | 7.30484 | 0.0089175 | 0.046611 |
| CD3EAP | 19 | 4.5E+07 | 4.5E+07 | + | 3661 | CD3e molecule | -0.2094 | 4.0994 | 7.30451 | 0.008919 | 0.046611 |
| DRD4 | 11 | 637293 | 640706 | + | 1477 | dopamine rece | -0.5313 | 1.47105 | 7.30415 | 0.0089205 | 0.046611 |
| ARHGAP2 | 1 | 9.4E+07 | 9.4E+07 | - | 18370 | Rho GTPase ac | -0.2737 | 3.31074 | 7.29798 | 0.0089478 | 0.046736 |
| KCNK6 | 19 | 3.8E+07 | 3.8E+07 | + | 6168 | potassium two | -0.1884 | 4.4555 | 7.29343 | 0.0089681 | 0.0468 |
| SEC14L5 | 16 | 4958330 | 5019157 | + | 6463 | SEC14 like lipid | -0.3137 | 2.95031 | 7.29315 | 0.0089693 | 0.0468 |
| DDX49 | 19 | 1.9E+07 | 1.9E+07 | + | 3306 | DEAD-box heli | 0.14329 | 5.42162 | 7.29297 | 0.0089701 | 0.0468 |
| AC069277 | 3 | 6490479 | 6736129 | + | 3023 | novel transcrip | 0.66719 | 0.95798 | 7.29232 | 0.008973 | 0.0468 |
| SKI | 1 | 2228319 | 2310213 | + | 6550 | SKI proto-onco | -0.0949 | 6.90042 | 7.29142 | 0.008977 | 0.046804 |
| RCN2 | 15 | 7.7E+07 | 7.7E+07 | + | 9901 | reticulocalbin 2 | 0.08178 | 7.50176 | 7.28673 | 0.0089979 | 0.046895 |
| GOLGA6L | SCHR15_5 | 8.2E+07 | 8.2E+07 | - | 4715 | golgin A6 famil | 0.57318 | 1.18706 | 7.2857 | 0.0090025 | 0.046902 |
| VEGFA | 6 | 4.4E+07 | 4.4E+07 | + | 14431 | vascular endot | -0.1243 | 5.78622 | 7.28386 | 0.0090107 | 0.046928 |
| FAHD2CP | 2 | 9.6E+07 | 9.6E+07 | + | 2140 | fumarylacetoa | 0.40592 | 2.12861 | 7.28293 | 0.0090148 | 0.046933 |
| TDRD7 | 9 | 9.7E+07 | 9.7E+07 | + | 4174 | tudor domain c | 0.35034 | 2.66106 | 7.28581 | 0.0090376 | 0.047034 |
| ARSD | X | 2903972 | 2929349 | - | 7026 | arylsulfatase D | 0.39489 | 2.17864 | 7.27621 | 0.0090449 | 0.047055 |
| KDELR3 | 22 | 3.8E+07 | 3.8E+07 | + | 1913 | KDEL endoplas | 0.35407 | 2.72381 | 7.33934 | 0.00905 | 0.047056 |
| ATP1A3 | 19 | 4.2E+07 | 4.2E+07 | - | 5989 | ATPase Na+/K- | -0.3265 | 2.81065 | 7.27473 | 0.0090515 | 0.047056 |
| PPP4R3A | 14 | 9.1E+07 | 9.2E+07 | - | 5912 | protein phosph | -0.087 | 7.11538 | 7.274 | 0.0090548 | 0.047056 |
| TRANK1 | 3 | 3.7E+07 | 3.7E+07 | - | 12174 | tetratricopepti | 0.17036 | 4.75997 | 7.27264 | 0.0090609 | 0.047057 |
| AP3M1 | 10 | 7.4E+07 | 7.4E+07 | - | 5912 | adaptor relate | 0.11044 | 6.16914 | 7.27236 | 0.0090622 | 0.047057 |
| PAIP2 | 5 | 1.4E+08 | 1.4E+08 | + | 3158 | poly(A) binding | -0.0927 | 6.79248 | 7.27176 | 0.0090649 | 0.047057 |
| GLB1L3 | 11 | 1.3E+08 | 1.3E+08 | + | 8945 | galactosidase b | 0.12498 | 5.74818 | 7.26776 | 0.0090829 | 0.047121 |
| ALPG | 2 | 2.3E+08 | 2.3E+08 | + | 2492 | alkaline phosp | -0.5436 | 1.40367 | 7.26717 | 0.0090855 | 0.047121 |
| IKBIP | 12 | 9.9E+07 | 9.9E+07 | - | 4163 | IKBKB interacti | 0.19666 | 4.26539 | 7.26684 | 0.009087 | 0.047121 |
| PPP2R3A | 3 | 1.4E+08 | 1.4E+08 | + | 7785 | protein phosph | -0.1673 | 4.86519 | 7.26577 | 0.0090919 | 0.047129 |
| CDK17 | 12 | 9.6E+07 | 9.6E+07 | - | 6467 | cyclin depende | 0.16919 | 4.80984 | 7.26381 | 0.0091007 | 0.047157 |
| TIMM23 | 10 | 4.6E+07 | 4.6E+07 | + | 1190 | translocase of | -0.098 | 6.54507 | 7.26226 | 0.0091077 | 0.047177 |
| DCX | X | 1.1E+08 | 1.1E+08 | - | 10203 | doublecortin [S | -0.2758 | 3.29067 | 7.25875 | 0.0091235 | 0.047242 |
| PCMTD1 | 8 | 5.2E+07 | 5.2E+07 | - | 7323 | protein-L-isoas | 0.15568 | 5.11745 | 7.25724 | 0.0091304 | 0.047255 |
| SARS | 1 | 1.1E+08 | 1.1E+08 | + | 3849 | seryl-tRNA syn | -0.0846 | 7.17945 | 7.25674 | 0.0091327 | 0.047255 |
| RCL1 | 9 | 4792944 | 4885917 | + | 4140 | RNA terminal p | 0.18911 | 4.51859 | 7.25596 | 0.0091362 | 0.047256 |
| AC009446 | 8 | 7.2E+07 | 7.2E+07 | + | 901 | novel transcrip | -0.1104 | 6.12892 | 7.24943 | 0.0091658 | 0.047393 |
| ASAP1 | 8 | 1.3E+08 | 1.3E+08 | - | 8294 | ArfGAP with SH | 0.12625 | 5.68746 | 7.24714 | 0.0091763 | 0.047414 |
| PSMB6 | 17 | 4796144 | 4798502 | + | 1447 | proteasome su | -0.0804 | 7.51597 | 7.24706 | 0.0091766 | 0.047414 |
| FEN1 | 11 | 6.2E+07 | 6.2E+07 | + | 2198 | flap structure-s | 0.0903 | 6.88319 | 7.24448 | 0.0091884 | 0.047451 |
| CNTNAP1 | 17 | 4.3E+07 | 4.3E+07 | + | 6306 | contactin asso | 0.27847 | 3.22276 | 7.24373 | 0.0091918 | 0.047451 |
| RCBTB1 | 13 | 5E+07 | 5E+07 | - | 4509 | RCC1 and BTB | 0.1164 | 6.00896 | 7.24331 | 0.0091937 | 0.047451 |
| ABCF1 | R6_MHC_s | 3.1E+07 | 3.1E+07 | + | 4496 | ATP binding ca | -0.0757 | 7.87637 | 7.24211 | 0.0091992 | 0.047462 |
| MAF | 16 | 8E+07 | 8E+07 | - | 6999 | MAF bZIP trans | -0.283 | 3.18171 | 7.23725 | 0.0092214 | 0.04756 |
| TRMT6 | 20 | 5937228 | 5950558 | - | 3783 | tRNA methyltr | -0.1704 | 4.77034 | 7.23069 | 0.0092515 | 0.047698 |

| | | | | | | | | | | | |
|---|---|---|---|---|---|---|---|---|---|---|---|
| ZNF689 | 16 | 3.1E+07 | 3.1E+07 | - | 4159 | zinc finger prot | 0.14967 | 5.14135 | 7.22822 | 0.0092628 | 0.047722 |
| ACAT1 | 11 | 1.1E+08 | 1.1E+08 | + | 5584 | acetyl-CoA ace | 0.15001 | 5.16252 | 7.22672 | 0.0092697 | 0.047723 |
| MAGI2 | 7 | 7.8E+07 | 7.9E+07 | - | 31624 | membrane ass | 0.20753 | 4.17275 | 7.22663 | 0.0092701 | 0.047723 |
| AC012513 | 2 | 2.2E+08 | 2.2E+08 | + | 2116 | TEC | 0.41468 | 2.15522 | 7.22601 | 0.009273 | 0.047723 |
| LRPAP1 | 4 | 3503612 | 3532446 | - | 11816 | LDL receptor re | 0.09817 | 6.56544 | 7.22321 | 0.0092859 | 0.047771 |
| RBMS3 | 3 | 2.9E+07 | 3E+07 | + | 18339 | RNA binding m | 0.354 | 2.47681 | 7.22255 | 0.0092889 | 0.047771 |
| GRAMD2A | 15 | 7.2E+07 | 7.2E+07 | - | 4990 | GRAM domain | 0.7117 | 0.68537 | 7.22162 | 0.0092932 | 0.047776 |
| TACC3 | 4 | 1712891 | 1745176 | + | 8202 | transforming a | 0.07584 | 7.94984 | 7.21937 | 0.0093036 | 0.047808 |
| AHCY | 20 | 3.4E+07 | 3.4E+07 | - | 3894 | adenosylhomo | -0.0664 | 8.95066 | 7.21885 | 0.009306 | 0.047808 |
| STK39 | 2 | 1.7E+08 | 1.7E+08 | - | 3701 | serine/threoni | 0.13248 | 5.59245 | 7.21795 | 0.0093101 | 0.047812 |
| LINC-ROR | 18 | 5.7E+07 | 5.7E+07 | - | 3891 | long intergenic | -0.4356 | 1.96702 | 7.21661 | 0.0093163 | 0.047827 |
| EPB41L4A | 5 | 1.1E+08 | 1.1E+08 | - | 8579 | erythrocyte me | 0.33347 | 2.65294 | 7.21553 | 0.0093213 | 0.047835 |
| ADAM11 | 17 | 4.5E+07 | 4.5E+07 | + | 4813 | ADAM metallo | 0.37959 | 2.31323 | 7.21389 | 0.0093289 | 0.047857 |
| TKT | 3 | 5.3E+07 | 5.3E+07 | - | 9502 | transketolase [ | 0.0597 | 10.4117 | 7.20494 | 0.0093705 | 0.048053 |
| MICAL3 | 22 | 1.8E+07 | 1.8E+07 | - | 22356 | microtubule as | -0.1517 | 5.14616 | 7.20335 | 0.0093779 | 0.048064 |
| PCDHB16 | 5 | 1.4E+08 | 1.4E+08 | + | 5001 | protocadherin | 0.39943 | 2.22928 | 7.20304 | 0.0093794 | 0.048064 |
| KDM1A | 1 | 2.3E+07 | 2.3E+07 | + | 5440 | lysine demethy | 0.06653 | 8.90659 | 7.19703 | 0.0094074 | 0.048191 |
| AC115223 | 4 | 6.6E+07 | 6.6E+07 | + | 870 | ribosomal prot | -0.2035 | 4.15602 | 7.1923 | 0.0094295 | 0.048287 |
| PQLC1 | 18 | 8E+07 | 8E+07 | - | 5218 | PQ loop repeat | 0.22962 | 3.82315 | 7.19111 | 0.0094351 | 0.048298 |
| MPV17L | SCHR16_1 | 1.5E+07 | 1.5E+07 | + | 6012 | MPV17 mitoch | -0.3625 | 2.63534 | 7.23238 | 0.0094775 | 0.048498 |
| FCF1 | 14 | 7.5E+07 | 7.5E+07 | + | 4747 | FCF1, rRNA-pro | -0.1151 | 6.05935 | 7.17802 | 0.0094967 | 0.048579 |
| ARF5 | 7 | 1.3E+08 | 1.3E+08 | + | 1825 | ADP ribosylatio | 0.09222 | 6.87277 | 7.17591 | 0.0095066 | 0.048612 |
| MAL2 | 8 | 1.2E+08 | 1.2E+08 | + | 3438 | mal, T cell diffe | 0.08848 | 6.96972 | 7.17167 | 0.0095267 | 0.048697 |
| KIF4A | X | 7E+07 | 7E+07 | + | 4573 | kinesin family | 0.09619 | 6.68175 | 7.16974 | 0.0095358 | 0.048727 |
| SHB | 9 | 3.8E+07 | 3.8E+07 | - | 6035 | SH2 domain co | -0.2486 | 3.55168 | 7.1669 | 0.0095493 | 0.048779 |
| KIF21A | 12 | 3.9E+07 | 3.9E+07 | - | 10117 | kinesin family | 0.13179 | 5.69283 | 7.16406 | 0.0095775 | 0.048899 |
| PDRG1 | 20 | 3.2E+07 | 3.2E+07 | - | 1957 | p53 and DNA d | -0.1757 | 4.6289 | 7.15996 | 0.0095823 | 0.048899 |
| CIB2 | 15 | 7.8E+07 | 7.8E+07 | - | 2146 | calcium and in | 0.13282 | 5.55401 | 7.1598 | 0.0095831 | 0.048899 |
| ACTR1B | 2 | 9.8E+07 | 9.8E+07 | - | 2892 | ARP1 actin rela | -0.1268 | 5.74501 | 7.1545 | 0.0096084 | 0.04901 |
| CKAP2L | 2 | 1.1E+08 | 1.1E+08 | - | 5464 | cytoskeleton a | -0.154 | 5.07209 | 7.1386 | 0.0096847 | 0.049382 |
| PRMT6 | 1 | 1.1E+08 | 1.1E+08 | + | 4335 | protein argini | 0.13408 | 5.48069 | 7.13691 | 0.0096928 | 0.049406 |
| TBL2 | 7 | 7.4E+07 | 7.4E+07 | - | 6466 | transducin bet | 0.20466 | 4.08249 | 7.13286 | 0.0097123 | 0.049481 |
| PKP3 | 11 | 392614 | 404908 | + | 3795 | plakophilin 3 [S | 0.12517 | 5.71279 | 7.13243 | 0.0097145 | 0.049481 |
| GNPTAB | 12 | 1E+08 | 1E+08 | - | 9774 | N-acetylglucos | -0.1005 | 6.54225 | 7.12808 | 0.0097355 | 0.049561 |
| LINC0236 | 12 | 1.3E+08 | 1.3E+08 | - | 1484 | long intergenic | 0.64731 | 0.92402 | 7.12776 | 0.009737 | 0.049561 |
| ARL16 | 17 | 8.2E+07 | 8.2E+07 | - | 2597 | ADP ribosylatio | 0.21533 | 3.98724 | 7.12657 | 0.0097428 | 0.049573 |
| FBLN5 | 14 | 9.2E+07 | 9.2E+07 | - | 4973 | fibulin 5 [Sourc | 0.3717 | 2.31837 | 7.12271 | 0.0097615 | 0.049651 |
| SEMA4C | 2 | 9.7E+07 | 9.7E+07 | - | 6264 | semaphorin 4C | -0.0859 | 7.09394 | 7.1185 | 0.009782 | 0.049715 |
| MYO5C | 15 | 5.2E+07 | 5.2E+07 | - | 10051 | myosin VC [Sou | 0.16218 | 4.87102 | 7.11802 | 0.0097843 | 0.049715 |
| CAPN11 | 6 | 4.4E+07 | 4.4E+07 | + | 3548 | calpain 11 [Sou | 0.55152 | 1.32347 | 7.118 | 0.0097845 | 0.049715 |
| MPP2 | 17 | 4.4E+07 | 4.4E+07 | - | 6114 | membrane pal | 0.12317 | 5.7426 | 7.11614 | 0.0097935 | 0.049741 |
| IRAK1 | X | 1.5E+08 | 1.5E+08 | - | 3917 | interleukin 1 re | 0.08708 | 7.06763 | 7.11553 | 0.0097965 | 0.049741 |

**Table S2. EC DEGs**

| geneid | chr | start | end | strand | length | description | logFC | logCPM | F | PValue | FDR |
|---|---|---|---|---|---|---|---|---|---|---|---|
| XIST | X | 73820651 | 73852723 | - | 25264 | X inactive s | 9.602614 | 9.923947 | 5362.733 | 1.61E-12 | 2.28E-08 |
| NID2 | 14 | 52004803 | 52069228 | - | 6392 | nidogen 2 [ | -1.297187 | 8.968 | 255.7269 | 2.55E-07 | 0.00180554 |
| LINC02381 | 12 | 54126098 | 54147485 | + | 4253 | long interge | 1.800401 | 3.178603 | 216.1175 | 4.88E-07 | 0.00230133 |
| CRHBP | 5 | 76953045 | 76981158 | + | 2468 | corticotrop | -2.993056 | 5.228362 | 214.0453 | 1.54E-06 | 0.00545654 |
| UNC5C | 4 | 95162504 | 95549206 | - | 10685 | unc-5 netri | 1.90636 | 5.697722 | 148.3631 | 3.09E-06 | 0.00807005 |
| CADM4 | 19 | 43622368 | 43639850 | - | 2257 | cell adhesio | 1.133709 | 5.222936 | 129.5218 | 3.42E-06 | 0.00807005 |
| CACNG8 | 19 | 53963040 | 53990215 | + | 8747 | calcium vol | -1.174138 | 4.261141 | 114.527 | 5.43E-06 | 0.0109715 |
| USP16 | 21 | 29024629 | 29054488 | + | 5694 | ubiquitin sp | -0.835512 | 5.48051 | 106.3141 | 7.16E-06 | 0.01266936 |
| ETS2 | 21 | 38805307 | 38824955 | + | 4497 | ETS proto-o | -0.892996 | 4.938304 | 97.78414 | 9.77E-06 | 0.01377666 |
| AK5 | 1 | 77282019 | 77559966 | + | 6948 | adenylate k | -1.204566 | 2.960036 | 93.26915 | 1.16E-05 | 0.01377666 |
| BTG3 | SCHR21_6_ | 17604145 | 17612947 | - | 2306 | BTG anti-pr | -0.984678 | 5.383749 | 91.97562 | 1.22E-05 | 0.01377666 |
| HOXA13 | 7 | 27193503 | 27200106 | - | 5086 | homeobox | -2.772199 | 1.950253 | 91.70662 | 1.24E-05 | 0.01377666 |
| COL6A1 | 21 | 45981737 | 46005050 | + | 5343 | collagen typ | -0.868296 | 9.718038 | 88.85193 | 1.39E-05 | 0.01377666 |
| TBX20 | 7 | 35202430 | 35254147 | - | 1875 | T-box 20 [S | -2.04082 | 1.326559 | 87.80064 | 1.45E-05 | 0.01377666 |
| CYP2S1 | 19 | 41193210 | 41207539 | + | 2713 | cytochrome | -1.089114 | 3.293752 | 87.6764 | 1.46E-05 | 0.01377666 |
| COL7A1 | 3 | 48564073 | 48595267 | - | 11071 | collagen typ | 0.91539 | 6.448358 | 85.75666 | 1.58E-05 | 0.01400825 |
| CFAP298 | 21 | 32592079 | 32612603 | - | 5403 | cilia and fla | -0.911062 | 3.903695 | 83.16083 | 1.77E-05 | 0.01409774 |
| LINC02593 | 1 | 916865 | 921016 | - | 4152 | long interge | 1.779661 | 2.808058 | 81.91434 | 1.87E-05 | 0.01409774 |
| RNASE1 | 14 | 20801228 | 20802855 | - | 1083 | ribonucleas | -1.476216 | 3.147799 | 81.679 | 1.89E-05 | 0.01409774 |
| TGM2 | 20 | 38127385 | 38166578 | - | 7948 | transglutam | -0.965558 | 8.391789 | 78.96888 | 2.14E-05 | 0.01514638 |
| CDH13 | 16 | 82626965 | 83800640 | + | 11025 | cadherin 13 | -0.84366 | 6.31602 | 71.4643 | 3.08E-05 | 0.01761807 |
| SUCNR1 | 3 | 1.52E+08 | 1.52E+08 | + | 4181 | succinate re | 3.357441 | 0.171343 | 70.50387 | 3.23E-05 | 0.01761807 |
| KRT19 | 17 | 41523617 | 41528308 | - | 2336 | keratin 19 [ | -0.714546 | 8.212065 | 69.3581 | 3.43E-05 | 0.01761807 |
| APP | 21 | 25880550 | 26171128 | - | 6316 | amyloid bet | -0.667404 | 11.08209 | 69.02051 | 3.49E-05 | 0.01761807 |
| BACE2 | 21 | 41167801 | 41282530 | + | 9631 | beta-secret | -1.761229 | 5.035582 | 81.52361 | 3.55E-05 | 0.01761807 |
| PKNOX2 | 11 | 1.25E+08 | 1.25E+08 | + | 6538 | PBX/knotte | 0.9897 | 3.644056 | 68.65601 | 3.55E-05 | 0.01761807 |
| KIAA1522 | 1 | 32741830 | 32774970 | + | 6020 | KIAA1522 [S | 0.867434 | 5.310574 | 68.38173 | 3.60E-05 | 0.01761807 |
| HOXA5 | 7 | 27141052 | 27143681 | - | 1720 | homeobox | 0.86514 | 3.751647 | 68.35746 | 3.61E-05 | 0.01761807 |
| TOX2 | 20 | 43914852 | 44069616 | + | 3235 | TOX high m | -1.01155 | 4.522393 | 68.34838 | 3.61E-05 | 0.01761807 |
| COL15A1 | 9 | 98943179 | 99070792 | + | 6653 | collagen typ | -1.685865 | 2.160932 | 67.06935 | 3.86E-05 | 0.01804457 |
| TMEM59L | 19 | 18607430 | 18621039 | + | 2626 | transmemb | 1.216745 | 2.789175 | 66.64563 | 3.95E-05 | 0.01804457 |
| TTC3 | 21 | 37073226 | 37203112 | + | 15883 | tetratricope | -0.714982 | 8.862235 | 64.31022 | 4.49E-05 | 0.01843821 |
| HMGN1 | 21 | 39342315 | 39349647 | - | 5042 | high mobilit | -0.69782 | 7.7626 | 63.20817 | 4.78E-05 | 0.01843821 |
| FAM207A | 21 | 44940012 | 44976989 | + | 1400 | family with | -0.864011 | 4.911231 | 63.18149 | 4.79E-05 | 0.01843821 |
| COL18A1 | 21 | 45405137 | 45513720 | + | 8492 | collagen typ | -0.796035 | 10.76584 | 62.60266 | 4.95E-05 | 0.01843821 |
| IFNGR2 | 21 | 33402896 | 33479348 | + | 2960 | interferon g | -0.692919 | 5.90559 | 62.08604 | 5.09E-05 | 0.01843821 |
| MLPH | 2 | 2.37E+08 | 2.38E+08 | + | 7435 | melanophil | -1.044879 | 4.50353 | 61.61881 | 5.23E-05 | 0.01843821 |
| CPE | 4 | 1.65E+08 | 1.65E+08 | + | 3119 | carboxypep | 0.70679 | 6.346678 | 61.25322 | 5.34E-05 | 0.01843821 |
| VPS26C | 21 | 37223420 | 37267919 | - | 9301 | VPS26 endo | -0.61939 | 5.598039 | 61.13394 | 5.38E-05 | 0.01843821 |
| HOXA11 | 7 | 27181510 | 27185223 | - | 2307 | homeobox | -0.883718 | 4.354471 | 61.07597 | 5.40E-05 | 0.01843821 |
| HLCS | 21 | 36750888 | 36990236 | - | 7889 | holocarbox | -0.761353 | 5.302137 | 60.94204 | 5.44E-05 | 0.01843821 |

| | | | | | | | | | | | |
|---|---|---|---|---|---|---|---|---|---|---|---|
| CSTB | 21 | 43772511 | 43776445 | - | 3935 | cystatin B [S | -0.82982 | 6.672631 | 60.43738 | 5.61E-05 | 0.01843821 |
| SLIT3 | 5 | 1.69E+08 | 1.69E+08 | - | 14260 | slit guidanc | 0.938133 | 6.717809 | 60.0964 | 5.72E-05 | 0.01843821 |
| CAPN6 | X | 1.11E+08 | 1.11E+08 | - | 3532 | calpain 6 [S | 1.879398 | 1.005937 | 59.49382 | 5.93E-05 | 0.01843821 |
| CCT8 | 21 | 29055805 | 29073797 | - | 3341 | chaperonin | -0.66806 | 7.619503 | 58.59624 | 6.26E-05 | 0.01843821 |
| ACE | 17 | 63477061 | 63498380 | + | 8365 | angiotensin | -1.29096 | 2.97703 | 58.4396 | 6.31E-05 | 0.01843821 |
| COL6A2 | 21 | 46098097 | 46132849 | + | 5350 | collagen typ | -0.951268 | 10.1327 | 58.4106 | 6.33E-05 | 0.01843821 |
| SULF2 | 20 | 47656348 | 47786616 | - | 6657 | sulfatase 2 | -0.642249 | 8.452569 | 58.3072 | 6.37E-05 | 0.01843821 |
| SUMO3 | 21 | 44805617 | 44818779 | - | 2732 | small ubiqu | -0.563645 | 6.712173 | 57.88517 | 6.53E-05 | 0.01843821 |
| PSMG1 | 21 | 39174769 | 39183851 | - | 2515 | proteasome | -0.744484 | 4.322245 | 57.29386 | 6.77E-05 | 0.01843821 |
| IFNAR1 | 21 | 33324429 | 33359864 | + | 6879 | interferon a | -0.722972 | 6.053269 | 57.26394 | 6.79E-05 | 0.01843821 |
| CYYR1 | 21 | 26466209 | 26573255 | - | 3240 | cysteine an | -1.096173 | 3.339893 | 57.13833 | 6.84E-05 | 0.01843821 |
| SEMA3F | 3 | 50155045 | 50189075 | + | 4826 | semaphorin | 0.904532 | 4.471098 | 56.72725 | 7.01E-05 | 0.01843821 |
| TNFRSF21 | 6 | 47231532 | 47309905 | - | 3595 | TNF recepto | 0.712185 | 7.816113 | 56.46634 | 7.13E-05 | 0.01843821 |
| NTF3 | 12 | 5432108 | 5521536 | + | 2923 | neurotroph | -2.570278 | 0.26251 | 56.38273 | 7.17E-05 | 0.01843821 |
| CD34 | 1 | 2.08E+08 | 2.08E+08 | - | 9219 | CD34 molec | -1.277426 | 2.773752 | 55.84927 | 7.41E-05 | 0.01872601 |
| PDE1C | 7 | 31751179 | 32299329 | - | 13821 | phosphodie | 1.611682 | 2.001829 | 54.88182 | 7.88E-05 | 0.01930131 |
| SOD1 | 21 | 31659622 | 31668931 | + | 2019 | superoxide | -0.837215 | 7.438872 | 54.82173 | 7.91E-05 | 0.01930131 |
| FMOD | 1 | 2.03E+08 | 2.03E+08 | - | 3034 | fibromodul | -2.390346 | 0.49223 | 54.37238 | 8.14E-05 | 0.01953093 |
| PAK3 | X | 1.11E+08 | 1.11E+08 | + | 10446 | p21 (RAC1) | 1.249442 | 3.37581 | 54.09317 | 8.29E-05 | 0.01955574 |
| SYNJ1 | 21 | 32628759 | 32728048 | - | 8070 | synaptojani | -0.853486 | 4.329595 | 53.54655 | 8.59E-05 | 0.01993287 |
| SBK1 | 16 | 28292525 | 28323849 | + | 4986 | SH3 domain | 1.168805 | 3.08201 | 53.10396 | 8.85E-05 | 0.01996607 |
| TRIB2 | 2 | 12716889 | 12742734 | + | 4588 | tribbles pse | 0.673092 | 4.508322 | 52.85823 | 8.99E-05 | 0.01996607 |
| CBR3 | 21 | 36135079 | 36146562 | + | 998 | carbonyl re | -1.643919 | 1.440552 | 52.67562 | 9.10E-05 | 0.01996607 |
| USP25 | 21 | 15730025 | 15880069 | + | 7823 | ubiquitin sp | -0.631331 | 4.956031 | 52.55929 | 9.17E-05 | 0.01996607 |
| THBS1 | 15 | 39581079 | 39599466 | + | 9158 | thrombospo | -1.222088 | 9.806965 | 56.38954 | 9.44E-05 | 0.0202439 |
| HOXA7 | 7 | 27153716 | 27157936 | - | 2411 | homeobox | 1.07429 | 2.382395 | 51.20191 | 1.01E-04 | 0.0212239 |
| NRIP1 | 21 | 14961235 | 15065936 | - | 8497 | nuclear rece | -0.672904 | 4.606573 | 50.49689 | 1.05E-04 | 0.021947 |
| TIAM1 | 21 | 31118416 | 31559977 | - | 9216 | T cell lymph | -0.778254 | 4.995907 | 50.22631 | 1.07E-04 | 0.02203723 |
| DIO3 | 14 | 1.02E+08 | 1.02E+08 | + | 2102 | iodothyron | -1.981798 | 1.506333 | 49.34576 | 1.14E-04 | 0.02290583 |
| CLDN6 | 16 | 3014712 | 3020071 | - | 1739 | claudin 6 [S | 2.534177 | 2.658506 | 56.55439 | 1.16E-04 | 0.02290583 |
| NCAM2 | 21 | 20998409 | 21543329 | + | 8286 | neural cell a | -0.991502 | 3.953905 | 49.06594 | 1.17E-04 | 0.02290583 |
| ITSN1 | 21 | 33642400 | 33899861 | + | 21748 | intersectin | -0.56934 | 6.584986 | 48.47185 | 1.22E-04 | 0.02356669 |
| RRP1B | 21 | 43659560 | 43696079 | + | 5750 | ribosomal R | -0.555051 | 6.415478 | 47.96896 | 1.26E-04 | 0.02364804 |
| N6AMT1 | 21 | 28872191 | 28885371 | - | 4865 | N-6 adenine | -0.793147 | 3.448802 | 47.79213 | 1.28E-04 | 0.02364804 |
| LHX1-DT | 17 | 36861674 | 36936661 | - | 2544 | LHX1 diverg | 2.155988 | 1.412431 | 47.69559 | 1.29E-04 | 0.02364804 |
| HECW1 | 7 | 43112629 | 43566001 | + | 13935 | HECT, C2 ar | 2.317517 | 0.960734 | 47.61071 | 1.29E-04 | 0.02364804 |
| HOXC4 | 12 | 54016931 | 54056030 | + | 2767 | homeobox | 1.009348 | 3.034032 | 47.43431 | 1.31E-04 | 0.02364804 |
| MORC3 | 21 | 36320189 | 36386148 | + | 6408 | MORC fami | -0.708146 | 4.69784 | 47.19768 | 1.33E-04 | 0.02364804 |
| AFF3 | 2 | 99545419 | 1E+08 | - | 12737 | AF4/FMR2 f | 1.342686 | 3.260577 | 47.11204 | 1.34E-04 | 0.02364804 |
| LDB2 | 4 | 16501541 | 16898678 | - | 5455 | LIM domain | -0.733979 | 4.159713 | 46.79304 | 1.37E-04 | 0.02364804 |
| LINC00205 | 21 | 45288050 | 45297806 | + | 7619 | long interge | -0.657019 | 6.949057 | 46.75581 | 1.38E-04 | 0.02364804 |
| CPM | 12 | 68842197 | 68971570 | - | 7535 | carboxypep | -1.063796 | 2.725545 | 46.65779 | 1.39E-04 | 0.02364804 |
| GPR37 | 7 | 1.25E+08 | 1.25E+08 | - | 3021 | G protein-c | 0.702985 | 4.919824 | 46.29817 | 1.42E-04 | 0.02399792 |

| | | | | | | | | | | | |
|---|---|---|---|---|---|---|---|---|---|---|---|
| AGPAT3 | 21 | 43865223 | 43987592 | + | 12315 | 1-acylglycer | -0.511277 | 6.661898 | 43.96344 | 1.70E-04 | 0.02825583 |
| PALMD | 1 | 99646113 | 99694541 | + | 8074 | palmdelphi | -1.218742 | 2.825156 | 43.84466 | 1.72E-04 | 0.02825583 |
| GALNT6 | 12 | 51351247 | 51392867 | - | 7347 | polypeptide | -0.702356 | 5.854735 | 43.34157 | 1.79E-04 | 0.02857273 |
| PMP22 | 17 | 15229777 | 15265326 | - | 3191 | peripheral r | -0.769242 | 5.996684 | 43.32689 | 1.79E-04 | 0.02857273 |
| PIANP | 12 | 6693792 | 6700800 | - | 3266 | PILR alpha a | 1.533498 | 1.426429 | 43.10929 | 1.82E-04 | 0.02857273 |
| EVA1C | 21 | 32412006 | 32515397 | + | 4353 | eva-1 homo | -2.234848 | 0.595446 | 42.93672 | 1.84E-04 | 0.02857273 |
| MSI1 | 12 | 1.2E+08 | 1.2E+08 | - | 3074 | musashi RN | 1.176801 | 1.923804 | 42.8545 | 1.86E-04 | 0.02857273 |
| SNTB1 | 8 | 1.21E+08 | 1.21E+08 | - | 6794 | syntrophin | 1.561475 | 0.858634 | 42.68724 | 1.88E-04 | 0.02857273 |
| MAFB | 20 | 40685848 | 40689236 | - | 3389 | MAF bZIP tr | -1.257652 | 3.154852 | 42.60205 | 1.89E-04 | 0.02857273 |
| ARHGAP22 | 10 | 48446034 | 48656265 | - | 6783 | Rho GTPase | -1.409798 | 3.864045 | 45.71107 | 1.90E-04 | 0.02857273 |
| CRYZL1 | 21 | 33589341 | 33643926 | - | 6177 | crystallin ze | -0.602909 | 4.343394 | 42.44471 | 1.92E-04 | 0.02857358 |
| CNTNAP1 | 17 | 42682531 | 42699993 | + | 6306 | contactin as | 0.946419 | 5.128659 | 42.54527 | 2.02E-04 | 0.02945782 |
| HOXB9 | 17 | 48621156 | 48626358 | - | 2586 | homeobox | 0.518217 | 7.638539 | 41.80905 | 2.02E-04 | 0.02945782 |
| LINC01614 | 2 | 2.16E+08 | 2.16E+08 | + | 648 | long interge | -1.161627 | 2.99334 | 41.29408 | 2.11E-04 | 0.03019118 |
| URB1 | 21 | 32311018 | 32393012 | - | 11447 | URB1 ribos | -0.609935 | 5.970208 | 41.25832 | 2.11E-04 | 0.03019118 |
| PFKL | 21 | 44300051 | 44327376 | + | 8926 | phosphofru | -0.663753 | 8.029651 | 40.27567 | 2.29E-04 | 0.03222335 |
| SPP1 | 4 | 87975650 | 87983426 | + | 2321 | secreted ph | 0.753617 | 4.854363 | 40.23328 | 2.30E-04 | 0.03222335 |
| HOXA11-AS | 7 | 27184507 | 27189298 | + | 3128 | HOXA11 an | -1.14883 | 2.301413 | 39.90789 | 2.36E-04 | 0.03237403 |
| KDR | 4 | 55078259 | 55125595 | - | 6563 | kinase inser | -0.76575 | 7.926168 | 39.81237 | 2.38E-04 | 0.03237403 |
| BRWD1 | 21 | 39184176 | 39321559 | - | 19794 | bromodom | -0.612883 | 5.070121 | 39.78421 | 2.39E-04 | 0.03237403 |
| RWDD2B | 21 | 29004384 | 29019360 | - | 3377 | RWD doma | -2.714312 | 2.246422 | 46.67437 | 2.40E-04 | 0.03237403 |
| FAP | 2 | 1.62E+08 | 1.62E+08 | - | 6968 | fibroblast a | -1.290492 | 1.928665 | 39.60644 | 2.43E-04 | 0.03237403 |
| KIF5A | 12 | 57550044 | 57586633 | + | 6216 | kinesin fam | 1.718364 | 2.094969 | 39.13871 | 2.53E-04 | 0.03324882 |
| PARVB | 22 | 43999211 | 44172949 | + | 7403 | parvin beta | -0.662715 | 5.501614 | 38.96673 | 2.56E-04 | 0.03324882 |
| MIS18A | 21 | 32268228 | 32279049 | - | 1694 | MIS18 kinet | -0.767492 | 3.735808 | 38.89002 | 2.58E-04 | 0.03324882 |
| CBR1 | 21 | 36069941 | 36073166 | + | 2450 | carbonyl re | -0.97619 | 4.754736 | 39.02842 | 2.60E-04 | 0.03324882 |
| PIGP | 21 | 37059170 | 37073170 | - | 7923 | phosphatid | -0.741148 | 3.467921 | 38.74363 | 2.61E-04 | 0.03324882 |
| IL17RD | 3 | 57089982 | 57170306 | - | 9561 | interleukin | 1.029122 | 2.702001 | 38.65591 | 2.63E-04 | 0.03324882 |
| CACNA1H | 16 | 1153106 | 1221771 | + | 8620 | calcium volt | 0.481259 | 5.444045 | 38.15046 | 2.75E-04 | 0.03444226 |
| HOPX | 4 | 56647988 | 56681899 | - | 7753 | HOP homeo | -0.911016 | 4.523441 | 38.00563 | 2.79E-04 | 0.03457798 |
| AC012531.1 | 12 | 54019910 | 54022589 | + | 2680 | novel trans | 1.11497 | 2.24408 | 37.86392 | 2.82E-04 | 0.03470875 |
| PTPRS | 19 | 5158495 | 5340803 | - | 8565 | protein tyro | 0.585845 | 6.753573 | 37.76466 | 2.85E-04 | 0.03471316 |
| CALCOCO1 | 12 | 53708517 | 53727745 | - | 8898 | calcium bin | 0.49089 | 6.262292 | 37.30277 | 2.97E-04 | 0.03569057 |
| SLC12A8 | 3 | 1.25E+08 | 1.25E+08 | - | 5586 | solute carri | 0.708215 | 5.595601 | 37.11445 | 3.02E-04 | 0.03569057 |
| ENTPD6 | 20 | 25195693 | 25226729 | + | 5327 | ectonucleo | -0.61909 | 5.872011 | 37.0857 | 3.02E-04 | 0.03569057 |
| GNG11 | 7 | 93921699 | 93928610 | + | 3055 | G protein su | -0.653248 | 6.729093 | 37.07405 | 3.03E-04 | 0.03569057 |
| RAC2 | 22 | 37225270 | 37244448 | - | 2523 | Rac family s | -0.655454 | 5.648819 | 36.75692 | 3.11E-04 | 0.03642412 |
| KCNMA1 | 10 | 76869601 | 77638369 | - | 35644 | potassium c | 0.854815 | 4.706294 | 36.50105 | 3.19E-04 | 0.03697582 |
| MRPS6 | 21 | 34073224 | 34143034 | + | 4273 | mitochondr | -0.747616 | 4.852105 | 36.16937 | 3.29E-04 | 0.03780574 |
| CEMIP | 15 | 80779343 | 80951776 | + | 7621 | cell migrati | -1.110952 | 1.873064 | 36.06449 | 3.32E-04 | 0.03786447 |
| MGAT5 | 2 | 1.34E+08 | 1.34E+08 | + | 8765 | alpha-1,6-m | -0.715812 | 7.493409 | 35.80008 | 3.40E-04 | 0.03796186 |
| ACSS1 | 20 | 25006230 | 25058980 | - | 8691 | acyl-CoA sy | -1.60556 | 1.200208 | 35.79933 | 3.40E-04 | 0.03796186 |
| PDXK | 21 | 43719094 | 43762307 | + | 11912 | pyridoxal ki | -0.571581 | 7.659764 | 35.73625 | 3.42E-04 | 0.03796186 |

| | | | | | | | | | | | |
|---|---|---|---|---|---|---|---|---|---|---|---|
| TPST2 | 22 | 26521996 | 26596717 | - | 6913 | tyrosylprot | -0.786321 | 5.361605 | 35.69354 | 3.43E-04 | 0.03796186 |
| MEIS3 | 19 | 47403124 | 47419523 | - | 5122 | Meis home | 0.900327 | 5.225387 | 36.53291 | 3.49E-04 | 0.03830172 |
| FXYD6 | 11 | 1.18E+08 | 1.18E+08 | - | 4986 | FXYD doma | 0.898717 | 4.82622 | 35.05548 | 3.67E-04 | 0.03987091 |
| SLC7A7 | 14 | 22773222 | 22829820 | - | 5131 | solute carri | -0.786271 | 3.664348 | 34.9232 | 3.69E-04 | 0.03987091 |
| SLITRK5 | 13 | 87672615 | 87696272 | + | 21103 | SLIT and NT | 1.866262 | 3.420782 | 41.13688 | 3.82E-04 | 0.03992422 |
| EGFL6 | X | 13569601 | 13633575 | + | 2611 | EGF like do | -3.915336 | 2.196298 | 43.85127 | 3.82E-04 | 0.03992422 |
| PAXBP1 | 21 | 32733899 | 32771792 | - | 7146 | PAX3 and P | -0.675741 | 4.848029 | 34.54865 | 3.83E-04 | 0.03992422 |
| EFNA2 | 19 | 1285873 | 1301431 | + | 2424 | ephrin A2 [ | 0.91467 | 2.5893 | 34.45169 | 3.86E-04 | 0.03992422 |
| KCNH2 | 7 | 1.51E+08 | 1.51E+08 | - | 5877 | potassium | 0.773255 | 4.001641 | 34.40936 | 3.88E-04 | 0.03992422 |
| B3GALT5 | 21 | 39556442 | 39673137 | + | 14319 | beta-1,3-ga | -1.186928 | 1.699041 | 34.3766 | 3.89E-04 | 0.03992422 |
| UBE2G2 | 21 | 44768580 | 44801826 | - | 7195 | ubiquitin c | -0.47988 | 6.421583 | 34.34332 | 3.90E-04 | 0.03992422 |
| SCUBE3 | 6 | 35213956 | 35253079 | + | 7819 | signal pept | 0.602905 | 5.272822 | 34.19663 | 3.96E-04 | 0.03992422 |
| ATP5PO | 21 | 33903453 | 33915814 | - | 3170 | ATP syntha | -0.650358 | 5.3567 | 34.15871 | 3.97E-04 | 0.03992422 |
| ZNF185 | X | 1.53E+08 | 1.53E+08 | + | 4812 | zinc finger | 0.525134 | 4.952141 | 34.06204 | 4.01E-04 | 0.03992422 |
| ETHE1 | 19 | 43506719 | 43527230 | - | 1590 | ETHE1, pers | -0.557167 | 4.824786 | 34.0076 | 4.03E-04 | 0.03992422 |
| B3GALT5-A | 21 | 39597147 | 39612821 | - | 2516 | B3GALT5 a | -1.461112 | 0.370577 | 33.92502 | 4.06E-04 | 0.03992422 |
| PTTG1IP | 21 | 44849585 | 44873903 | - | 3537 | PTTG1 inte | -0.542603 | 9.184563 | 33.92141 | 4.06E-04 | 0.03992422 |
| TUBB2B | 6 | 3224277 | 3231730 | - | 2096 | tubulin bet | 0.878615 | 5.518534 | 34.89976 | 4.13E-04 | 0.04033206 |
| HOXB4 | 17 | 48575507 | 48578350 | - | 2002 | homeobox | 0.626649 | 3.974124 | 33.45569 | 4.25E-04 | 0.04094476 |
| LRRN2 | 1 | 2.05E+08 | 2.05E+08 | - | 5686 | leucine rich | 0.896879 | 2.841328 | 33.42616 | 4.26E-04 | 0.04094476 |
| PPL | 16 | 4882507 | 4960741 | - | 7699 | periplakin [ | 1.331979 | 1.999673 | 33.33244 | 4.30E-04 | 0.04094476 |
| EDARADD | 1 | 2.36E+08 | 2.37E+08 | + | 3985 | EDAR asso | -2.066778 | 0.63462 | 33.31359 | 4.31E-04 | 0.04094476 |
| SSC4D | 7 | 76389334 | 76409697 | - | 2821 | scavenger | 1.184188 | 2.311663 | 33.14341 | 4.38E-04 | 0.04134359 |
| CHN1 | 2 | 1.75E+08 | 1.75E+08 | - | 8898 | chimerin 1 | 0.670858 | 3.335326 | 33.0527 | 4.42E-04 | 0.04134359 |
| FOXO6 | 1 | 41361922 | 41383590 | + | 2398 | forkhead b | 1.46732 | 1.219466 | 33.01368 | 4.44E-04 | 0.04134359 |
| NPNT | 4 | 1.06E+08 | 1.06E+08 | + | 6117 | nephronect | 2.898355 | 4.269156 | 48.64342 | 4.67E-04 | 0.0428562 |
| LRRN1 | 3 | 3799437 | 3847703 | + | 4264 | leucine rich | 1.666778 | 2.491408 | 34.52334 | 4.69E-04 | 0.0428562 |
| NSG1 | 4 | 4348140 | 4419058 | + | 4355 | neuronal v | -0.978232 | 2.695139 | 32.45812 | 4.69E-04 | 0.0428562 |
| PRR29 | 17 | 63998351 | 64004304 | + | 4142 | proline rich | -0.881229 | 2.163442 | 32.30779 | 4.77E-04 | 0.04322654 |
| RASGRF2 | 5 | 80960363 | 81230162 | + | 11032 | Ras protein | -0.764511 | 3.801764 | 32.1735 | 4.83E-04 | 0.04322654 |
| MGAT4C | 12 | 85955666 | 86838904 | - | 27763 | MGAT4 fam | 2.50969 | -0.22337 | 32.16923 | 4.83E-04 | 0.04322654 |
| NDUFV3 | 21 | 42879644 | 42913304 | + | 6342 | NADH:ubiq | -0.451908 | 6.145823 | 32.06154 | 4.89E-04 | 0.04322654 |
| FAM19A5 | 22 | 48489460 | 48850912 | + | 3173 | family with | -1.640042 | 1.42834 | 32.05889 | 4.89E-04 | 0.04322654 |
| HOXA6 | 7 | 27145396 | 27150603 | - | 1285 | homeobox | 0.869738 | 2.319718 | 31.9853 | 4.92E-04 | 0.04328075 |
| TIMP3 | 22 | 32801701 | 32863043 | + | 4603 | TIMP meta | -1.197518 | 8.026477 | 38.77168 | 4.98E-04 | 0.04352929 |
| C2CD2 | 21 | 41885112 | 41954018 | - | 8730 | C2 calcium | -0.882159 | 4.241266 | 31.65407 | 5.09E-04 | 0.04422197 |
| COL13A1 | 10 | 69801867 | 69964275 | + | 5888 | collagen ty | -0.680709 | 4.534927 | 31.5805 | 5.13E-04 | 0.04428583 |
| ZBTB21 | 21 | 41986831 | 42010387 | - | 8062 | zinc finger | -0.781573 | 3.958925 | 31.50168 | 5.17E-04 | 0.04437614 |
| SAMD11 | 1 | 923928 | 944581 | + | 4173 | sterile alph | 1.18176 | 3.849364 | 33.35434 | 5.23E-04 | 0.04455697 |
| CLEC11A | 19 | 50723364 | 50725718 | + | 1434 | C-type lect | -0.747769 | 7.652576 | 31.42376 | 5.31E-04 | 0.04455697 |
| CPT1A | 11 | 68754620 | 68844410 | - | 6571 | carnitine p | -0.593864 | 3.667329 | 31.25289 | 5.31E-04 | 0.04455697 |
| NPR1 | 1 | 1.54E+08 | 1.54E+08 | + | 4952 | natriuretic | -1.343115 | 2.850903 | 31.82696 | 5.32E-04 | 0.04455697 |
| SLC27A3 | HSCHR1_1_ | 1.54E+08 | 1.54E+08 | + | 5470 | solute carri | -0.76354 | 3.707778 | 31.13075 | 5.38E-04 | 0.04475787 |

| | | | | | | | | | | | |
|---|---|---|---|---|---|---|---|---|---|---|---|
| DNMT3A | 2 | 25227855 | 25342590 | - | 12020 | DNA methy | 0.44788 | 6.520561 | 31.00012 | 5.45E-04 | 0.04510628 |
| SCAF4 | 21 | 31671033 | 31732075 | - | 5901 | SR-related ( | -0.483941 | 5.587616 | 30.64421 | 5.66E-04 | 0.04655051 |
| ADGRE2 | 19 | 14732392 | 14778560 | - | 8642 | adhesion G | -0.98577 | 1.893516 | 30.32268 | 5.85E-04 | 0.04788424 |
| GSE1 | 16 | 85169525 | 85676204 | + | 13376 | Gse1 coiled | 0.674459 | 5.843361 | 30.23881 | 5.91E-04 | 0.04803602 |
| SAMSN1 | 21 | 14485228 | 14658821 | - | 5185 | SAM domai | -1.349256 | 0.892413 | 29.99428 | 6.06E-04 | 0.04902696 |
| RRP1 | 21 | 43789513 | 43805293 | + | 4251 | ribosomal F | -0.525435 | 5.6233 | 29.72356 | 6.24E-04 | 0.04999913 |
| HOXC10 | 12 | 53985065 | 53990279 | + | 2371 | homeobox | 1.264712 | 1.514275 | 29.68927 | 6.27E-04 | 0.04999913 |
| PCDHGB6 | 5 | 1.41E+08 | 1.42E+08 | + | 6782 | protocadhe | 0.821628 | 4.543208 | 29.65442 | 6.29E-04 | 0.04999913 |
